# Short tandem repeats bind transcription factors to tune eukaryotic gene expression

**DOI:** 10.1101/2022.05.24.493321

**Authors:** Connor A. Horton, Amr M. Alexandari, Michael G. B. Hayes, Emil Marklund, Julia M. Schaepe, Arjun K. Aditham, Nilay Shah, Avanti Shrikumar, Ariel Afek, William J. Greenleaf, Raluca Gordân, Julia Zeitlinger, Anshul Kundaje, Polly M. Fordyce

**Affiliations:** Department of Genetics, Stanford University, Stanford, CA 94305; Department of Computer Science, Stanford University, Stanford, CA 94305; Department of Bioengineering, Stanford University, Stanford, CA 94305; Stowers Institute for Medical Research, Kansas City, MO, USA; Stanford Biophysics Program, Stanford University, Stanford, CA 94305; Center for Genomic and Computational Biology, Duke University School of Medicine, Durham, NC, USA; Department of Biostatistics and Bioinformatics, Duke University School of Medicine, Durham, NC, USA; Department of Computer Science, Duke University, Durham, NC, USA; Department of Molecular Genetics and Microbiology, Duke University School of Medicine, Durham, NC, USA; The University of Kansas Medical Center, Kansas City, KS, USA; ChEM-H Institute, Stanford University, Stanford, CA 94305; Chan Zuckerberg Biohub, San Francisco, CA 94110

## Abstract

Short tandem repeats (STRs) are enriched in eukaryotic *cis*-regulatory elements and their polymorphisms alter gene expression, yet how they regulate transcription remains unknown. We find that STRs can modulate transcription factor (TF)-DNA affinities and on rates by up to 70-fold by directly binding TF DNA-binding domains, with energetic impacts approaching or exceeding mutations to consensus sites. STRs maximize the number of weakly preferred microstates near target sites, thereby increasing TF density near motifs to speed target search. Confirming that STRs also impact TF binding in cells, neural networks trained only on *in vivo* occupancies predict identical effects to those observed *in vitro*. Approximately 90% of TFs preferentially bind STRs that need not resemble known motifs, providing a novel *cis*-regulatory mechanism to target TFs to cognate sites.

## Introduction

Activation and repression of eukaryotic transcription depends on sequence-specific interactions between transcription factor proteins (TFs) and DNA *cis*-regulatory elements (CREs). Although *in vitro* and *in vivo* studies of TFs over decades have identified preferred DNA sequence ‘motifs’ (*1*–*8*), these motifs alone are not sufficient to quantitatively predict genomic TF occupancies (*9*–*13*). Chromatin immunoprecipitation (ChIP) assays reveal many TFs bound at loci without motifs as well as an absence of TFs at motifs within accessible genomic loci (*14*, *15*). Additionally, TF paralogs with near-identical motif preferences bind and regulate distinct target genes (*16*–*22*). Finally, it remains extremely challenging to accurately design synthetic CREs with a specified output amplitude or kinetics (*23*–*25*). As a potential explanation for why motif-based models are insufficient, *in vitro* measurements of TF binding to motifs embedded within varying DNA sequences establish that sequence context can have dramatic impacts on binding that is not predicted by motif-centric models (*26*–*28*).

Short tandem repeats (STRs, consisting of 1-6 bp units repeated consecutively) comprise approximately 5% of the human genome (**Fig. S1-3**)—compared to just 1.5% for protein-coding genes (*29*, *30*)—and are enriched in CREs across eukaryotic genomes (*31*), including in humans (^~^¼ of enhancers contain an STR (*32*, *33*); **Fig. S4**). STRs can activate or repress transcription in *H. sapiens* (*33–46), M. musculus (*47*, *48*), S. cerevisiae* (*31*), *D. melanogaster* (*49*, *50*), and others (*51*). Dinucleotide STRs are associated with broad activity of CREs across cell types in *D. melanogaster* (*52*), and sequence variation in STRs has been proposed to account for ‘missing heritability’ in GWAS studies (*33*, *53*). Finally, population-level genomic studies have linked non-coding STR polymorphisms to autism (*54*, *55*), schizophrenia (*56*), height (*56*), and Crohn’s disease (*33*).

Despite the widespread prevalence of STRs in CREs and their documented effects on gene expression, the physical mechanism by which they alter transcription remains unclear. STRs have been proposed to modulate transcription by directly altering the intrinsic affinity of histone proteins for DNA, thereby changing nucleosome occupancy (*31*, *47*, *49*, *57*). However, STRs have not been directly shown to alter chromatin accessibility besides the unique example of nucleosome-disfavoring poly-A tracts (*58*). Alternatively, polymorphisms in STR length could alter distances between multiple motifs or between motifs and core promoter elements, thereby disrupting regulatory grammar (*59*–*61*). However, this hypothesis is at odds with genome-wide studies suggesting that the syntax of cooperative TF interactions at enhancers is unlikely to be perturbed by small changes in motif spacing (*10*, *62*, *63*). As a final hypothesis, theoretical work has suggested “sequence symmetries” (*i.e*. repetitiveness) alone contribute to non-specific TF binding, with maximum effects for homopolymer sequences (*64*, *65*). *In vitro* binding measurements and bioinformatic analyses have corroborated these theoretical predictions to suggest that STRs impact TF-DNA binding in the absence of specific base-pair recognition (*28*, *64*, *66*–*69*). Nevertheless, the interplay between specific and non-specific binding and the relative magnitudes of their thermodynamic effects remain unexplored.

Here, we use multiple high-throughput microfluidic binding assays (MITOMI (*70*, *71*), *k*-MITOMI (*72*), and STAMMP (*73*)) to systematically interrogate how STRs influence equilibrium binding and kinetics for two different basic helix-loop-helix TFs. Measured binding constants (K_d_s) for 595 distinct TF-DNA combinations establish that STRs are directly bound by TF DNA-binding domains (DBDs) and can alter binding affinities by >70-fold, approaching or exceeding effects associated with mutating the consensus binding motif. Observed effects differed for the basic helix-loop-helix TFs Pho4 (from *S. cerevisiae*) and MAX (from *H. sapiens*), which share a consensus motif, demonstrating that motif information is insufficient to predict repeat preferences. Measured dissociation rates (*k*_off_s) and calculated association rates (*k*_on_s) for 118 TF/DNA combinations establish that STRs surrounding consensus motifs primarily alter macroscopic on rates, and kinetic models and stochastic simulations based on these measurements establish that STRs increase the local density of TFs near motifs to speed target search. Neural networks trained only on *in vivo* genome-wide ChIP data predict identical effects to those measured *in vitro* across a wide variety of sequences, suggesting that STR preferences play a substantial role in correctly localizing TFs in cells. By analyzing existing protein binding microarray (PBM) data, we find that preferential binding to STRs is surprisingly widespread, that models accounting for these weak preferences significantly improve occupancy predictions, and that differential STR preferences could target TF paralogs to distinct regulatory regions. As STRs are highly mutable, we suggest that STRs should be considered an easily evolvable and novel class of *cis-*regulatory elements that tune gene expression.

## Results

### Quantitative measurements establish that STRs alter TF binding affinities

The basic helix-loop-helix TFs Pho4 (a *S. cerevisiae* TF involved in phosphate starvation response (*74*, *75*)) and MAX (a human TF involved in cell proliferation, differentiation, and apoptosis (*76*, *77*)) each bind an E-box regulatory element (**Fig. 1A**). To test the impact of STRs on binding, we quantified binding of each TF to 17 DNA sequences containing either an extended consensus E-box motif (GTCACGTGAC) or a random sequence (‘no motif’) flanked by 13 bp of either random sequence or STRs previously shown to enhance binding (*28*) (‘Library 1’; **Fig. 1B**, **Table S1**) via MITOMI microfluidic binding assays (**Figs. 1C, S5**). In these assays, valved microfluidic devices containing 1,568 reaction chambers are aligned to glass slides printed with arrays of fluorescently labeled doublestranded DNA (**Fig. S6**). Following alignment, C-terminally eGFP-tagged TFs are specifically recruited to antibody-patterned surfaces and buffer is introduced, solubilizing printed spots and allowing DNA to interact with and bind to surface-immobilized TFs (**Fig. S7**). Measured binding for each DNA sequence over multiple concentrations can be combined with calibration curves (**Figs. S8-9**) to allow quantification of concentration-dependent TF binding and global fitting of Langmuir isotherms to extract *K*_d_s (**Fig. 1C;** see Supplementary Methods).

**Fig. 1.**
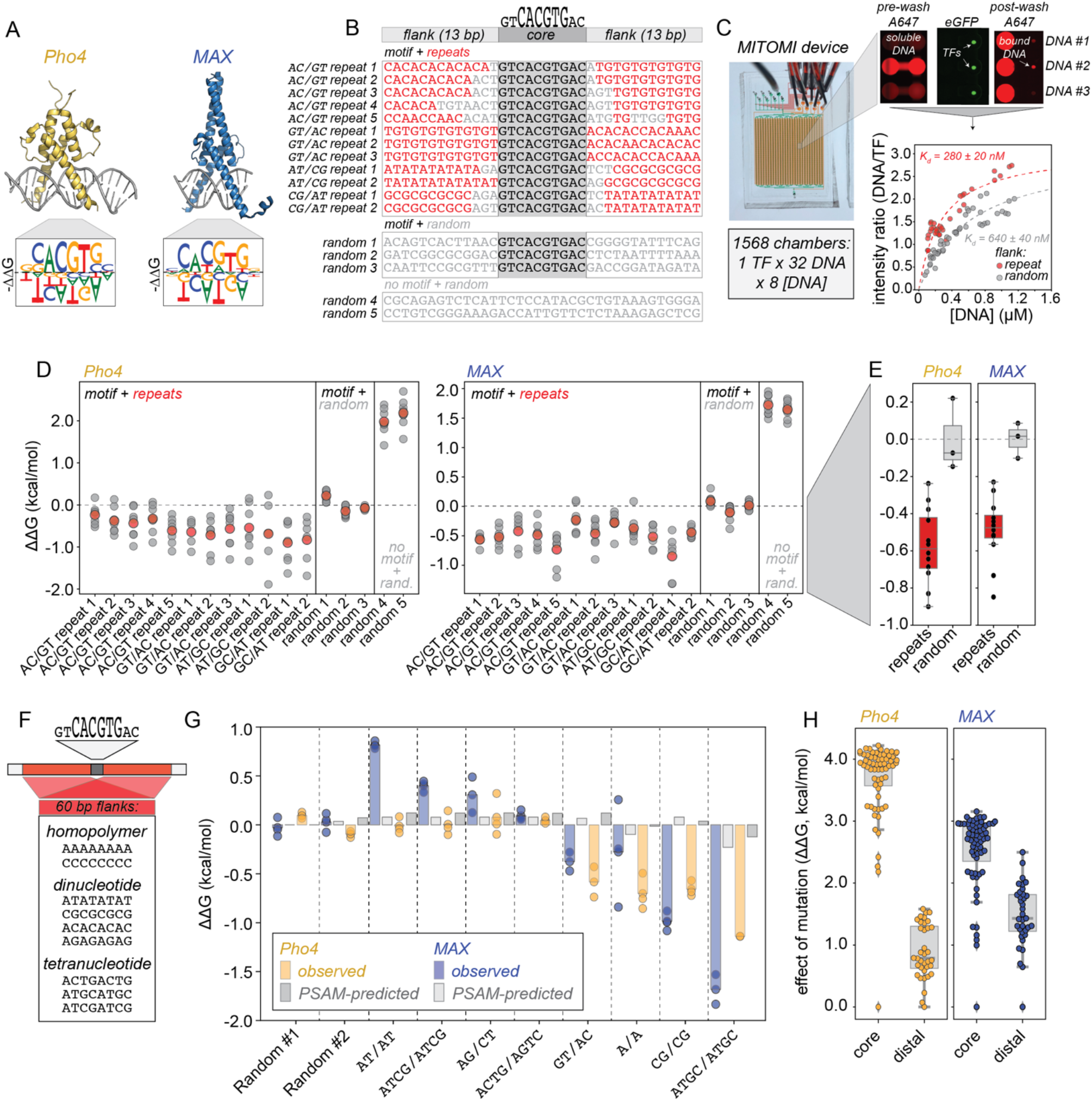
Repetitive flanking sequences alter TF-DNA binding affinities in a sequence-specific manner. **(A)** Crystal structures and position-specific affinity matrices (PSAMs) (*41*) for Pho4 (gold; PDB ID: 1a0a) and MAX (blue; PDB ID: 1hlo). **(B)** Library #1: 17 DNA sequences with either an extended (10 bp) E-box motif (dark gray) or random (light gray) sequence surrounded by 13 bp on either side of repetitive (red) or random (light gray) sequence. **(C)** MITOMI microfluidic device (left) and zoomed-in view of 3 chambers (top right) showing solubilized DNA during incubation (‘pre-wash A647’), immobilized TFs (‘eGFP’), and TF-bound DNA after washing (‘post-wash A647’). Bottom right shows representative concentration-dependent binding for DNA sequences containing an extended E-box surrounded by either repetitive (red) or random (gray) flanks. **(D)** Measured ΔΔG values across all Library #1 sequences for Pho4 (left) and MAX (right); ΔΔGs are calculated relative to the overall median value for oligonucleotides bearing an E-box consensus surrounded by random flanking sequence. Light gray dots show all measurements; darker circles indicate median values per oligo. **(E)** Median values (black markers and box plots) for all sequences containing either repetitive (red) or random (gray) flanking sequences for Pho4 (left) and MAX (right). **(F)** Library #2: 10 DNA sequences containing a central extended (10 bp) E-box motif surrounded by 60 bp on either side of listed homopolymeric, dinucleotide, or tetranucleotide repeats. **(G)**. Measured ΔΔG values across all Library #2 sequences for Pho4 (gold) and MAX (blue); ΔΔGs are again calculated relative to the overall median value for oligonucleotides bearing an E-box consensus surrounded by random flanking sequence. Gray bars indicate magnitude of effects predicted by PSAMs. **(H)** Observed effects on ΔΔG for mutating single nucleotides within the CACGTG core E-box (‘core’) (*70*) *vs*. altering flanking sequence within Library 2 (‘distal’) for Pho4 (left, gold) and MAX (right, blue) overlaid on boxplots (gray).

Measured Library 1 ΔΔGs spanned ^~^2.6 and 3.1 kcal/mol with a mean RMSE between replicates of ^~^0.53 and 0.31 kcal/mol for Pho4 and MAX, respectively (**Figs. 1D, S10-18**). DNA sequences with a motif surrounded by STRs were consistently bound 0.23-0.90 kcal/mol tighter than those with a motif surrounded by random sequences, corresponding to a ^~^1.5-4.6-fold change in predicted affinity (**Figs. 1D-E**), and the magnitude of these effects scaled with the length of repetitive sequence (**Fig. S19**). Measured ΔΔGs did not change with ^~^5-fold differences in protein concentration, confirming that DNA was in vast excess of available protein (**Figs. S20-21**). Measured ΔΔGs were also consistent when using either wheat germ extract or TBS as binding buffer (**Fig. S22**), and negative control experiments assessing binding to eGFP alone showed no variability above the background RMSE (maximum deviation of ±0.5 kcal/mol; **Fig. S23**). Linear mononucleotide (*e.g*. position-specific affinity matrix, PSAM) specificity models predict a <0.1 kcal/mol effect for all flanking sequences but 1 (‘Motif + GT/AC repeat 2’) (**Fig. S24**), establishing that measured effects are not due to cryptic consensus sites distal to the core motif.

### The magnitude of STR effects on affinity depend on STR nucleotide sequence

To test how the nucleotide sequence of STRs alters binding, we designed a DNA library containing either an extended consensus E-box motif or random sequence surrounded on each side by 60-nucleotide flanks comprised of homopolymer, dinucleotide, or tetranucleotide STRs or random sequence (‘Library 2’; **Fig. 1F, Table S2;** CG/AT indicates a CG repeat on one side of the motif and an AT repeat on the other). As extension of repetitive sequences can be technically challenging, we visualized extension via denaturing gel electrophoresis and then quantified binding affinities only for those sequences that extended successfully (**Figs. 1G, S25-32**). Observed effects ranged from increasing affinity by 1.7 kcal/mol (18-fold affinity increase) to reducing affinity by 0.8 kcal/mol (4-fold affinity decrease); while ATGC STRs enhanced binding for both Pho4 and MAX, other STRs (AT/AT, ATCG/ATCG, and AG/CT) were deleterious for MAX only (**Fig. 1G**). As for Library 1, results did not change with surface protein density (**Figs. S33-34**), no sequence-specific binding was detected for an eGFP-only negative control (**Fig. S35**), and observed effects were inconsistent with PSAM-based models of specificity (**Fig. 1G**). Observed effects diverged significantly for Pho4 and MAX (**Fig. 1G**), signifying that ‘consensus’ binding motifs are not sufficient to predict STR preferences. The energetic contributions of these flanking sequences approach or exceed those associated with mutating core consensus residues (*70*), particularly for MAX, suggesting that STRs could play a significant role in proper TF localization *in vivo* (**Fig. 1H**).

### STRs alter affinities by directly recruiting TFs

These observed STR effects suggest two possible mechanistic models (**Fig. 2A**). STRs could enhance TF binding to the core consensus site, perhaps by altering local DNA ‘shape’ (*78*–*82*) (**Fig. 2A**, top). This model predicts that STRs should only alter binding in the presence of a core motif and that TF/DNA stoichiometry should not depend on flanking sequence. Alternatively, STRs could themselves represent additional binding sites (**Fig. 2A**, bottom). This second model predicts that STRs should enhance binding whether or not they flank a consensus motif and that multiple TFs will bind oligonucleotides containing STRs.

**Fig. 2:**
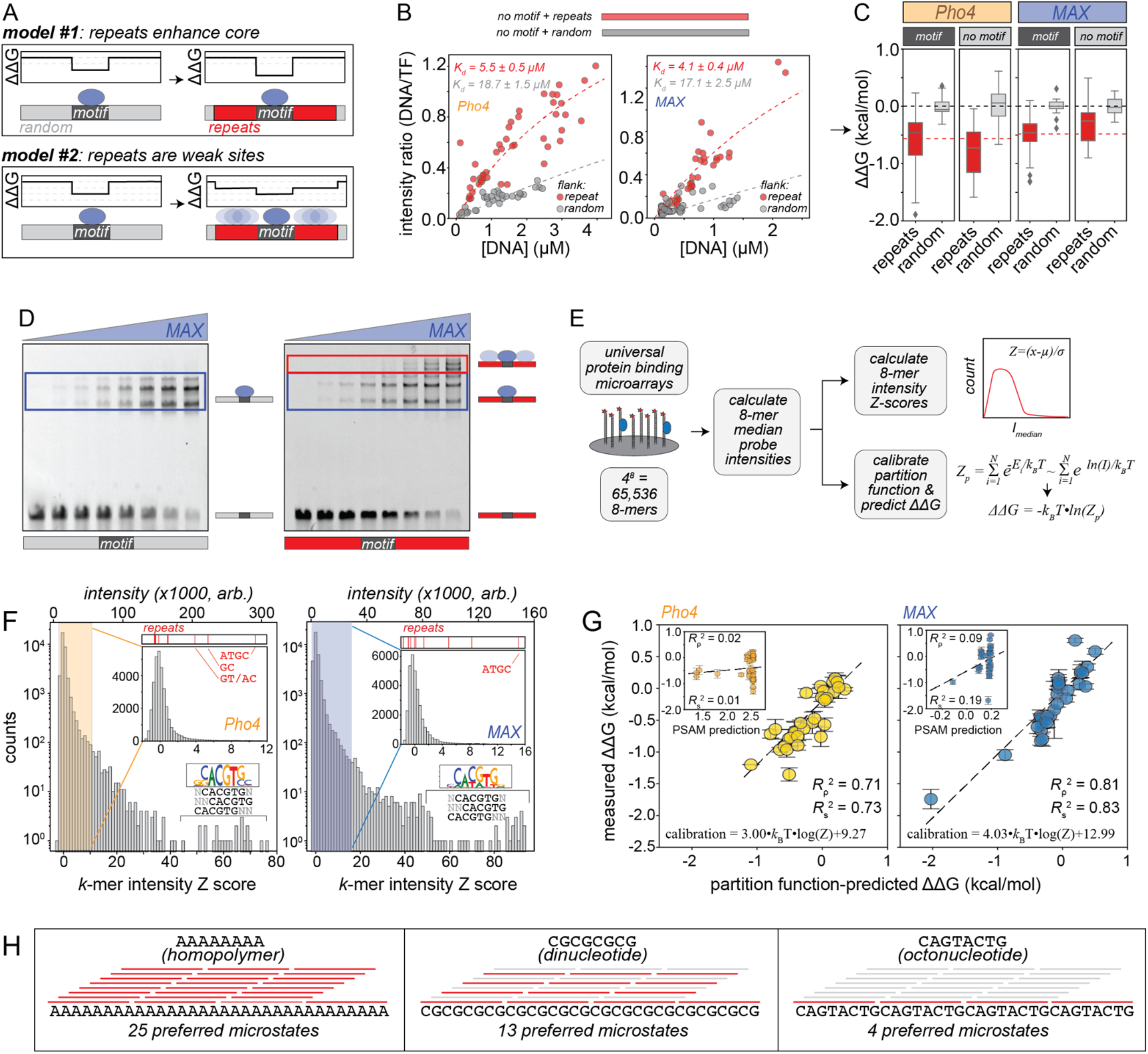
STRs are directly bound by transcription factors with observed affinities that can be accurately predicted via statistical mechanics. **(A)** Cartoon schematic of models explaining how repetitive flanking sequence could enhance TF binding affinities. **(B)** Representative concentration-dependent binding for Pho4 (left) and MAX (right) interacting with DNA sequences containing either repetitive (red) or random (gray) sequences in the absence of an E-box motif. **(C)** Box plots of relative affinities (ΔΔGs) for Pho4 and MAX binding to oligonucleotides with repetitive (red) or random (gray) sequence flanking an extended E-box consensus (dark gray) or random sequence (light gray); black and red dashed lines indicate median overall affinities. **(D)** Electromobility shift assays (EMSAs) for increasing concentrations of eGFP-tagged MAX interacting with Alexa647-labeled dsDNA duplexes containing a central extended E-box surrounded by random (left) or repetitive (right) sequences. Blue boxes highlight TF complexes bound to the core motif; red boxes highlight supershifted species with additional bound TFs. Native gel electrophoresis reveals MAX alone runs as 3 bands, likely representing MAX homodimers, MAX monomers, and eGFP-only truncation constructs (**Fig. S39**). **(E)** Pipeline for calculating 8-mer intensity *Z*-scores from universal PBM data and calibrating partition function scores to predict binding (see Methods). **(F)** Log-linear histograms of affinity *Z*-scores for all 8-mers for Pho4 (left) and MAX (right); inset linear-linear plots highlight background binding distributions and Z scores of STRs measured in this study (red bars, top). **(G)** Scatter plots, linear regressions, and correlation coefficients for measured ΔΔGs *vs*. calibrated partition function-predicted scores across all measured repeats for Pho4 (left) and MAX (right). (**H**) Schematic showing possible microstates as a function of sequence.

Contradicting a model in which STRs alter motif-proximal ‘shape’, concentration-dependent binding for both Pho4 and MAX was clearly stronger for sequences containing favorable STRs even in the absence of a motif (**Figs. 2B-C**, data from Library 2). Moreover, energetic effects of STRs did not correlate significantly with any predicted shape parameters (*i.e*. minor groove width, helical twist, propeller twist, roll, and electrostatic potential) (**Figs. S36-37**), and circular dichroism (CD) spectroscopy ruled out enhanced binding resulting from STR-dependent structural transitions between B- and Z-form DNA (**Fig. S38**). Finally, electrophoretic mobility shift assays (EMSAs) using Alexa-647-labeled dsDNA and increasing concentrations of eGFP-tagged MAX TFs (**Figs. 2D, S39**) revealed supershifted bands and a secondary linear increase for DNA sequences containing STRs at higher MAX concentrations, consistent with increased TF recruitment. Together, these experiments are consistent with a model in which STR flanks enhance DNA binding via direct recruitment of TFs *in vitro*.

### Statistical mechanical models integrating data across platforms accurately predict STR effects

Universal protein-binding microarray (uPBM) experiments measure binding of fluorescently-tagged TFs to surface-immobilized DNA duplexes containing all possible 8-mer DNA sequences, providing comprehensive measurements of TF-DNA specificity in an alternate (flipped) experimental configuration (*2*, *3*, *5*, *83*). To test if previously published uPBM measurements also reveal enhanced binding of Pho4 and MAX to specific STRs, we calculated the median intensity for all probes containing each of the 65,538 possible DNA 8-mers and then calculated a Z-score for each 8-mer relative to this distribution (**Figs. 2E,F**). As expected, probes containing 8-mer variants of the known E-box CACGTG consensus were bound very strongly by Pho4 and MAX, with intensity distribution Z-scores of 40-80 (**Fig. 2F**). Consistent with MITOMI results, favorable repeats were bound statistically significantly above background for both MAX (ATGC, Z=15.1, p=4·10^-127^; CG, Z=8.3, p=5·10^-40^; and AC, Z=5.0, p=1·10^-15^) and Pho4 (ATGC, Z=10.7, p=7·10^-72^; GC, Z=3.9, p=3·10^-11^; and AC, Z=5.4, p=9·10^-20^; **Fig. 2F;** see Supplementary Methods).

Next, we combined information from PBM and MITOMI experiments to test if statistical mechanics models improve binding predictions by accurately accounting for effects of flanking sequence (**Figs. 2E,G**). Comparisons between MITOMI-measured ΔΔGs (**Figs. S40-43, Table S3**) and log-transformed gcPBM intensities for MAX binding to 32 probes were strongly anticorrelated (*R*_p_^2^ = 0.89) over a wide dynamic range (2.5 kcal/mol) (**Fig. S44**), confirming prior reports that PBM intensities can report on affinities (*3*, *19*, *20*, *84*, *85*). This property allowed us to compute a partition function from intensities and predict binding ΔΔGs for Pho4 and MAX binding to all oligonucleotide sequences from DNA Libraries 1 and 2 (see **Supplementary Methods**; **Fig. 2G, S45**). For DNA Library 1 (which contains intact, mutated, or ablated E-box consensus sequences surrounded by 13 bp variable flanking sequences), partition function-based predictions significantly improve agreement with measured ΔΔGs over standard PSAM predictions (*R*_p_^2^ = 0.91 *vs*. 0.66 and *R*_p_^2^ = 0.93 *vs*. 0.74 for Pho4 and MAX, respectively) (**Fig. S46**). For DNA Library 2, in which all sequences contain an E-box but differences in flanking sequences can change measured ΔΔGs by up to 1.6 and 2.5 kcal/mol for Pho4 and MAX, respectively, partition function-based calculations yielded even more dramatic improvements (*R*_p_^2^ = 0.71 *vs*. 0.09 and *R*_p_^2^ = 0.81 *vs*. 0.02 for Pho4 and MAX, respectively) (**Fig. 2G**). Returned fit parameters from these linear regressions allow calibration of partition function-based predictions in energetic space with as few as 9 thermodynamic measurements (*K_d_s* or ΔΔGs) (see **Supplementary Methods**; **Fig. S47**).

### Even weakly preferred STRs enhance binding by increasing the number of preferred microstates

Preferred repeats for Pho4 and MAX (*e.g*. CG, ATGC) do not show strong sequence similarity to the known E-box consensus, as evidenced by a failure of PSAM-based models to predict observed effects (**Figs. 1H, 2G, S24, S46**). Why, then, do repeats recruit TFs? By virtue of being repetitive, STRs create multiple identical binding sites that are equally probable binding microstates (**Fig. 2H**), and an analytical treatment establishes that STRs (in particular, homopolymers) maximize binding entropy and minimize binding energy (see **Supplementary Discussion**). To estimate the energetic magnitude of this statistical effect, we conducted Monte Carlo simulations that randomly sample from the observed energy distributions to simulate either random or homopolymeric sequences (**Fig. S48**, see **Supplementary Methods**) and find that increasing repetitiveness alone can contribute up to 0.3 kcal/mol mean binding energy via entropic effects (**Fig. S48**). However, these effects are considerably more pronounced for binding sites with affinities even only slightly stronger than background binding: dinucleotide STRs with intensity Z scores between 1 and 2 or between 5 and 10 are predicted to enhance binding by 0.6 and 1.4 kcal/mol (10-fold), respectively, for a 57-bp DNA sequence, and different STRs reveal similar results (**Figs. S48**). We note that this partition function model is purely additive, establishing that additional mechanisms of cooperativity (*e.g*. allostery, avidity, allovalency) are not necessary to explain the effects of STRs on *in vitro* binding.

### STRs are directly bound by TF DNA-binding domains

Results thus far establish that TFs directly bind STRs but do not identify which portion of the TF recognizes them. STRs may be recognized by intrinsically disordered regions outside of TF DNA-binding domains (DBDs) (*86*); alternatively, STRs may be bound by DBDs themselves. To distinguish between these possibilities, we used STAMMP (*73*) (**Fig. 3A-B**) to: (1) recombinantly express and purify 221 Pho4 variants containing systematic amino acid mutations within and surrounding the DBD (**Fig. 3C, Table S4**), and (2) quantify concentration-dependent binding for each variant interacting with DNA sequences containing a motif flanked by either random sequence or GC dinucleotide STRs previously shown to enhance binding (**Fig. 1G**). Across 9 STAMMP experiments, 214/221 variants showed strong expression (**Figs. 3B, 3C, S49-50**) and binding intensities as a function of concentration were well-fit by a two-state model across both DNA sequences (**Figs. 3D, S51-56**). This process yielded 6139 individual TF/DNA *K*_d_ measurements; after normalization between experiments, measured energetic effects were consistent across experiments (<0.48 kcal/mol RMSE) and spanned >4 kcal/mol (**Figs. 3D, S51-56**).

**Fig. 3:**
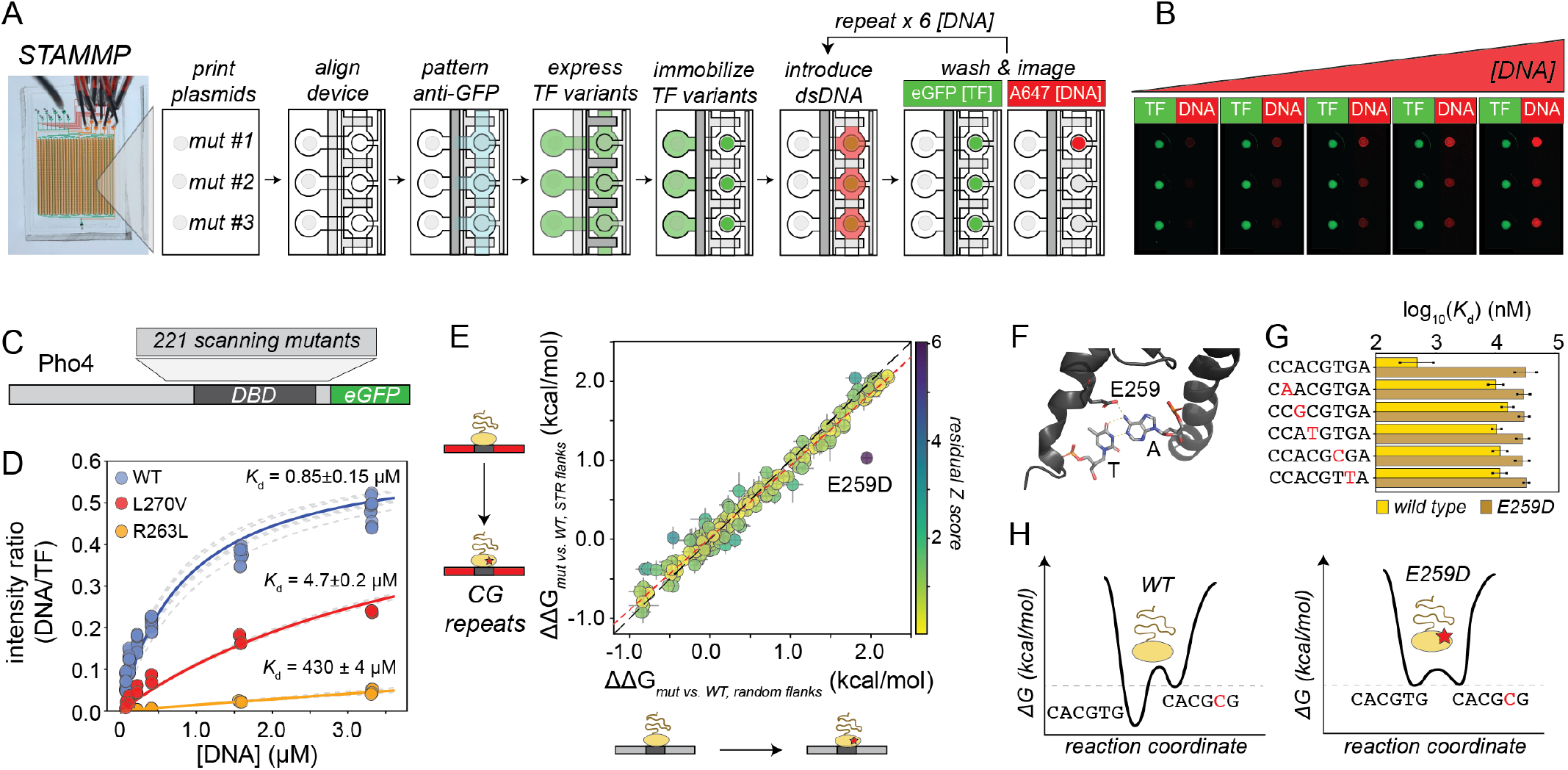
Mutations within TF DBDs alter repeat sensitivity. **(A)** Experimental pipeline for STAMMP illustrating steps for recombinant protein expression, surface-immobilization, purification, and measurement of concentration-dependent binding behavior. **(B)** Example zoomed-in fluorescence images showing immobilized TFs and concentration-dependent DNA binding. **(C)** Schematic of C-terminally eGFP-tagged Pho4 and location of scanning mutants. **(D)** Example concentration-dependent binding measurements and Langmuir isotherm fits for WT Pho4 and 2 mutants (L270V and R263L) interacting with ‘Motif + random 1’. **(E)** Effects of TF mutations on relative DNA binding affinity for an extended E-box consensus flanked by CG repeats *vs*. random sequence. Black dashed line indicates 1:1 relationship, red dashed line shows linear regression; color bar indicates Z score of residuals from linear regression. **(F)** Zoomed-in crystal structure showing contacts between the WT E259 and E-box consensus (PDB ID: 1a0a). **(G)** Affinities for Pho4 WT and E259D mutants interacting with consensus E-box and 5 single nucleotide variants. **(H)** Reaction coordinate diagram of binding specificity landscape for Pho4 WT and E259D.

We then compared measured ΔΔGs for each mutant relative to the WT TF across DNA sequences, as any residues involved in STR recognition should differentially impact affinity upon mutation (**Fig. 3E**). Nearly all mutants altered binding affinities equally across DNA sequences, but E259D showed significantly enhanced binding to a sequence with flanking sequences comprised of CG dinucleotide repeats as compared with random flanks (**Figs. 3E, S57**; residual Z score = 6.0, p = 1.7·10^-9^, ΔΔΔG ≈ 0.73 kcal/mol). In the Pho4 crystal structure, E259 directly contacts nucleotides from both strands at the CACG**T**G position (*87*) (**Fig. 3F**), and comparisons of measured affinities for WT Pho4 and E259D reveal that while WT Pho4 shows a strong preference for the canonical E-box motif (CACGTG), E259D shows equal, weak (100-fold lower) binding to the canonical E-box and CACG**C**G (*73*) (**Fig. 3G, H**). These observations are consistent with a model in which the increased promiscuity of the E259D binding energy landscape leads to an effective increase in preference for CG dinucleotide repeats relative to the WT motif (**Fig. 3H**) and establish that the DBD alone is sufficient for repeat recognition.

### STRs increase macroscopic association rates

To investigate how flanking sequences alter TF binding kinetics, we leveraged *k*-MITOMI (*72*) (**Fig. 4A**) to quantify dissociation rates for Pho4 and MAX interacting with DNA sequences containing an extended E-box motif (GTCACGTGAC) surrounded by 60 bp flanks comprised of either random sequence or 8 different STRs that extended properly (*homopolymer: A/A; dinucleotide: AT/AT*, *AG/CT*, *GT/AC; tetranucleotide: ACGT/ACGT*, *ATCG/ATCG*, *ACTG/AGTC*, *ATGC/ATGC*, where *AG/CT* indicates an AG dinucleotide repeat on one side of the motif and a CT dinucleotide repeat on the other side of the motif). To quantify dissociation, we iteratively: (1) closed valves to trap TF-bound DNA, (2) introduced a high-affinity unlabeled DNA competitor, (3) opened valves for 1-4 seconds to allow fluorescently-labeled DNA to dissociate, (4) closed valves and washed out unbound material, and then (5) imaged all device chambers (**Fig. 4A**); excess unlabeled DNA competitor was included to outcompete rebinding and ensure accurate rate measurements (see **Supplementary Methods**). Decreases in the measured Alexa-647/eGFP (DNA/TF) intensity ratio over time were well-fit by a single exponential for both Pho4 and MAX (**Figs. 4B**, **S58-59, Supplementary Data**); measured rates typically varied by <3-fold across experiments prior to normalization (**Fig. S60-61**; see **Supplementary Methods**). For both Pho4 and MAX, different 60 bp flanking STRs changed dissociation rates only slightly (<1.7-fold, less than noise between experiments) (**Figs. 4C, S60-61**). By contrast, inferred on rates (*k*_on_ = *k_off_/K_d_*, calculated assuming a two-state model in which DNA is either bound or unbound) were dramatically altered (**Figs. 4C, S62-71**): favorable STRs increased macroscopic on-rates by 7- to 54-fold for Pho4 and MAX, respectively, suggesting that observed changes in affinity are primarily due to altered TF macroscopic association rates (**Figs. 4C, S62-71**).

**Fig. 4:**
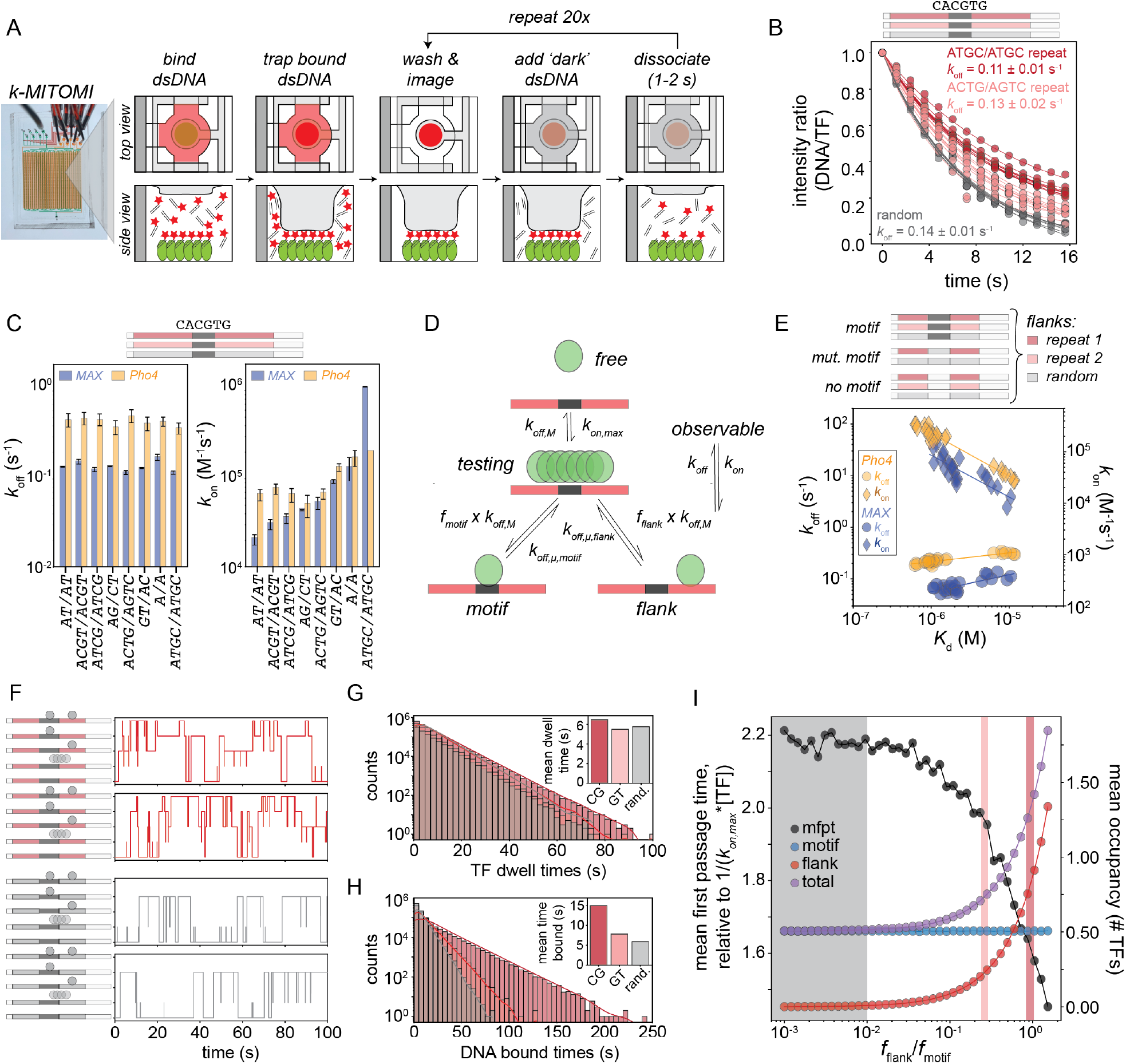
Repetitive flanking sequences increase macroscopic association rates and reduce mean first passage time. **(A)** Experimental pipeline for *k*-MITOMI (see Supplementary Methods). **(B)** Example dissociation curves for MAX interacting with DNA Library 2 sequences showing per-chamber measurements (markers), per-chamber single-exponential fits (lines), and the average of returned fit parameters (annotation) for each sequence. **(C)** Measured *k_off_* (left) and calculated *k_on_* (right) values as a function of flanking sequence for Pho4 (yellow) and MAX (blue) interacting with DNA library 2 sequences (all of which contain a core motif). **(D)** Proposed 4-state model and associated microscopic rate constants for TF binding to sequences with a central core ‘motif’ surrounded by different flanking sequences. **(E)** Average measured *k_off_* (circle markers, left axis) and calculated *k*_on_ (diamond markers, right axis) values *vs*. measured affinities (*K*_d_s) for Pho4 (yellow) and MAX (blue) interacting with all sequences from DNA Library 1. **(F)** Sample TF trajectories from Gillespie simulations modeling 2600 TFs interacting with a single DNA sequence containing a consensus motif flanked by either repetitive (top, red) or random (bottom, gray) flanks; DNA can be unbound, associated with TFs in a ‘testing’ state, bound by a TF at the motif, bound by a TF in flanking sequence, or bound by TFs at the motif and flanking sequence simultaneously. **(G)** Log-linear distribution of TF dwell times across 1000 simulations for sequences with a consensus motif flanked by CG repeats, GT repeats, or random sequence; inset shows mean dwell times by sequence. **(H)** Log-linear distribution of the fraction of time a DNA sequence is bound across 1000 simulations for sequences with a consensus motif flanked by GC repeats, GT repeats, or random sequence; inset shows mean time occupied by sequence. **(I)** Mean first passage time (black markers, left axis; units relative to fastest possible search time, 1/(k_on,max_*[TF])), mean motif occupancy (blue markers, right axis), mean flank occupancy (red markers, right axis), and mean total DNA occupancy (purple markers, right axis) as a function of the likelihood of binding flanking sequence; gray box indicates range of affinities for random flanks; pink and red boxes correspond to *f_flank_* values for GT and CG repeats, respectively.

### STRs create a pool of weakly bound TFs enriched near target motifs

STRs are enriched near binding sites of stress-response TFs in budding yeast that likely require a rapid transcriptional response (*31*), suggesting that STRs could reduce search times *in vivo*. To model how changes to motifs and flanking sequences alter search behavior, we turned to a 4-state continuous-time Markov Chain (CTMC) model in which TFs may either be: (1) *free* (nonspecifically diffusing in the nucleoplasm), (2) *testing* (nonspecifically bound to DNA), (3) bound to a *motif*, or (4) bound to the *flanks* (**Fig. 4D,** see **Supplementary Methods**). The rate constant for transitioning between the *free* and *testing* states is given by *k*_on,max_ (the theoretical upper bound for the on-rate if all non-specific TF-DNA interactions result in specific binding); rate constants for transitioning from the *motif-* or *flank-bound* state to the *testing* state are given by *k*_off,μ,motif_ and *k*_off,μ,flank_; and the probabilities of transitioning to the motif or flanks depend on the probability of binding either sequence (*f_flank_* or *f_motif_*) and the rate at which TFs transition from *testing* back to the *free* state (*k*_off,M_). Together, this yields a simple expression for the transition probability from the testing state to either the flank or motif (*p*_testing,x_ = *f*_x_/(1 + *f*_flank_ + *f*_motif_); x ∈ {flank,motif}). Assuming time spent in the *testing* state is negligible, this 4-state model can determine these microscopic rate constants from macroscopic measurements of affinities and dissociation rates for sequences containing either a consensus E-box, weak E-box, or scrambled sequence surrounded by 13 bp flanks comprised of either GT/AC or CG/AT dinucleotides or random sequence (DNA Library 1; **Fig. 4E, S72-82;** see **Supplementary Methods**). Consistent with recent work on *E. coli* LacI binding to various operator sequences (*88*), mean microscopic dissociation rates (*k_off,μ_*) for sequences with a consensus E-box or a weak E-box were similar but affinities and microscopic association probabilities differed by 12- or 16-fold (**Fig. S82**). Using these microscopic rate parameters in Gillespie stochastic simulations to predict binding trajectories for individual TFs showed that sequences with flanking STRs were frequently occupied by multiple TFs (**Fig. 4F, S83-86**). While the DNA dwell time for any individual TF was largely independent of flanking sequence identity (**Fig. 4G**), as expected with the absence of an observed macroscopic off-rate effect (**Figs. 4C, 4E, S82**), the fraction of time that a DNA sequence was occupied by TFs was dramatically longer for DNA sequences with preferred flanking STRs (**Fig. 4H**). Mean behavior across 100 simulations showed that as the relative affinity for flanking STRs increases, total DNA occupancy increases, thereby creating a locally concentrated pool of TFs available to bind the consensus (**Fig. 4I, S83-84**). Though mean first passage time and occupancy depend on TF concentration and estimated on rates, the magnitudes of simulated effects were invariant across a range of these parameters (**Figs. S85-87**). Even for this simple model that does not consider the proximity between the motif and the flanks, favorable STRs thereby reduce the mean first passage time to the total DNA site of motif and flanking sequences across 10,000 simulations (**Fig. 4I, S88**), consistent with a hypothesized role for STRs in regulating stress responses (*31*) and with previous work showing that favorable flanking sequences can act as “antennae” to enhance TF target search (*89*).

### STRs alter gene expression by tuning TF occupancies in vivo

While STRs have repeatedly been associated with changes in gene expression in cells and the length of STRs in the genome exceeds the length required for an *in vitro* effect (**Fig. S89**), results thus far do not elucidate if STRs change gene expression by altering TF occupancies *in vivo*. Directly quantifying impacts of STRs on TF binding in cells is technically challenging, as chromatin immunoprecipitation assays often struggle to detect the low-affinity and transient binding expected for STRs. Instead, we trained the BPNet (*10*) neural network (NN) on *in vivo* ChlP-seq data to predict TF binding profiles from DNA sequence with nucleotide resolution, and then applied AffinityDistillation (AD (*13*)) to predict log-transformed mean read counts (Δlog(counts)) previously shown to correlate with measured thermodynamic energies (ΔΔGs). If STRs alter gene expression *in vivo* by changing TF occupancies, we expect BPNet to learn that they impact TF binding and AD to predict sequence-dependent read count changes that mirror ΔΔGs measured *in vitro*.

After training on high-quality MAX ChIP-seq data (*1*, *90*) (**Fig. 5A**, see Supplementary Methods), BPNet accurately predicted log-transformed read counts for held-out data (*R^2^* = 0.52) (*13*) with base-pair-resolution binding profiles that reproduced those observed experimentally (**Fig. 5A**). Returned contribution weight matrices (CWMs), which identify short subsequences most predictive of TF binding, revealed E-box-like motifs (CACGTG) that sometimes included a flanking preference for CG dinucleotides, consistent with *in vitro* preferences (**Figs. 5A, 1G, 2G-H**); some CWMs also included an AP1 binding motif (TGACTCA), consistent with AP1 acting as a pioneer factor to increase chromatin accessibility for MAX (**Fig. 5A**) (*91*). AD-predicted log-transformed read counts (Δlog(counts)) for DNA Library 1 sequences containing either a consensus or mutated E-box motif flanked by either STRs or random sequences (**Fig. 5B**, Supplementary Data) were strongly correlated with measured ΔΔGs (*R*^2^ = 0.78). Strikingly, AD consistently predicted tighter binding to consensus motifs flanked by preferred repeats (**Fig. 5B**), and importance scores from DeepSHAP (*92*, *93*), which identify single base pair contributions to the observed model output, confirmed that enhanced binding was due to the flanking STRs in these synthetic sequences (**Figs. 5C,D**). Together, these analyses establish that observed *in vivo* effects of polymorphic STRs on gene expression can be explained by differential TF binding.

**Fig. 5:**
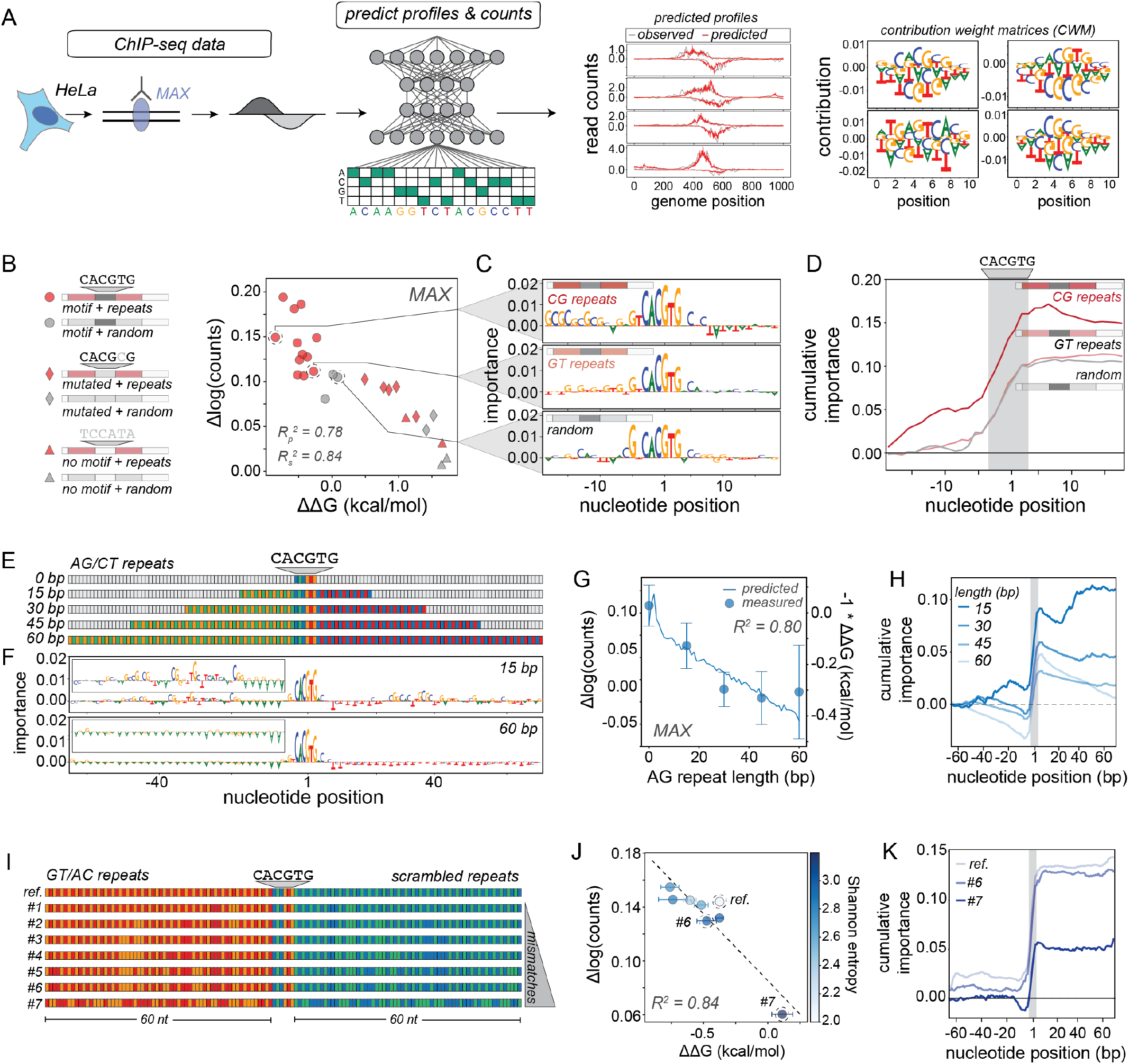
NNs trained on *in vivo* datasets correctly recapitulate repeat effects observed *in vitro* and return predictions similar to statistical mechanics models. **(A)** Experimental pipeline: AffinityDistillation (AD) neural network models trained on MAX ChIP-seq data predict bp-resolution binding profiles and return hypothetical contribution weight matrices (CWMs) representing binding preferences; positive and negative numbers represent nucleotides that favor and disfavor binding, respectively. **(B)** AD-predicted binding (Δlog(counts)) *vs*. MITOMI-measured ΔΔGs for 26 DNA sequences containing either an intact motif, a mutated motif, or scrambled sequence surrounded by either repetitive (red markers) or random (gray markers) flanking sequence. **(C)** DeepSHAP interpretations for a motif surrounded by either a favored repeat (CG, top), a disfavored repeat (GT, middle), or random sequence (bottom). The sum of importance scores across a sequence are equal to the count prediction output of the NN. **(D)** Cumulative importance scores as a function of position for a favored repeat (CG, dark red), a disfavored repeat (GT, light red), or random sequence (gray); gray box indicates motif location. **(E)** Schematic of sequences with E-box and 15, 30, 45, or 60 bp of disfavored AG/CT repeats. **(F)** DeepSHAP interpretations for 15 and 60 bp sequences from (E). **(G)** AD-predicted change in log(counts) (blue line, left axis) and −1*MITOMI measured ΔΔGs (blue markers, right axis) as a function of repeat length (relative to a sequence with a motif and random flanks). Markers and error bars show median and standard deviation across replicates. **(H)** Cumulative importance scores as a function of position for sequences with E-box and 15, 30, 45, or 60 bp of AG/CT repeats; gray box indicates motif position. **(I)** Schematic of sequences with E-box and increasingly scrambled GT/AC repeats. **(J)** AD-predicted change in log(counts) (blue line, left axis) and −1*MITOMI measured ΔΔGs (blue markers, right axis) for sequences shown in (I) (calculated relative to ‘reference’ sequence); color indicates Shannon entropy. Markers and error bars show median and standard deviation across replicates. **(K)** Cumulative importance scores as a function of position for reference sequence and sequences #6 and #7; gray box indicates motif position.

### STR impacts extend over tens of nucleotides and mismatches reduce effects

To determine the distance over which STRs impact binding, we quantified MAX binding affinities for DNA containing an E-box motif surrounded by increasing lengths (15, 30, 45, or 60 bp) of either disfavored (AG/CT) or favored (GT/AC) repeats via MITOMI (**Table S2**); in parallel, we used AD to predict MAX occupancies and binding profiles for the same sequences (**Figs. 5E-G; S90**). For disfavored AG/CT repeats, both MITOMI and AD revealed that increasing lengths of repetitive sequence monotonically reduce binding, with effects saturating after ^~^40 nucleotides (**Fig. 5G;** *R*^2^ = 0.80 between predictions and measured ΔΔGs); returned DeepSHAP interpretations and cumulative importance scores confirmed a negative contribution from STRs on either side of the motif (**Figs. 5F,H**). Favored GT/AC repeats showed more complex behavior, with short repeats (15-30 bp) increasing binding and longer repeats having only minor effects, but predictions were again consistent with experimental observations (**Fig. S90**; *R*^2^ = 0.93).

Nearly 80% of repeated units within the median human STR match the consensus repeat exactly, with the remaining 20% containing an indel or mismatched base(s) (**Supplementary Methods**). To investigate how imperfections within STRs alter binding, we applied MITOMI and AD to measure and predict MAX binding to 7 increasingly scrambled (GT/AC) repeat sequences (**Fig. 5I**). Even though the relationship between measured affinities and repeat imperfection (as quantified by Shannon entropy) was complex and non-monotonic, AD accurately predicted energetic measurements (*R*^2^ = 0.84) with predicted binding very similar to the full partition function model (*R*^2^ = 0.77; **Fig. S91**), suggesting that the algorithm had learned that the increased multiplicity of even weakly-preferred STRs can enhance binding (**Fig. 5J,K**).

### TF binding STRs is widespread across structural families and organisms

To determine if STR binding is unique to Pho4 and MAX or widespread, we analyzed PBM data for 1291 TFs from 114 species, including *S. cerevisiae, A. thaliana, D. melanogaster, C. elegans, M. musculus*, and *H. sapiens* (*2*, *5*, *94*). For each experiment for each TF, we iterated through all 65,536 (4^8^) 8-mers, computed median intensities for all probes with a given 8-mer, and calculated Z-scores relative to this distribution for all 39 non-redundant homopolymeric, dinucleotide repeat, and tetranucleotide 8-mer STRs (**Figs. 6A, S92; Supplementary Methods**). TF preference for STRs was ubiquitous, with 90% (1158/1291) of all TFs binding at least one STR with p < 1.3·10^-3^ (the Bonferroni-corrected threshold for significance) (**Figs. 6A, S92-95**), and STR preferences varied widely across TF families. Some families (*e.g*. nuclear hormone receptors, T-box, and bZIP) show little preference for any STRs, while others (*e.g*. AT hook, E2F, and ARID/BRIGHT families) prefer STRs simply because they resemble the known consensus (**Figs. 6B, 6C, S96**). More interestingly, members of multiple families (*e.g*. AP2, Forkhead, GATA, homeodomain, Myb/SANT, zinc fingers, and bHLH) weakly prefer particular STRs (**Figs. 6B, 6C, S96**) that often have little sequence similarity to the known motif (as quantified by Levenshtein distance, **Fig. 6D**). Across all TFs, AATT and CCGG repeats are most preferred, largely because these STRs resemble known motifs for the two most abundant TF families (homeodomain and zinc finger TFs, respectively (*6*)) (**Fig. 6E**); C homopolymers are most disfavored (**Fig. S97**).

**Fig. 6:**
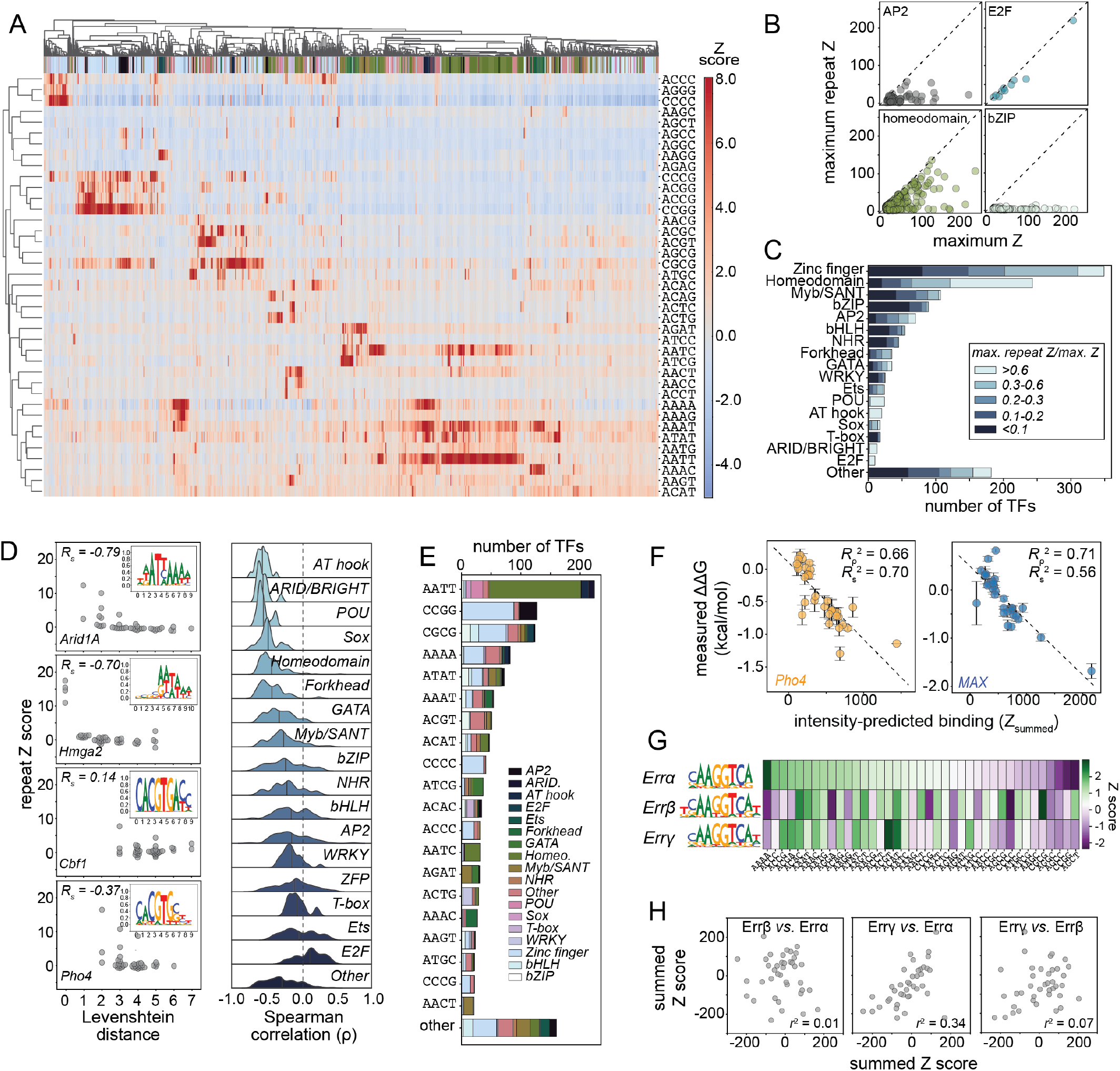
Most TFs show statistically significant binding to repetitive sequences. **(A)** Heat map showing calculated 8-mer intensity Z score for 1,291 TFs (columns) interacting with the 39 non-redundant tetrameric repeat sequences (*i.e*. reverse complements are considered a single sequence). **(B)** Maximum repeat Z score *vs*. maximum overall Z score for TFs from 4 different structural families (AP2, E2F, homeodomain, and bZIP). **(C)** Distributions of ratios of maximum repeat Z scores relative to maximum overall Z scores across TF families. **(D)** Left: Repeat Z score as a function of Levenshtein distance from preferred consensus sequence for Arid1a, Hmga2, Cbf1, and Pho4 (left); insets show PWM representations of preferred consensus sequences (downloaded from CIS-BP (*2*)). Right: Distributions of Spearman correlation coefficients between repeat Z score and Levenshtein distance from consensus across 17 different TF structural families. **(E)** Histogram showing the number of TFs that prefer a particular tetranucleotide repeat, shaded by TF family. **(F)** Scatter plots, linear regressions, and correlation coefficients for measured ΔΔGs *vs*. summed Z scores (intensity-predicted binding) across all measured repeats for Pho4 (left) and MAX (right). **(G)** PSAM (top) and heat map showing 8-mer Z scores for 3 NHR paralogs from *M. musculus* (Errα, Errβ, and Errγ). **(H)** Pairwise comparisons of predicted binding (calculated by summing Z scores) for consensus motifs surrounded by 50 bp on either side of tetranucleotide repeats Errα, Errβ, and Errγ.

### Differential STR preferences could allow closely related paralogs to target distinct genes

Many closely-related paralogs with conserved DBDs and near-identical consensus motif preferences bind and regulate distinct gene targets *in vivo*, and this differential binding has been attributed to either subtle differences in motif (*19*) or flanking nucleotide (*14*, *26*, *27*, *79*, *84*) preferences or direct binding by poorly conserved regions outside of the DBD (*86*). As an alternate hypothesis, differential STR preferences could drive paralog-specific localization. Global comparisons of preferred STRs and preferred motifs across paralogs within a species (quantified via cosine similarity, see **Supplementary Methods**) revealed many TF pairs with highly similar motifs but divergent STR preferences (**Figs. S98-103**), particularly for bHLH and nuclear hormone receptor (NHR) TF paralogs in *A. thaliana* and *M. musculus* (**Figs. S104-106**).

Uncalibrated summed 8-mer Z scores for Pho4 and MAX binding to DNA Library 2 sequences correlated well with measured ΔΔGs (*R*^2^ = 0.66 and 0.71 for Pho4 and MAX, only slightly worse than for calibrated partition function-based predictions) (**Fig. 6F**), suggesting that existing PBM measurement can be used to estimate binding to arbitrary sequences even without quantitative affinity measurements. Predicted binding of the Errα, Errβ, and Errγ NHR TFs from *M. musculus* (which have near-identical PWMs but distinct STR preferences) to sequences containing the consensus surrounded by 50 bp (on either side) of random sequence or STRs showed significant differences (*R*^2^ = 0.01, 0.34, and 0.07; **Figs. 6G,H**), consistent with the hypothesis that sensitivity to STRs could differentially localize paralogs.

### STRs are associated with active enhancers and high mutation rates

STRs can either enhance or decrease TF binding energies; however, the lower bound of affinity imposed by non-specific, electrostatic-mediated interactions skews STR effects to predominantly enhance binding (**Fig. S107**). Consistent with a primarily activating role, STRs are most enriched within the most active enhancers (R_S_^2^ = 0.67, as measured by CAGE-seq, p300 ChIP, GRO-seq, or similar enhancer activity assay (*95*); control datasets shuffling enhancer sequences and measured activity show no significant correlation (R_S_^2^ = 0.16)) (**Fig. 7A**). We also find that STRs are preferentially enriched in enhancers that are broadly active across 278 human cell types (R_S_^2^ = 0.85; shuffled negative control datasets show no enrichment (R_S_^2^ = 0.02)) **Fig. 7B**). Across various eukaryotic genomes, mutations in STRs occur several orders of magnitude more frequently than short insertions and deletions (indels, 1-3bp) and base substitutions (**Fig. 7C**), suggesting that STRs can provide an easily evolvable mechanism to tune transcription (*31*, *40*, *96*).

**Fig. 7:**
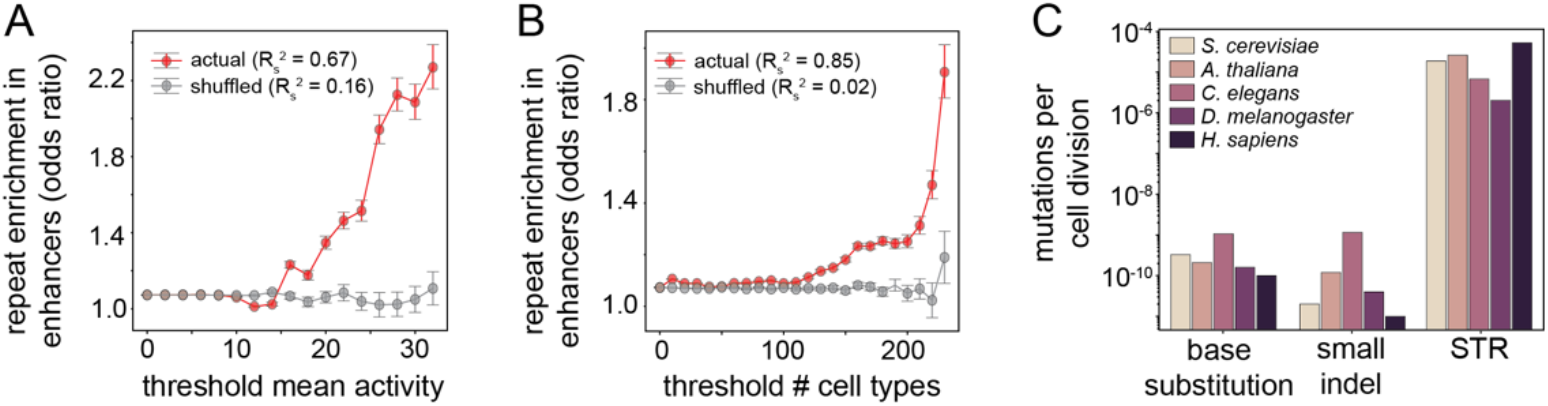
Binding to repetitive sequences is a broad phenomenon across many TFs and TF families. **(A)** Enrichment of STRs in enhancers (red) *vs*. shuffled negative controls (gray) as a function of mean enhancer activity. Error bars are 95% confidence intervals. **(B)** Enrichment of STRs in enhancers (red) *vs*. shuffled negative controls (gray) as a function of the number of cell types within which an enhancer is active. Error bars are 95% confidence intervals. **(C)** Calculated rate of mutation (per cell division) for base substitutions, small indels, and STRs in 5 different model organisms.

## Discussion

The role of STRs in transcriptional regulation has been thoroughly documented, yet the mechanism by which they alter gene expression is poorly understood. Here, we present a model in which STRs directly recruit TFs, thus establishing STRs as a novel class of regulatory elements. Our model is consistent with prior work suggesting that STRs tune gene expression by modulating nucleosome occupancy (*31*), as TF binding, especially that of pioneer factors, is the primary determinant of chromatin accessibility (*68*, *97*–*100*). However, this model allows for more sophisticated regulation: rather than uniformly altering chromatin accessibility, STRs can differentially impact binding for even closely related TFs, serving as rheostats to precisely tune TF binding at a specific locus (*101*–*105*). Moreover, with relatively few types of STRs relative to the number of different TFs, STRs in the absence of known motifs can recruit a diverse set of TFs, thereby functioning as general regulatory elements, in line with observations that STR-enriched enhancers are broadly active across cell types ((*52*), **Fig. 7**). Finally, STRs need not surround a TF consensus motif to have a regulatory effect. Rather, STRs may sequester TFs for precise temporal control of transcription, as hypothesized for pericentromeric satellites regulating the timing of chromosomal replication (*106*).

In contrast to the canonical model that long residence times confer specificity and function while TF search is non-specific and diffusion-limited (*107*), we find that favorable STRs surrounding target motifs alter affinities primarily by increasing macroscopic TF association rates. These results contradict prior microfluidic measurements suggesting that DNA sequence variation primarily impacts dissociation rates; however, we note that prior experiments did not include dark competitor and therefore likely observed a convolved process of dissociation and rebinding (*107*, *108*). Thus, we join other recent work in challenging the canonical view that protein-nucleic acid binding affinities are primarily determined by dissociation rates (*88*). Our measurements can be explained by a simple 4-state model that suggests STRs enhance affinities by increasing the rate of DNA association, in line with prior work suggesting that degenerate recognition sites may serve as “DNA antennae” to attract TFs to a particular regulatory site (*89*, *109*–*111*). However, this 4-state model likely underestimates the true impacts of STRs on target search, as it does not explicitly consider whether TFs can move from flanking STRs to a central motif via one-dimensional sliding, hopping, and intersegmental transfer (*112*–*115*), rather than dissociate, diffuse, and rebind. Future experiments will be required to deconvolve the kinetic contributions of non-specific, electrostatic-mediated binding from other “testing” states for different TF structural classes.

As TFs recruit transcriptional co-activators via “fuzzy,” multivalent (*116*–*118*), and allovalent (*119*) interactions, the finding that STRs enhance the local concentration and residence times of TFs near genomic target sites raises the intriguing possibility that dense clusters of loosely bound TFs could enhance recruitment of co-activator proteins to ensure fast transcriptional response kinetics. This hypothesis is supported by the observation that STRs in budding yeast are enriched near binding sites of stress response TFs (*31*) for which a rapid transcriptional response may be especially advantageous.

This case study of STRs further underscores the limits of motif-centric models in predicting TF occupancy from sequence, as STRs composed of overlapping instances of even low-affinity sites bearing little resemblance to the known motif can dramatically alter binding. Binding of the same TF to dissimilar motifs has previously been reported and attributed to alternate binding modes driven by either entropic or enthalpic effects (*120*–*122*). Here, we show that statistical mechanical models that explicitly account for weak affinity binding dramatically improve quantitative binding predictions for arbitrary DNA sequences relative to motif-based approaches. In future work, small sets of absolute affinity measurements across many TFs could be combined with statistical mechanical and machine learning models to enable quantitative predictions of how changes in nuclear TF concentration alter cooperation and competition between TFs to drive unique transcriptional programs.

As our statistical mechanics framework is agnostic to the identity of binding partners and considers only a distribution of binding energies, we anticipate that the same physical considerations by which DNA-binding proteins recognize STRs may also apply to RNA-binding proteins (RBPs). Evidence in the literature already points to a role for intronic STRs in regulating splicing (*123*–*133*) or promoting the formation of RNP compartments (*134*–*136*). These observations raise the intriguing possibility that STR-enriched enhancers could serve a dual function to bind TFs to regulate transcription and to subsequently recruit RBPs once transcribed into enhancer RNAs.

STRs are highly evolvable (*96*, *137*), requiring only slipped-strand mispairing during replication, repair, or recombination to mutate (*138*, *139*), and may therefore serve as the raw material for evolving new *cis*-regulatory elements (*31*, *96*, *140*) and fine-tuning existing regulatory modules for sensitive transcriptional programs, such as those in development (*141*). This work may motivate future efforts to assess evolution of regulatory networks across species by considering not only conservation of nucleotides within motifs, but also the types and lengths of STRs surrounding them. Evolution of regulatory STRs is likely complemented by co-evolution of TF binding preferences, consistent with a model in which DNA-binding domains exist as a conformational ensemble of partially folded states where single residue substitutions alter the distribution of states within the ensemble and therefore tune the specificity or promiscuity of binding (*73*, *142*–*145*). The observation that STRs disrupt gene expression by directly altering TF binding may provide new clinical insights and therapeutic directions for a variety of STR-associated diseases, from autism (*54*, *55*) to microsatellite instability-associated cancers (*146*, *147*) and others yet to be discovered.

## Materials and methods summary

A complete description of Materials and Methods is included in the Supplementary Information. Briefly, microfluidic devices were fabricated and aligned to printed oligonucleotide or plasmid DNA arrays as described previously (*71*, *142*). Microfluidic devices were controlled by a custom pneumatic manifold (*148*) and imaged with a fully automated microscope and custom software (*73*, *142*). Single-stranded DNA oligonucleotide libraries were synthesized by Integrated DNA Technologies (IDT) and fluorescently labeled and duplexed with a primer extension step. For MITOMI assays, eGFP-tagged TFs were expressed off-chip and purified with anti-eGFP antibodies on the device. Printed fluorescent DNA was solubilized in TBS or wheat germ extract and allowed to bind to immobilized TF for 90 minutes before washing unbound species and imaging. For STAMMP assays, eGFP-tagged TFs were expressed and purified on-chip and increasing concentrations of fluorescently labeled dsDNA were flowed over the chip and allowed to bind for 50 minutes before washing and imaging. Binding was quantified as a ratio of DNA fluorescence to TF fluorescence and the resulting data for multiple concentrations of DNA were fit to a Langmuir isotherm to extract *K_d_* and ΔΔG values. For kinetic measurements, excess unlabeled (“dark”) dsDNA was iteratively introduced in solution and button valves were opened to allow dissociation. Macroscopic dissociation rates (*k_off_*) were fit to the ratio of DNA fluorescence to TF fluorescence over several timepoints to an exponential decay. Macroscopic association rates (*k_on_*) were inferred by *k_on_*=*k_off_/K_d_*, assuming a two-state macroscopic binding model. ChIP-seq data for MAX were downloaded from the ENCODE portal (*1*, *90*) with accession numbers ENCSR000EZM (control) and ENCSR000EZF (experiment). Neural net architecture was adapted from BPNet (*10*) and trained on IDR peaks, with regions from chromosomes 8 and 9 used as the test set and regions from chromosomes 16, 17, and 18 used as the tuning set for hyperparameter tuning. All neural network models were implemented and trained in Keras (v.2.2.4; TensorFlow backend v.1.14) (*149*, *150*) using the Adam optimizer (*151*).

AffinityDistillation scores (Δlog(counts)) were calculated by inserting a given sequence at the center of 100 different background sequences and computing the mean of the differences between the log(count) predictions for query sequence and background sequence alone, as described in (*13*). Universal protein-binding microarray (uPBM) data and associated Z-scores for all possible 8-mers were downloaded from CIS-BP (*2*) and filtered for data quality. STRs in the human genome were identified using Tandem Repeats Finder (*152*). Genome annotations used to calculate enrichment of STRs in enhancers were downloaded from Enhancer Atlas (*95*), FANTOM 5 (*153*), and HACER (*154*) databases. Mutation rates per cell division were cited or calculated from (*137*, *155*–*157*).

## Author contributions

CAH and MGBH measured transcription factor-DNA affinities and kinetics in MITOMI microfluidic assays, and CAH performed EMSA assays. AKA performed STAMMP assays for Pho4 mutant library. AMA, AS, and AK developed, trained, and tested neural net models on genomic binding data. CAH and JMS designed and implemented Gillespie algorithm simulation of TF binding kinetics. EM derived and fit microscopic rate model. CAH, AKA, and MGBH fabricated microfluidic molds and devices. CAH performed bioinformatic analyses and statistical mechanical modeling. NS and JZ guided analysis of chromatin immunoprecipitation data. CAH, AA, RG, JZ, AK, WJG, and PMF conceived of the study. CAH and PMF wrote the manuscript, with input from all authors. PMF supervised the project.

## Acknowledgements

We thank Craig Markin, Nicole DelRosso, Peter Suzuki, Samuel Thompson, Daniel Le, Darach Miller, Sasha Levy, Daniel Herschlag, Jarod Rutledge, and members of the Fordyce Lab for technical assistance, valuable discussions, and feedback on the manuscript. **Funding:** P.M.F. is a Chan Zuckerberg Biohub Investigator and this work was supported by NIH R01-GM117106-01 awarded to R.G., NIH DP2-GM123641 awarded to PMF, and NSF CAREER 2142336 awarded to PMF. A.K.A. was supported by the ChEM-H Chemistry-Biology Interface predoctoral training program. E.M. was supported by the Swedish Research Council grant 2020-06459. J.M.S., A.K.A. and M.G.B.H. were supported by the NSF GRFP. **Data and materials availability:** all data acquired in this study are available in an Open Science Foundation Repository (https://osf.io/gbxhz/). All code required to reproduce analyses in this paper are available on GitHub (https://github.com/FordyceLab/STR_analysis).

## Notes

https://osf.io/gbxhz/

https://github.com/FordyceLab/STR_analysis

### Competing Interest Statement

The authors have declared no competing interest.

### Summary of Updates

This version of the manuscript has been revised to include citations to important prior work.

## References

1. ENCODE Project Consortium, Nature. 489, 57–74 (2012).

2. M. T. Weirauch et al., Cell. 158, 1431–1443 (2014).

3. G. Badis et al., Science. 324, 1720–1723 (2009).

4. C. Zhu et al., Genome Res. 19, 556–566 (2009).

5. M. F. Berger et al., Nat. Biotechnol. 24, 1429–1435 (2006).

6. S. A. Lambert et al., Cell. 172, 650–665 (2018).

7. A. Jolma et al., Cell. 152, 327–339 (2013).

8. J. Yan et al., Nature. 591, 147–151 (2021).

9. M. Slattery et al., Trends Biochem. Sci. 39, 381–399 (2014).

10. Ž. Avsec et al., Nat. Genet. 53, 354–366 (2021).

11. Ž. Avsec et al., Nat. Biotechnol. 37, 592–600 (2019).

12. T. Kaplan et al., PLoS Genet. 7, e1001290 (2011).

13. A. M. Alexandari et al., Manuscr. Prep.

14. X. Zhou, E. K. O’Shea, Mol. Cell. 42, 826–836 (2011).

15. A. Tanay, Genome Res. 16, 962–972 (2006).

16. S. Feng et al., “Transcription factor paralogs orchestrate alternative gene regulatory networks by contextdependent cooperation with multiple cofactors” (preprint, Molecular Biology, 2022),, doi:10.1101/2022.02.26.482133.

17. T. Gera, F. Jonas, R. More, N. Barkai, “Evolution of binding preferences among whole-genome duplicated transcription factors” (preprint, Genomics, 2021),, doi:10.1101/2021.07.27.453962.

18. C. A. Shively, J. Liu, X. Chen, K. Loell, R. D. Mitra, Proc. Natl. Acad. Sci. 116, 16143–16152 (2019).

19. N. Shen et al., Cell Syst. 6, 470–483.e8 (2018).

20. T. Siggers, M. H. Duyzend, J. Reddy, S. Khan, M. L. Bulyk, Mol. Syst. Biol. 7, 555 (2011).

21. J. Crocker et al., Cell. 160, 191–203 (2015).

22. J. F. Kribelbauer, C. Rastogi, H. J. Bussemaker, R. S. Mann, Annu. Rev. Cell Dev. Biol. 35, 357–379 (2019).

23. C. G. de Boer et al., Nat. Biotechnol. 38, 56–65 (2020).

24. B. J. Vincent, J. Estrada, A. H. DePace, Integr. Biol. 8, 475–484 (2016).

25. H. Redden, H. S. Alper, Nat. Commun. 6, 7810 (2015).

26. D. D. Le et al., Proc. Natl. Acad. Sci. U. S. A. 115, E3702–E3711 (2018).

27. M. Levo et al., Genome Res. 25, 1018–1029 (2015).

28. A. Afek, J. L. Schipper, J. Horton, R. Gordân, D. B. Lukatsky, Proc. Natl. Acad. Sci. U. S. A. 111, 17140–17145 (2014).

29. S. Nurk et al., “The complete sequence of a human genome” (preprint, Genomics, 2021),, doi:10.1101/2021.05.26.445798.

30. E. S. Lander et al., Nature. 409, 860–921 (2001).

31. M. D. Vinces, M. Legendre, M. Caldara, M. Hagihara, K. J. Verstrepen, Science. 324, 1213–1216 (2009).

32. S. Sawaya et al., PloS One. 8, e54710 (2013).

33. M. Gymrek et al., Nat. Genet. 48, 22–29 (2016).

34. A. Contente, A. Dittmer, M. C. Koch, J. Roth, M. Dobbelstein, Nat. Genet. 30, 315–320 (2002).

35. H. Hamada, M. Seidman, B. H. Howard, C. M. Gorman, Mol. Cell. Biol. 4, 2622–2630 (1984).

36. F. Gebhardt, K. S. Zänker, B. Brandt, J. Biol. Chem. 274, 13176–13180 (1999).

37. S. Shimajiri et al., FEBS Lett. 455, 70–74 (1999).

38. K. M. Warpeha et al., FASEB J. Off. Publ. Fed. Am. Soc. Exp. Biol. 13, 1825–1832 (1999).

39. R. I. Richards, K. Holman, S. Yu, G. R. Sutherland, Hum. Mol. Genet. 2, 1429–1435 (1993).

40. A. Sulovari et al., Proc. Natl. Acad. Sci. U. S. A. 116, 23243–23253 (2019).

41. A. J. Hannan, Nat. Rev. Genet. 19, 286–298 (2018).

42. A. C. Johnson, Y. Jinno, G. T. Merlino, Mol. Cell. Biol. 8, 4174–4184 (1988).

43. B. Wang, J. Ren, L. L. P. J. Ooi, S. S. Chong, C. G. L. Lee, Oncogene. 24, 3999–4008 (2005).

44. A. Heidari et al., Gene. 492, 195–198 (2012).

45. R. Meloni, V. Albanèse, P. Ravassard, F. Treilhou, J. Mallet, Hum. Mol. Genet. 7, 423–428 (1998).

46. Y.-H. Chen et al., Hum. Genet. 111, 1–8 (2002).

47. R. Srinivasan et al., Sci. Adv. 6, eaaz9115 (2020).

48. M. Goldshtein et al., Biophys. J. 118, 2015–2026 (2020).

49. Q. Lu, L. L. Wallrath, H. Granok, S. C. Elgin, Mol. Cell. Biol. 13, 2802–2814 (1993).

50. A. Orian et al., Genes Dev. 17, 1101–1114 (2003).

51. J. T. Streelman, T. D. Kocher, Physiol. Genomics. 9, 1–4 (2002).

52. J. O. Yáñez-Cuna et al., Genome Res. 24, 1147–1156 (2014).

53. A. J. Hannan, Trends Genet. TIG. 26, 59–65 (2010).

54. I. Mitra et al., Nature. 589, 246–250 (2021).

55. B. Trost et al., Nature. 586, 80–86 (2020).

56. S. F. Fotsing et al., Nat. Genet. 51, 1652–1659 (2019).

57. T. Raveh-Sadka et al., Nat. Genet. 44, 743–750 (2012).

58. V. Iyer, K. Struhl, EMBO J. 14, 2570–2579 (1995).

59. C. Fiore, B. A. Cohen, Genome Res. 26, 778–786 (2016).

60. F. Liu, J. W. Posakony, PLoS Genet. 8, e1002796 (2012).

61. J. Erceg et al., PLoS Genet. 10, e1004060 (2014).

62. L. M. Liberman, A. Stathopoulos, Dev. Biol. 327, 578–589 (2009).

63. R. W. Lusk, M. B. Eisen, PLoS Genet. 6, e1000829 (2010).

64. I. Sela, D. B. Lukatsky, Biophys. J. 101, 160–166 (2011).

65. A. Afek, D. B. Lukatsky, Biophys. J. 105, 1653–1660 (2013).

66. A. R. Iglesias, E. Kindlund, M. Tammi, C. Wadelius, Gene. 341, 149–165 (2004).

67. A. Afek, H. Cohen, S. Barber-Zucker, R. Gordân, D. B. Lukatsky, PLoS Comput. Biol. 11, e1004429 (2015).

68. A. Afek, I. Sela, N. Musa-Lempel, D. B. Lukatsky, Biophys. J. 101, 2465–2475 (2011).

69. M. Goldshtein, D. B. Lukatsky, Biophys. J. 112, 2047–2050 (2017).

70. S. J. Maerkl, S. R. Quake, Science. 315, 233–237 (2007).

71. P. M. Fordyce et al., Nat. Biotechnol. 28, 970–975 (2010).

72. M. Geertz, D. Shore, S. J. Maerkl, Proc. Natl. Acad. Sci. U. S. A. 109, 16540–16545 (2012).

73. A. K. Aditham, C. J. Markin, D. A. Mokhtari, N. DelRosso, P. M. Fordyce, Cell Syst. 12, 112–127.e11 (2021).

74. E. M. O’Neill, A. Kaffman, E. R. Jolly, E. K. O’Shea, Science. 271, 209–212 (1996).

75. K. Vogel, W. Hörz, A. Hinnen, Mol. Cell. Biol. 9, 2050–2057 (1989).

76. C. Bouchard, P. Staller, M. Eilers, Trends Cell Biol. 8, 202–206 (1998).

77. A. Cascón, M. Robledo, Cancer Res. 72, 3119–3124 (2012).

78. R. Rohs et al., Nature. 461, 1248–1253 (2009).

79. R. Gordân et al., Cell Rep. 3, 1093–1104 (2013).

80. M. A. H. Samee, B. G. Bruneau, K. S. Pollard, Cell Syst. 8, 27–42.e6 (2019).

81. S. Pal, J. Hoinka, T. M. Przytycka, Nucleic Acids Res. 47, 6632–6641 (2019).

82. T. Zhou et al., Proc. Natl. Acad. Sci. U. S. A. 112, 4654–4659 (2015).

83. M. F. Berger, M. L. Bulyk, Nat. Protoc. 4, 393–411 (2009).

84. T. Siggers, J. Reddy, B. Barron, M. L. Bulyk, Mol. Cell. 55, 640–648 (2014).

85. Y. Zhang, T. D. Ho, N. E. Buchler, R. Gordân, Genome Res. 31, 1216–1229 (2021).

86. S. Brodsky et al., Mol. Cell. 79, 459–471.e4 (2020).

87. T. Shimizu et al., EMBO J. 16, 4689–4697 (1997).

88. E. Marklund et al., Science. 375, 442–445 (2022).

89. M. Castellanos, N. Mothi, V. Muñoz, Nat. Commun. 11, 540 (2020).

90. C. A. Davis et al., Nucleic Acids Res. 46, D794–D801 (2018).

91. M. B. Gerstein et al., Nature. 489, 91–100 (2012).

92. S. M. Lundberg, S.-I. Lee, in Advances in Neural Information Processing Systems, I. Guyon et al., Eds. (Curran Associates, Inc., 2017; https://proceedings.neurips.cc/paper/2017/file/8a20a8621978632d76c43dfd28b67767-Paper.pdf), vol. 30.

93. A. Shrikumar, P. Greenside, A. Kundaje, in Proceedings of the 34th International Conference on Machine Learning, D. Precup, Y. W. Teh, Eds. (PMLR, 2017; https://proceedings.mlr.press/v70/shrikumar17a.html), vol. 70 of *Proceedings of Machine Learning Research*, pp. 3145–3153.

94. M. A. Hume, L. A. Barrera, S. S. Gisselbrecht, M. L. Bulyk, Nucleic Acids Res. 43, D117–D122 (2015).

95. T. Gao, J. Qian, Nucleic Acids Res. 48, D58–D64 (2020).

96. R. Gemayel, M. D. Vinces, M. Legendre, K. J. Verstrepen, Annu. Rev. Genet. 44, 445–477 (2010).

97. E. Segal, J. Widom, Nat. Rev. Genet. 10, 443–456 (2009).

98. P. Korber, T. Luckenbach, D. Blaschke, W. Hörz, Mol. Cell. Biol. 24, 10965–10974 (2004).

99. A. Valouev et al., Genome Res. 18, 1051–1063 (2008).

100. S. L. Klemm, Z. Shipony, W. J. Greenleaf, Nat. Rev. Genet. 20, 207–220 (2019).

101. S. Meinhardt, M. W. Manley, D. J. Parente, L. Swint-Kruse, PLoS ONE. 8, e83502 (2013).

102. F. M. V. Rossi, A. M. Kringstein, A. Spicher, O. M. Guicherit, H. M. Blau, Mol. Cell. 6, 723–728 (2000).

103. S. Bensmihen et al., Plant Cell. 14, 1391–1403 (2002).

104. M. D. Ilsley et al., Nucleic Acids Res. 45, 6572–6588 (2017).

105. A. Rizzino, Biochem. J. 411, e5–e7 (2008).

106. A. K. Csink, S. Henikoff, Trends Genet. 14, 200–204 (1998).

107. O. G. Berg, R. B. Winter, P. H. von Hippel, Biochemistry. 20, 6929–6948 (1981).

108. T. Paramanathan, D. Reeves, L. J. Friedman, J. Kondev, J. Gelles, Nat. Commun. 5, 5207 (2014).

109. T. Jana, S. Brodsky, N. Barkai, Trends Genet. 37, 421–432 (2021).

110. J. Iwahara, Y. Levy, Transcription. 4, 58–61 (2013).

111. I. Dror, R. Rohs, Y. Mandel-Gutfreund, BioEssays. 38, 605–612 (2016).

112. S. Redding, E. C. Greene, Chem. Phys. Lett. 570, 1–11 (2013).

113. A. B. Kolomeisky, Phys Chem Chem Phys. 13, 2088–2095 (2011).

114. A. Bhattacherjee, Y. Levy, Nucleic Acids Res. 42, 12404–12414 (2014).

115. M. Bauer, R. Metzler, PloS One. 8, e53956 (2013).

116. L. M. Tuttle et al., Cell Rep. 22, 3251–3264 (2018).

117. K. Shrinivas et al., Mol. Cell. 75, 549–561.e7 (2019).

118. A. L. Sanborn et al., eLife. 10, e68068 (2021).

119. P. Klein, T. Pawson, M. Tyers, Curr. Biol. 13, 1669–1678 (2003).

120. E. Morgunova et al., eLife. 7, e32963 (2018).

121. J. M. Rogers et al., Mol. Cell. 74, 245–253.e6 (2019).

122. L. Zhang et al., Genome Res. 28, 111–121 (2018).

123. D. E. Riley, J. N. Krieger, Gene. 344, 203–211 (2005).

124. K. Sathasivam et al., Proc. Natl. Acad. Sci. 110, 2366–2370 (2013).

125. M. Baralle, T. Pastor, E. Bussani, F. Pagani, Am. J. Hum. Genet. 83, 77–88 (2008).

126. A. A. Shishkin et al., Mol. Cell. 35, 82–92 (2009).

127. J. Hui, K. Stangl, W. S. Lane, A. Bindereif, Nat. Struct. Biol. 10, 33–37 (2003).

128. Ł. J. Sznajder, M. S. Swanson, Int. J. Mol. Sci. 20, 3365 (2019).

129. Ł. J. Sznajder et al., Proc. Natl. Acad. Sci. 115, 4234–4239 (2018).

130. C. A. Nutter et al., Genes Dev. 33, 1635–1640 (2019).

131. C. S. Shelley, F. E. Baralle, Nucleic Acids Res. 15, 3787–3799 (1987).

132. H. Cuppens et al., J. Clin. Invest. 101, 487–496 (1998).

133. J. Hui et al., EMBO J. 24, 1988–1998 (2005).

134. K. Yap et al., Mol. Cell. 72, 525–540.e13 (2018).

135. G. V. Echeverria, T. A. Cooper, Brain Res. 1462, 100–111 (2012).

136. K. Ninomiya, T. Hirose, Non-Coding RNA. 6, 6 (2020).

137. M. Lynch et al., Proc. Natl. Acad. Sci. 105, 9272–9277 (2008).

138. C. E. Pearson, K. N. Edamura, J. D. Cleary, Nat. Rev. Genet. 6, 729–742 (2005).

139. G. Levinson, G. A. Gutman, Mol. Biol. Evol. 4, 203–221 (1987).

140. I. Tirosh, N. Barkai, K. J. Verstrepen, J. Biol. 8, 95 (2009).

141. E. K. Farley et al., Science. 350, 325–328 (2015).

142. C. J. Markin et al., Science. 373, eabf8761 (2021).

143. R. K. Das, S. L. Crick, R. V. Pappu, J. Mol. Biol. 416, 287–299 (2012).

144. N. Lyle, R. K. Das, R. V. Pappu, J. Chem. Phys. 139, 121907 (2013).

145. C. G. Kalodimos et al., Science. 305, 386–389 (2004).

146. I. Cortes-Ciriano, S. Lee, W.-Y. Park, T.-M. Kim, P. J. Park, Nat. Commun. 8, 15180 (2017).

147. R. Wooster et al., Nat. Genet. 6, 152–156 (1994).

148. K. Brower et al., HardwareX. 3, 117–134 (2018).

149. F. Chollet, Keras (2018; https://github.com/fchollet/keras).

150. M. Abadi et al., ArXiv160304467 Cs (2016) (available at http://arxiv.org/abs/1603.04467).

151. D. P. Kingma, J. Ba, ArXiv14126980 Cs (2017) (available at http://arxiv.org/abs/1412.6980).

152. G. Benson, Nucleic Acids Res. 27, 573–580 (1999).

153. R. Andersson et al., Nature. 507, 455–461 (2014).

154. J. Wang et al., Nucleic Acids Res. 47, D106–D112 (2019).

155. S. Ossowski et al., Science. 327, 92–94 (2010).

156. T. N. Marriage et al., Heredity. 103, 310–317 (2009).

157. J. M. Watson et al., Proc. Natl. Acad. Sci. U. S. A. 113, 12226–12231 (2016).

