## Supplementary information for "Short tandem repeats bind transcription factors to tune eukaryotic gene expression"

2022

**3 Supplemental tables** **46**

**4 Supplemental figures** **62**

### 1 Supplemental methods

#### 1.1 Fabrication of microfluidic molds and devices

Flow and control molds were fabricated as described previously ([18, 27]) and all design files are available on the Fordyce Lab website (http://www.fordycelab.com/microfluidic-design-files). We fabricated two-layer MITOMI devices from these molds using polydimethylsiloxane (PDMS) polymer (RS Hughes, RTV615) in the Stanford Microfluidics Foundry. To fabricate the control layer, we combined $\sim$ 55 g of PDMS (1:5 ratio of cross-linker to base), mixed and degassed the components within a centrifugal mixer at 2000 and 2200 rpm, respectively, for 3 minutes each (THINKY). We then poured the mixture onto the molds, degassed them in a vacuum chamber for 45 minutes under house vacuum, and baked them in an 80°C convection oven for 45 minutes. We then cut control layers for individual devices from the cast PDMS and punched fluid inlet lines using a drill press (Technical Innovations) with a mounted catheter hole punch (SYNEO, CR0350255N20R4). To fabricate the flow layer, we combined PDMS at a 1:20 ratio (cross-linker to base) and mixed and degassed the components within a centrifugal mixer at 2000 and 2200 rpm, respectively, for 3 minutes each. We then spin-cast the PDMS onto molds for 10 s at 500 rpm followed by 1750–1850 rpm for 65 s. Spin-cast layers were allowed to relax on a flat surface at room temperature for 5 minutes before baking at 80°C for 35–40 minutes. We then manually aligned individual control layers to the partially cured flow layer and baked the aligned devices for an additional 45 minutes at 80°C. Bonded two-layer devices were cut from the flow mold with a scalpel and the flow-layer fluid inlet lines were punched as described above.

#### 1.2 Preparation of MITOMI oligonucleotide arrays

Single-stranded DNA (ssDNA) oligonucleotides in Tables S1, S2 were synthesized by Integrated DNA Technologies (IDT). For each library of 32 or 48 sequences, the library was replicated 3 or 2 times, respectively, to fill a 96-well plate.

##### 1.2.1 DNA labeling and duplexing via Klenow extension

To generate fluorescently-labeled double-stranded DNA, we annealed each ssDNA oligo to a universal primer (IDT) with the following sequence: `5‘-/FLUOR/-GTC ATA CCG CCG GA-3‘`, where FLUOR represents a covalently-attached Alexa 647 or Atto 647N fluorophore. This annealing process took place within 40 $\mu$L reaction volumes in a 96 well plate:

- 12 $\mu$L of 100 $\mu$M single-stranded oligonucleotide DNA
- 12 $\mu$L of 100 $\mu$M fluorescently-labeled primer
- 12 $\mu$L of dNTPs (10 mM each)
- 4 $\mu$L of NEBuffer 2 (10x) (NEB B7002S)

To anneal primers, we incubated using a decreasing temperature ramp:

- Melting: 94°C for 3 minutes
- Annealing: 94 to 37°C with 57 cycles of 50 seconds with -1°C per cycle

We then extended annealed primers using Klenow exo- polymerase (expressed and purified in-house; can substitute NEB, M0212S). The following reagents were added to each well for a total volume of 60 $\mu$L per well:

- 2 $\mu$L of Klenow exo- polymerase
- 2 $\mu$L of NEBuffer 2 (10x)
- 16 $\mu$L of MilliQ water

Extension took place using the following temperature ramp:

- Extension: 37°C for 60 minutes
- Enzyme deactivation: 72°C for 20 minutes
- Slow cool-down: 72 to 10°C with 41 cycles of 1 minute with -1.5°C per cycle

###### 1.2.2 Klenow extension check

We assessed the efficiency of the annealing and extension protocol described above by running oligos on a denaturing gel using the following reaction mixture:

- 1 $\mu$L of 0.94 $\mu$M double-stranded DNA
- 5 $\mu$L of 2x TBE-urea (see below)
- 2 $\mu$L of 5x Orange G loading dye (5x is 0.25% w/v orange G; 1x is 0.05%; Sigma Life Sciences, O7252-25G)
- 2 $\mu$L of MilliQ water

20 mL of 2x TBE-urea was prepared by mixing 4 mL of 10x TBE buffer (tris-borate EDTA; Thermo Scientific, B52), 8.41 g of solid urea (Sigma Life Sciences, U5378-100G), and MilliQ water to 20 mL.

We heated this mixture for 10 minutes at 70°C to promote denaturation and then loaded samples onto a Novex 10% TBE-urea gel (Thermo Scientific, EC68752BOX). The gel was run for 50 minutes at 180V in 1x TBE running buffer (10x buffer diluted in MilliQ water). For short oligos (50-60 bp), we ran gels at room temperature; for long oligos ($>$ 100 bp), we preheated the running buffer to $\sim$55°C to ensure the DNA remained melted. We found that heating the buffer was necessary to keep long oligos from reannealing after removal from the thermal cycler. The un-extended fluorescent primer (`5‘-/FLUOR/-GTC ATA CCG CCG GA-3‘`) was also loaded on the gel as a negative control. Gels were imaged on the Typhoon FLA 9500 (GE Healthcare) with the LPR-Ch2 filter cube (665 nm long pass filter; GE Healthcare) and examined to ensure that oligonucleotides were fully extended and the correct length.

###### 1.2.3 Library dilution

Prior to library spotting, we generated serial dilutions of each oligonucleotide. We first prepared a 5x concentrated stock of "spotting solution" comprised of 5% (w/v) lyophilized bovine serine albumin (Sigma Life Sciences, B4287-25G) and 62.5 mg/mL trehalose dihydrate (Sigma Life Science, T9531-25G) in 15x SSC (saline-sodium citrate). We then made a 1x stock by diluting this 5x stock in MilliQ water and filter-sterilized both stocks before use.

To generate libraries of oligonucleotides at 8 different concentrations for printing, we prepared the following serial dilution series in eight 96-well plates and then transferred 15 $\mu$L of each well of each plate to two 384-well plates:

1. 16 $\mu$M DNA

- 48 $\mu$L Klenow Product
- 12 $\mu$L 5x spotting solution

2. 10.7 $\mu$M DNA
   - 40 $\mu$L 16 $\mu$M DNA (dilution #1)
   - 20 $\mu$L 1x spotting solution
3. 7.1 $\mu$M DNA
   - 40 $\mu$L 10.7 $\mu$M DNA (dilution #2)
   - 20 $\mu$L 1x spotting solution
4. 4.7 $\mu$M DNA
   - 40 $\mu$L 7.1 $\mu$M DNA (dilution #3)
   - 20 $\mu$L 1x spotting solution
5. 3.2 $\mu$M DNA
   - 40 $\mu$L 4.7 $\mu$M DNA (dilution #4)
   - 20 $\mu$L 1x spotting solution
6. 2.1 $\mu$M DNA
   - 40 $\mu$L 3.2 $\mu$M DNA (dilution #5)
   - 20 $\mu$L 1x spotting solution
7. 1.4 $\mu$M DNA
   - 40 $\mu$L 2.1 $\mu$M DNA (dilution #6)
   - 20 $\mu$L 1x spotting solution
8. 0.94 $\mu$M DNA
   - 40 $\mu$L 1.4 $\mu$M DNA (dilution #7)
   - 20 $\mu$L 1x spotting solution

##### 1.2.4 DNA library printing

This library of serial dilutions was then spotted onto 2"x3" epoxysilane-coated glass slides (Thermo Scientific, UCSF2X3-C50-20) or 1"x3" epoxysilane-coated glass slides (Arrayit, SuperEpoxy 2) using either a custom-built microarrayer robot with silicon pins (Parallel Synthesis Technologies, SMT-S75) or the sciFLEXARRAYER S3 (Scienion AG) with a single PDC 70 nozzle with a Type 1 coating. Oligonucleotide arrays were printed in 28 columns of 56 spots/column with spacings of 643 $\mu$m between columns and 321 $\mu$m between spots within a column. For the larger 2"x3" slides, two arrays were deposited on each slide with 31 mm spacing between arrays.

When using the custom-built microarrayer robot, pins were washed twice for 10 seconds in scalding hot water and vacuum dried for 8 seconds between samples in order to prevent contamination.

The sciFLEXARRAYER S3 microarrayer was operated largely according to manufacturer's instructions. Briefly, the instrument and nozzle were primed with particle-free, degassed water. We then optimized water droplets and print buffer droplets for consistency and volume. Finally, we programmed the machine to iteratively aspirate and deposit each well in a plate, as follows:

- Move to next well in plate and take 3 uL of fluorescent oligonucleotide
- Wash outside of nozzle twice
- Wait 32 seconds to allow nozzle exterior to dry
- Pre-spot 1500 drops to improve droplet consistency
- Perform droplet QC (based on displacement from nozzle and presence of a single droplet)
- If QC passed, deposit oligonucleotide solution on slides
- QC droplet again
- Flush nozzle interior and exterior to prevent DNA cross-contamination

After printing and drying arrays at least overnight, we manually aligned MITOMI devices to the DNA array such that each chamber contained one oligonucleotide spot. Devices were then bonded to the glass slide by baking on a hot plate at 95°C for approximately 4 hours (and sometimes up to 12 hours).

#### 1.3 Transcription factor expression for MITOMI

##### 1.3.1 Plasmids for wheat germ extract expression (Pho4 and MAX MITOMI experiments)

The coding sequence for each full-length transcription factor was inserted into the pTNT vector backbone (Promega, L5610) separated from a monomeric eGFP coding sequence [48] by a 10 amino acid gly-ser linker (GGGSGGGGSG). The plasmid backbone contains an ampicillin resistance cassette (beta lactamase gene), T7 promoter and terminator, and Kozak sequence.

**Plasmid backbone preparation for Gibson assembly.** The plasmid backbone, already containing the Gly-Ser linker and eGFP coding sequence, was linearized and amplified by PCR using the following mixture:

- 25 $\mu$L, Q5 Hot Start 2x master mix (New England Biolabs, M0494S)
- 1$\mu$l, plasmid template (5ng/$\mu$l)
- 2.5 $\mu$L, 10 $\mu$M forward primer (final concentration 500 nM)
- 2.5 $\mu$L, 10 $\mu$M reverse primer (final concentration 500 nM)
- 29 $\mu$L, MilliQ water

For amplification, we used the following thermal cycler protocol:

- Initial melting: 98°C, 60 s
- 32 cycles:
  - Melting: 98°C, 10 s
  - Annealing: 67.5°C, 15 s
  - Extension: 72°C, 105 s
- Final extension: 72°C, 120 s

We cleaned up the PCR products with a Zymo DNA Clean and Concentrator-25 kit (Zymo Research, D4005) as per manufacturer's guidelines and verified their lengths with gel electrophoresis. We then treated cleaned products with DpnI to digest any template plasmid by preparing the following mixture and incubating the digestion reaction at 37°C for 20 minutes:

- 15 $\mu$L, plasmid backbone cleaned PCR prodcut
- 1 $\mu$L, DpnI enzyme (New England Biolabs, R0176S)
- 2 $\mu$L, 10x CutSmart buffer (New England Biolabs, B7204S)
- 1 $\mu$L, MilliQ water

**Transcription factor insert preparation for Gibson assembly.** Coding sequences for transcription factors (excluding the stop codon) were amplified by PCR from the yeast genome following a genomic DNA extraction. All transcription factor coding sequences used in this work were prepared in this way, except for MAX, whose coding sequence was ordered as a gBlock gene fragment (Integrated DNA Technologies). To perform the genomic DNA extraction, we grew BY4741 strain yeast overnight in YPD and diluted the cultures 1:2 (with YPD) in the morning (protocol adapted from [28]). We then added 200 $\mu$L of diluted culture to 200 $\mu$L of lysis solution, which we prepared in MilliQ water as follows:

- 200 mM lithium acetate (Sigma-Aldrich, L4158-100G)
- 1% sodium dodecyl sulfate (from 10% stock; Fisher Bioreagents, BP2436-1)
- 10 mM Tris-HCl, pH 8.0 (from 1 M stock; Fisher Bioreagents, BP1758-500)
- 1 mM EDTA, pH 8.0 (from 0.5 M stock; Invitrogen, AM9260G)

The yeast lysis mixture was incubated for 5 minutes at 70°C. We then added 1 $\mu$L (100 $\mu$g) of RNAse A (Qiagen, 19101) to the solution and incubated for an additional 5 minutes at room temperature. Next, we added 600 $\mu$L of 200 proof ethanol (Gold Shield Distributors, 412804) and briefly vortexed the mixture. We centrifuged the mixture at 15,000 x g for 3 minutes to form a compact pellet and then removed the supernatant. The pellet was then resuspended in 950 $\mu$L of 70% ethanol (diluted from 200 proof stock) and centrifuged again at 15,000 x g for 3 minutes. Following removal of the supernatant, we resuspended the pellet in 200 $\mu$L TE (Invitrogen, AM9849). The solution was once again centrifuged for 15 seconds at 15,000 x g to pellet cell debris, and then we used 1 $\mu$L of the supernatant for PCR.

To perform PCR amplification, we prepared the following mixture for each reaction:

- 5 $\mu$L, Q5 Hot Start 2x master mix (New England Biolabs, M0494S)
- 1 $\mu$L, gDNA template
- 0.5 $\mu$L, 10 $\mu$M forward primer (final concentration: 500 nM)
- 0.5 $\mu$L, 10 $\mu$M reverse primer (final concentration: 500 nM)
- 3 $\mu$L, MilliQ water

We then amplified using the following thermal cycler protocol:

- Initial melting: 98°C, 60 s
- 32 cycles:
  - Melting: 98°C, 10 s
  - Annealing: 67.5°C, 15 s
  - Extension: 72°C, 65 s
- Final extension: 72°C, 120 s

We cleaned up the PCR products with a Zymo DNA Clean and Concentrator-25 kit (Zymo Research, D4005) as per manufacturer's guidelines and verified their lengths with gel electrophoresis.

**Gibson assembly.** Transcription factor coding sequences were inserted into the plasmid backbone using the HiFi DNA Assembly kit (New England Biolabs, E5520S). We prepared the following reaction mixture for each assembly:

- 25 fmol plasmid
- 75 fmol insert
- x $\mu$L of 2x NEBuilder HiFi DNA Assembly Cloning Kit (NEB, E5520S) equal to combined volume of plasmid and insert solutions.

The assembly reactions were incubated at 50°C for 1 hour.

**Bacterial transformation.** To transform cells, we added 0.8 $\mu$L of the assembly reaction to 20 $\mu$L of E. coli DH5$\alpha$ cells (New England Biolabs, C2987I), which then sat on ice for 30 minutes. We heat-shocked the cells at 42°C for 30 seconds and transferred them to ice for 5 minutes. We added 380 $\mu$L of SOC rescue media (NEB, B9020S) to each reaction, which then grew for 1 hour at 37°C while shaking at 350 rpm. We added 10 $\mu$L of the culture to 90 $\mu$L of SOC median and plated the 100 $\mu$L on LB+ampicillin plate. Plates were incubated overnight at 37°C. We picked colonies the next day, grew them overnight in LB+ampicillin cultures, and then performed minipreps the following day (Qiagen). Plasmid sequences were verified with Sanger sequencing.

##### 1.3.2 Preparation of linear templates for *in vitro* transcription and translation

We found that linear templates prepared via PCR yielded higher expression than plasmid templates during *in vitro* transcription and translation in wheat germ extract. We therefore generated linear templates from plasmids prior to expression as follows:

- 25 $\mu$L Q5 Hot Start 2x master mix (New England Biolabs, M0494S)
- 2 $\mu$L plasmid template (at 5 ng/$\mu$L concentration for 10 ng total)
- 3.5 $\mu$L of 10 $\mu$M forward primer (see below)
- 3.5 $\mu$L of 10 $\mu$M reverse primer (see below)
- 16 $\mu$L of MilliQ water

We used the following thermal cycling protocol:

- Initial melting: 98°C, 30 s
- 30 cycles:
  - Melting: 98°C, 5 s
  - Annealing: 66.5°C, 10 s
  - Extension: 72°C, 60 s
- Final extension: 72°C, 120 s

We cleaned up the PCR products with a Zymo DNA Clean and Concentrator-25 kit (Zymo Research, D4005) as per manufacturer's guidelines and verified construct lengths with gel electrophoresis.

| Primer | Sequence |
|---|---|
| IVTT linear template forward primer | CCCCTCAAGACCCGTTTAGA |
| IVTT linear template reverse primer | AGGCTAGAGTACTTAATACGACTCACTAT |

##### 1.3.3 Off-chip *in vitro* transcription and translation

For MITOMI assays, transcription factor-eGFP fusions were expressed off-chip (*i.e.* in a microcentrifuge tube, rather than on the microfluidic device) using a wheat germ-based *in vitro* transcription and translation mix (Promega).

For each device, we prepared the following 100 $\mu$L 1x reaction mixture:

- 50 $\mu$L wheat germ extract (Promega)
- 4 $\mu$L TnT buffer (Promega)
- 0.66 $\mu$L amino acids -Cys
- 0.66 $\mu$L amino acids -Met
- 0.66 $\mu$L amino acids -Leu
- 2 $\mu$L recombinant RNAse inhibitor
- 2 $\mu$L T7 polymerase
- 1–2 $\mu$g of plasmid template or 200–300 ng of linear template
- MilliQ water to a final volume of 100 $\mu$L

We scaled reactions by multiplying each reagent volume by the number of devices used on a particular day. To express transcription factors, we incubated the IVTT reaction at 30°C for 3 hours in a shaking dry heat block at 300 rpm. Following expression, the reaction was spun in a tabletop microcentrifuge for 5 minutes at 10000 x g to precipitate aggregates and other insoluble components prior to loading onto the device.

#### 1.4 Microscopy instrumentation

For all MITOMI and STAMMP experiments, devices were mounted on a Nikon Ti-S microscope with an automated XY stage (Applied Scientific Instrumentation, MS-2000 XYZ stage), a solid-state light source (Lumencor, SOLA SE Light Engine), a cMOS camera (Oxford Instruments, Andor Zyla 4.2 cMOS), and an automated filter turret with an eGFP filter set (Chroma Technology Corp., part no. 49002) and a Cy5 filter set (Chroma Technology Corp., part no. 39007). The microscope stage, filter, and camera were controlled using Micro-Manager [17]; all images were acquired with a 4x objective lens (CFI Plan Apochromat $\lambda$ 4X NA 0.20, Nikon) and 2x2 binning (1024x1024 pixels).

#### 1.5 MITOMI device operation and experimental pipeline

MITOMI devices were operated using a custom-built pneumatics manifold, as described in [12]. Reagents were loaded into Tygon tubing (inner diameter 0.02”; Saint-Gobain, AAD04103) via suction from a 1 mL syringe (BD, 309628) at one end connected via 23-gauge luer connector (McMaster-Carr, 75165A684) and a blunt-end steel tube at the other (0.013” ID x 0.025” OD x 0.5” length, New England Small Tube Corporation, NE-1310-02). After loading reagents into the tubing, we inserted the blunt end tube into one of the inlet ports. We then disconnected the other end from the syringe and connected it to the pneumatic manifold, outfitted with another 23-gauge luer connector. Reagent lines were pressurized at 3.0–4.1 psi, depending on the height of features in the flow layer on a particular device. To prepare devices for surface patterning, we dead-end filled control lines with MilliQ H2O at 28–35 psi.

##### 1.5.1 Surface patterning

The surfaces of epoxysilane-coated glass slides were functionalized with eGFP-tagged transcription factor in a manner similar to that described previously [2, 27] and illustrated in Fig. S7. First, we introduced biotinylated bovine serum albumin (bBSA; 2 mg/mL, ThermoFisher Pierce, 29130) into the device with the inlet and outlet valves open until a droplet of liquid formed at the outlet. We then closed the outlet valve and dead-end filled the device for about 5–10 minutes to remove all air. Following removal of air from the device, we flowed biotinylated BSA (bBSA, same as above) through the device for 5 minutes with button valves closed and 30 minutes with button valves open. We then flushed the flow lines with 1x phosphate-buffered saline (PBS, diluted from 10x stock, Corning 46-013-CM) for 10 minutes. After flushing, we flowed NeutrAvidin (1 mg/mL, Thermo Scientific, 31000) through the device for 30 minutes with button valves open, followed by another PBS wash for 10 minutes. Next, we introduced biotinylated BSA for an additional 35 minutes with button valves closed to passivate all surfaces except for those protected by the closed button valve, followed by an additional PBS wash for 10 minutes. Next, we flowed biotinylated anti-eGFP antibody (Abcam, ab6658; diluted in 1x PBS to 100 $\mu$g/mL from a stock concentration of 1 mg/mL) through the device for 2 minutes with button valves closed to allow the antibody solution to perfuse the device. We then opened button valves and flowed this antibody solution through the device for an additional 13 minutes and 20 seconds, followed by another PBS wash for 10 minutes.

##### 1.5.2 Transcription factor loading and immobilization

To immobilize transcription factors to antibody-patterned surfaces within device chambers, we introduced transcription factors expressed off-chip into devices with button valves open. The transcription factor protein solution was allowed to flow through the device either until antibodies became saturated—that is, eGFP fluorescence no longer increased even with continued protein flow—or until all of the protein solution had been loaded onto the device.

##### 1.5.3 Trypsin wash to eliminate non-specifically bound transcription factors

To remove TF-eGFP fusions bound non-specifically on the surface (*i.e.* not under the button valve), we flushed devices with PBS for 10 minutes and then washed with TrypLE enzyme (1X stock, ThermoFisher, 12604-013) for 15 minutes with button valves closed. Devices were then again flushed with PBS for 10 minutes. Next, we replenished blocking proteins on device walls that may have become digested during the TrypLE wash by flowing biotinylated BSA for 15 minutes (again, with buttons closed). Finally, we washed the inlet manifold with PBS for 10 minutes to remove any residual trypsin.

##### 1.5.4 MITOMI equilibrium DNA binding measurements

**DNA solubilization.** To prepare devices for equilibrium binding measurements, we flowed assay buffer onto the device for 10 minutes. In most experiments, we used wheat germ extract (Promega

L4380) as the assay "buffer" to mimic the cellular environment. For control experiments designed to ensure that components within the wheat germ extract were not responsible for repeat binding, we used the following tris-buffered saline solution with dithiothreitol (DTT):

- 10 mM Tris-HCl pH 7.5 (from 1 M stock; Fisher Bioreagents, BP1757-500)
- 150 mM NaCl (Sigma-Aldrich, 71376–1KG)
- 1 mM DTT (Sigma-Aldrich, D9779)

To solubilize the DNA deposited by the arrayer within DNA chambers (see Section 1.2), we introduced assay buffer into the DNA chambers by opening neck valves (with button valves still closed) until the DNA chambers were approximately 95% full. We then closed the neck valves and flushed the device with assay buffer for 10 minutes to wash away any DNA that leaked during the solubilization. Finally, we closed the sandwich valves to separate adjacent reaction chambers and opened neck valves to allow DNA to diffuse from DNA chambers into the binding chamber (again, with button valves closed). DNA was allowed to diffuse into the binding chamber for 15–20 minutes before proceeding to equilibrium binding measurements.

**DNA binding and incubation.** To begin the equilibrium binding measurement, we opened button valves to allow DNA in solution to bind to the surface-immobilized transcription factors. The binding reaction was allowed to proceed for 90 minutes, far exceeding the time needed to reach equilibrium estimated by the following equation [24]:

$$k_{equil} = k_{off} + k_{on}[TF] \tag{1}$$

For the limiting (slowest) case where $[TF] \approx 0$, this can be simplified to [24]:

$$k_{equil} \approx k_{off} \tag{2}$$

From this simplification, reaction half-life can be approximated as:

$$t_{1/2} = \frac{\ln(2)}{k_{equil}} \approx \frac{\ln(2)}{k_{off}} \tag{3}$$

If we approximate our slowest dissociation rate as $k_{off} = 10^{-2}$ (consistent with kinetic measurements presented later), then $t_{1/2} \approx 69.3\ s$. By this formalism, we estimate that the reaction will have reached 99.9% of its equilibrium state in 10 half-lives, or in 693 seconds (∼11.5 minutes).

At the end of the 90 minutes, we first imaged devices in the Cy5 channel to quantify the concentration of DNA in the chamber available for binding ("pre-wash" images). We then closed the button valves to mechanically trap any Cy5-labeled DNA bound to surface-immobilized eGFP-tagged transcription factors and flushed devices with PBS for 10 minutes to remove unbound DNA. After washing, we again imaged devices in the eGFP channel (to quantify surface-immobilized TF concentration) and in the Cy5 channel (to quantify TF-bound DNA); we term these "post-wash" images. Microfluidic devices were imaged with a gridded acquisition, with adjacent images overlapping by 10% to facilitate image stitching (see below for more details). See Section 1.7 below for details regarding how these images were used to estimate $K_d$ values of binding.

##### 1.5.5 eGFP-only negative control binding measurements

To ensure that observed binding resulted from a specific interaction between TFs of interest and DNA and not from other proteins within the wheat germ extract, we repeated binding measurements (as described in Section 1.5) but using eGFP alone instead of eGFP-tagged TFs. For each experiment, we flowed approximately 30 nM purified eGFP (BioVision #4999, diluted in PBS) across devices for approximately 30 minutes.

##### 1.5.6 MITOMI image processing

**Image stitching and flatfield correction.** To account for variations in excitation intensities and emitted photon detection efficiencies across the field of view, we applied a flatfield correction to all images taken in the eGFP and Cy5 channels, as described previously [37]. Following flatfield correction, images taken from each gridded acquisition were stitched together by an in-house imaging stitching program (https://github.com/FordyceLab/ImageStitcher). For equilibrium binding, image stitching yields: (1) a "pre-wash" Cy5 image of the entire device that can be used to calculate the intensity of free DNA available for binding in each chamber, (2) a "post-wash" GFP image of the device that can be used to calculate the intensity of surface-immobilized TF-eGFP molecules in each chamber, and (3) a "post-wash" Cy5 image of the device indicating the intensity of bound Alexa647- or Atto647N-tagged DNA recruited to surface-immobilized TFs. These intensities can then be converted to effective per-chamber TF and DNA concentrations using eGFP and Alexa 647/Atto647N calibration curves (as described below).

**Image quantification.** To quantify DNA and TF concentrations within each chamber, we used a custom in-house software package (https://github.com/FordyceLab/ProcessingPack) as described in [2]. To process "pre-wash" Cy5 images, the software identifies the center of each chamber with a Hough Transform and then quantifies median fluorescence in each chamber. To process "post-wash" images, the software first identifies the centroid position of each button within the eGFP "post-wash" image (which should have relatively constant intensities across buttons) using a grid search algorithm that maximizes the total fluorescence within a circle of fixed size. These centroid button coordinates are then used to quantify the sum of the total fluorescence intensity for each circle within both "post-wash" eGFP and Cy5 images (where intensities vary considerably). To account for differences in local background fluorescence across the device, we subtracted the total intensity of an annulus surrounding the circle (normalized by the ratio of areas of the annulus and button) from each calculated button intensity across both images.

##### 1.5.7 Data quality control

Before analyzing data, we first examined stitched images from each device to exclude any columns of the device with unusually low transcription factor deposition (due to device defects and associated restricted flows). We also excluded any chambers with Cy5 signal in the DNA chambers below local background fluorescence intensities, as these signal errors in oligonucleotide deposition. Finally, we excluded chambers with obvious debris or protein aggregates.

#### 1.6 DNA fluorescence standard curve

To allow conversion of measured Alexa or Atto fluorophore intensities to absolute DNA concentrations, we measured observed fluorescence for a series of fluorescently-labeled DNA "standard" solutions of known concentration. First, we prepared fluorescently-labeled dsDNA constructs as described above in Section 1.2.1 and then passed the Klenow-extended DNA through a 10 kDa molecular weight cutoff filter (Millipore Sigma UFC501096) at 8000 x g for 8 minutes in a tabletop microcentrifuge to filter out free Alexa dye and unincorporated dNTPs. This filtering step was applied three times, each time increasing the volume of input sample to 300 $\mu$L with PBS before centrifugation to facilitate efficient filtering.

Next, we diluted DNA into PBS to yield the following approximate concentrations (assuming a 20 $\mu$M concentration of Klenow-extended DNA): 2.50 $\mu$M, 1.25 $\mu$M, 0.625 $\mu$M, 0.313 $\mu$M, 0.156 $\mu$M, 0.0781 $\mu$M, 0.0391 $\mu$M. Finally, we quantified the concentration of each dilution by measuring absorbance at 260 nm with the DeNovix and using the following formula to calculate concentration (mass/volume):

$$\text{concentration } (\mu g/mL) = A_{260} \cdot 50\ \mu g/mL \tag{4}$$

The mass/volume concentration was then converted to molar concentration:

$$\text{concentration } (M) = \frac{conc_{mass/vol}}{36 + 618 \cdot d} \cdot \frac{10^6 \mu g}{g} \cdot \frac{10^3 mL}{L} \tag{5}$$

where $d$ is the length of the DNA molecule in base pairs.

To measure DNA chamber fluorescence as a function of known concentration, we used the same MITOMI devices and pneumatic setup as described above. Briefly, control lines were pressurized with MilliQ water and reagents were introduced into the device by first loading them into Tygon tubing via syringe suction and then flowing them onto the device with pneumatic pressure. To passivate device surfaces prior to measuring the calibration curve, we introduced biotinylated bovine serum albumin (bBSA; 2 mg/mL, ThermoFisher Pierce, 29130) as described for the first surface patterning step of MITOMI experiments. To introduce each DNA concentration onto the device, we then flowed DNA across the device for 10 minutes with the button valves closed (to ensure equal distribution). After opening button valves, we imaged devices in the Cy5/Alexa 647/Atto647N channel with a gridded acquisition, as described above. Camera exposure times were adjusted to maximize dynamic range without saturating any pixels. This process was repeated for all concentrations from lowest to highest with a 10 minute PBS washing step between each concentration to prevent contamination between samples.

Using the same image stitching and processing pipeline described above, we quantified median fluorescence intensities for each chamber of the device for each concentration of DNA and camera exposure (excluding chambers with obvious technical errors such as dust particles or an absence of fluid flow; data for each experiment are shown in Figs. S8 and S9). We then fit a line ($y = mx + b$) to the relationship between fluorescence intensity and DNA concentration, using all chambers, constraining the intercept (b) to be non-negative:

$$[DNA] = f(fluor) = m \cdot fluor + b,\ b \geq 0 \tag{6}$$

To apply this standard curve to fluorescence intensity measurements taken at different exposures, we used the following formula:

$$[DNA] = g(ex, fluor) = (m \cdot fluor + b) \cdot \frac{ex_0}{ex} \tag{7}$$

where $fluor$ is the fluorescence intensity, $ex$ is the exposure time used in the experiment, and $ex_0$ is the exposure time used in the calibration experiment and $m$ and $b$ are the coefficients relating fluorescence and [DNA] as defined above in Eqn. 6.

##### 1.7 Determination of equilibrium binding constants

To determine equilibrium binding constants from MITOMI TF/DNA concentration-dependent binding curves, we first converted measurements of soluble DNA intensities in "pre-wash images" to effective concentrations available for TF binding using the DNA fluorescence calibration curves described in Section 1.6.

Next, we quantified differences in DNA bound by surface immobilized TFs by calculating the ratio (R) of "postwash" DNA fluorescence (Alexa 647 or Atto 647N) to "postwash" TF fluorescence (GFP).

Finally, we collated concentration-dependent binding data by oligonucleotide sequence within each experiment and fit to a Langmuir isotherm:

$$R = \frac{R_{max} \cdot [DNA]}{K_d + [DNA]} \tag{8}$$

where $R_{max}$ is globally fit for all data on a device (as in [18, 31]) and where $K_d$ is fit for each oligo.

Having observed empirically and in simulations that errors in estimation of $R_{max}$ (the binding ratio saturation value) can lead to systematic errors in estimated affinities ($K_d$s), we sought to fit $K_d$ values in manner robust to outliers. Toward this end, we randomly selected half of the available chambers on a device (e.g. if 1,200 chambers were left after data QC, we selected 600 chambers without replacement) and performed a fit on this subset of data. We then repeated this process 1,000 times per dataset in order to build a distribution of estimated $K_d$ values per oligo per experiment. Reported $K_d$ values and errors represent the mean and standard deviations of the estimated $K_d$ distributions per oligo per experiment calculated via this iterative sampling.

##### 1.8 Data culling and normalization for MITOMI measurements

For each Langmuir isotherm fit in a library (across all experiments and all oligonucleotides), we calculated root mean squared error (rmse) of the fit, normalized by the saturating value ($y_{max}$). We then eliminated all binding measurements with a normalized rmse greater than 1.5 standard deviations above the mean rmse. These data are denoted in the binding curve supplementary files as **CULLED** and were excluded from further analyses.

To account for variation between experiments, we normalized $\Delta\Delta G$ measurements for a given protein across all experiments. First, we chose a reference experiment based on the following criteria: (1) evenness of protein deposition across the device, (2) high number of chambers in a MITOMI device remaining after culling, (3) low average rmse across all curve fits, (4) high number of oligos passing QC based on fits (see above), and (5) observation of expected affinity relationships (as assessed by ensuring that sequences with a consensus motif were bound more tightly than sequences without a motif). After selecting a reference experiment, we normalized all other experiments for a given library to this reference via a linear transformation (setting the slope to 1 and intercept to 0). This normalization procedure is diagrammed in Fig. S10.

#### 1.9 Comparison of observed binding energies with predicted DNA shape parameters

We predicted DNA shape parameters (minor groove width, helical twist, propeller twist, roll, and electrostatic potential) for all sequences in DNA Library 2 using DNAshapeR [13]. The predicted shape parameters for either the 5 bp or 20 bp sequences surrounding the extended consensus site (`GTCACGTGAC`) of 11 oligonucleotides are shown in the left-hand column of Figs. S36 and S37. We then compared measured energy ($\Delta\Delta G$) with the mean value for each of predicted shape parameters for the 5 and 20 bp on either side of the extended consensus site for each of the 11 sequences (Figs. S36 and S37).

#### 1.10 DNA circular dichroism measurements

The following DNA sequences were synthesized by Integrated DNA Technologies (IDT):

| **Name** | **Sequence** |
|---|---|
| Motif + CG/AT repeat 1 | CGCGCGCGCGCGCAGAGTCACGTGAC TCTATATATATATGTCCGGCGGTATGAC |
| Motif + CG/AT repeat 1 reverse complement | GTCATACCGCCGGACATATATATATAGAGTCAC GTGACTCTGCGCGCGCGCGCG |
| Motif + random 1 | CGCACAGTCACTTAACGTCACGTGAC CGGGGTATTTCAGGTCCGGCGGTATGAC |
| Motif + random 1 reverse complement | GTCATACCGCCGGACCTGAAATACCCCGGTCACGTGAC GTTAAGTGACTGTGCG |
| GC hairpin | GCGCGCGCGCGCGCGCGCGAAAAAA CGCGCCGCGCGCGCGCGCGGTCCGGCGGTATGAC |

Complementary strands were mixed in equimolar amounts to a final concentration of 1 mg/mL in TE buffer (pH 8.0). To anneal strands, the mixture was heated to 94°C for 3 minutes and ramped down to 37°C at a rate of -1°C/minute. The GC hairpin sequence was solubilized at 1 mg/mL in TE buffer and annealed to a hairpin via the same protocol. The annealed DNA was then dialyzed into one of the following buffers:

- 1 M $MgCl_2$ (all GC hairpin samples were measured in this buffer)
- 100 mM $CaCl_2$
- 200 mM $CaCl_2$
- 1x TE (10 mM Tris-HCl, 1 mM EDTA, pH 8.0) + 5 $\mu$M spermine
- 1x TE (10 mM Tris-HCl, 1 mM EDTA, pH 8.0) + 50 $\mu$M spermine

The buffer conditions above were chosen based on previous observations that poly(dG-dC) tracts can spontaneously form Z-DNA in the presence of high concentrations of polyamines or divalent cations (see [25] and [43]).

Circular dichroism (CD) spectra were obtained using a Jasco J-815 spectropolarimeter and 10 mm cuvettes. Each measurement was performed on 1.5 mL of sample. Spectra were collected at a constant temperature of 20°C with a scanning speed of 100 nm/min and a data pitch of 1 nm.

#### 1.11 Electrophoretic mobility shift assays (EMSAs)

We expressed MAX-eGFP with PURExpress (NEB E6800L) by first adding 15 $\mu$L of part B to 20 $\mu$L of part A. We then allowed these components to sit on ice for 30–45 minutes. We next added 500 ng of MAX-eGFP plasmid, 0.5 $\mu$L of rRNAsin recombinant ribonuclease inhibitor (Promega N2515), and MilliQ water to 50 $\mu$L. The reaction was incubated at 30°C for 3 hours for expression and then 90 minutes at 23°C to promote fluorophore maturation.

Protein yields were assessed by measuring fluorescence intensity with the DeNovix basic fluorimeter. Briefly, we diluted 2 $\mu$L of the expression mixture into a total volume of 200 $\mu$L 1x PBS. The measured fluorescence intensity of a PBS blank was then subtracted from the fluorescence intensity of the diluted protein sample and used to calculate molar concentration as follows:

$$\Delta F = fluorescence_{sample} - fluorescence_{blank} \tag{9}$$

$$\text{molar concentration (nM)} = \Delta F \cdot \frac{\text{dilution factor}}{\text{conversion factor}} = \Delta F \cdot \frac{100}{50} = 2 \cdot \Delta F \tag{10}$$

where the conversion factor was obtained from a standard curve of known concentrations of eGFP.

Single-stranded DNA oligonucleotides were synthesized by IDT and extended with a custom Alexa-647 labeled primer with a Klenow exo- polymerase as described in Section 1.2.1.

We prepared each 15 $\mu$L binding reaction with the following conditions and allowed them to equilibrate for 60–90 minutes prior to gel electrophoresis:

- 10 mM Tris-HCl pH 7.5
- 150 mM NaCl
- 1 mM DTT
- 33 ng/$\mu$L BSA
- 0.7 ng/$\mu$L poly(dI-dC)
- 1.33 nM fluorescently-labeled DNA
- variable [MAX-eGFP]

MAX-eGFP concentrations within binding reactions ranged from ∼1–80 nM, in two-fold dilutions, depending on expression yield. For each assay, we also set up a reaction without MAX-eGFP as a negative control.

EMSAs were run on pre-cast 10% TBE Novex gels (Thermo Fisher, EC62752BOX) with 1x TBE running buffer in a 4°C room. Before adding binding reactions, we pre-ran gels for 20–30 minutes to flush residual polymerization reagents from the gel matrix. Samples were then added to the gel while running. The gel was then run for 90–120 minutes and imaged on the Typhoon FLA 9500 (GE Healthcare) with the LPR-Ch2 filter cube (665nm long pass filter; GE Healthcare); gel band intensities were quantified using FIJI [40].

#### 1.12 MITOMI comparison to universal protein-binding microarray data

##### 1.12.1 Universal protein-binding microarray data

Universal protein-binding microarray (uPBM) data for Pho4 (*S. cerevisiae*) and Max (*M. musculus*) were obtained from [49] and [8], respectively. Both data sets were downloaded from CIS-BP [46], including contiguous 8-mer intensities and Z-scores. As detailed in [46], CIS-BP provides 8-mer intensities for all non-redundant 8-mers (*i.e.* combining reverse complementary 8-mers), calculated by taking the median protein fluorescence intensity for all probes containing a given 8-mer. Protein fluorescence intensities were normalized by DNA fluorescence to account for variability in probe surface density and in duplexing efficiency [11]. As detailed in [46], Z-scores were calculated for each 8-mer by taking the number of standard deviations above the median 8-mer intensity:

$$Z_i = \frac{x_i - \widetilde{x}}{\sigma}, \tag{11}$$

where $\widetilde{x}$ is the median 8-mer intensity and $x_i$ is the intensity for 8-mer $i$. We used mean Z-scores across experimental replicates for all calculations.

###### 1.12.2 Partition function binding prediction

For a given sequence, we calculated a statistical mechanics-based binding score using the following equation:

$$\Delta G_{pred} = \beta \log(Z), \tag{12}$$

where $Z$ is the partition function and $\beta = 1/k_B T$, where $k_B$ is Boltzmann's constant and $T$ is temperature of the binding reaction in Kelvin. $Z$ can then be calculated from the energy of microstates in the system:

$$Z = \sum_i e^{-\beta E_i}, \tag{13}$$

where $E_i$ is the energy of microstate $i$. We assume that a microstate in our system is the energy of a protein bound to a specific $k$-mer. We omit the energy of all unbound states $\Delta G_{unbound}$ in our calculations, as we assume that this quantity is the same for a given protein across all sequences.

We next assume that the energy of a microstate $i$ is related to the negative log of the $k$-mer binding intensity provided by a uPBM experiment. We can rewrite our binding score:

$$\Delta\Delta G_{pred} \propto \beta \log\left(\sum_j e^{\beta \log(I_j)}\right), \tag{14}$$

where $I_j$ is the binding intensity for $k$-mer $j$.

To apply this calculation to Pho4 and MAX in Fig. 2G, we considered binding to each possible 8-mer in a sequence as a separate microstate. For example, macroscopic binding to the following 12 bp sequence would be broken into 5 binding microstates:

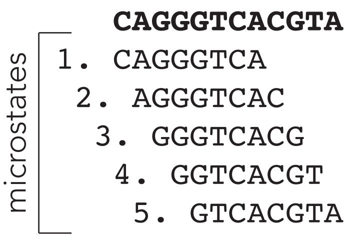

Here, each binding microstate has an energy linearly related to the logarithm of the 8-mer binding intensity ($\log(I_j)$) from uPBM measurements (as described above in 1.12.1) and used in Eqn. 14).

Code to reproduce these calculations is provided in a Github repository (https://github.com/FordyceLab/STR_analysis) and predicted binding scores for all sequences used in this work are available in the OSF repository (https://osf.io/gbxhz/).

##### 1.12.3 Calibration of predicted $\Delta\Delta G$

Partition function-based predictions of binding (described in Eqn. 14 and calculated from uPBM 8-mer intensities) are linearly related to measured $\Delta\Delta G$ (Fig. S45). We performed a linear regression on predicted and observed $\Delta\Delta G$ values in DNA Library 2 and used the regression parameters to calibrate additional partition function-based predictions (as in Fig. S46 , S48 and Sections 1.13 and 1.14).

##### 1.12.4 Estimating the number of measurements needed to calibrate predicted $\Delta\Delta G$.

In Fig. S45, we used measurements for 32 oligos to calibrate partition function-derived predictions to energetic quantities (kcal/mol). To estimate the minimum number of measurements required to reliably calibrate a binding model, we: (1) sampled $n$ measurements from the 32-member library, (2) regressed the $n$ measurements against the partition function-derived predictions to determine calibration parameters, (3) applied this calibration (a linear transformation from the regression of $n$ items) to the full library, and then (4) calculated the rmse for the measured vs. predicted $\Delta\Delta G$ for the full library using the calibration determined from $n$ measurements. We applied this sampling and fitting procedure over 10,000 iterations to capture a broad range of sampling outcomes. We set our rmse threshold to 0.5 kcal/mol, meaning that binding could be predicted within a range of approximately 2-fold for any arbitrary sequence. The results of these simulations are shown in Fig. S47.

#### 1.13 Entropy simulations

To estimate the effect of STRs on the entropy of binding (regardless of intrinsic TF-STR binding energy), we applied a Monte Carlo simulation as follows (Fig. S48):

1. Define distribution, $X$, of $k$-mer binding intensities.
2. Simulate binding entropy to 10,000 random sequences. For each sequence:
    - Sample 100 $k$-mer binding intensities from distribution $X$ with replacement to populate array of 100 microstate energies
    - Calculate $T\Delta S$ from array of microstate energies
3. Simulate binding entropy to 10,000 repetitive sequences with repeat unit length, $\lambda$. For each sequence,
    - Sample $\lambda$ $k$-mer binding intensities from distribution $X$ with replacement
    - Replicate $\lambda$ binding intensities to populate array of 100 microstate energies
    - Calculate $T\Delta S$ from array of microstate energies

4. Normalize $T\Delta S$ values to median $T\Delta S$ of random sequences

Code to reproduce simulation results is provided at https://github.com/FordyceLab/STR_analysis. We also include an analytical treatment of the entropic contribution of STRs in Section 2.1.

###### 1.13.1 Simulation of random sequences

To simulate a random sequence with 100 binding microstates, we select 100 observations from the distribution of Pho4 or MAX 8-mer binding intensities. Sampling is performed with the following Python command: `numpy.random.choice(X, size=(N,seq_len))`, where `N` is the number of sequences simulated (10,000) and `seq_len` is the number of microstates per sequence (100); `random.choice` defaults to sampling with replacement.

###### 1.13.2 Simulation of repetitive sequences

To simulate a repetitive sequence with 100 binding microstates and a repeat unit of length $\lambda$, we select $\lambda$ observations from the distribution of Pho4 or MAX 8-mer binding intensities. Sampling is performed with the following Python command: `numpy.random.choice(X, size=(N,lambda))`, where `N` is the number of sequences simulated (10,000) and `lambda` is the repeat unit size. To create an array with N microstates from an array of $\lambda$ observations, we then replicate the array of length $\lambda$ until it reaches length $N$. When $N/\lambda$ is a non-integer, we use the first $n$ elements of the array (where $n = N \,\%\, \lambda$ and % is the modulo operator).

For the simulation in Fig. S48, we use $\lambda = 1, 2, 3, 4, 5, \text{or}\, 6$, as STRs are defined as having repeated units of 1–6 nucleotides.

###### 1.13.3 Calculation and normalization of $T\Delta S$

The entropy of a system, $S$, is defined as follows:

$$S = k_B \sum_i \left(p_i \log(p_i)\right), \tag{15}$$

where $p_i$ is defined as the probability of occupying microstate $i$:

$$p_i = \frac{e^{-\beta E_i}}{\sum_j \left(e^{-\beta E_j}\right)} = \frac{e^{-\beta E_i}}{Z}, \tag{16}$$

where $Z = \sum_j \left(e^{-\beta E_j}\right)$ is the partition function and $E_i$ is the energy of microstate $i$. $\beta = 1/k_B T$, where $k_B$ is Boltzmann's constant and $T$ is temperature of the binding reaction in Kelvin. Given the relationship between $E_i$ and uPBM $k$-mer binding intensities ($I_j$) described in Eqn. 14 and shown in Fig. S48, we rewrite Eqn. 16 as:

$$p_i = \frac{e^{\beta \log(I_i)}}{\sum_j \left(e^{\beta \log(I_j)}\right)} = \frac{e^{\beta \log(I_i)}}{Z} \tag{17}$$

Simulated binding predictions were again converted to energetic quantities (kcal/mol) using the linear transformation derived for calibration (described above in 1.12.3 and shown in Fig. S45). We calculate a $T\Delta S$ value by subtracting all computed $T \cdot S$ values from the median $T \cdot S$ value of 10,000 simulated random sequences. We report a $\Delta S$ value rather than an absolute $S$ value, because we assume that entropic contributions from the unbound state, protein conformation, etc. are similar for all DNA sequences. Most importantly, this simplification allows direct comparison of configurational entropy for different DNA sequences.

#### 1.14 Gibbs free energy simulations

We next apply a Monte Carlo simulation to estimate the effect of intrinsic TF-STR binding affinity on overall TF-DNA binding energy (Fig. S48). The simulation procedure, below, is similar to that described above in Section 1.13:

1. Define distribution, $X$, of $k$-mer binding intensities
   - Calculate Z-scores for all $k$-mers in distribution
2. Simulate binding to 10,000 repeat sequences with repeat unit length, $\lambda$ in a defined Z-score range. For each sequence,
   - Sample $\lambda$ $k$-mer binding intensities from distribution $X$ with replacement. Sample only within defined Z-score range.
   - Replicate $\lambda$ binding intensities to populate array of 100 microstate energies
   - Calculate $\Delta H$, $T\Delta S$, and $\Delta G$ from array of microstate energies
3. Normalize $\Delta H$, $T\Delta S$, and $\Delta G$ values

##### 1.14.1 Simulation of repetitive sequences in Z-score range

To investigate the role of intrinsic TF-STR sequence preference, we simulate binding to sequences with the same repeat unit ($\lambda$) sampled from different regions of the same microstate energy distribution. We first define a subset of the distribution of Pho4 or MAX 8-mer binding intensities, constrained by upper and lower Z-score bounds. We select $\lambda$ observations from this subset distribution with the following Python command: `numpy.random.choice(subset, size=(N,lambda))`, where `N` is the number of sequences simulated (10,000) and `lambda` is the repeat unit size. Similar to entropy simulations described above (Section 1.13), we replicate the array of length $\lambda$ until it reaches length $N$ to create an array with N microstates from an array of $\lambda$ observations. When $N/\lambda$ is a non-integer, we use the first $n$ elements of the array (where $n = N \,\%\, \lambda$ and % is the modulo operator). This sampling procedure is repeated for all Z-score ranges to be interrogated.

##### 1.14.2 Calculation and normalization of $\Delta H$, $T\Delta S$, and $\Delta G$

The enthalpy of a binding reaction, $\Delta H$, is defined as follows:

$$\Delta H = \langle E \rangle, \tag{18}$$

where $\langle E \rangle$ is the mean microstate energy. As we are using log median intensity values in lieu of microstate energies, we instead calculate:

$$\Delta H \propto \langle -\log(I) \rangle, \tag{19}$$

$T\Delta S$ is calculated as in Section 1.13.3.

$\Delta G$ is calculated as:

$$\Delta G = \Delta H - T\Delta S \tag{20}$$

or as in Eqns. 12–14. These methods are equivalent.

Simulated binding predictions were calibrated to energetic quantities (kcal/mol) with the linear transformation as described above (Section 1.13). Given Eqn. 20, the linear transformation applied for calibration of $\Delta G$ values is also valid for $\Delta H$ and $T\Delta S$. $T\Delta S$ values are normalized by subtracting all computed $T \cdot S$ values from the median $T \cdot S$ value of 10,000 simulated random sequences. $\Delta H$ values are normalized by subtracting the median $\Delta H$ value of 10,000 simulated repetitive sequences selected from the $Z < 0$ range of the distribution. $\Delta G$ values are normalized by consequence of having normalized $T\Delta S$ and $\Delta H$.

#### 1.15 STAMMP Experiments

##### 1.15.1 Generation of Pho4 WT construct for expression via PURExpress

STAMMP experiments used Pho4 mutants expressed via *in vitro* transcription translation in a recombinantly expressed and purified cell-free expression mix (NEB PURExpress). The plasmid backbone (from the PURExpress control expression plasmid) contains an ampicillin resistance cassette, T7 promoter and terminator, and a Shine-Dalgarno sequence; the Pho4 construct was inserted into this plasmid via Gibson assembly (as described in Section 1.3.1 and in [2]).

Plasmids coding for single residue mutants of Pho4 were generated by QuikChange mutagenesis as described in [2]. Briefly, we designed primers for QuikChange mutagensis with a custom script (https://github.com/FordyceLab/designQuickChangePrimers). Candidate primers generated by the script were scored according to manufacturer guidelines. Primers selected from the pool of candidates were ordered from IDT with forward and reverse primers for each mutant in the same well at 6 nmol normalized yield, shipped dry. We used the PURExpress Pho4-eGFP plasmid described above as the template for mutagenesis, and mutagenized the library to yield the Pho4 mutants listed in Table S4 via QuickChange mutagenesis (Agilent).

For each mutagenesis reaction, we prepared the following mix:

- 14 $\mu$L MilliQ water
- 2.5 $\mu$L 10X Pfu buffer (Agilent)
- 0.25 $\mu$L Pho4 plasmid template (100 ng/$\mu$L)
- 1.25 $\mu$L DMSO (5% v/v)
- 0.5 $\mu$L dNTPs (final concentration: 200 $\mu$M)
- 0.5 $\mu$L Pfu turbo polymerase (Agilent) (0.05 Units)
- 6 $\mu$L primers, mixed forward and reverse (300 nM)

We then performed PCR as per manufacturer's guidelines (Agilent QuikChange Manual). We subsequently treated each reaction with DpnI (NEB, R0176S) for 3 hours at 37°C to digest wild-type template plasmids, followed by a 20-minute enzyme inactivation at 80°C. We used 1 $\mu$L of each reaction to transform 5 $\mu$L of E. coli DH5$\alpha$ cells (New England Biolabs, C2987I). To transform cells, we kept the cell and plasmid mixture on ice for 30 minutes, heat-shocked it at 42°C for 30 seconds, and recovered it for 1 hour in 300 $\mu$L of SOC medium (NEB, B9020S). We then plated cells on LB+ampicillin plated and allowed them to grow overnight at 37°C. Colonies were picked, miniprepped (Qiagen), and verified with Sanger sequencing. Full details can be found in the supplementary information of [2], as the same mutant library was used for this work.

##### 1.15.2 STAMMP plasmid library printing

Plasmid arrays for STAMMP (Simultaneous Transcription Factor Affinity Measurements via Microfluidic Protein Arrays) were printed as described in [2]. Briefly, 10 $\mu$L of each plasmid preparation was transferred to one of two 384-well plates with a Biomek FX Automated Workstation (Beckman Coulter, A31843). The liquid in all wells was then evaporated and plasmids were resolubilized in 12 uL of the following print solution prepared in MilliQ and then filter-sterilized:

- 1% (w/v) bovine serum albumin (Sigma Life Science, B4287-25G)
- 200 mM NaCl (Sigma Life Science, 71376-1KG)
- 12 mg/mL trehalose dihydrate (Sigma Life Science, T9531-25G)

Plasmid arrays were deposited onto 2"x3" epoxysilane-coated slides as described in Section 1.2.4. Microfluidic devices were also aligned and bonded as described above.

##### 1.15.3 STAMMP oligonucleotide preparation

Fluorescently-labeled dsDNA oligonucleotides were prepared for STAMMP binding assay by annealing and extending a fluorescently-labeled primer using the same protocol described above in Section 1.2.1. Following extension, the Klenow reaction product was passed through a 0.45 $\mu$m spin filter column (Merck Millipore, UFC30HVNB) in a tabletop microcentrifuge for 5 minutes at 5,000 rpm to remove any precipitate from the Klenow polymerase stock. Finally, the buffer of the DNA preparation was exchanged to match the "assay buffer" (10 mM Tris-HCl pH 7.5, 100 mM NaCl, 1 mM DTT) by diluting to a volume of 300 $\mu$L in assay buffer. The solution was iteratively passed through a 10 kDa molecular weight cutoff filter (Amicon Ultra, UFC501096) via centrifugation at 8,000 rpm for 10 minutes and then re-diluted to 300 $\mu$L in assay buffer for a total of five centrifugation steps. The flow-through was discarded after each centrifugation step. Following the fifth centrifugation, the remaining solution was eluted by inverting the column and centrifuging the column and its contents into a new tube for 4 minutes at 3000 rpm. A dilution series was prepared from this eluted DNA solution to yield the following approximate final concentrations: 5 $\mu$M, 2.5 $\mu$M, 625 nM, 315 nM, 155 nM, and 80 nM. To quantify DNA concentrations more precisely, we quantified absorbance at 260 nm with the DeNovix instrument to quantify DNA concentrations more precisely and calculated molar concentration (in units of nanomolar) from A260 as follows:

$$\text{molar concentration (nM)} = \frac{(A_{260} \cdot 50) \cdot 10^6}{618 \cdot d + 36} \tag{21}$$

where $d$ is the length of dsDNA in base pairs.

##### 1.15.4 STAMMP on-chip *in vitro* transcription, translation, and purification

For STAMMP assays, all TF-eGFP fusions were expressed on the device using the PURExpress in vitro transcription and translation system (New England Biolabs, E6800L). For each device, we added 7.5 $\mu$L of PURExpress component B to 10 $\mu$L of component A and mixed gently by pipetting. The mixture was then kept on ice for 30–45 minutes, as this was found to increase expression yield.

After the 30–45 minutes, 1.5 $\mu$L of RNAsin (Promega, N2515) and MilliQ water were added to a final volume of 25 $\mu$L. This reaction was scaled to the number of devices, as each 25 $\mu$L preparation is sufficient for one device.

For STAMMP assays involving on-chip expression, the control lines operating button, sandwich, and neck valves were pressurized with 0.55 M NaCl to prevent the hyperosmotic plasmid spot from absorbing water. Following surface patterning (performed as described for MITOMI experiments in Section 1.5.1), we flowed the PURExpress mix through the device for 10 minutes with neck and button valves closed. We then closed the outlet valve and opened the neck valves and flowed PURExpress for an additional 2–3 minutes to ensure that DNA chambers were filled. Subsequently, we closed sandwich valves (to isolate adjacent chambers) and opened neck valves (to allow expression to take place across the entire chamber). Devices were then transferred to a hot plate at 37°C for 45 minutes for protein expression. To promote proper protein folding and maturation, devices were removed from the hot plate and allowed to incubate for an additional 2 hours at room temperature.

After the room temperature-incubation, button valves were opened to expose the antibody-patterned surface to the TF-eGFP fusions. The binding of TFs to the antibody surface was allowed to proceed for 2 hours, during which time periodic images of the device were taken to monitor fluorescence buildup.

##### 1.15.5 STAMMP equilibrium binding measurements

Following on-chip expression, purification, and trypsin wash, we substituted the PBS line with "assay buffer", made in MilliQ water with the following final concentrations:

- 10 mM Tris-HCl pH 7.5 (from 1 M stock; Fisher Bioreagents, BP1757-500)
- 100 mM NaCl (Sigma-Aldrich, 71376-1KG)
- 1 mM DTT (Sigma-Aldrich, D9779)

We next "equilibrated" the device and the surface-immobilized transcription factors in the new buffer by flowing the buffer for 10 minutes and then opening button valves for 50 minutes. Finally, we measured binding of all TF variants on the device to a single sequence per device at six different concentrations, prepared as described above in Section 1.15.3. For each concentration, we (1) flowed each through the device for 10 minutes (with button valves closed), (2) stopped flow by closing sandwich and outlet valves, (3) opened button valves for 50 minutes to allow transcription factor-DNA binding, (4) took images of the device to quantify the DNA concentration available for binding ("pre-wash images"), (5) closed button valves to trap interactions, (6) washed away unbound oligonucleotides with assay buffer, and then (7) imaged the device again ("post-wash images") to quantify amounts of immobilized transcription factor (GFP channel) and bound DNA (Cy5 channel). When processing data for STAMMP assays, the same chamber coordinates were used for all 6 concentrations.

##### 1.15.6 STAMMP data normalization and culling

STAMMP affinity measurements were culled by rmse as described in Section 1.8. In addition, because multiple $K_d$ measurements for a given protein-DNA sequence pair were collected on a single device, we also calculated the CV (coefficient of variation) of $K_d$ values for each protein-DNA pair. We then eliminated any protein-DNA pairs from a given experiment with a CV greater than 1.5 standard deviations above the mean CV.

STAMMP affinity data were also normalized largely as described in Section 1.8. A reference experiment for each oligonucleotide was chosen based on (1) average rmse across all curve fits, (2) number of chambers in a MITOMI device remaining after culling, (3) number of mutants passing QC, and (4) evenness of protein deposition across the device.

#### 1.16 k-MITOMI dissociation rate measurements

##### 1.16.1 k-MITOMI experimental procedure

All k-MITOMI experiments used the same device and initially followed the same protocol described for MITOMI equilibrium binding assays (see Section 1.5). To quantify dissociation rates after binding, we iteratively allowed fluorescently-labeled bound DNA to dissociate for 1–4 seconds, mechanically trapped remaining bound DNA, washed devices, and imaged. The inclusion of non-fluorescent (dark) competitor at high concentrations during dissociation is critical to prevent rebinding of labeled material, which leads to systematic underestimation of dissociation rates.

To acquire dissociation rate data after equilibrium binding, we first flushed each device with non-fluorescent (dark) competitor dsDNA oligonucleotides containing an E-box motif at a concentration of 2 $\mu$M diluted in PBS for 10 minutes with button valves closed after the acquisition of the "post-wash" MITOMI image. These oligos were prepared with Klenow polymerase largely as described above in Section 1.2.1 but with unlabeled primers.

After stopping flow of unlabeled competitor dsDNA and closing sandwich valves, we then opened the buttons for 1.2, 2.0, or 4.0 seconds to allow dissociation of bound fluorescent DNA from surface-immobilized TF. Next, we closed buttons, flushed the device, and imaged in both the Cy5 and eGFP channels to quantify loss of DNA binding and surface-bound TF, respectively. For each experiment, we iterated this process for 30, 20, or 10 iterations for button duty cycles of 1.2, 2.0, or 4.0 seconds, respectively.

##### 1.16.2 k-MITOMI image acquisition and processing

For dissociation rate measurements, image stitching yields a Cy5 and eGFP image of the entire device for each timepoint (*i.e.* for each button opening and closing cycle) indicating bound DNA concentration and immobilized TF-eGFP, as some protein loss is inevitable during the experiment. As for MITOMI experiments, button centroid coordinates acquired from TF eGFP "post-wash" images were used to compute DNA intensities at each button for all time points.

##### 1.16.3 Determination of kinetic constants

As for MITOMI experiments, the ratio (R) of "post-wash" DNA fluorescence (Alexa 647 or Atto 647N) to "post-wash" GFP fluorescence per chamber provides a proxy for the number of fluorescently-labeled DNA molecules bound to surface-immobilized TFs. To quantify labeled DNA dissociation over time, we calculated this ratio for each chamber of the device at each of the time points collected (*i.e.* after each pulse of the button valves).

As noted above (Section 1.16.1), the addition of dark competitor allows the assumption of negligible rebinding:

$$\frac{\mathrm{d}[TF \cdot DNA]}{\mathrm{d}t} = -k_{off}[TF \cdot DNA] + k_{on}[TF][DNA] \rightarrow -k_{off}[TF \cdot DNA] \tag{22}$$

This differential equation can be solved analytically, yielding a single exponential to which data were fit:

$$R(t) = R(0) \cdot e^{-kt} + c \tag{23}$$

Here, $R(t)$ is the fluorescence ratio as a function of time, $k$ is the dissociation constant, and $c$ is a constant term which accounts for background fluorescence or non-specific sticking of DNA. Dissociation curves were fit only to the highest concentration chambers (*i.e.* those corresponding to the highest 2 or 3 concentrations in the dilution series), as these curves started with higher R values and thus had more time points before decaying to the noise floor. Furthermore, only the first 16 seconds of the binding curve were used to fit a dissociation rate as this was found to improve consistency across different button pulses.

##### 1.16.4 Data culling and normalization for k-MITOMI kinetic measurements

Kinetic data were culled based on the rmse of each curve fit ($> 2\sigma$ above the mean rmse across all curve fits) and CV of all fits across a device ($> 2\sigma$ above the mean across all experiments). In addition, dissociation curves were culled for unusually low or high asymptote values (outside $2\sigma$).

Similar to above-described normalizations for MITOMI and STAMMP experiments, kinetic measurements were normalized relative to a reference experiment. A reference experiment was chosen based on the following criteria: (1) low average rmse across all curve fits, (2) high number of chambers in a MITOMI device remaining after culling, (3) high number of oligos passing QC, and (4) low CV of $k_{off}$ values for a given oligo within an experiment. For normalization, $k_{off}$ values in a given experiment were linearly transformed such that the median $k_{off}$ value for that experiment matched the median $k_{off}$ of the reference experiment. This normalization procedure is diagrammed in Fig. S10.

##### 1.16.5 Alternative strategy for k-MITOMI measurement normalization

A previous study employing k-MITOMI observed a relationship between button valve pulse times and measured $k_{off}$ values [20]. We observed no relationship between button pulse time and measured $k_{off}$ values for DNA Library 1 (see Figs. S74, S76). However, we observed a linear relationship between button pulse time and measured $k_{off}$ for DNA Library 2 (Figs. S68, S69). To correct for this dependency on button pulse time, imposed by the mechanical shear of valve opening, we performed a linear regression of $k_{off}$ *vs.* 1/button pulse, where 1 second was subtracted from the button pulse time to account for the delay in button opening (see below). The $y$-intercept of the linear fit corresponds to an "infinite" button pulse, without mechanical shear from valve actuation. The results of these linear regressions and the computed $k_{off}$ and $k_{on}$ values are in Figs. S68–S70. We observed strong agreement between $k_{on}$ values determined with this correction method and those described above in Section 1.16.4 (see Fig. S71).

**Measuring button actuation time.** To determine the time for button valves to open fully, we first flowed fluorescent dye onto a BSA-passivated MITOMI device, with button valves closed. We then opened button valves and continuously imaged 16 button valves at 1000 frames per second and monitored buildup of fluorescent signal as a proxy for button opening. We observed that buttons took approximately 1 second to fully open, as shown in Fig. S67.

##### 1.16.6 Calculating inferred $k_{on}$ values from MITOMI and k-MITOMI measurements

The following chemical equation describes a general macroscopic TF-DNA binding reaction:

$$TF + DNA \rightleftharpoons TF \cdot DNA, \tag{24}$$

We can model the rate change of species in this reaction as:

$$\frac{\mathrm{d}[TF]}{\mathrm{d}t} = \frac{\mathrm{d}[DNA]}{\mathrm{d}t} = -\frac{\mathrm{d}[TF \cdot DNA]}{\mathrm{d}t} = k_{off}[TF \cdot DNA] - k_{on}[TF][DNA], \tag{25}$$

where the $k_{on}$ is the forward rate constant and $k_{off}$ is the reverse rate constant in chemical equation 24. At equilibrium, there is no net change in concentrations of species:

$$\frac{\mathrm{d}[TF]}{\mathrm{d}t} = \frac{\mathrm{d}[DNA]}{\mathrm{d}t} = \frac{\mathrm{d}[TF \cdot DNA]}{\mathrm{d}t} = 0, \tag{26}$$

which permits us to rewrite Eqn. 25:

$$\frac{k_{off}}{k_{on}} = \frac{[TF][DNA]}{[TF \cdot DNA]} = K_d \tag{27}$$

As we have directly measured both $K_d$ and $k_{off}$, we can infer $k_{on}$:

$$k_{on} = \frac{k_{off}}{K_d} \tag{28}$$

Normalized $k_{off}$ values (see Section 1.16.4) are used in Eqn. 28. We applied no further normalization to reported $k_{on}$ values.

#### 1.17 Four-state model and kinetic simulations

##### 1.17.1 Four-state model derivation

The model considered is the one drawn in Fig. 4D of the main-text, where the free, unbound state has state index 1, the testing state has state index 2, the motif-bound state has state index 3, and the flank-bound state has state index 4. Formulas for model $K_{d,model}$ and $k_{off,model}$ were obtained by deriving the relevant mean first passage times of the model, with a similar methodology as in [33]. $K_{d,model}$ and $k_{off,model}$ can be expressed in first mean first passage times as

$$K_{d,model} = \frac{k_{off,initial}}{k_{on,initial}} = \frac{t_{on,initial}[DNA]}{t_{off,initial}} \tag{29}$$

and

$$k_{off,model} = k_{off,equilibrium} = \frac{1}{t_{off,equilibrium}}, \tag{30}$$

where $t_{off,initial}$ is the mean time for the model to dissociate, *i.e.* to go from one of the protein-DNA bounds states (state 2, 3 or 4) to the free state (state 1), after just having entered a protein-DNA bound state. Correspondingly, $t_{off,equilibrium}$ is the mean time for the model to dissociate after it has reached steady-state in an association process. That $k_{off,model} = k_{off,equilibrium}$ is a consequence of how $k_{off,experiment}$ is measured and estimated. $\frac{1}{k_{off,experiment}}$ is an estimate of the mean time for dissociation after having reached equilibrium in an association process, which is reflected in the $k_{off,model}$ that tries to model this quantity. We note, however, that $k_{off,initial}$ and $k_{off,equilibrium}$ should not be interpreted as true rate constants (*i.e.* exponential factors in some single exponential decay, but rather as inverses of average times for dissociation). In a general scenario, dissociation out of a group of aggregated states in a Markov model (in this case state 2, 3 and 4) is a sum of multiple exponentials. One single exponential cannot describe the process, but it is still possible to define a mean time for dissociation, which is what achieve in the following derivation.

The mean first passage time $t_{i,j}$ for a continuous time Markov chain to go from state $i$ to state $j$ can be calculated according to

$$t_{i,j} = t_{i,k \neq i} + \sum_{k \neq j} p_{ik} t_{k,j}, \tag{31}$$

where $t_{i,k \neq i}$ is the mean time for the model to leave state $i$ for any of the other states, and $p_{i,k}$ is the probability for the model to transition to state $k$ given that it is currently in state $i$. The probability $p_{i,k}$ to transition from the testing state (state 2) to the motif (state 3) or the flank (state 4) can in turn be expressed in terms of the relevant $f$ parameters, which for the transition to the motif is

$$p_{2,3} = \frac{f_{2,3}}{1 + f_{2,3} + f_{2,4}} = \frac{f_{motif}}{1 + f_{motif} + f_{flank}}. \tag{32}$$

The re-parameterization (or re-normalization) of $p_{i,k}$ to $f$ in the model fitting is made to avoid doing conversions between different $p_{i,k}$ for different model architectures (with or without flanks for example) at every step in the optimization algorithm. $p_{i,k}$ will be different and have to be converted for different model architectures, but $f$ type parameters do not need to be converted when the model architecture changes. However, in the derivation of the mean first passage times, $p_{i,k}$ is the more natural parameter to consider.

We first consider the association process, i.e. the transitions towards state 2, 3 and 4, to obtain an expression giving $t_{on,initial}$. For the association process we can lump states 3 and 4 together, and only consider the time to transition to either of these states. Evaluating Eqn. 31 then gives the linear system of equations

$$\begin{pmatrix} -1 & 1 \\ 1 - (p_{23} + p_{24} & ) - 1 \end{pmatrix} \begin{pmatrix} t_{1,3+4} \\ t_{2,3+4} \end{pmatrix} = \begin{pmatrix} -t_{1,k \neq 1} \\ -t_{2,k \neq 2} \end{pmatrix}, \tag{33}$$

where $t_{i,3+4}$ is the mean time for the model to transition from state $i$ to state 3 or 4. After solving Eqn. 33 for $t_{1,3+4}$ we get

$$t_{1,3+4} = \frac{t_{1,k \neq 1} + t_{1,k \neq 2}}{p_{2,3} + p_{2,4}}. \tag{34}$$

We now assume that the time spent in the testing state $t_{2,k \neq 2}$ is very small compared to the time spent in any of the other states of the model, so that Eqn. 34 simplifies to

$$t_{on,initial} = t_{1,3+4} = \frac{t_{1,k \neq 1}}{p_{2,3} + p_{2,4}} = \frac{1/(k_{on,max}[DNA])}{p_{2,3} + p_{2,4}}. \tag{35}$$

We now consider the dissociation process, *i.e.* the transitions towards state 1, to obtain expressions giving $t_{off,initial}$ and $t_{off,equilibrium}$. Evaluating Eqn. 31 now gives the linear system of equations

$$\begin{pmatrix} -1 & p_{2,3} & p_{2,4} \\ 1 & -1 & 0 \\ 1 & 0 & -1 \end{pmatrix} \begin{pmatrix} t_{2,1} \\ t_{3,1} \\ t_{4,1} \end{pmatrix} = \begin{pmatrix} -t_{2,k\neq 2} \\ -t_{3,k\neq 3} \\ -t_{4,k\neq 4} \end{pmatrix}. \tag{36}$$

After again assuming that $t_{2,k\neq 2}$ is very small, Eqn. 36 has the solution

$$t_{3,1} = \frac{(1-p_{2,4})t_{3,k\neq 3} + p_{2,4}t_{4,k\neq 4}}{1-p_{2,3}-p_{2,4}} = \frac{(1-p_{2,4})\frac{1}{k_{off,\mu,motif}} + p_{2,4}\frac{1}{k_{off,\mu,flank}}}{1-p_{2,3}-p_{2,4}} \tag{37}$$

and

$$t_{4,1} = \frac{(1-p_{2,3})t_{4,k\neq 4} + p_{2,3}t_{3,k\neq 3}}{1-p_{2,3}-p_{2,4}} = \frac{(1-p_{2,3})\frac{1}{k_{off,\mu,flank}} + p_{2,4}\frac{1}{k_{off,\mu,motif}}}{1-p_{2,3}-p_{2,4}}. \tag{38}$$

The average dissociation times $t_{off,initial}$ and $t_{off,equilibrium}$ can in turn be calculated as weighted averages of $t_{3,1}$and $t_{4,1}$, where the weights are set according to the relative probabilities for the model to be in state 3 and 4 (after just having entered state 3 or 4) or at steady-state after an association process, respectively. $t_{off,initial}$ and $t_{off,equilibrium}$ can thus be calculated according to

$$t_{off,initial} = \frac{p_{2,3}}{p_{2,3}+p_{2,4}}t_{3,1} + \frac{p_{2,4}}{p_{2,3}+p_{2,4}}t_{4,1} \tag{39}$$

and

$$t_{off,equilibrium} = \frac{\Pi_3}{\Pi_3+\Pi_4}t_{3,1} + \frac{\Pi_4}{\Pi_3+\Pi_4}t_{4,1} = \frac{1}{1+\frac{p_{2,4}}{p_{2,3}}\frac{k_{off,\mu,motif}}{k_{off,\mu,flank}}}t_{3,1} + \left(1 - \frac{1}{1+\frac{p_{2,4}}{p_{2,3}}\frac{k_{off,\mu,motif}}{k_{off,\mu,flank}}}\right)t_{4,1}. \tag{40}$$

Here, $\Pi_i$ is the steady-state probability for the model to be in state $i$. Eqns. 39 and 40 were validated in an orthogonal manner, by solving the system of linear ordinary differential equations (ODEs) defining this model instead of using mean first passage times. The average dissociation time was then calculated by integrating the resulting dissociation curve (survival function). Initial conditions when solving the ODEs were chosen either so that the model had just entered state 3 or 4 (to obtain $t_{off,initial}$) or so that the model had reached steady-state after an association process (to obtain $t_{off,equilibrium}$). This was done for a few numerical examples, which all gave exactly the $t_{off,initial}$ and $t_{off,equilibrium}$ predicted by equations 39 and 40.

Inserting Eqns. 35, 37–40 into Eqns. 29 and 30 gives the final expressions

$$K_{d,model} = \Bigg( k_{on,max}(p_{2,3}+p_{2,4}) \times \Bigg( \frac{p_{2,3}}{p_{2,3}+p_{2,4}} \frac{(1-p_{2,4})\frac{1}{k_{off,\mu,motif}} + p_{2,4}\frac{1}{k_{off,\mu,flank}}}{1-p_{2,3}-p_{2,4}} + \frac{p_{2,4}}{p_{2,3}+p_{2,4}} \frac{(1-p_{2,3})\frac{1}{k_{off,\mu,flank}} + p_{2,4}\frac{1}{k_{off,\mu,motif}}}{1-p_{2,3}-p_{2,4}} \Bigg)\Bigg)^{-1}. \tag{41}$$

and

$$k_{off,model} = \Bigg( \frac{1}{1+\frac{p_{2,4}}{p_{2,3}}\frac{k_{off,\mu,motif}}{k_{off,\mu,flank}}} \frac{(1-p_{2,4})\frac{1}{k_{off,\mu,motif}} + p_{2,4}\frac{1}{k_{off,\mu,flank}}}{1-p_{2,3}-p_{2,4}} + \Bigg(1 - \frac{1}{1+\frac{p_{2,4}}{p_{2,3}}\frac{k_{off,\mu,motif}}{k_{off,\mu,flank}}}\Bigg) \frac{(1-p_{2,3})\frac{1}{k_{off,\mu,flank}} + p_{2,4}\frac{1}{k_{off,\mu,motif}}}{1-p_{2,3}-p_{2,4}} \Bigg)^{-1} \tag{42}$$

##### 1.17.2 Model fitting to estimate parameters

The dataset used for fitting the model (Fig. 4D of the main text) consists of $K_d$ and $k_{off}$ values for the consensus and mutated motif with random flanks:

- $K_d$ and $k_{off}$ values for Motif + AT/CG repeat 1 (`CGCATATATATATAGAGTCACGTGACTCTCGCGCGCGCGGTCCGGCGGTATGAC`)
- $K_d$ and $k_{off}$ values for Motif + AC/GT repeat 1 (`CGCCACACACACACATGTCACGTGACATGTGTGTGTGTGGTCCGGCGGTATGAC`)
- $K_d$ and $k_{off}$ values for the Mutated motif + AT/CG repeat 1 (`CGCATATATATATAGAGTCACGCGACTCTCGCGCGCGCGGTCCGGCGGTATGAC`)
- $K_d$ values for No motif + AT/CG repeat 3 (`CGCATATATATATATATATATCGCGCGCGCGCGCGCGCGGTCCGGCGGTATGAC`)
- $K_d$ values for No motif + AC/GT repeat 5 (`ACGCCCAACCAACACATGTCACGTGACATGTGTTGGTGTGGTCCGGCGGTATGAC`)

We assume that these repeats behave identically to the repetitive flanks in Motif + AT/CG repeat 1 and Motif + AC/GT repeat 1, respectively, and thus have $n_{motifs} = 2$ and $n_{flanks} = 2$, resulting in a total of $1+2n_{motifs}+2n_{motifs} = 9$ free parameters in the model. These free parameters can be uniquely estimated using the 12 non-redundant data points (7 $K_d$ and 5 $k_{off}$) from DNA Library 1. The parameters of the model were fitted to the experimental data by minimizing the sum squared deviation between the experimental and model-predicted $K_d$ and $k_{off}$ values:

$$s^2 = \sum \left( \frac{K_{d,model,i} - K_{d,experiment,i}}{max(K_{d,experiment})} \right)^2 + \sum \left( \frac{k_{off,model,i} - k_{off,experiment,i}}{max(k_{off,experiment})} \right)^2, \tag{43}$$

Here, we normalized $max(K_{D,experiment})$ and $max(k_{off,experiment})$ to account for the fact that the experimental $K_D$ and $k_{off}$ have different magnitudes. Eqn. 43 was minimized using the Interior Point Algorithm (`fmincon` in MATLAB), with 1000 randomized start guesses, and then taking the smallest of the 1000 obtained local minima as the global minimum. Start guesses were generated on the interval $10^{-6}$–$10^{-2} s^{-1} nM^{-1}$ for $k_{on,max}$, on the interval $10^{-3}$–$10^{3}$ for $f$ type parameters, and on the interval $10^{-3}$–$10^{3}$ $s^{-1}$ for $k_{off,\mu}$ type parameters, all using log-uniform distributions. The fit was constrained so that all parameters were positive, $k_{off,\mu}$ for the two different motifs differed at most by a factor of 10, $k_{off,\mu}$ for the two different flanks differed at most by a factor of 10, $f$ for the two different motifs differed at most by a factor of 10 and $f$ for the two different flanks differed at most by a factor of 10. Error bars for model $K_D$, $k_{off}$ and fitted parameters are 68% confidence intervals, obtained by resampling the experimental $K_D$ and $k_{off}$ 200 times and re-running the fitting algorithm on these samples. The samples of $K_D$ and $k_{off}$ were drawn from a normal distribution with the average equal to the mean of the current data point ($K_D$ or $k_{off}$) and with the standard deviation equal to the standard error of the current data point. In the fitting on the re-sampled values, we used 500 randomized start guesses for each re-sample fit. The optimization algorithm was validated to return similar fitted parameter values when it was re-run twice with different random number generator streams.

#### 1.18 Gillespie model of TF search with repetitive flanks

Gillespie algorithms are stochastic simulations based on reaction rates that use discrete molecule counts and variable time steps [21]. Here, we simulated TF search trajectories using a Gillespie model based on the four states from the kinetic model depicted in Fig. 4D. At each time step, we compute: (1) How long until the next reaction occurs? and (2) Which reaction happens? First, we calculated reaction propensities ($a$) from reaction probabilities ($c$) and the number of reactants available for each reaction. Reaction probabilities can be derived from the kinetic rate constants as follows:

- For first-order reactions, $c = k$ and $a = c * X_1$ where $X_1$ represents the number of molecules of the reactant.
- For second-order reactions, $k$ is converted from 1/M/s to 1/(molecules/L)/s and $c = k/V$, where $V$ represents the volume of the system. $a = c * X_1 * X_2$, where $X_1$ and $X_2$ represent the number of molecules of each of the two reactants.

The time step $\tau$ is calculated by drawing a random number to pull a time step from an exponential distribution with a mean of $1/a_{total}$, where $a_{total}$ is the sum of all the reaction propensities, using the following equation:

$$\tau = \frac{1}{a_{total}} \ln\left(\frac{1}{rand_1}\right)$$

Next, the reaction that occurs is decided by drawing a second random number and using the following equation, where q represents the index of the chosen reaction:

$$\sum_{j=1}^{q} a_j > a_{total} * rand_2$$

In this model, we set the volume of the yeast nucleus to 3 $\mu m^2$ [35]. We initialized with 2600 molecules of Pho4 and 1 DNA “molecule” [23]. $k_{off,M}$ was set to $10^6$. We used the microscopic rates fit for Pho4 interacting with Motif + AT/CG repeat 1 (`CGCATATATATATAGAGTCACGTGACTCTCGCGC`

GCGCGGTCCGGCGGTATGAC) and Motif + AC/GT repeat 1 (CGCCACACACACACATGTCACGTGACATGTGT GTGTGTGGTCCGGCGGTATGAC) calculated from the CTMC model (Section 1.17). For random flanks, we estimated that $f_{randomflank}$ was 20-fold lower than $f_{AC/GTflank}$. Specifically, we used:

- $k_{on,MAX} = 2.67 \cdot 10^{-4}\, nM^{-1}s^{-1}$
- $k_{off,M} = 1 \cdot 10^{6}\, s^{-1}$
- $k_{off,\mu,motif} = 0.317\, s^{-1}$
- $k_{off,\mu,flank} = 0.373\, s^{-1}$
- $f_{motif} = 0.846$
- $f_{AT/CGflank} = 0.789$
- $f_{AC/GTflank} = 0.223$
- $f_{randomflank} = 0.011$

We impose a limit of 1 TF bound to the motif and 9 TFs bound to the flanks (we note that this limit for the flanks is arbitrary, but that we do not observe more than 6 TFs bound to the flanks at a time in our simulations). Sensitivity analysis was conducted for a range of TF concentrations and $k_{on}$ values and flank affinities, with 10 simulations per parameter combination (see Figs. S85,S86). Code to reproduce these simulations is available at https://github.com/FordyceLab/STR_analysis.

#### 1.19 Affinity Distillation neural network model

##### 1.19.1 Model architecture

Our neural network model architectures are adapted from BPNet, a sequence-to-profile convolutional neural network with one-hot-encoded DNA sequence inputs and base-resolution read count profile outputs [7]. We use the output of the final convolutional layer (bottleneck activation map) as input for two output heads: (1) a deconvolutional layer for the profile prediction; and (2) a global average pooling layer followed by the fully connected layer for total read count prediction. Full details of BPNet, its implementation, and extraction of CWMs and binding profiles are publicly available [7].

The model takes 1346 bp sequences as input. The first convolutional layer uses 64 filters of width 21 bp, followed by 6 dilated convolutional layers (each with 64 filters of width 3). The deconvolutional layer has a filter width of 75 bp and the output profile length is 1000 bp. We set the count prediction head relative weight compared to the profile prediction head to 100 for all models. All convolutional and deconvolutional layers are reverse-complement layers [42].

##### 1.19.2 ChIP-seq data

MAX ChIP-seq data were obtained from the ENCODE portal with accession numbers ENCSR000EZF (experiment, HeLa-S3 cells) and ENCSR000EZM (control) [15, 16]. We processed data using the ENCODE pipeline: https://github.com/ENCODE-DCC/chip-seq-pipeline2 (v1.3.6).

##### 1.19.3 Model training

IDR peaks were divided for training, validation and testing as follows:

- **Test set:** regions from chromosomes 8 and 9 (approx. 10% of the genome), for model performance evaluation

- **Validation set:** regions from chromosomes 16, 17, and 18 (approx. 10% of the genome), for tuning hyperparameters
- **Training set:** regions from the remaining chromosomes (chromosomes 1–7, 10–15, 19–22, X; approx. 80% of the genome).

We performed five-fold cross-validation, tuning hyperparameters to optimize performance. We employed a uniform jitter during training with a maximum shift of 200 bp of the regions in each batch. Neural network models were implemented and trained in Keras (v.2.2.4) (TensorFlow backend v.1.14) [14, 1] using the Adam optimizer [26] with learning rate 0.001 and early stopping with patience of ten epochs.

###### 1.19.4 Model cross-validation

In addition to model training described above, we evaluated models with five-fold cross-validation. Each fold divides the genome into non-overlapping regions (separated by at least 2 kb between sets), assigning 10% of peaks to the test set and 10% of peaks to the validation set. The remaining 80% were used for training.

###### 1.19.5 DeepSHAP contribution scores

We used the deep explainer implementation of SHAP (DeepSHAP) [29], which is an updated version of DeepLIFT [41], for model interpretation. At each position in the input sequence, we iterate over the one-hot encoding possibilities and compute the hypothetical difference-from-reference in each case. We then multiply the hypothetical differences-from-reference with the multipliers to get the hypothetical contributions. We used a shuffled reference with 20 random shuffles. For each of the one-hot encoding possibilities, the hypothetical contributions are then summed across the ACGT axis to estimate the total hypothetical contribution of each position. This per-position hypothetical contribution is then projected onto whichever base was present in the hypothetical sequence.

###### 1.19.6 *In silico* marginalization

A detailed description of methods for *in silico* marginalization is provided at [4]. Briefly, we performed *in silico* marginalization for a sequence of interest by:

1. generating 100 background sequences by dinucleotide shuffling DNA sequences from held-out genomic peaks
2. computing a model prediction for each of the background sequences ($\log(counts_{bg})$)
3. inserting the sequence of interest at the center of the background sequences
4. computing a model prediction for the sequence of interest inserted in the background ($\log(counts_{insert})$)
5. computing the mean difference between the background sequences before and after insertion to compute the marginalization score: $\Delta \log(counts) = mean(\log(counts_{insert})) - mean(\log(counts_{bg}))$

#### 1.20 Calculation of Shannon entropy

To calculate Shannon entropy, we broke DNA sequences into overlapping 3-nucleotide "words." For example, a sequence `ATGGCGA` would be broken into the following words: `ATG`, `TGG`, `GGC`, `GCG`, and `CGA`. Following this procedure, we calculated Shannon entropy ($H$) with the following equation:

$$H = -\sum_{i=0}^{k} \left(\frac{w_i}{l}\right) \ln\left(\frac{w_i}{l}\right), \tag{44}$$

where $w_i$ is the number of instances of word $i$, $l$ is the total number of words, and $k$ is the "alphabet size," or the total number of possible words (in this case $k = 4^3 = 64$). This calculation was implemented in Python with the following code:

```python
def calculate_entropy(sequence, block=3):
    words = defaultdict(int)
    l = len(sequence) - block + 1 # total number of words
    Shannon_score = 0
    # count words
    for m in range(l):
        key = sequence[m:m+block]
        words[key] += 1
    # calculate entropy
    for word in words:
        Shannon_score -= words[word]/l * np.log(words[word]/l)
    return Shannon_score
```

#### 1.21 Analysis of uPBM data

##### 1.21.1 Data filtering and aggregation

Universal protein-binding microarray (uPBM) data were downloaded from CIS-BP with accompanying Z-scores for all contiguous 8-mers [46]. The following data were excluded from further analysis:

- Artificial or designed protein constructs (e.g. chimeric proteins)
- Mutated transcription factors (as in [9])
- Non-eukaryotic proteins (e.g. Epstein-Barr Virus)
- Methylated DNA (as in [47])

In addition, data were filtered to eliminate low signal measurements, according to the following criteria:

- **Distribution skew**. We expect an approximately Gaussian distribution of noise and a long rightward tail representing strong binding events. We therefore calculated skew for each uPBM data set and filtered out data sets with a skew in the bottom 1/3 of the skew distribution.

- **Distribution maxima**. We filtered data sets with abnormally small (below 10th %ile) or large (above 95th %ile) maximum values.
- **Data reproduciblity**. We filtered data sets where $k$-mer Z-scores across replicates had a Pearson $r < 0.616$ and Spearman $\rho < 0.446$. These thresholds represent the 20th and 30th percentiles of correlation coefficients, respectively.

Following data filtering, we computed mean 8-mer Z-scores across all measurements for a given transcription factor, regardless of whether measurements originated from different labs or publications. Data were not averaged across species, even for well-conserved TFs. 8-mer Z-scores for 1,291 TFs across 114 species were used for subsequent analyses.

We manually re-classified TFs into a smaller list of 17 domain families and grouped all other TFs into an "Other" category. For example, $C_2H_2$ zinc finger TFs and zinc knuckle (CCHC) TFs were combined into a general "zinc finger" category.

Binding intensity Z-scores for all TFs passing QC are available on OSF (https://osf.io/gbxhz/). TF family classifications are also included in this repository.

###### 1.21.2 Repeat Z-score aggregation

We aggregated Z-scores for 1,291 TFs interacting with 39 repeat types by taking the mean of all 8-mers within a repeat type. We define a repeat type to include all 8-mers, which, when repeated in tandem, would produce the same STR or its reverse complement. For example, `ACACACAC`, `GTGTGTGT`, and `CACACACA` fall within the same repeat type. Given that Z-scores are computed for 8-mer binding intensities, we consider only repeat units of 1, 2, or 4 bp. Binding intensity Z-scores for 39 repeat types for all TFs passing QC are available on OSF (https://osf.io/gbxhz/).

###### 1.21.3 Transcription factor binding motifs

Position weight matrices (PWM) for all TFs with uPBM data were downloaded from CIS-BP [46]. To define a binding motif for a given TF, we included all nucleotides at a given position which had at least 60% of the maximum binding affinity. For the example of Pho4 (Fig. 6D, bottom left panel), the binding motif would be `CACGTGSNN` where `S` is the IUPAC degenerate nucleotide for `G` or `C`.

###### 1.21.4 Calculation of motif-repeat Levenshtein distance

To calculate the Levenshtein distance (LD) between a given repeat 8-mer (*i.e.* an 8-mer which, when repeated in tandem, would form an STR with unit 1, 2, or 4 bp) and a binding motif, we implemented the following algorithm:

- **Compute the LD between each pairwise motif and repeat variation.** We used the `python-Levenshtein` package (source code: https://github.com/ztane/python-Levenshtein/) to compute Levenshtein distances.
  - **Iterate over each degenerate letter in a binding motif.** We define a list of motifs for each TF. For each position where multiple bases are tolerated, each one is included. For the example of Pho4, the binding motif is `CACGTGSNN`, where `S` is the IUPAC degenerate nucleotide for `G` or `C`, and we thereby generate a list of motifs: `CACGTGCAA`, `CACGTGGAA`, `CACGTGCCA`, and so forth.

- **Iterate over each frame and reverse complement in repeat 8-mer.** We generate a list of synonymous repeats and their reverse complements. For example, the following patterns would be synonymous: `AC`, `CA`, `GT`, and `TG`. A Pho4 motif, `CACGTGSNN`, would be only 2 edits away from a `CGCGCGCG` but 4 edits away from `GCGCGCGC` even though they represent the same pattern.

- **Return the minimum LD.**

This iterative approach avoids the use of an alignment algorithm. Code to replicate these calculations can be found at https://github.com/FordyceLab/STR_analysis.

###### 1.21.5 Estimation of "significant" binding above background

To determine whether a given repeat is bound by a given TF "significantly" above background, we first define a background distribution of binding. For each TF, we fit a Gaussian to the negative portion of the Z-score distribution, reflected over $x = 0$. We then define "significantly" bound 8-mers as all 8-mers with a Z-score greater than a predetermined quantile in the Gaussian fit of the background distribution. To account for multiple hypothesis testing of 39 repeat types, we use a Bonferroni correction to set a significance threshold of $p = 0.05/39 = 0.0013$. Sample Gaussian fits to the background distribution with annotated significance thresholds are shown in Fig. S94. Given the use of a conservative Bonferroni correction, we suspect that our estimate of TFs with significant binding to repeats is an underestimate.

##### 1.22 Paralog analysis

To identify paralogous TFs where low affinity interactions with repeats may confer binding specificity, we computed the cosine similarity in repeat preferences and motif preferences for each pair in the same TF structural family and species.

###### 1.22.1 Repeat similarity calculations

For each paralogous TF pair, we computed the cosine similarity between the vectors for their respective repeat Z-scores with the `cosine_similarity` function from `sklearn.metrics.pairwise`. Z-scores were computed and aggregated as described above (Section 1.21.2). Aggregated Z-scores for 1,291 TFs binding to 39 repeat types are available in an OSF repository (https://osf.io/gbxhz/). Code to replicate these calculations can be found at https://github.com/FordyceLab/STR_analysis.

###### 1.22.2 Motif similarity calculations

For each paralogous TF pair, we computed the cosine similarity between the flattened PWM matrices with the `cosine_similarity` function from `sklearn.metrics.pairwise`. In cases where one PWM is longer than the other, we compare each possible register. For example, comparison of `CCACGTGGC` has much greater similarity with `CACGTGGC` starting at the second position than at the first. We also computed the cosine similarity with the reverse complement of one matrix. The code to generate a PWM reverse complement is included below and on GitHub (https://github.com/FordyceLab/ STR_analysis), along with all other code required to compute motif similarity:

```python
import numpy as np

# pwm is an N x 4 numpy array
def compute_rc_matrix(pwm):
    pwm_rc = np.zeros(pwm.shape)

    # swap + reverse A position (0) and T position (3)
    pwm_rc[:,0] = pwm[::-1,3]
    pwm_rc[:,3] = pwm[::-1,0]

    # swap + reverse C position (1) and G position (2)
    pwm_rc[:,1] = pwm[::-1,2]
    pwm_rc[:,2] = pwm[::-1,1]

    return pwm_rc
```

##### 1.22.3 Sliding Z-score binding prediction

The use of 8-mer Z-scores for binding prediction relies on the same principles of statistical mechanics outlined in Section 1.12.2. To predict binding from uPBM-derived Z-scores, we applied the following formula:

$$\Delta\Delta G \propto \sum_j Z_j, \tag{45}$$

where $Z_j$ is the binding intensity Z-score for a given uPBM 8-mer. For the Pho4 and MAX comparisons shown in Fig. 6F, we considered binding to each possible 8-mer in a sequence as a separate microstate, as for the partition function calculation in Section 1.12.2.

#### 1.23 Bioinformatic analysis of repeat content within the human genome

STRs in the human genome were identified with Tandem Repeats Finder (TRF) version 4.09 [10] (source code: https://github.com/Benson-Genomics-Lab/TRF). We used the following command for each chromosome: `trf [chr.fa] 2 3 5 80 10 30 6 -d -h -l 6`. We used the following parameters:

- match score: +2
- mismatch penalty: −3
- indel penalty: −5
- match probability: 80%
- mismatch probability: 10%
- minimum score to report: 30
- maximum repeat period: 6

We chose generous parameters for repeat identification, as slightly-scrambled repeats had similar affinities as perfect repeats (Fig. 5J). For general calculations of repeat content in the human genome, we used the most complete human genome reference from the CHM13hTERT cell line released by the T2T consortium [38]. For repeat enrichment calculations involving genome annotations, we used the hg38 reference genome, as no liftOver tools yet exist for the CHM13 reference genome to our knowledge.

##### 1.23.1 Repeat perfection

For each STR identified, Tandem Repeats Finder reports several statistics, including `%match`, or the percentage of bases in the identified STR which match the consensus repeat pattern exactly. To identify the "perfection" of the median repeat in the genome, we took the median value of the `%match` statistic across all repeats in the genome.

##### 1.23.2 Human genome enhancer annotations

We used enhancer annotations for the human genome from the following 3 sources:

- **Enhancer Atlas**. We downloaded genome annotations and activity measurements of enhancers across 278 human cell types from EnhancerAtlas 2.0 (enhanceratlas.org [19]). Enhancer coordinates were lifted from hg19 to hg38 with UCSC genome browser's liftOver tool [22].
- **FANTOM**. We downloaded enhancer annotations from FANTOM 5 (https://fantom.gsc.riken.jp/5/ [6]) for the GRCh38 (hg38) build.
- **HACER**. We downloaded enhancer annotations from HACER for all available cell types/assays (http://bioinfo.vanderbilt.edu/AE/HACER/download.html [44]). Enhancer coordinates were lifted from hg19 to hg38 with UCSC genome browser's liftOver tool [22].

##### 1.23.3 Identifying enhancers containing STRs

For each of the 3 enhancer annotation databases, we counted the number of enhancers which overlap with an STR. We iteratively checked whether the overlap between each enhancer and the list of short tandem repeats for hg38 (each defined programmatically as an array of genome coordinates) was $> 0$ bp. If so, we considered that enhancer to contain or overlap with an STR.

##### 1.23.4 Calculation of STR enrichment in enhancers

For each of the 3 enhancer annotation databases, we calculated enrichment of repeats in enhancers as an odds ratio. We construct a contingency table for each calculation:

| | bp in enhancers | bp not in enhancers |
|---|---|---|
| bp in STRs | a | b |
| bp not in STR | c | d |

$$\text{Odds ratio (OR)} = \frac{a/b}{c/d} = \frac{a \cdot d}{b \cdot c} \tag{46}$$

To populate the contingency table, we count the number of bp in the genome that fall into each of the 4 categories (e.g. $b$ is the number of bp in STRs but not enhancers).

**Matched negatives.** For each of the 3 enhancer annotation databases, we calculated enrichment of repeats in non-enhancer regions of the genome with similar nucleotide and length composition. Code to generate a set of matched negatives is available on GitHub (https://github.com/FordyceLab/STR_analysis).

**STR enrichment in strong enhancers.** We repeated STR enrichment calculations for enhancers with a normalized strength above a given threshold according to Enhancer Atlas. We used enhancer strength data from Enhancer Atlas, calculated according to the overall signal of the enhancer-identification assay (CAGE-seq, p300 ChIP, GRO-seq, etc.) [19]. A modified contingency table used in these calculations is below:

| | bp in enhancers with signal $\geq 6$ | bp not in enhancers or in enhancers with signal $<6$ |
|---|---|---|
| bp in STRs | a | b |
| bp not in STR | c | d |

To assess whether we would observe an enrichment of STRs in the strongest enhancers through random chance with our thresholding procedure, we shuffled the relationships between enhancers and the number of cell types. We repeated this shuffling procedure 1,000 times and calculated enrichment for each of these shuffled arrangements, and we report the mean and standard deviation across the 1,000 shuffled trials in Fig. 7A. Code to replicate these calculations is available at https://github.com/FordyceLab/STR_analysis.

**STR enrichment in broadly active enhancers.** We repeated STR enrichment calculations for enhancers active in $x$ cell types, where $x$ is some threshold number of cell types, according to Enhancer Atlas. A modified contingency table used in these calculations is below:

| | bp in enhancers active in $\geq 15$ cell types | bp not in enhancers or in enhancers active in $<15$ cell types |
|---|---|---|
| bp in STRs | a | b |
| bp not in STR | c | d |

As above, to assess whether we would observe an enrichment of STRs in broadly active enhancers through random chance with our thresholding procedure, we shuffled the relationships between enhancers and the number of cell types. We repeated this shuffling procedure 1,000 times and calculated enrichment for each of these shuffled arrangements, and we report the mean and standard deviation across the 1,000 shuffled trials in Fig. 7B. Code to replicate these calculations is available at https://github.com/FordyceLab/STR_analysis.

#### 1.24 Calculating mutation rates

Mutation rates displayed in Fig. 7C were calculated by dividing the number of mutations per generation by the number of germ-line cell divisions per generation. Reported values for *S. cerevisiae*, *C. elegans*, *D. melanogaster*, and *H. sapiens* were cited from [30]. Mutation rates per generation for *A. thaliana* were quoted from [39] for single base substitutions and small indels (1–3 bp) and from [34] for STRs. We assumed 34 germ-line cell divisions per generation as reported for long-day *A. thaliana* growth in [45].

### 2 Supplemental calculations

#### 2.1 Statistical mechanics basis for increased entropy of TF-STR binding

To ask how sequence complexity (repetitiveness) contributes to sequence configurational entropy, we can constrain the enthalpy of the system ($\Delta H$) and determine what sequence configuration yields maximum entropy for the system. We define sequence configurational entropy as the entropy arising from the binding of an arbitrary nucleic acid-binding protein to some sequence with `N` binding sites given an arbitrary sequence-to-binding energy mapping.

We can define the change in Gibbs free energy upon binding:

$$\Delta G = -k_B T \ln(Z), \tag{47}$$

where $k_B$ is the Boltzmann constant, and $Z$ is the partition function:

$$Z = \sum_{i=1}^{N} e^{-E_i/k_B T}, \tag{48}$$

where $e^{-E_i/k_B T}$ is the Boltzmann weight for state $i$. The Boltzmann weight is proportional to the probability that state $i$ (with energy $E_i$) will be occupied. We can define the probability of being in state $i$:

$$p_i = \frac{e^{-E_i/k_B T}}{Z} = \frac{e^{-E_i/k_B T}}{\sum_{j=1}^{N} e^{-E_j/k_B T}} \tag{49}$$

We next define enthalpy of the system, $\Delta H$:

$$\Delta H = \langle E \rangle = \sum_{i=1}^{N} E_i \cdot p_i = \frac{\sum_{i=1}^{N} E_i \cdot e^{-E_i/k_B T}}{Z}, \tag{50}$$

Finally, we define entropy:

$$S = -k_B \sum_{i=1}^{N} p_i \cdot \ln(p_i) \tag{51}$$

We can now ask, with some fixed $\langle E \rangle$, under what sequence configuration is the minimum energy ($\Delta G$) obtained? We note that this constraint on the energy distribution of the system is required for meaningful analysis of this system: without it, optimization will yield $N \to \infty$ and $E_i \to -\infty$. Given the monotonic relationship between $\Delta G$ and $Z$, we can minimize $\Delta G$ by maximizing $Z$. We can make use of Lagrange multipliers to maximize $Z$ under the constraint $\langle E \rangle = C$.

$$\mathcal{L}(\vec{E}, \lambda) = f(\vec{E}) - \lambda \cdot g(\vec{E}), \tag{52}$$

where $f(\vec{E}) = Z$ and $g(\vec{E}) = \langle E \rangle = C$ and $\vec{E}$ is the vector of energies for all states.

##### 2.1.1 Toy example: two binding sites

In the simplest case, we can imagine two binding sites, $A$ and $B$ with Boltzmann weights $e^{-E_A/k_B T}$ and $e^{-E_B/k_B T}$, respectively, for an arbitrary nucleic acid-binding protein. For convenience, let $x = E_A/k_B T$ and $y = E_B/k_B T$. We can rewrite Eqn. 52 as:

$$\mathcal{L}(x, y, \lambda) = f(x, y) - \lambda \cdot g(x, y) \tag{53}$$

$$f(x,y) = Z = \sum_{i=1}^{N} e^{-E_i/k_BT} = e^{-x} + e^{-y} \tag{54}$$

$$g(x,y) = \langle E \rangle = \sum_{i=1}^{N} E_i \cdot p_i = \frac{k_BTxe^{-x} + k_BTye^{-y}}{e^{-x} + e^{-y}} = C \tag{55}$$

Finally, this yields our Langrangian for a two-state system:

$$\mathcal{L}(x,y,\lambda) = e^{-x} + e^{-y} - \lambda \left( \frac{k_BTxe^{-x} + k_BTye^{-y}}{e^{-x} + e^{-y}} \right) \tag{56}$$

We now take the set the gradient of $\mathcal{L}$ equal to zero to find minima/maxima:

$$\nabla \mathcal{L}(x,y,\lambda) = 0 \implies \nabla f(x,y) = \lambda \cdot \nabla g(x,y) \tag{57}$$

Computing the partial derivatives, we get:

$$-e^{-x} = \lambda k_BTe^{-x} \left[ \frac{(1-x)(e^{-x} + e^{-y}) + xe^{-x} + ye^{-y}}{(e^{-x} + e^{-y})^2} \right] \tag{58}$$

$$-e^{-y} = \lambda k_BTe^{-y} \left[ \frac{(1-y)(e^{-x} + e^{-y}) + xe^{-x} + ye^{-y}}{(e^{-x} + e^{-y})^2} \right] \tag{59}$$

Dividing Eqn. 58 by $e^{-x}$ and Eqn. 59 by $e^{-y}$ and simplifying, we can write:

$$\lambda k_BT \left[ \frac{(1-x)(e^{-x} + e^{-y}) + xe^{-x} + ye^{-y}}{(e^{-x} + e^{-y})^2} \right] = \lambda k_BT \left[ \frac{(1-y)(e^{-x} + e^{-y}) + xe^{-x} + ye^{-y}}{(e^{-x} + e^{-y})^2} \right] \tag{60}$$

which yields $x = y$. Physically, we can interpret the equality $x = y$ that the energy of states $A$ and $B$ are exactly equal, which implies that the binding states themselves are identical. $A$ and $B$ are identical if the nucleic acid sequence for these two binding sites are the same. This relationship can be easily visualized by plotting entropy as a function of $p_A$ (the probability state $A$ will be occupied). Given that there are only two states in our system, there is only one free parameter, $p_A$.

###### 2.1.2 General case: $N$ binding sites

Having demonstrated that entropy is maximized when two states are equally likely in a two-state system, we can show that sequence configurational entropy is always maximized when *all* states are equally likely. Physically, all states are equally likely when a sequence is "perfectly repetitive" (*i.e.* a homopolymeric DNA sequence). We begin by writing a general form of Eqn. 57

$$\nabla \mathcal{L}(\vec{x},\lambda) = 0 \implies \nabla f(\vec{x}) = \lambda \cdot \nabla g(\vec{x}) \tag{61}$$

We compute the partial derivative of $g(\vec{x})$ for arbitrary element in $\vec{x}$, $x_i$:

$$g_{x_i} = k_BTe^{-x_i} \left[ \frac{(1-x_i)(\sum_j e^{-x_j}) + \sum_j x_ie^{-x_j}}{(\sum_j e^{-x_j})^2} \right] \tag{62}$$

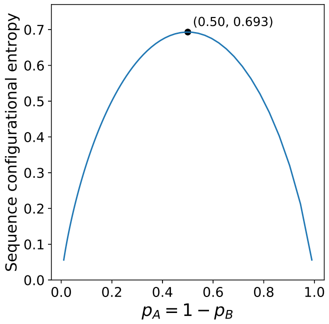

Figure A: Sequence configurational entropy as a function of $p_A$, where $p_B$ is constrained by $p_A$. Sequence configurational entropy is maximized $p_A = p_B = 1/2$, *i.e.* when the two binding sites are identical.

We can rewrite Eqn. 62, substituting Eqns. 48 and 50:

$$g_{x_i} = k_B T e^{-x_i} \left[ \frac{1 - x_i + \langle E \rangle}{Z} \right] \tag{63}$$

Similarly, the partial derivative of partial derivative of $f(\vec{x})$ for arbitrary element in $\vec{x}$, $x_i$:

$$f_{x_i} = -e^{-x_i} \tag{64}$$

From Eqns. 63 and 64, we get the Lagrangian:

$$\begin{bmatrix} -e^{-x_1} \\ -e^{-x_2} \\ \vdots \\ -e^{-x_N} \end{bmatrix} = \frac{\lambda k_B T}{Z} \begin{bmatrix} e^{-x_1}(1 - x_1 + \langle E \rangle) \\ e^{-x_2}(1 - x_2 + \langle E \rangle) \\ \vdots \\ e^{-x_N}(1 - x_N + \langle E \rangle) \end{bmatrix} \tag{65}$$

Finally, we write the following equality from Eqn. 65:

$$e^{-x_1}(1 - x_1 + \langle E \rangle) = e^{-x_2}(1 - x_2 + \langle E \rangle) = \cdots = e^{-x_N}(1 - x_N + \langle E \rangle) \tag{66}$$

which yields the general solution:

$$x_1 = x_2 = \cdots = x_N \tag{67}$$

For the case $N = 3$, we can visualize the entropy landscape on a 3D surface plot, where the $x$- and $y$- axes represent the two free parameters, $p_A$ and $p_B$, and the $z$-axis gives configurational entropy. The probability of occupying the third state, $p_C$ is constrained by the free parameters, $p_A$ and $p_B$.

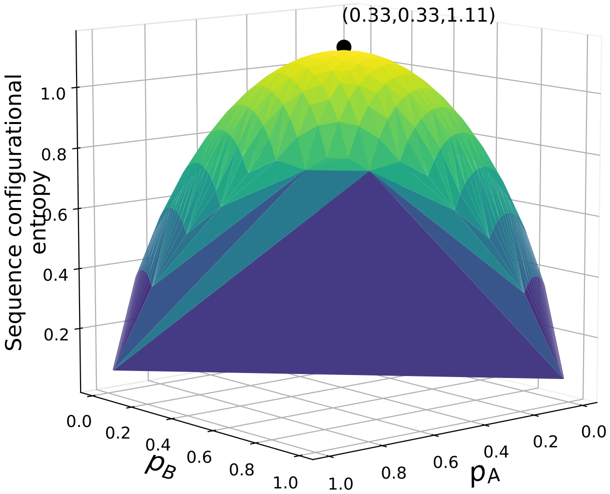

Figure B: Sequence configurational entropy as a function of $p_A$ and $p_B$, where $p_C$ is constrained by $p_A$ and $p_B$. Sequence configurational entropy is maximized $p_A = p_B = p_C = 1/3$, *i.e.* when all three sites are identical.

#### 2.2 Surface density of transcription factors on the MITOMI device

The average distance between two transcription factors ($d$) can be approximated by computing the total number of TFs under the button valve. $d = 2r_m$, where $A_{mean} = \pi r_m^2$, the mean area per TF. The mean area per TF can be given by:

$$A_{mean} = \pi r_m^2 = \frac{\pi r_b^2}{N_{molecules}}, \tag{68}$$

where $r_b$ is the radius of the circular area under the button valve where TFs are patterned. The number of molecules under the button valve is given by:

$$N_{molecules} = x \cdot N_A \cdot V_{button} = x \cdot N_A \cdot \pi r_b^2 h, \tag{69}$$

where $x$ is the molar concentration of TFs under the button valve, and $h$ is the height of the microfluidic chamber under the button valve. Combining Eqns. 68 and 69 to solve for $r_m$ yields:

$$r_m = \sqrt{\frac{r_b^2}{x \cdot N_A \cdot \pi r_b^2 h}} = \frac{1}{\sqrt{x \cdot N_A \cdot \pi h}} \tag{70}$$

The height of the MITOMI chambers is $\sim 15\,\mu m$ [18], and the concentration of TF monomer is $< 20\,nM$ [32]. To estimate the lower bound of distance between neighboring TFs, we assume [TF dimer] $= 10\,nM = 10^{-23}\,mol/\mu m^3$. We plug these estimates into Eqn. 70:

$$r_m = \frac{1}{\sqrt{(10^{-23}\,mol/\mu m^3) \cdot (6.022 \times 10^{23}\,molecules/mol) \cdot \pi \cdot (15\,\mu m)}} = 59\,nm. \tag{71}$$

The mean distance between neighboring TFs on the MITOMI device surface is therefore $\sim 118\,nm$. We emphasize that this estimate is likely to be a lower bound on the mean distance between neighboring TFs, as [TF] on the device rarely reaches 20 nM. We assume all TFs on the surface are dimerized, as monomeric TFs are unlikely to contribute substantially to binding at equilibrium; 10 nM is the maximum *effective* concentration.

Assuming a mean distance between DNA base pairs of 0.34 nm [36], the 54 bp oligonucleotides in DNA Library 1 would span $\sim 18\,nm$ and the 149 bp oligonucleotides in DNA Library 2 would span $\sim 51\,nm$. We conclude that it is unlikely for two neighboring TFs to bind the same DNA oligo, as the mean distance between neighboring TFs exceeds the length of DNA oligos. We therefore assume a two-state binding model, where DNA oligos can be either bound or unbound. We note, however, that multiple TFs can and do bind a single DNA oligo in solution-phase assays, such as EMSAs (Fig. 2D).

### 3 Supplemental tables

| **Sequence name** | **Full sequence** |
|---|---|
| Motif + AC/GT repeat 1 | CGCCACACACACACATGTCACGTGACATGTGTGTGTGT GGTCCGGCGGTATGAC |
| Motif + AC/GT repeat 2 | CGCCACACACACAACTGTCACGTGACATGTGTGTGTGT GGTCCGGCGGTATGAC |
| Motif + AC/GT repeat 3 | CGCCACACACACAACTGTCACGTGACAGTTGTGTGTGT GGTCCGGCGGTATGAC |
| Motif + AC/GT repeat 4 | CGCCACACATGTAACTGTCACGTGACAGTTGTGTGTGT GGTCCGGCGGTATGAC |
| Motif + AC/GT repeat 5 | CGCCCAACCAACACATGTCACGTGACATGTGTTGGTGT GGTCCGGCGGTATGAC |
| Motif + GT/AC repeat 1 | CGCTGTGTGTGTGTGTGTCACGTGACACACACCACAAA CGTCCGGCGGTATGAC |
| Motif + GT/AC repeat 2 | CGCTGTGTGTGTGTGTGTCACGTGACACACAACACACA CGTCCGGCGGTATGAC |
| Motif + GT/AC repeat 3 | CGCTGTGTGTGTGTGTGTCACGTGACACCACACCACAA AGTCCGGCGGTATGAC |
| Motif + AT/CG repeat 1 | CGCATATATATATAGAGTCACGTGACTCTCGCGCGCGC GGTCCGGCGGTATGAC |
| Motif + AT/CG repeat 2 | CGCTATATATATATATGTCACGTGACAGGCGCGCGCGC GGTCCGGCGGTATGAC |
| Motif + CG/AT repeat 1 | CGCGCGCGCGCGCAGAGTCACGTGACTCTATATATATA TGTCCGGCGGTATGAC |
| Motif + CG/AT repeat 2 | CGCCGCGCGCGCGAGTGTCACGTGACACTATATATATA TGTCCGGCGGTATGAC |
| Motif + short GT/AC repeat 2 | CGCGTGCGTTGTGTGTGTCACGTGACACACAACTCAAC AGTCCGGCGGTATGAC |
| Motif + short GT/AC repeat 3 | CGCAAGGCTTGTGTGTGTCACGTGACACCACACATCGA TGTCCGGCGGTATGAC |
| Motif + short GT/AC repeat 4 | CGCACAGGTGTGTGTTGTCACGTGACACACACACACTG TGTCCGGCGGTATGAC |
| Motif + short CG/AT repeat 1 | CGCAGTGACGCGCAGAGTCACGTGACTCTATATTACGC TGTCCGGCGGTATGAC |
| Motif + short A/T repeat 1 | CGCCGGAACAAAAGGGGTCACGTGACCCCTTTTTTCAG TGTCCGGCGGTATGAC |
| Motif + random 1 | CGCACAGTCACTTAACGTCACGTGACCGGGGTATTTCA GGTCCGGCGGTATGAC |
| Motif + random 2 | CGCGATCGGCGCGGACGTCACGTGACCTCTAATTTTAA AGTCCGGCGGTATGAC |
| Motif + random 3 | CGCCAATTCCGCGTTTGTCACGTGACGACCGGATAGAT AGTCCGGCGGTATGAC |

Table 1: Library 1 sequences

| Sequence name | Full sequence |
|---|---|
| Mutated motif + GT/AC repeat 1 | CGCTGTGTGTGTGTGTGTCACG**C**GACACACACCACAAA<br>CGTCCGGCGGTATGAC |
| Mutated motif + AT/CG repeat 1 | CGCATATATATATAGAGTCACG**C**GACTCTCGCGCGCGC<br>GGTCCGGCGGTATGAC |
| Mutated motif + CG/AT repeat 1 | CGCGCGCGCGCGCAGAGTCACG**C**GACTCTATATATATA<br>TGTCCGGCGGTATGAC |
| Mutated motif + CG/AT repeat 2 | CGCCGCGCGCGCGAGTGTCACG**C**GACACTATATATATA<br>TGTCCGGCGGTATGAC |
| Mutated motif + random 1 | CGCACAGTCACTTAACGTCACG**C**GACCGGGGTATTTCA<br>GGTCCGGCGGTATGAC |
| Mutated motif + random 2 | CGCGATCGGCGCGGACGTCACG**C**GACCTCTAATTTTAA<br>AGTCCGGCGGTATGAC |
| Mutated motif + random 3 | CGCCAATTCCGCGTTTGTCACG**C**GACGACCGGATAGAT<br>AGTCCGGCGGTATGAC |
| No motif + AT/CG repeat 3 | CGCATATATATATATATATATCGCGCGCGCGCGCGCGCG<br>GTCCGGCGGTATGAC |
| No motif + AC/GT repeat 6 | CGCACACACACACACACACACACTGTGTGTGTGTGTG<br>TGTGGTCCGGCGGTATGAC |
| No motif + random 4 | CGCCGCAGAGTCTCATTCTCCATACGCTGTAAAGT<br>GGGAGTCCGGCGGTATGAC |
| No motif + random 5 | CGCCCTGTCGGGAAAGACCATTGTTCTCTAAAGAG<br>CTCGGTCCGGCGGTATGAC |

Table 1: Library 1 sequences (continued)

| Sequence name | Sub-library | Full sequence |
|---|---|---|
| Random 1 | Type | CGCCCCAGCACTGCCAAGCCGACGTTAAAACGGGTTGC TTCATCAATCGAATGTCAATACATAGTCACGTGACACG CGGTAGGCTCGCTATCGGCACTTGCGCTTGAGTGCATC GAATAGTTCGGTTTATGAGCGTCCGGCGGTATGAC |
| Random 2 | Type | CGCGTTTAACGCGGACGCGTAATCCTTTCCACATCGTG GCGCCTGCCGATCCCGACTGGACTTGTCACGTGACGAA TGGAATGGTTCCGACAATAGATCAACAAGAAGTGGTAA GTAGGCTCACATTGCTTTCCGTCCGGCGGTATGAC |
| A/A 60bp | Type | CGCAAAAAAAAAAAAAAAAAAAAAAAAAAAAAAAAAAA AAAAAAAAAAAAAAAAAAAAAAAAAGTCACGTGACAAA AAAAAAAAAAAAAAAAAAAAAAAAAAAAAAAAAAAAAA AAAAAAAAAAAAAAAAAAACGTCCGGCGGTATGAC |
| AT/AT 60bp | Type | CGCATATATATATATATATATATATATATATATATATA TATATATATATATATATATATATATGTCACGTGACATA TATATATATATATATATATATATATATATATATATATA TATATATATATATATATATCGTCCGGCGGTATGAC |
| C/C 60bp * | Type | CGCCCCCCCCCCCCCCCCCCCCCCCCCCCCCCCCCCCC CCCCCCCCCCCCCCCCCCCCCCCCCGTCACGTGACCCC CCCCCCCCCCCCCCCCCCCCCCCCCCCCCCCCCCCCCC CCCCCCCCCCCCCCCCCCCCGTCCGGCGGTATGAC |
| CG/CG 60bp | Type | CGCGCGCGCGCGCGCGCGCGCGCGCGCGCGCGCGCGCG CGCGCGCGCGCGCGCGCGCGCGCGCGTCACGTGACGCG CGCGCGCGCGCGCGCGCGCGCGCGCGCGCGCGCGCGCG CGCGCGCGCGCGCGCGCGCCGTCCGGCGGTATGAC |
| ACGT/ACGT 60bp | Type | CGCGTACGTACGTACGTACGTACGTACGTACGTACGTA CGTACGTACGTACGTACGTACGTACGTCACGTGACGTA CGTACGTACGTACGTACGTACGTACGTACGTACGTACG TACGTACGTACGTACGTACCGTCCGGCGGTATGAC |
| ACTG/AGTC 60bp | Type | CGCACTGACTGACTGACTGACTGACTGACTGACTGACT GACTGACTGACTGACTGACTGACTGGTCACGTGACCAG TCAGTCAGTCAGTCAGTCAGTCAGTCAGTCAGTCAGTC AGTCAGTCAGTCAGTCAGTCGTCCGGCGGTATGAC |
| ATCG/ATCG 60bp | Type | CGCATCGATCGATCGATCGATCGATCGATCGATCGATC GATCGATCGATCGATCGATCGATCGGTCACGTGACCGA TCGATCGATCGATCGATCGATCGATCGATCGATCGATC GATCGATCGATCGATCGATCGTCCGGCGGTATGAC |
| ATGC/ATGC 60bp | Type | CGCATGCATGCATGCATGCATGCATGCATGCATGCATG CATGCATGCATGCATGCATGCATGCGTCACGTGACGCA TGCATGCATGCATGCATGCATGCATGCATGCATGCATG CATGCATGCATGCATGCATCGTCCGGCGGTATGAC |
| AG/CT 15bp | Length | CGCCCCAGCACTGCCAAGCCGACGTTAAAACGGGTTGC TTCATCAATCGGAGAGAGAGAGAGAGTCACGTGACTCT CTCTCTCTCTCTTATCGGCACTTGCGCTTGAGTGCATC GAATAGTTCGGTTTATGAGCGTCCGGCGGTATGAC |
| AG/CT 30bp | Length | CGCCCCAGCACTGCCAAGCCGACGTTAAAACGGGAGAG AGAGAGAGAGAGAGAGAGAGAGAGAGTCACGTGACTCT CTCTCTCTCTCTCTCTCTCTCTCTCTCTTGAGTGCATC GAATAGTTCGGTTTATGAGCGTCCGGCGGTATGAC |

Table 2: Library 2 sequences

| Sequence name | Sub-library | Full sequence |
|---|---|---|
| AG/CT 45bp | Length | CGCCCCAGCACTGCCAAGGGAGAGAGAGAGAGAGAGAGAG<br>AGAGAGAGAGAGAGAGAGAGAGAGAGTCACGTGACTCT<br>CTCTCTCTCTCTCTCTCTCTCTCTCTCTCTCTCTCTCT<br>CTCTAGTTCGGTTTATGAGCGTCCGGCGGTATGAC |
| AG/CT 60bp | Type/length | CGCGAGAGAGAGAGAGAGAGAGAGAGAGAGAGAGAGAG<br>AGAGAGAGAGAGAGAGAGAGAGAGAGTCACGTGACTCT<br>CTCTCTCTCTCTCTCTCTCTCTCTCTCTCTCTCTCTCT<br>CTCTCTCTCTCTCTCTCTCTCCGTCCGGCGGTATGAC |
| CG/AT 15bp * | Length | CGCCCCAGCACTGCCAAGCCGACGTTAAAACGGGTTGC<br>TTCATCAATCCGCGCGCGCGCGCGCGTCACGTGACATA<br>TATATATATATATATATCGGCACTTGCGCTTGAGTGCATC<br>GAATAGTTCGGTTTATGAGCGTCCGGCGGTATGAC |
| CG/AT 30bp * | Length | CGCCCCAGCACTGCCAAGCCGACGTTAAAACGCGCGCG<br>CGCGCGCGCGCGCGCGCGCGCGCGCGTCACGTGACATA<br>TATATATATATATATATATATATATATATTTGAGTGCATC<br>GAATAGTTCGGTTTATGAGCGTCCGGCGGTATGAC |
| CG/AT 45bp * | Length | CGCCCCAGCACTGCCAAGCGCGCGCGCGCGCGCGCGCG<br>CGCGCGCGCGCGCGCGCGCGCGCGCGTCACGTGACATA<br>TATATATATATATATATATATATATATATATATATATA<br>TATAAGTTCGGTTTATGAGCGTCCGGCGGTATGAC |
| CG/AT 60bp * | Type/length | CGCGCGCGCGCGCGCGCGCGCGCGCGCGCGCGCGCGCGCG<br>CGCGCGCGCGCGCGCGCGCGCGCGCGTCACGTGACATA<br>TATATATATATATATATATATATATATATATATATATA<br>TATATATATATATATATATATCGTCCGGCGGTATGAC |
| GT/AC 15bp | Length | CGCCCCAGCACTGCCAAGCCGACGTTAAAACGGGTTGC<br>TTCATCAATCTGTGTGTGTGTGTGTGTCACGTGACACA<br>CACACACACACATATCGGCACTTGCGCTTGAGTGCATC<br>GAATAGTTCGGTTTATGAGCGTCCGGCGGTATGAC |
| GT/AC 30bp | Length | CGCCCCAGCACTGCCAAGCCGACGTTAAAACGGGTGTG<br>TGTGTGTGTGTGTGTGTGTGTGTGTGTCACGTGACACA<br>CACACACACACACACACACACACACACTTGAGTGCATC<br>GAATAGTTCGGTTTATGAGCGTCCGGCGGTATGAC |
| GT/AC 45bp | Length | CGCCCCAGCACTGCCAAGTGTGTGTGTGTGTGTGTGTG<br>TGTGTGTGTGTGTGTGTGTGTGTGTGTCACGTGACACA<br>CACACACACACACACACACACACACACACACACACACA<br>CACAAGTTCGGTTTATGAGCGTCCGGCGGTATGAC |
| GT/AC 60bp | Type/length | CGCGTGTGTGTGTGTGTGTGTGTGTGTGTGTGTGTGTGTG<br>TGTGTGTGTGTGTGTGTGTGTGTGTGTCACGTGACACA<br>CACACACACACACACACACACACACACACACACACACA<br>CACACACACACACACACACACCGTCCGGCGGTATGAC |
| AG/CT 10bp distance | Distance | CGCCCCAGCACTGCCAAGCCGACGAGAGAGAGAGAGAG<br>AGAGAGAGAGAGAGATCAATACATAGTCACGTGACACG<br>CGGTAGGCTCTCTCTCTCTCTCTCTCTCTCTCTCTCTC<br>GAATAGTTCGGTTTATGAGCGTCCGGCGGTATGAC |
| AG/CT 20bp distance | Distance | CGCCCCAGCACTGGAGAGAGAGAGAGAGAGAGAGAGAG<br>AGAGACAATCGAATGTCAATACATAGTCACGTGACACG<br>CGGTAGGCTCGCTATCGCTCTCTCTCTCTCTCTCTCTC<br>TCTCTCTCTGGTTTATGAGCGTCCGGCGGTATGAC |

Table 2: Library 2 sequences (continued)

| **Sequence name** | **Sub-library** | **Full sequence** |
|---|---|---|
| AG/CT 30bp distance | Distance | CGCGAGAGAGAGAGAGAGAGAGAGAGAGAGAGAGAGTTGC<br>TTCATCAATCGAATGTCAATACATAGTCACGTGACACG<br>CGGTAGGCTCGCTATCGGCACTTGCGCCTCTCTCTCTC<br>TCTCTCTCTCTCTCTCTCTCGTCCGGCGGTATGAC |
| AG/CT 45bp distance | Distance | CGCGAGAGAGAGAGAGAGCCGACGTTAAAACGGGTTGC<br>TTCATCAATCGAATGTCAATACATAGTCACGTGACACG<br>CGGTAGGCTCGCTATCGGCACTTGCGCTTGAGTGCATC<br>GAATTCTCTCTCTCTCTCTCGTCCGGCGGTATGAC |
| CG/AT 10bp distance * | Distance | CGCCCCAGCACTGCCAAGCCGACGCGCGCGCGCGCGCG<br>CGCGCGCGCGCGCGCTCAATACATAGTCACGTGACACG<br>CGGTAGGTATATATATATATATATATATATATATATATAC<br>GAATAGTTCGGTTTATGAGCGTCCGGCGGTATGAC |
| CG/AT 20bp distance * | Distance | CGCCCCAGCACTGGCGCGCGCGCGCGCGCGCGCGCGCG<br>CGCGCCAATCGAATGTCAATACATAGTCACGTGACACG<br>CGGTAGGCTCGCTATCGTATATATATATATATATATATAT<br>ATATATATAGGTTTATGAGCGTCCGGCGGTATGAC |
| CG/AT 30bp distance * | Distance | CGCGCGCGCGCGCGCGCGCGCGCGCGCGCGCGCGCGTTGC<br>TTCATCAATCGAATGTCAATACATAGTCACGTGACACG<br>CGGTAGGCTCGCTATCGGCACTTGCGCTATATATATAT<br>ATATATATATATATATATATACGTCCGGCGGTATGAC |
| CG/AT 45bp distance * | Distance | CGCGCGCGCGCGCGCGCGCCGACGTTAAAACGGGTTGC<br>TTCATCAATCGAATGTCAATACATAGTCACGTGACACG<br>CGGTAGGCTCGCTATCGGCACTTGCGCTTGAGTGCATC<br>GAATTATATATATATATATATCGTCCGGCGGTATGAC |
| GT/AC 10bp distance | Distance | CGCCCCAGCACTGCCAAGCCGACGTGTGTGTGTGTGTG<br>TGTGTGTGTGTGTGTTCAATACATAGTCACGTGACACG<br>CGGTAGGACACACACACACACACACACACACACACACACC<br>GAATAGTTCGGTTTATGAGCGTCCGGCGGTATGAC |
| GT/AC 20bp distance | Distance | CGCCCCAGCACTGGTGTGTGTGTGTGTGTGTGTGTGTG<br>TGTGTCAATCGAATGTCAATACATAGTCACGTGACACG<br>CGGTAGGCTCGCTATCGACACACACACACACACACACA<br>CACACACACGGTTTATGAGCGTCCGGCGGTATGAC |
| GT/AC 30bp distance | Distance | CGCGTGTGTGTGTGTGTGTGTGTGTGTGTGTGTGTGTTGC<br>TTCATCAATCGAATGTCAATACATAGTCACGTGACACG<br>CGGTAGGCTCGCTATCGGCACTTGCGCCACACACACAC<br>ACACACACACACACACACACACGTCCGGCGGTATGAC |
| GT/AC 45bp distance | Distance | CGCGTGTGTGTGTGTGTGTGCCGACGTTAAAACGGGTTGC<br>TTCATCAATCGAATGTCAATACATAGTCACGTGACACG<br>CGGTAGGCTCGCTATCGGCACTTGCGCTTGAGTGCATC<br>GAATACACACACACACACACACGTCCGGCGGTATGAC |
| GT/AC scrambled 1 | Fidelity | CGCGTGTGTGTGTGTGTGTGTGTGTGTGTGTGTGTGTGTG<br>TGTGTGTGTGTTTGTGGGTGTGTGTGTCACGTGACACA<br>CACACCCACAAACACACACACACACACACACACACACA<br>CACACACACACACACACACACCGTCCGGCGGTATGAC |
| GT/AC scrambled 2 | Fidelity | CGCGTGTGTGTGTGTGTGTGTGTGTGTGTGGGTGTGTGTG<br>TGTTTGTGTGTTTGTGGGTGTGTGTGTCACGTGACACA<br>CACACCCACAAACACACAAACACACACACACCCACACA<br>CACACACACACACACACACACCGTCCGGCGGTATGAC |

Table 2: Library 2 sequences (continued)

| **Sequence name** | **Sub-library** | **Full sequence** |
|---|---|---|
| GT/AC scrambled 3 | Fidelity | CGCGTGTGTGTGTGTGTGTGTGTGGGTGGGTGTGTGTG<br>TGTTTGTGTGTTTGTTGGTGTGTGTGTCACGTGACACA<br>CACACCAACAAACACACAAACACACACACACCCACCCA<br>CACACACACACACACACACCGTCCGGCGGTATGAC |
| GT/AC scrambled 4 | Fidelity | CGCGTGTGTGTGTGTGTGTGTGTGGGGGGGTGTGTGTT<br>TGTTTGTGTGTTTGTTGGTGTGTGTGTCACGTGACACA<br>CACACCAACAAACACACAAACAAACACACACCCCCCCA<br>CACACACACACACACACACCGTCCGGCGGTATGAC |
| GT/AC scrambled 5 | Fidelity | CGCGTTTGTGTGTGTGTGTGTGTGGGGTGGTGTGTGGT<br>TGTTTGGGTGTTTGTTGGTGTGTGTGTCACGTGACACA<br>CACACCAACAAACACCCAAACAACCACACACCACCCCA<br>CACACACACACACACAAACCGTCCGGCGGTATGAC |
| GT/AC scrambled 6 | Fidelity | CGCGTTTGTGTGTGTGTGTGTGTGGGGTGGTGTGTGGT<br>TGTTTGGGTGTTTGTTGGTGTGTGTGTCACGTGACACA<br>CACACCAACAAACACCCAAACAACCACACACCACCCCA<br>CACACACACACACACAAACCGTCCGGCGGTATGAC |
| GT/AC scrambled 7 | Fidelity | CGCTGGTTTGGTTTGTGGGTGGGGTTTGTTTTGGTGTT<br>TGTTGTGGGTGGTTTGGGTGGGGTTGTCACGTGACAAC<br>CCCACCCAAACCACCCACAACAAACACCAAAACAAACC<br>CCACCCACAAACCAAACCACGTCCGGCGGTATGAC |
| Motif + random 1 | Calibration | CGCACAGTCACTTAACGTCACGTGACCGGGG<br>TATTTCAGGTCCGGCGGTATGAC |
| Motif + random 3 | Calibration | CGCCAATTCCGCGTTTGTCACGTGACGACCG<br>GATAGATAGTCCGGCGGTATGAC |
| Motif + GT/AC repeat 2 | Calibration | CGCTGTGTGTGTGTGTGTCACGTGACACACA<br>ACACACACGTCCGGCGGTATGAC |
| Motif + CG/AT repeat 1 | Calibration | CGCGCGCGCGCGCAGAGTCACGTGACTCTAT<br>ATATATATGTCCGGCGGTATGAC |

Table 2: Library 2 sequences (continued)

| Sequence ID | Full sequence |
|---|---|
| 1 | GTGACTGGAGTTCAGACGTGTGCTCTTCCGATCGCGGAGCCTGGG AGGCACGCGAGGGCCGGGCCGGGCGTCTTGTCCGGCGGTATGAC |
| 2 | GTGACTGGAGTTCAGACGTGTGCTCTTCCGATCAGGAGGTCAAAG TCACACGTGCTGAGCCACTGACATGTCTTGTCCGGCGGTATGAC |
| 3 | GTGACTGGAGTTCAGACGTGTGCTCTTCCGATCCATTCGCAGGGA CACCACGTGGTTTCCAGAACTTGGGTCTTGTCCGGCGGTATGAC |
| 4 | GTGACTGGAGTTCAGACGTGTGCTCTTCCGATCTGGCCCAGGAGC CACCACGTGGTGCCTCCTAGGGCTGTCTTGTCCGGCGGTATGAC |
| 5 | GTGACTGGAGTTCAGACGTGTGCTCTTCCGATCGCGCCCGCCCAC CGGCACGCGGAGGGGCTTCCTGGAGTCTTGTCCGGCGGTATGAC |
| 6 | GTGACTGGAGTTCAGACGTGTGCTCTTCCGATCGGTGTAAAGATGC CCCACGTGCCAGGGGCCCAGACCGTCTTGTCCGGCGGTATGAC |
| 7 | GTGACTGGAGTTCAGACGTGTGCTCTTCCGATCAATTGAACCAGC CACCACGTGGTCCCCGGAGGCGCTGTCTTGTCCGGCGGTATGAC |
| 8 | GTGACTGGAGTTCAGACGTGTGCTCTTCCGATCTGTGGCTTATCA AACCACGTGGTTTATCAAAATAATGTCTTGTCCGGCGGTATGAC |
| 9 | GTGACTGGAGTTCAGACGTGTGCTCTTCCGATCCCGCCAGCTCTG CCCCATGCGCATGAGCGGCCGCCGGTCTTGTCCGGCGGTATGAC |
| 10 | GTGACTGGAGTTCAGACGTGTGCTCTTCCGATCGCGACCTGGACA AGGCACGTGGTTTAGTGAGCCCCTGTCTTGTCCGGCGGTATGAC |
| 11 | GTGACTGGAGTTCAGACGTGTGCTCTTCCGATCGAATTTGACCAA AACCACGTGGTTTGGCTGCAAGTTGTCTTGTCCGGCGGTATGAC |
| 12 | GTGACTGGAGTTCAGACGTGTGCTCTTCCGATCCGGGGAGGAGGA GGGCACGCGCCGGGAGGACCGCGAGTCTTGTCCGGCGGTATGAC |
| 13 | GTGACTGGAGTTCAGACGTGTGCTCTTCCGATCACCGACTTCATAG GGCACGAGGAAGATGAACGGAGAGTCTTGTCCGGCGGTATGAC |
| 14 | GTGACTGGAGTTCAGACGTGTGCTCTTCCGATCCCTCCTCTTCTGA AACACGTGTGGCGCATACAGCTGGTCTTGTCCGGCGGTATGAC |
| 15 | GTGACTGGAGTTCAGACGTGTGCTCTTCCGATCCCCCGGCGCCGGA ACCACGTGGTGCCTGTTTTGTCGGTCTTGTCCGGCGGTATGAC |
| 16 | GTGACTGGAGTTCAGACGTGTGCTCTTCCGATCGGGGCGGGGCGG CGCCACGCGCGAGGCCTCTCTTGCGTCTTGTCCGGCGGTATGAC |
| 17 | GTGACTGGAGTTCAGACGTGTGCTCTTCCGATCGGTGTTTCTTCTC CCCACATGCTGGAACCCAGGAGGGTCTTGTCCGGCGGTATGAC |
| 18 | GTGACTGGAGTTCAGACGTGTGCTCTTCCGATCGCAAGGCTCTTT GCCCACGTGTGCGCGCGCTCCAGTGTCTTGTCCGGCGGTATGAC |
| 19 | GTGACTGGAGTTCAGACGTGTGCTCTTCCGATCTTTAAAAAAAAA AATCACATGCTGGGGCCAGAGATAGTCTTGTCCGGCGGTATGAC |
| 20 | GTGACTGGAGTTCAGACGTGTGCTCTTCCGATCCAGAAGGCGAGG CCCCACGCGGGAAGACAGTCGAACGTCTTGTCCGGCGGTATGAC |
| 21 | GTGACTGGAGTTCAGACGTGTGCTCTTCCGATCGGCCTACTGCCC TCGCACATGCTCTGCCCACTTCTTGTCTTGTCCGGCGGTATGAC |
| 22 | GTGACTGGAGTTCAGACGTGTGCTCTTCCGATCTTGGTCCATGCCG CCCACGTGCTCCCCGGGGCTCCGGTCTTGTCCGGCGGTATGAC |

Table 3: gcPBM calibration library

| Sequence ID | Full sequence |
|---|---|
| 23 | GTGACTGGAGTTCAGACGTGTGCTCTTCCGATCCGGGCTGCCCCG GGTCACGTGACTCGGACACCGCCCGTCTTGTCCGGCGGTATGAC |
| 24 | GTGACTGGAGTTCAGACGTGTGCTCTTCCGATCTAGAACTGCACG AGCCACGCGGCCGAGCTGTTGGCGGTCTTGTCCGGCGGTATGAC |
| 25 | GTGACTGGAGTTCAGACGTGTGCTCTTCCGATCTCCCCAGGTGTA CAGCACATGGGAGGCTTAGGACTCGTCTTGTCCGGCGGTATGAC |
| 26 | GTGACTGGAGTTCAGACGTGTGCTCTTCCGATCACCACAGATCAG GACCACGTGCTCCCACGTCACCACGTCTTGTCCGGCGGTATGAC |
| 27 | GTGACTGGAGTTCAGACGTGTGCTCTTCCGATCGCCAAAAAGACC AACCACGTGATTTAGGGTAGGGCCGTCTTGTCCGGCGGTATGAC |
| 28 | GTGACTGGAGTTCAGACGTGTGCTCTTCCGATCAGACAGAAGTTT GCCCACGTGCTCCGGCCGCCTGAGGTCTTGTCCGGCGGTATGAC |
| 29 | GTGACTGGAGTTCAGACGTGTGCTCTTCCGATCGTAGTTCCTGGT CCGCACATGGTTAGGAGGTTCTCGGTCTTGTCCGGCGGTATGAC |
| 30 | GTGACTGGAGTTCAGACGTGTGCTCTTCCGATCTTGTTGCCTGCG GCCCACGTGGGAGAACTGCTTACTGTCTTGTCCGGCGGTATGAC |
| 31 | GTGACTGGAGTTCAGACGTGTGCTCTTCCGATCTCGCATCCGCAC GTGCACGTGCACCCGCTGCTGCACGTCTTGTCCGGCGGTATGAC |
| 32 | GTGACTGGAGTTCAGACGTGTGCTCTTCCGATCCCTGGACCTCCT GACCACGTGGTACTGCAGCTTCAAGTCTTGTCCGGCGGTATGAC |

Table 3: gcPBM calibration library (continued)

| Position | WT residue | Mutated residue | Protein region |
|---|---|---|---|
| 230 | Gly | Ala | Outside of DNA-binding domain |
| | | Val | |
| 231 | Ser | Ala | |
| | | Val | |
| 232 | Ser | Ala | |
| | | Val | |
| 233 | His | Ala | |
| | | Val | |
| 234 | Ser | Ala | |
| | | Val | |
| 235 | Arg | Ala | |
| | | Val | |
| 236 | Ser | Ala | |
| | | Val | |
| 237 | Leu | Ala | |
| | | Val | |
| 238 | Ser | Ala | |
| | | Val | |
| 239 | Lys | Ala | |
| | | Val | |
| 240 | Arg | Ala | |
| | | Val | |
| 241 | Arg | Ala | |
| | | Val | |
| 242 | Ser | Ala | |
| | | Val | |
| 243 | Ser | Ala | |
| | | Glu | |
| | | Val | |
| 244 | Gly | Ala | |
| | | Val | |
| 245 | Ala | Val | |
| 246 | Leu | Ala | |
| | | Val | |
| 247 | Val | Ala | |
| 248 | Asp | Ala | |
| | | Val | |
| 249 | Asp | Ala | |
| | | Val | |
| 250 | Asp | Ala | Basic region |
| | | Glu | |
| | | His | |
| | | Lys | |
| | | Gln | |
| | | Ser | |
| | | Val | |

Table 4: List of Pho4 mutants

| Position | WT residue | Mutated residue | Protein region |
|---|---|---|---|
| 251 | Lys | Ala | Basic region |
| | | Arg | |
| | | Ser | |
| | | Val | |
| 252 | Arg | Ala | |
| | | Asp | |
| | | Lys | |
| | | Gln | |
| | | Thr | |
| | | Val | |
| 253 | Glu | Ala | |
| | | Asp | |
| | | Arg | |
| | | Val | |
| 254 | Ser | Ala | |
| | | Val | |
| 255 | His | Ala | |
| | | Asn | |
| | | Arg | |
| | | Val | |
| 256 | Lys | Ala | |
| | | Glu | |
| | | Arg | |
| | | Val | |
| 257 | His | Ala | |
| | | Pro | |
| | | Thr | |
| | | Val | |
| 258 | Ala | Leu | |
| | | Arg | |
| | | Val | |
| 259 | Glu | Ala | |
| | | Asp | |
| | | Lys | |
| | | Asn | |
| | | Val | |
| 260 | Gln | Ala | |
| | | Glu | |
| | | Lys | |
| | | Asn | |
| | | Arg | |
| | | Val | |
| 261 | Ala | Val | |

Table 4: List of Pho4 mutants (continued)

| Position | WT residue | Mutated residue | Protein region |
|---|---|---|---|
| 262 | Arg | Ala | Basic region |
| | | His | |
| | | Lys | |
| | | Leu | |
| | | Gln | |
| | | Val | |
| | | Tyr | |
| 263 | Arg | Ala | |
| | | Lys | |
| | | Leu | |
| | | Met | |
| | | Asn | |
| | | Gln | |
| | | Val | |
| | | Trp | |
| | | Tyr | |
| 264 | Asn | Ala | |
| | | Glu | |
| | | Gly | |
| | | Ser | |
| | | Val | |
| 265 | Arg | Ala | |
| | | Lys | |
| 266 | Leu | Ala | Helix 1 |
| 267 | Ala | Val | |
| 268 | Val | Ala | |
| 269 | Ala | Val | |
| 270 | Leu | Ala | |
| | | Val | |
| 271 | His | Ala | |
| | | Val | |
| 272 | Glu | Ala | |
| | | Val | |
| 273 | Leu | Ala | |
| | | Val | |
| 274 | Ala | Val | |
| 275 | Ser | Ala | |
| | | Val | |
| 276 | Leu | Ile | |
| | | Ser | |
| | | Val | |
| 277 | Ile | Ala | |
| | | Val | |
| 278 | Pro | Ala | Loop |
| | | Gln | |
| | | Ser | |
| | | Val | |

Table 4: List of Pho4 mutants (continued)

| **Position** | **WT residue** | **Mutated residue** | **Protein region** |
|---|---|---|---|
| 279 | Ala | Val | Loop |
| 280 | Glu | Ala | |
| | | Val | |
| 281 | Trp | Ala | |
| | | Val | |
| 282 | Lys | Ala | |
| | | Gly | |
| | | Arg | |
| | | Val | |
| 283 | Gln | Ala | |
| | | Gly | |
| | | Val | |
| 284 | Gln | Ala | |
| | | Gly | |
| | | Val | |
| 285 | Asn | Ala | |
| | | Val | |
| 286 | Val | Ala | |
| | | Gly | |
| 287 | Ser | Ala | |
| | | Gly | |
| | | Val | |
| 288 | Ala | Gly | |
| | | Val | |
| 289 | Ala | Gly | |
| | | Lys | |
| | | Arg | |
| 290 | Pro | Ala | |
| | | Gly | |
| | | Val | |
| 291 | Ser | Ala | |
| | | Gly | |
| | | Val | |
| 292 | Lys | Ala | Helix 2 |
| | | Glu | |
| | | Gln | |
| | | Arg | |
| | | Val | |
| 293 | Ala | Val | |
| 294 | Thr | Ala | |
| | | Val | |
| 295 | Thr | Ala | |
| | | Val | |
| 296 | Val | Ala | |
| 297 | Glu | Ala | |
| | | Val | |
| 298 | Ala | Val | |

Table 4: List of Pho4 mutants (continued)

| **Position** | **WT residue** | **Mutated residue** | **Protein region** |
|---|---|---|---|
| 299 | Ala | Asp | Helix 2 |
| | | Val | |
| 300 | Cys | Ala | |
| 301 | Arg | Ala | |
| | | Val | |
| 302 | Tyr | Ala | |
| | | His | |
| | | Val | |
| 303 | Ile | Ala | |
| | | Leu | |
| | | Val | |
| 304 | Arg | Ala | |
| | | Val | |
| 305 | His | Ala | |
| | | Val | |
| 306 | Leu | Ala | |
| | | Val | |
| 307 | Gln | Ala | Outside of DNA-binding domain (C-terminus) |
| | | Val | |
| 308 | Gln | Ala | |
| | | Val | |
| 309 | Asn | Ala | |
| | | Val | |
| 310 | Val | Ala | |
| | | Asp | |
| | | Leu | |
| | | Asn | |
| 311 | Ser | Ala | |
| | | Val | |
| 312 | Thr | Ala | |
| | | Val | |

Table 4: List of Pho4 mutants (continued)

<table>
<tr><th>Date (yymmdd)</th><th>setup</th><th>fluor</th><th>slope</th><th>intercept</th><th>exposure (ms)</th><th>device</th><th>mean slope</th><th>mean intercept</th></tr>
<tr><td>181113</td><td>4</td><td>Alexa647</td><td>18.43</td><td>0</td><td>250</td><td>d1</td><td>18.43</td><td>0</td></tr>
<tr><td>190118</td><td>4</td><td>ATTO647</td><td>9.59</td><td>445</td><td>300</td><td>d1</td><td>9.59</td><td>445</td></tr>
<tr><td rowspan="2">190515</td><td>4</td><td>Alexa647</td><td>3.90</td><td>197</td><td>300</td><td>d1</td><td rowspan="2">3.92</td><td rowspan="2">179</td></tr>
<tr><td>4</td><td>Alexa647</td><td>3.95</td><td>162</td><td>300</td><td>d3</td></tr>
<tr><td rowspan="2">190529</td><td>4</td><td>ATTO647</td><td>5.65</td><td>0</td><td>300</td><td>d1</td><td rowspan="2">5.46</td><td rowspan="2">0</td></tr>
<tr><td>4</td><td>ATTO647</td><td>5.26</td><td>0</td><td>300</td><td>d3</td></tr>
<tr><td rowspan="2">200704</td><td>2</td><td>Alexa647</td><td>15.55</td><td>0</td><td>300</td><td>d1</td><td rowspan="2">15.51</td><td rowspan="2">0</td></tr>
<tr><td>2</td><td>Alexa647</td><td>15.47</td><td>0</td><td>300</td><td>d2</td></tr>
<tr><td>200708</td><td>4</td><td>Alexa647</td><td>7.90</td><td>0</td><td>200</td><td>d2</td><td>7.90</td><td>0</td></tr>
<tr><td>210623</td><td>2</td><td>Alexa647</td><td>15.60</td><td>0</td><td>100</td><td>d2</td><td>15.60</td><td>0</td></tr>
</table>

Table 5: Calibration curve experiments

| Date (yymmdd) | setup | device | fluor | protein | buffer | calibration curve date (yymmdd) | prewash exposure (ms) | Rmax | button duty cycle |
|---|---|---|---|---|---|---|---|---|---|
| 181114 | 4 | d1 | Alexa | MAX | WGE | 181113 | 250 | 2.36 | N/A |
| 181114 | 4 | d3 | Alexa | Pho4 | WGE | 181113 | 250 | 5.85 | N/A |
| 181206 | 4 | d1 | Alexa | MAX | WGE | 181113 | 200 | 2.93 | 2 s |
| 181206 | 4 | d3 | Alexa | Pho4 | WGE | 181113 | 200 | 5.91 | 2 s |
| 181217 | 4 | d3 | Alexa | Pho4 | WGE | 181113 | 250 | 3.30 | 2 s |
| 190110 | 4 | d3 | Alexa | MAX | WGE | 181113 | 300 | 3.11 | N/A |
| 190503 | 4 | d1 | ATTO | MAX | WGE | 190529 | 200 | 1.87 | 4 s |
| 190510 | 4 | d1 | ATTO | MAX | WGE | 190529 | 300 | 0.80 | N/A |
| 190510 | 4 | d3 | ATTO | MAX | WGE | 190529 | 300 | 1.29 | 1.2 s |
| 200703 | 2 | d1 | Alexa | Pho4 | TBS | 200704 | 250 | 2.34 | N/A |
| 200703 | 2 | d2 | Alexa | Pho4 | TBS | 200704 | 250 | 3.52 | N/A |
| 200708 | 2 | d1 | Alexa | Pho4 | WGE | 200704 | 200 | 3.67 | N/A |
| 200709 | 4 | d2 | Alexa | MAX | WGE | 200708 | 300 | 1.71 | N/A |
| 200710 | 4 | d1 | Alexa | Pho4 | TBS | 200708 | 300 | 2.66 | N/A |
| 200710 | 4 | d2 | Alexa | Pho4 | WGE | 200708 | 300 | 2.91 | 2 s |
| 200731 | 4 | d2 | Alexa | MAX | TBS | 200708 | 500 | 0.28 | N/A |
| 201023 | 4 | d1 | Alexa | eGFP | WGE | 200708 | 250 | 2.785* | N/A |
| 201111 | 4 | d2 | Alexa | eGFP | WGE | 200708 | 250 | 2.785* | N/A |

Table 6: Library 1 experiments

| Date (yymmdd) | setup | device | fluor | protein | buffer | calibration curve date (yymmdd) | prewash exposure (ms) | Rmax | button duty cycle |
|---|---|---|---|---|---|---|---|---|---|
| 191010 | 4 | d3 | Alexa | MAX | WGE | 190515 | 300 | 0.75 | 4 s |
| 191023 | 4 | d3 | Alexa | MAX | WGE | 190515 | 250 | 0.90 | 2 s |
| 191025 | 4 | d1 | Alexa | MAX | WGE | 190515 | 1000 | 0.70 | 1.2 s |
| 191031 | 4 | d1 | Alexa | Pho4 | WGE | 190515 | 300 | 1.76 | 2 s |
| 191031 | 4 | d3 | Alexa | Pho4 | WGE | 190515 | 300 | 1.57 | 2 s |
| 191104 | 4 | d1 | Alexa | Pho4 | WGE | 190515 | 300 | 1.49 | 4 s |
| 201111 | 2 | d1 | Alexa | eGFP | WGE | 200704 | 200 | 1.195* | N/A |
| 201111 | 2 | d3 | Alexa | eGFP | WGE | 200704 | 200 | 1.195* | N/A |

Table 7: Library 2 experiments

| **Date (yymmdd)** | **setup** | **device** | **fluor** | **protein** | **buffer** | **calibration curve date (yymmdd)** | **prewash exposure (ms)** | **Rmax** |
|---|---|---|---|---|---|---|---|---|
| 190325 | 4 | d3 | ATTO | MAX | WGE | 190118 | 250 | 2.11 |
| 190411 | 4 | d3 | ATTO | MAX | WGE | 190118 | 250 | 1.19 |

Table 8: gcPBM-MITOMI calibration experiments

| Date (yymmdd) | Oligo | DNA A260 | GFP threshold | Rmax |
|---|---|---|---|---|
| 190321 | random 5 | 1.35 | 8e+5 | 0.172 |
| 190321 | repeat 14 | 1.65 | 5e+5 | 0.250 |
| 190402 | repeat 14 | 2.11 | 1e+6 | 0.977 |
| 190407 | repeat 14 | 2.20 | 5e+5 | 0.787 |
| 190409 | random 5 | 2.53 | 1.2e+6 | 0.643 |
| 190409 | repeat 14 | 1.90 | 1.2e+6 | 0.951 |
| 190418 | random 5 | 2.73 | 1.2e+6 | 0.405 |
| 190419 | repeat 14 | 2.10 | 1.2e+6 | 0.676 |
| 190423 | random 5 | 2.60 | 2e+6 | 0.632 |
| 190423 | repeat 14 | 2.00 | 1.2e+6 | 0.633 |

Table 9: STAMMP experiments

### 4 Supplemental figures

5 Cartoon schematic of flow channels (blue and green) and control channels (orange) for the devices used here for all MITOMI, STAMMP, and $k$-MITOMI experiments. . 77

6 Cartoon schematic showing a detailed view of microfluidic valves and compartments within 3 unit cells. . . . . 78

7 Cartoon schematic showing experimental pipeline for MITOMI experiments. **Top panel:** eGFP-tagged transcription factor is immobilized under the button valve with the sandwich valve (dark gray) closed. **Middle panel:** Alexa647-labeled dsDNA is solubilized and allowed to bind to immobilized TF. Adjacent unit cells are segregated by closing sandwich valves (dark gray). **Bottom panel:** Finally, button valves (dark gray) are closed to trap TF-DNA interactions. Unbound DNA is washed from the device prior to imaging. . . . . 79

8 Calibration curves quantifying measured median Alexa-647 fluorescence intensity per chamber as a function of labeled DNA concentration. Light gray markers indicate per-chamber intensities, black markers indicate median value across all chambers, error bars indicate standard deviation, and red line shows the best fit linear regression. Fit parameters and exposure times are indicated within each panel. . . . . 80

9 Calibraton curves quantifying measured median ATTO-647 fluorescence intensity per chamber as a function of labeled DNA concentration. Light gray markers indicate per-chamber intensities, black markers indicate median value across all chambers, error bars indicate standard deviation, and red line shows the best fit linear regression. Fit parameters and exposure times are indicated within each panel. . . . . 81

10 Cartoon schematic depicting computational pipeline for normalizing data across experimental replicates. . . . . 82

11 Example binding curves for MAX binding to DNA Library 1 from experiment on 01/10/2019. Markers indicate per-chamber fluorescence intensity ratios (Alexa-647/eGFP); black dashed line indicates fit to a Langmuir isotherm. Binding curves for other experiments are available on OSF (https://osf.io/gbxhz/). . . . . 83

12 Comparisons between $\Delta\Delta$G measurements for MAX binding to DNA Library 1 across 8 different experiments before (blue markers) and after (red markers) linear normalization (see Supplementary Methods). Blue line shows a linear regression to data prior to normalization; blue text indicates pre-normalization regression parameters; black dashed line indicates the 1:1 identity line. . . . . . . . . . . . . . . . . . . . . . . 84
13 Pairwise comparisons of normalized $\Delta\Delta$G values between all 8 experiments quantifying MAX binding to DNA Library 1. . . . . . . . . . . . . . . . . . . . . . . . . . 85
14 Example binding curves for Pho4 binding to DNA Library 1 from experiment on 07/10/2020 (device #2). Markers indicate per-chamber fluorescence intensity ratios (Alexa-647/eGFP); black dashed line indicates fit to a Langmuir isotherm. Binding curves for other experiments are available on OSF (https://osf.io/gbxhz/). . . . . . . 86
15 Comparisons between $\Delta\Delta$G measurements for Pho4 binding to DNA Library 1 across 8 different experiments before (blue markers) and after (red markers) linear normalization (see Supplementary Methods). Blue line shows a linear regression to data prior to normalization; blue text indicates pre-normalization regression parameters; black dashed line indicates the 1:1 identity line. . . . . . . . . . . . . . . . . . . . . . . 87
16 Pairwise comparisons of normalized $\Delta\Delta$G values between all 8 experiments quantifying Pho4 binding to DNA Library 1. . . . . . . . . . . . . . . . . . . . . . . . . . 88
17 Normalized $\Delta\Delta$Gs across all experiments for MAX binding to each sequence from DNA Library 1. Gray dots indicate all measured $\Delta\Delta$Gs across all experiments; red dots indicate overall median. All $\Delta\Delta$Gs are calculated relative to the median affinity for all sequences containing a motif surrounded by random flanking sequences. . . . 89
18 Normalized $\Delta\Delta$Gs across all experiments for Pho4 binding to each sequence from DNA Library 1. Gray dots indicate all measured $\Delta\Delta$Gs across all experiments; red dots indicate overall median. All $\Delta\Delta$Gs are calculated relative to the median affinity for all sequences containing a motif surrounded by random flanking sequences. . . . 90
19 Distribution and box plot of median $\Delta\Delta$Gs for Pho4 (left) and MAX (right) binding all sequences with a motif surrounded by 'long repeats' (Motif + AC/GT repeat 1–5, Motif + GT/AC repeat 1–3, Motif + AT/CG repeat 1–2, Motif + CG/AT repeat 1–2; red, 13 bp repeats), 'short repeats' (Motif + short GT/AC repeat 2–4, Motif + short CG/AT repeat 1, and Motif + short A/T repeat 1; pink, 6–7 bp repeats and 6–7 bp random), or random sequence (Motif + random 1–3; gray, 13 bp random). . . 91
20 Measured $\Delta\Delta$G values for MAX do not depend on surface-immobilized TF concentration. Each panel shows measured $\Delta\Delta$G values *vs.* eGFP button intensities across all experiments for a given DNA Library 1 sequence. Markers indicate mean for all replicates within a given experiment; error bars indicate standard deviation. . . . . . 92
21 Measured $\Delta\Delta$G values for Pho4 do not depend on surface-immobilized TF concentration. Each panel shows measured $\Delta\Delta$G values *vs.* eGFP button intensities across all experiments for a given DNA Library 1 sequence. Markers indicate median for all replicates within a given experiment; error bars indicate standard deviation. . . . 93
22 Measured $\Delta\Delta$G values for DNA Library 1 in tris-buffered saline (TBS, $y$-axis) *vs.* wheat germ extract (WGE, $x$-axis) for MAX (left) and Pho4 (right). Markers indicate median values across all replicates for a given sequence; error bars indicate standard deviation; black dashed line indicates the 1:1 identity line. . . . . . . . . . . . . . . 94

23 $\Delta\Delta$Gs across all experiments for an eGFP-only negative control binding to each sequence from DNA Library 1. Gray dots indicate all measured $\Delta\Delta$Gs across all experiments; red dots indicate overall median. All $\Delta\Delta$Gs are calculated relative to the median affinity for all sequences containing a motif surrounded by random flanking sequences. . . . . . . . . . . . . . . . . . . . . . . . . . . . . . . . . . . . . 95
24 Detailed comparison of measured $\Delta\Delta$Gs *vs.* predicted binding calculated using a position-specific affinity matrix (PSAM) model. **(A)** Measured (blue bars) and predicted (gray bars) binding for MAX interacting with all DNA Library 1 sequences; error bars indicate the standard deviation across all experiments; translucent gray box indicates energy range highlighted at right. **(B)** Measured (yellow bars) and predicted (gray bars) binding for Pho4 interacting with all DNA Library 1 sequences; error bars indicate the standard deviation across all experiments; translucent gray box indicates energy range highlighted at right. . . . . . . . . . . . . . . . . . . . . . 96
25 Example binding curves for MAX binding to DNA Library 2 from experiment on 10/10/2019. Markers indicate per-chamber fluorescence intensity ratios (Alexa-647/eGFP); dashed line indicates fit to a Langmuir isotherm; data that failed quality controls are shown in gray and marked as **CULLED**. Binding curves for other experiments are available on OSF (https://osf.io/gbxhz/). . . . . . . . . . . . . . . . 97
26 Comparisons between $\Delta\Delta$G measurements for MAX binding to DNA Library 2 across 3 different experiments before (blue markers) and after (red markers) linear normalization (see Supplementary Methods). Blue line shows a linear regression to data prior to normalization; blue text indicates pre-normalization regression parameters; black dashed line indicates the 1:1 identity line. . . . . . . . . . . . . . . . . . . . . . 98
27 Pairwise comparisons of normalized $\Delta\Delta$G values between all 3 experiments quantifying MAX binding to DNA Library 2. . . . . . . . . . . . . . . . . . . . . . . . . . 99
28 Example binding curves for Pho4 binding to DNA Library 2 from experiment on 11/04/2019. Markers indicate per-chamber fluorescence intensity ratios (Alexa-647/eGFP); dashed line indicates fit to a Langmuir isotherm; data that failed quality controls are shown in gray and marked as **CULLED**. Binding curves for other experiments are available on OSF (https://osf.io/gbxhz/). . . . . . . . . . . . . . . . 100
29 Comparisons between $\Delta\Delta$G measurements for Pho4 binding to DNA Library 2 across 3 different experiments before (blue markers) and after (red markers) linear normalization (see Supplementary Methods). Blue line shows a linear regression to data prior to normalization; blue text indicates pre-normalization regression parameters; black dashed line indicates the 1:1 identity line. . . . . . . . . . . . . . . . . . . . . . 101
30 Pairwise comparisons of normalized $\Delta\Delta$G values between all 3 experiments quantifying Pho4 binding to DNA Library 2. . . . . . . . . . . . . . . . . . . . . . . . . . 102
31 Normalized $\Delta\Delta$Gs across all experiments for MAX binding to each sequence from DNA Library 2. Gray dots indicate per-chamber $\Delta\Delta$Gs across all experiments; red dots indicate overall median. All $\Delta\Delta$Gs are calculated relative to the median affinity for all sequences containing a motif surrounded by random flanking sequences. . . . 103
32 Normalized $\Delta\Delta$Gs across all experiments for Pho4 binding to each sequence from DNA Library 2. Gray dots indicate per-chamber $\Delta\Delta$Gs across all experiments; red dots indicate overall median. All $\Delta\Delta$Gs are calculated relative to the median affinity for all sequences containing a motif surrounded by random flanking sequences. . . . 104

33 Measured DNA Library 2 $\Delta\Delta$G values for MAX do not depend on surface-immobilized TF concentration. Each panel shows measured $\Delta\Delta$G values *vs.* eGFP button intensities across all experiments for a given DNA Library 2 sequence. Markers indicate median for all replicates within a given experiment; error bars indicate standard deviation. . . . . . . . . . . . . . . . . . . . . . . . . . . . . . . . . . . . . . . . 105
34 Measured DNA Library 2 $\Delta\Delta$G values for Pho4 do not depend on surface-immobilized TF concentration. Each panel shows measured $\Delta\Delta$G values *vs.* eGFP button intensities across all experiments for a given DNA Library 2 sequence. Markers indicate median for all replicates within a given experiment; error bars indicate standard deviation. . . . . . . . . . . . . . . . . . . . . . . . . . . . . . . . . . . . . . . . 106
35 $\Delta\Delta$Gs across all experiments for an eGFP-only negative control binding to each sequence from DNA Library 2. Gray dots indicate per-chamber $\Delta\Delta$Gs across all experiments; red dots indicate overall median. All $\Delta\Delta$Gs are calculated relative to the median affinity for all sequences containing a motif surrounded by random flanking sequences. . . . . . . . . . . . . . . . . . . . . . . . . . . . . . . . . . . 107
36 DNA shape parameters do not correlate with observed effects on DNA binding affinity for MAX. **(A)** Heat maps displaying calculated minor groove width (MGW), helical twist (HelT), propeller twist (ProT), roll (Roll), and electrostatic potential (EP) as a function of flanking nucleotide position for DNA Library 2 sequences. **(B)** Scatter plots showing measured $\Delta\Delta$Gs *vs.* calculated DNA shape parameters for each DNA Library 2 sequence. Left and right columns show calculations considering either the proximal 5 or 20 nucleotides on either side of the consensus motif, respectively. Sequences in heatmaps are sorted from top to bottom in order of highest to lowest affinity. . . . . . . . . . . . . . . . . . . . . . . . . . . . . . . . . . . . . . . . . . 108
37 DNA shape parameters do not correlate with observed effects on DNA binding affinity for Pho4. **(A)** Heat maps displaying calculated minor groove width (MGW), helical twist (HelT), propeller twist (ProT), roll (Roll), and electrostatic potential (EP) as a function of flanking nucleotide position for DNA Library 2 sequences. **(B)** Scatter plots showing measured $\Delta\Delta$Gs *vs.* calculated DNA shape parameters for each DNA Library 2 sequence. Left and right columns show calculations considering either the proximal 5 or 20 nucleotides on either side of the consensus motif, respectively. Sequences in heatmaps are sorted from top to bottom in order of highest to lowest affinity. . . . . . . . . . . . . . . . . . . . . . . . . . . . . . . . . . . . . . . . . . 109
38 CD spectra for double-stranded DNA sequences containing a central E-box motif surrounded by either random (green lines) or repetitive (blue lines) flanking sequences; representative spectra for a GC hairpin sequence predicted to form Z-DNA are shown in black for comparison. . . . . . . . . . . . . . . . . . . . . . . . . . . . . . . 110

39 Additional EMSAs and quantification of gel band intensities. **(A)** EMSA for sequence containing a motif surrounded by random sequence, imaged in the Cy5 (DNA) channel. Concentration of MAX increases left to right. **(B)** EMSA for sequence containing a motif flanked on one side by GT repeats and on the other by AC repeats, imaged in the Cy5 (DNA) channel. GT/AC repeat flanks are favored over random flanks. Concentration of MAX increases left to right. Red bracket indicates DNA species bound by multiple TFs. **(C)** EMSA for sequence containing flanked on one side by CG repeats and on the other by AT repeats, imaged in the Cy5 (DNA) channel. CG/AT repeat flanks are favored over random flanks. Concentration of MAX increases left to right. Red bracket indicates DNA species bound by multiple TFs. **(D)** MAX-eGFP on a native protein gel runs with 3 bands, likely corresponding to MAX-eGFP dimers, MAX-eGFP monomers, and eGFP alone. Imaged in the eGFP channel. **(E)** Quantification of Alexa-647 signal in gel bands corresponding to MAX-bound DNA. Motif + repeat 1 and Motif + ATGC/ATGC repeat quantification correspond to the EMSAs shown in the main text, **Fig. 2D.** . . . . . . . . . . . 111
40 Binding curves for MAX binding to a selection of fluorescently-labeled dsDNA gcPBM probe sequences from Afek *et al.* [3]. Data from experiment on 03/25/2019. Markers indicate per-chamber fluorescence intensity ratios (Alexa-647/eGFP); dashed line indicates fit to a Langmuir isotherm. . . . . . . . . . . . . . . . . . . . . . . . . . . 112
41 Binding curves for MAX binding to a selection of fluorescently-labeled dsDNA gcPBM probe sequences from Afek *et al.* [3]. Data from experiment on 04/11/2019. Markers indicate per-chamber fluorescence intensity ratios (Alexa-647/eGFP); dashed line indicates fit to a Langmuir isotherm. Binding curves for other experiments are available on OSF (https://osf.io/gbxhz/). . . . . . . . . . . . . . . . . . . . . . . . . . . 113
42 Comparisons between $\Delta\Delta$G measurements for MAX binding to 32 gcPBM dsDNA sequences across 2 experimental replicates before (blue markers) and after (red markers) normalization (see Supplementary Methods). Blue line shows a linear regression to data prior to normalization; blue text indicates pre-normalization regression parameters; black dashed line indicates the 1:1 identity line. . . . . . . . . . . . . . . . 114
43 Pairwise comparisons of normalized $\Delta\Delta$G values between experiments quantifying MAX binding to 32 gcPBM dsDNA sequences. Markers indicate median across all replicates within a given experiment; error bars indicate standard deviation of measured values. . . . . . . . . . . . . . . . . . . . . . . . . . . . . . . . . . . 114
44 Comparison between MAX intensities in a gcPBM experiment *vs.* $\Delta\Delta$Gs measured via MITOMI. Markers indicate median intensities and $\Delta\Delta$G values within a given experiment; error bars indicate standard deviation. Red line indicates a linear regression; returned fit parameters can be used to convert intensities to $\Delta\Delta$Gs for all remaining gcPBM probes. . . . . . . . . . . . . . . . . . . . . . . . . . . . . . . 115
45 Measured binding of MAX (left) and Pho4 (right) binding to DNA Library 2 *vs.* uncalibrated partition function-based predictions of binding. Data are shown as mean ± standard deviation across experimental replicates. Dashed line indicates best fit, with slope and intercept of linear regression used for subsequent calibrates indicated. . . . . . . . . . . . . . . . . . . . . . . . . . . . . . . . . . . . . 116

46 PSAM and partition-function based predictions of binding to DNA Library 1. **(A)** Measured binding of MAX (left) and Pho4 (right) binding to DNA Library 1 *vs.* PSAM-derived binding predictions. Data are shown as mean ± standard deviation across experimental replicates. Dotted line indicates best fit. **(B)** Measured binding of MAX (left) and Pho4 (right) binding to DNA Library 1 *vs.* partition function-derived predictions. Partition function-derived predictions were calibrated using linear regressions in Fig. S45. Data are shown as mean ± standard deviation across experimental replicates. Dotted line indicates best fit. . . . . . . . . . . . . . 117
47 Results of simulation to estimate how many measurements are required for accurate calibration of partition function-derived binding energy predictions. For each value indicated on the $x$-axis, $n$ measurements were randomly sampled and used to calibrate predictions over 10,000 iterations. The rmse of predictions *vs.* measured binding is shown in a boxplot or outlier point for each of 10,000 iterations per $n$ measurements sampled. The rmse calculated from calibrating predictions with all 32 measurements is indicated in blue. An accuracy threshold of 0.5 kcal/mol is indicated in red. . . . . . . . . . . . . . . . . . . . . . . . . . . . . . . . . . . . . . . 118
48 **(A)** Pipeline for simulating effects of repetitive *vs.* random flanking sequences on Gibbs free energy ($\Delta\Delta G$), entropy ($T\Delta S$), and enthalpy ($\Delta H$) as described in Sections 1.13–1.14. **(B)** Simulated entropy distributions as a function of repeat unit; box plots denote distribution median while triangles denote distribution mean. **(C)** Simulated entropy and enthalpy distributions for homopolymer repeat sequences (repeat unit = 1) for sequences with different Z-score ranges. **(D)** Simulated entropy and enthalpy distributions for dinucleotide repeat sequences (repeat unit = 2) for sequences with different Z-score ranges. **(E)** Simulated entropy and enthalpy distributions for tetranucleotide repeat sequences (repeat unit = 4) for sequences with different Z-score ranges. . . . . . . . . . . . . . . . . . . . . . . . . . . . . . . 119
49 Distribution of measured per-chamber eGFP ‘button’ intensities for plasmid-containing (green bars) and empty (gray bars) chambers after on-chip *in vitro* transcription/-translation in 5 STAMMP experiments quantifying binding of eGFP-tagged Pho4 to dsDNA containing an E-box motif surrounded by random sequence. Black dashed line indicates intensity threshold used to identify chambers with expressed Pho4 protein for downstream analysis. . . . . . . . . . . . . . . . . . . . . . . . . . . 120
50 Distribution of measured per-chamber eGFP ‘button’ intensities for plasmid-containing (green bars) and empty (gray bars) chambers after on-chip *in vitro* transcription/-translation in 4 STAMMP experiments quantifying binding of eGFP-tagged Pho4 to dsDNA containing an E-box motif surrounded by CG/AT repeats. Black dashed line indicates intensity threshold used to identify chambers with expressed Pho4 protein for downstream analysis. . . . . . . . . . . . . . . . . . . . . . . . . . . . . . . . 121
51 Example binding curves for STAMMP experiments assessing binding of 214 Pho4 variants passing QC to a DNA sequence containing a CACGTG E-box surrounded by random sequence from experiment on 04/23/2019. Markers indicate per-chamber fluorescence intensity ratios (Alexa-647/eGFP); dashed lines indicates per-chamber fits to a Langmuir isotherm. Binding curves for other experiments are available on OSF (https://osf.io/gbxhz/). . . . . . . . . . . . . . . . . . . . . . . . . . . . . . 122
51 STAMMP binding curves (continued) . . . . . . . . . . . . . . . . . . . . . . 123
51 STAMMP binding curves (continued) . . . . . . . . . . . . . . . . . . . . . . 124

51 STAMMP binding curves (continued) . . . . . . . . . . . . . . . . . . . . . . . . . . . . 125
51 STAMMP binding curves (continued) . . . . . . . . . . . . . . . . . . . . . . . . . . . . 126
52 Comparisons between ΔΔG measurements for 214 Pho4 variants binding to a DNA sequence containing an E-box motif surrounded by random sequence across 5 different experiments before (blue markers) and after (red markers) linear normalization (see Supplementary Methods). Blue line shows a linear regression to data prior to normalization; blue text indicates pre-normalization regression parameters; black dashed line indicates the 1:1 identity line. . . . . . . . . . . . . . . . . . . . . . . . . 127
53 Pairwise comparisons of normalized ΔΔG values between 5 STAMMP experiments quantifying Pho4 mutants binding to a dsDNA sequence containing an E-box motif surrounded by random sequence. Markers indicate median across all replicates within a given experiment; error bars indicate standard deviation of measured values. . . . 128
54 Example binding curves for STAMMP experiments assessing binding of 214 Pho4 variants passing QC to a DNA sequence containing a CACGTG E-box surrounded by CG/AT repeats sequence from experiment on 04/23/2019. Markers indicate per-chamber fluorescence intensity ratios (Alexa-647/eGFP); dashed lines indicates per-chamber fits to a Langmuir isotherm. Binding curves for other experiments are available on OSF (https://osf.io/gbxhz/). . . . . . . . . . . . . . . . . . . . . . . . . 129
54 STAMMP binding curves (continued) . . . . . . . . . . . . . . . . . . . . . . . . . . . . 130
54 STAMMP binding curves (continued) . . . . . . . . . . . . . . . . . . . . . . . . . . . . 131
54 STAMMP binding curves (continued) . . . . . . . . . . . . . . . . . . . . . . . . . . . . 132
54 STAMMP binding curves (continued) . . . . . . . . . . . . . . . . . . . . . . . . . . . . 133
55 Comparisons between ΔΔG measurements for 214 Pho4 variants binding to a DNA sequence containing an E-box motif surrounded by CG/AT repeats across 4 different experiments before (blue markers) and after (red markers) linear normalization (see Supplementary Methods). Blue line shows a linear regression to data prior to normalization; blue text indicates pre-normalization regression parameters; black dashed line indicates the 1:1 identity line. . . . . . . . . . . . . . . . . . . . . . . . . . . . 134
56 Pairwise comparisons of normalized ΔΔG values between 4 STAMMP experiments quantifying Pho4 mutants binding to a dsDNA sequence containing an E-box motif surrounded by GC repeats. Markers indicate median across all replicates within a given experiment; error bars indicate standard deviation of measured values. . . . . 135
57 Calculated residuals **(A)** and residual Z-scores **(B)** from the 1:1 identity line for measured ΔΔGs for all Pho4 mutant variants interacting with DNA sequences containing a central E-box surrounded by either repetitive or random flanking sequences. . . . . 136
58 Example dissociation measurements for MAX interacting with DNA Library 2 sequences from experiment on 10/25/2019. Markers indicate per-chamber fluorescence ratio of Alexa647-labeled dsDNA to GFP-tagged TF; dashed lines indicate single exponential decay fits for data from each chamber. Sequences with no corresponding $K_d$ measurement from this experiment have fit lines shown in red. Dissociation curves for other experiments are available on OSF (https://osf.io/gbxhz/). . . . . . 137

59 Example dissociation measurements for Pho4 interacting with DNA Library 2 sequences from experiment on 11/04/2019. Markers indicate per-chamber TF-bound Alexa-647 intensities; lines indicate single exponential decay fits for data from each chamber. Sequences with no corresponding $K_d$ measurement from this experiment have fit lines shown in red. Dissociation curves for other experiments are available on OSF (https://osf.io/gbxhz/). . . . . . . . . . . . . . . . . . . . . . . . . . . . 138
60 Comparisons between $k_{off}$ measurements for MAX binding to DNA Library 2 across 3 different experiments before and after linear normalization (see Supplementary Methods). Black points with red outline represent median $k_{off}$ across all experiments. 139
61 Comparisons between $k_{off}$ measurements for Pho4 binding to DNA Library 2 across 3 different experiments before (blue markers) and after (red markers) linear normalization (see Supplementary Methods). Black points with red outline represent median $k_{off}$ across all experiments. . . . . . . . . . . . . . . . . . . . . . . . . . . . 139
62 Comparisons between inferred $k_{on}$ values for MAX binding to DNA Library 2 across 3 different experiments before and after linear normalization of $k_{off}$ values (see Supplementary Methods). Black points with red outline represent median $k_{on}$ across all experiments. . . . . . . . . . . . . . . . . . . . . . . . . . . . . . . . . . . . . 140
63 Comparisons between inferred $k_{on}$ values for Pho4 binding to DNA Library 2 across 3 different experiments before and after linear normalization of $k_{off}$ values (see Supplementary Methods). Black points with red outline represent median $k_{on}$ across all experiments. . . . . . . . . . . . . . . . . . . . . . . . . . . . . . . . . . . . . 140
64 Pairwise comparisons of calculated $k_{on}$ values for MAX by sequence across 3 experiments quantifying MAX association rate to DNA Library 2. Markers indicate mean across all replicates within a given experiment; error bars indicate standard deviation of measured values. Lower bounds of error bars adjusted for log scale. . . . . . . . 141
65 Pairwise comparisons of calculated $k_{on}$ values for Pho4 by sequence across 3 experiments quantifying Pho4 association rate to DNA Library 2. Markers indicate mean across all replicates within a given experiment; error bars indicate standard deviation of measured values. . . . . . . . . . . . . . . . . . . . . . . . . . . . . . . . . . . . 142
66 Average measured $k_{off}$ (circle markers, left axis) and calculated $k_{on}$ (diamond markers, right axis) values *vs.* measured affinities ($K_d$s) for Pho4 (yellow/orange) and MAX (blue) interacting with all sequences from DNA Library 2. . . . . . . . . . . 143
67 Quantification of MITOMI button valve opening kinetics demonstrates that button valves take approximately 1 second to open completely. As described in Section 1.16.5, 16 button valves were imaged continuously for 1 second at 1000 frames per second after sending a near-instantaneous signal to depressurize valves. Relative mean intensity is a measure of button state, where a value of 1 corresponds to a completely open button valve and a value of 0 corresponds to a completely closed valve. Each point is a measurement for a single button valve in a single imaging frame. 144

68 Measured $k_{off}$ values for MAX dissociating from oligonucleotides in DNA Library 2 *vs.* 1/button pulse, where button pulse is the length of time button valves are left open and oligos are allowed to dissociate. 1 second is subtracted from the programmed button pulse time to account for button opening delay, as shown in Figure S67. Points and error bars show median $k_{off}$ fits ± standard deviation for a given oligo per experiment. Red dashed lines show linear fits, where the $y$-intercept (indicated in top-left corner of each plot) is interpreted as the 'true' $k_{off}$ value in the absence of mechanical shear. This normalization procedure is described in Section 1.16.5. . . . . . . . . . . . . . . . . . . . . . . . . . . . . . . . . . . . . . . . 145
69 Measured $k_{off}$ values for Pho4 dissociating from oligonucleotides in DNA Library 2 *vs.* 1/button pulse, where button pulse is the length of time button valves are left open and oligos are allowed to dissociate. 1 second is subtracted from the programmed button pulse time to account for button opening delay, as shown in Figure S67. Points and error bars show median $k_{off}$ fits ± standard deviation for a given oligo per experiment. Red dashed lines show linear fits, where the $y$-intercept (indicated in top-left corner of each plot) is interpreted as the 'true' $k_{off}$ value in the absence of mechanical shear. This normalization procedure is described in Section 1.16.5. . . . . . . . . . . . . . . . . . . . . . . . . . . . . . . . . . . . . . . . 146
70 Measured $k_{off}$ (left, circle markers) and calculated $k_{on}$ (right, diamond markers) values as a function of flanking sequence for Pho4 (yellow) and MAX (blue) using an alternative normalization procedure described in Section 1.16.5. . . . . . . . . . . 147
71 Different normalization strategies yield nearly identical inferred $k_{on}$ values for MAX (left) and Pho4 (right) binding to DNA Library 2. $x$-axis shows $k_{on}$ values calculated from normalization method described in Section 1.16.4 and shown in main text Fig. 4C. $y$-axis shows $k_{on}$ values calculated from normalization method described in Section 1.16.5 and shown in Fig. S70. . . . . . . . . . . . . . . . . . . . . . . . . . 148
72 Example dissociation measurements for MAX interacting with DNA Library 1 sequences from experiment on 12/06/2018. Markers indicate per-chamber fluorescence ratio of Alexa647-labeled dsDNA to GFP-tagged TF; lines indicate single exponential decay fits for data from each chamber. Sequences with no corresponding $K_d$ measurement from this experiment have fit lines shown in red. Dissociation curves for other experiments are available on OSF (https://osf.io/gbxhz/). . . . . . . . . . . 149
73 Example dissociation measurements for Pho4 interacting with DNA Library 1 sequences from experiment on 12/17/2018. Markers indicate per-chamber TF-bound Alexa-647 intensities; lines indicate single exponential decay fits for data from each chamber. Sequences with no corresponding $K_d$ measurement from this experiment have fit lines shown in red. Dissociation curves for other experiments are available on OSF (https://osf.io/gbxhz/). . . . . . . . . . . . . . . . . . . . . . . . . . . 150
74 Comparisons between $k_{off}$ measurements for MAX binding to DNA Library 1 across 3 different experiments before and after linear normalization (see Supplementary Methods). Black points with red outline represent median $k_{off}$ across all experiments. 151
75 Comparisons between inferred $k_{on}$ values for MAX binding to DNA Library 1 across 3 different experiments before and after linear normalization of $k_{off}$ values (see Supplementary Methods). Black points with red outline represent median $k_{on}$ across all experiments. . . . . . . . . . . . . . . . . . . . . . . . . . . . . . . . . . . . . . . 151

76 Comparisons between $k_{off}$ measurements for Pho4 binding to DNA Library 1 across 3 different experiments before and after linear normalization (see Supplementary Methods). Black points with red outline represent median $k_{off}$ across all experiments. 152
77 Comparisons between inferred $k_{on}$ values for Pho4 binding to DNA Library 1 across 3 different experiments before and after linear normalization of $k_{off}$ values (see Supplementary Methods). Black points with red outline represent median $k_{on}$ across all experiments. . . . . . . . . . . . . . . . . . . . . . . . . . . . . . . . . . . . . . 152
78 Pairwise comparisons of calculated $k_{on}$ values for MAX by sequence across 3 experiments quantifying MAX association rate to DNA library1. Markers indicate mean across all replicates within a given experiment; error bars indicate standard deviation of measured values. . . . . . . . . . . . . . . . . . . . . . . . . . . . . . . . . . . . 153
79 Pairwise comparisons of calculated $k_{on}$ values for Pho4 by sequence across 3 experiments quantifying Pho4 association rate to DNA Library 1. Markers indicate mean across all replicates within a given experiment; error bars indicate standard deviation of measured values. . . . . . . . . . . . . . . . . . . . . . . . . . . . . . . . . . . . 154
80 Measured vs. model-predicted affinities ($K_d$s, left) and dissociation rates ($k_{off}$s, right) across all sequences for Pho4 (top row) and MAX (bottom row) . . . . . . . . 155
81 Continuous-time Markov Chain (CTMC) model fitting performance for train/test data sets. **(A)** Cartoon schematic of all sequence types with kinetic and equilibrium data used to train and test CTMC model. **(B)** Cartoon schematic of two train/test splits. Left: motif surrounded by random flank used for test (dark gray points in **(C)** and **(D)**). Right: mutated motif (CACGTG) surrounded by random flank used for test (light gray points in **(C)** and **(D)**). **(C)** Performance of model on test data for Pho4 $K_d$ and $k_{off}$. Error bars indicate standard deviation of bootstrapped distribution of fits. Differences between experimental and model values are likely due to known systematic over-estimation of $k_{off}$ and $K_d$ in MITOMI assays. **(D)** Performance of model on test data for MAX $K_d$ and $k_{off}$. Error bars indicate standard deviation of bootstrapped distribution of fits. . . . . . . . . . . . . . . . . . 156
82 Parameters fit from continuous-time Markov Chain model for **(A)** Pho4 and **(B)** MAX. . . . . . . . . . . . . . . . . . . . . . . . . . . . . . . . . . . . . . . . . . 156
83 Example occupancy traces of a genomic binding site containing a motif and either CG repeats (maroon, left column), GT repeats (medium red, middle column), or random flanks (gray, right column) from individual iterations of a Gillespie simulation over 100 s. Each case models 2600 TF molecules. Note that flanking sequences can be occupied by up to 9 TFs at a time, while a motif can be occupied by only a single TF. 157
84 Example traces showing the number of TFs bound to flanking sequences surrounding a motif from individual iterations of a Gillespie simulation over 100 s. Flanking sequences are either CG repeats (maroon, left column), GT repeats (medium red, middle column), or random (gray, right column). Each case models 2600 TF molecules. Note that flanking sequences can be occupied by up to 9 TFs at a time, while a motif can be occupied by only a single TF. . . . . . . . . . . . . . . . . . . . . 158

85 Sensitivity analysis for number of TFs acroos a range of flank/motif affinity ratios in Gillespie simulations. **(A)** Mean first passage time as a function of flank/motif affinity ratio and number of TFs. **(B)** Motif occupancy as a function of flank/motif affinity ratio and number of TFs. **(C)** Flank occupancy as a function of flank/motif affinity ratio and number of TFs. Each depicted value is averaged across 10 iterations of the simulation. . . . . . . . . . . . . . . . . . . . . . . . . . . . . . . . . . . . . . 158
86 Sensitivity analysis for $k_{on}$ across a range of flank/motif affinity ratios in Gillespie simulations. **(A)** Mean first passage time as a function of flank/motif affinity ratio and $k_{on}$. **(B)** Motif occupancy as a function of flank/motif affinity ratio and $k_{on}$. **(C)** Flank occupancy as a function of flank/motif affinity ratio and $k_{on}$. Each depicted value is averaged across 10 iterations of the simulation. . . . . . . . . . . . . . . . . . . 159
87 Supplement to Fig. 4G-I in the main text, where simulations were conducted with $k_{on,max} = 2.67{\times}10^5\ M^{-1}s^{-1}$. Here the same simulations are repeated with $k_{on,max} = 2.67 \times 10^4\ M^{-1}s^{-1}$. **(A)** Log-linear distribution of TF dwell times across 1000 simulations for sequences with a consensus motif flanked by CG repeats, GT repeats, or random sequence; inset shows mean dwell times by sequence. **(B)** Log-linear distribution of the fraction of time a DNA sequence is bound across 1000 simulations for sequences with a consensus motif flanked by GC repeats, GT repeats, or random sequence; inset shows mean time occupied by sequence. **(C)** Mean first passage time (black markers, left axis; units relative to fastest possible search time, 1/(kon,max*[TF])), mean motif occupancy (blue markers, right axis), mean flank occupancy (red markers, right axis), and mean total DNA occupancy (purple markers, right axis) as a function of the likelihood of binding flanking sequence; gray box indicates range of affinities for random flanks; pink and red boxes correspond to fflank values for GT and CG repeats, respectively. . . . . . . . . . . . . . . . . . . . . . . . 160
88 Mean first passage time computed from continuous-time Markov Chain model for Pho4 searching in the yeast nucleus as a function of $f_{flank}$, the likelihood of binding flanking sequence surrounding a motif. . . . . . . . . . . . . . . . . . . . . . . . . . 161
89 Distribution of all repeat sizes in the human genome with mean and median repeat sizes indicated. . . . . . . . . . . . . . . . . . . . . . . . . . . . . . . . . . . . . . . 162
90 AffinityDistillation predictions for the effects of GT repeats on binding (trained on MAX ChIP-seq data) mirror *in vitro* measurements of MAX binding. **(A)** Schematic of sequences with E-box and 15, 30, 45, or 60 bp of favored GT repeats, measured in DNA Library 2. **(B)** AffinityDistillation-predicted change in log(counts) (blue line, left axis) and -1*MITOMI measured $\Delta\Delta G$s (blue markers, right axis) as a function of repeat length (relative to a sequence with a motif and random flanks). Markers and error bars show median and standard deviation across replicates. **(C)** DeepLIFT interpretations for sequences from **(A)** demonstrate non-uniform and subtle contributions of GT repeats to AffinityDistillation log($counts$) predictions. Gray box indicates motif position. **(D)** Cumulative importance scores as a function of position for sequences with E-box and 15, 30, 45, or 60 bp of GT repeats; gray box indicates motif position. . . . . . . . . . . . . . . . . . . . . . . . . . . . . . . . . . . . . . . 163
91 AffinityDistillation-predicted binding ($\Delta \log(counts)$) *vs.* partition-function predicted $\Delta\Delta G$s for 26 DNA sequences containing either an intact motif, a mutated motif, or scrambled sequence surrounded by either repetitive (red markers) or random (gray markers) flanking sequence. . . . . . . . . . . . . . . . . . . . . . . . . . . . . . . . . 164

92 Heatmap showing the intensity Z-score for 1291 TFs interacting with 39 repeat types with TFs ordered by family. . . . . . . . . . . . . . . . . . . . . . . . . . . . . . . . . . 165
93 Distribution of the maximum repeat median intensity Z score across all 1291 TFs analyzed. . . . . . . . . . . . . . . . . . . . . . . . . . . . . . . . . . . . . . . . . . 165
94 Median 8-mer intensity Z score distributions for uPBM experiments across 4 representative TFs (Pho4, Nrg1, Gata3, and Hoxa1). Background intensity Z-score distributions (left) were well-fit to a Gaussian centered around 0; 8-mer Z-scores with p-value less than Bonferroni-corrected threshold of 0.05/39 = 0.0013 (indicated by the solid red light, right) were considered to be bound significantly above background. 39 is the number of repeat types and therefore the number of hypothesis tests performed. . . . . . . . . . . . . . . . . . . . . . . . . . . . . . . . . . . . . . . 166
95 Number (top) and proportion (bottom) of TFs within TF families that show stronger binding to repeats than would be expected purely by chance. . . . . . . . . . . . . . 167
96 Distributions showing the maximum intensity Z score for a repetitive sequence *vs.* the maximum intensity Z score for all sequences for 18 different TF families; markers are colored according to the Levenshtein distance between the highest-scoring repeat and the highest-scoring sequence. . . . . . . . . . . . . . . . . . . . . . . . . . . 168
97 Repeats with lowest 8-mer Z-scores for each of 1,291 TFs. Mononucleotide repeats are shown in red, dinucleotide repeats are shown in yellow, and tetranucleotide repeats are shown in blue. All other repeats with fewer than 10 TFs are grouped into "Other." 169
98 Pairwise comparison of motif and STR preferences, as measured by TF binding on universal PBMs, across basic helix-loop-helix (bHLH) paralogs within 4 different species. Here, we define paralogs as all TFs of the same structural class within a single species. Each point indicates the cosine similarity of motifs and of repeat preferences for a pairwise paralog comparison, with higher values indicating greater similarity. Each subplot depicts paralogs within a different species. . . . . . . . . . . 169
99 Pairwise comparison of motif and STR preferences, as measured by TF binding on universal PBMs, across basic leucine zipper (bZIP) paralogs within 7 different species. Here, we define paralogs as all TFs of the same structural class within a single species. Each point indicates the cosine similarity of motifs and of repeat preferences for a pairwise paralog comparison, with higher values indicating greater similarity. Each subplot depicts paralogs within a different species. . . . . . . . . . . . . . . . . . . . 170
100 Pairwise comparison of motif and STR preferences, as measured by TF binding on universal PBMs, across homeodomain paralogs within 10 different species. Here, we define paralogs as all TFs of the same structural class within a single species. Each point indicates the cosine similarity of motifs and of repeat preferences for a pairwise paralog comparison, with higher values indicating greater similarity. Each subplot depicts paralogs within a different species. . . . . . . . . . . . . . . . . . . . . . . . . 171
101 Pairwise comparison of motif and STR preferences, as measured by TF binding on universal PBMs, across Myb/SANT paralogs within 6 different species. Here, we define paralogs as all TFs of the same structural class within a single species. Each point indicates the cosine similarity of motifs and of repeat preferences for a pairwise paralog comparison, with higher values indicating greater similarity. Each subplot depicts paralogs within a different species. . . . . . . . . . . . . . . . . . . . . . . . . 172

102 Pairwise comparison of motif and STR preferences, as measured by TF binding on universal PBMs, across zinc finger paralogs within 14 different species. Here, we define paralogs as all TFs of the same structural class within a single species. Each point indicates the cosine similarity of motifs and of repeat preferences for a pairwise paralog comparison, with higher values indicating greater similarity. Each subplot depicts paralogs within a different species. . . . . . . . . . . . . . . . . . . . . . . . . 173
103 Pairwise comparison of motif and STR preferences, as measured by TF binding on universal PBMs, across various species and structural classes. Here, we define paralogs as all TFs of the same structural class within a single species. Each point indicates the cosine similarity of motifs and of repeat preferences for a pairwise paralog comparison, with higher values indicating greater similarity. Each subplot depicts paralogs of a specific structural class within a different species. . . . . . . . . 174
104 Pairwise comparison of STR preferences, as measured by TF binding on universal PBMs, across 4 pairs of basic helix-loop-helix (bHLH) paralogs in *Arabidopsis thaliana* with similar motif preferences. Heat maps show 8-mer Z-scores for 39 different repeat types. PWM logos show consensus motif preferences for each TF. . . . 175
105 Pairwise comparison of STR preferences, as measured by TF binding on universal PBMs, across 2 pairs of basic helix-loop-helix (bHLH) paralogs in *Mus musculus* with similar motif preferences. Heat maps show 8-mer Z-scores for 39 different repeat types. PWM logos show consensus motif preferences for each TF. . . . . . . . . . . . 176
106 Comparison of STR preferences, as measured by TF binding on universal PBMs, across 6 pairs nuclear hormone receptor (NHR) paralogs in *Mus musculus* with similar motif preferences. Heat map shows 8-mer Z-scores for 39 different repeat types. PWM logos show consensus motif preferences for each TF. . . . . . . . . . . . . . . . 176
107 Distribution of all repeat Z-scores aggregated across 1,291 TFs. . . . . . . . . . . . . 177

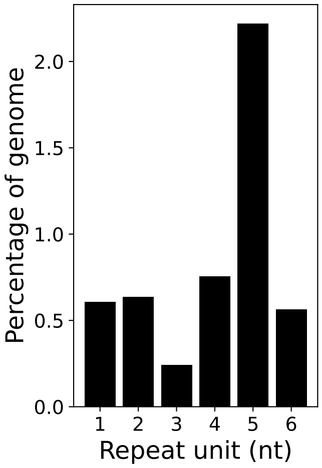

Figure 1: Percentage short tandem repeat (STR) content by repeat unit in the T2T-CHM13 reference genome [38]. The high proportion of 5 nt STRs is due to human satellite 3 (HSat3, [5]).

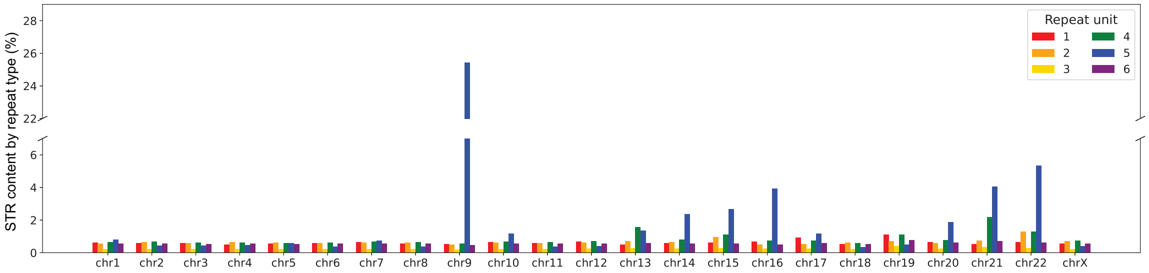

Figure 2: Percentage short tandem repeat (STR) content across each human chromosome by repeat unit in the T2T-CHM13 reference genome [38]. The high proportion of STRs on chromosome 9 is due to human satellite 3 (HSat3, [5]).

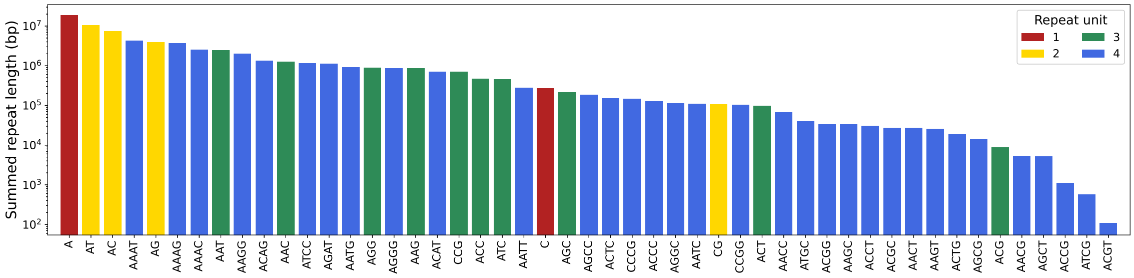

Figure 3: Distribution showing overall length of repeats in the human genome by sequence type. Mononucleotide repeats are shown in red, dinucleotide repeats are shown in yellow, trinucleotide repeats are shown in green, and tetranucleotide repeats are shown in blue.

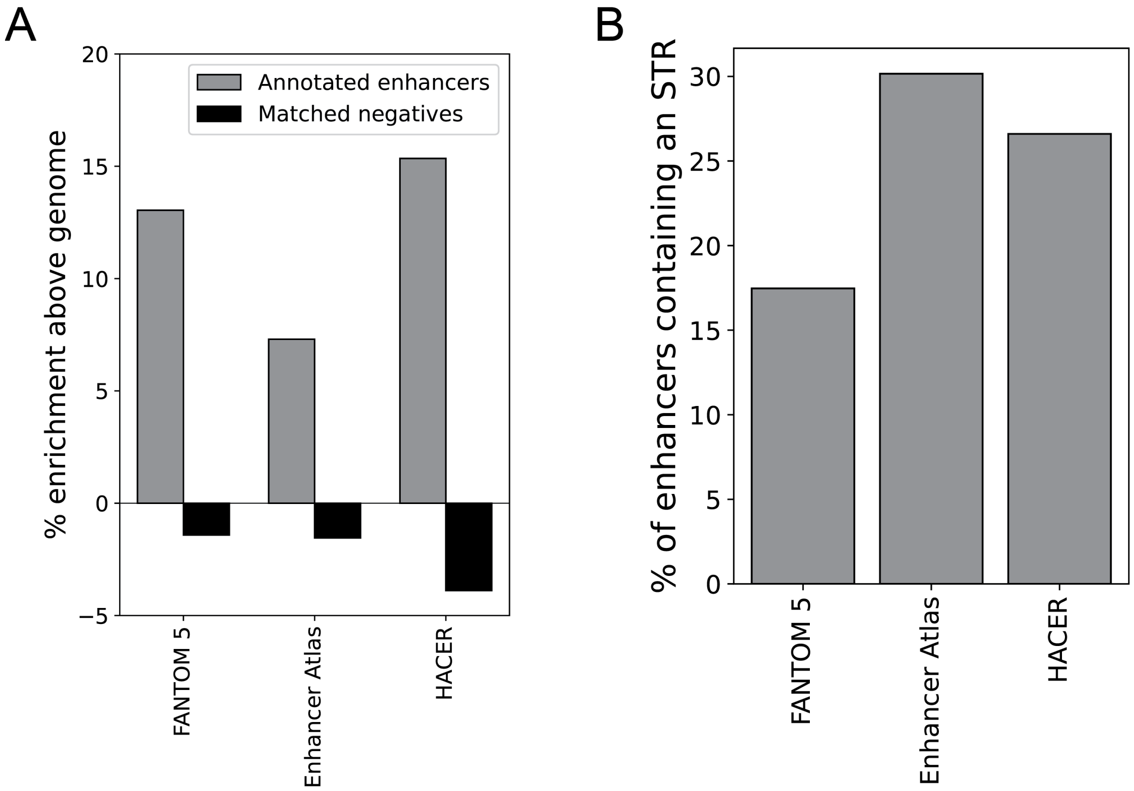

Figure 4: **(A)** enrichment of STRs above genomic background in enhancers (gray) compared to nucleotide composition-matched genomic non-enhancer loci (black). $x$-axis labels indicate source of enhancer annotations. **(B)** percentage of enhancers containing an STR; $x$-axis labels indicate source of enhancer annotations.

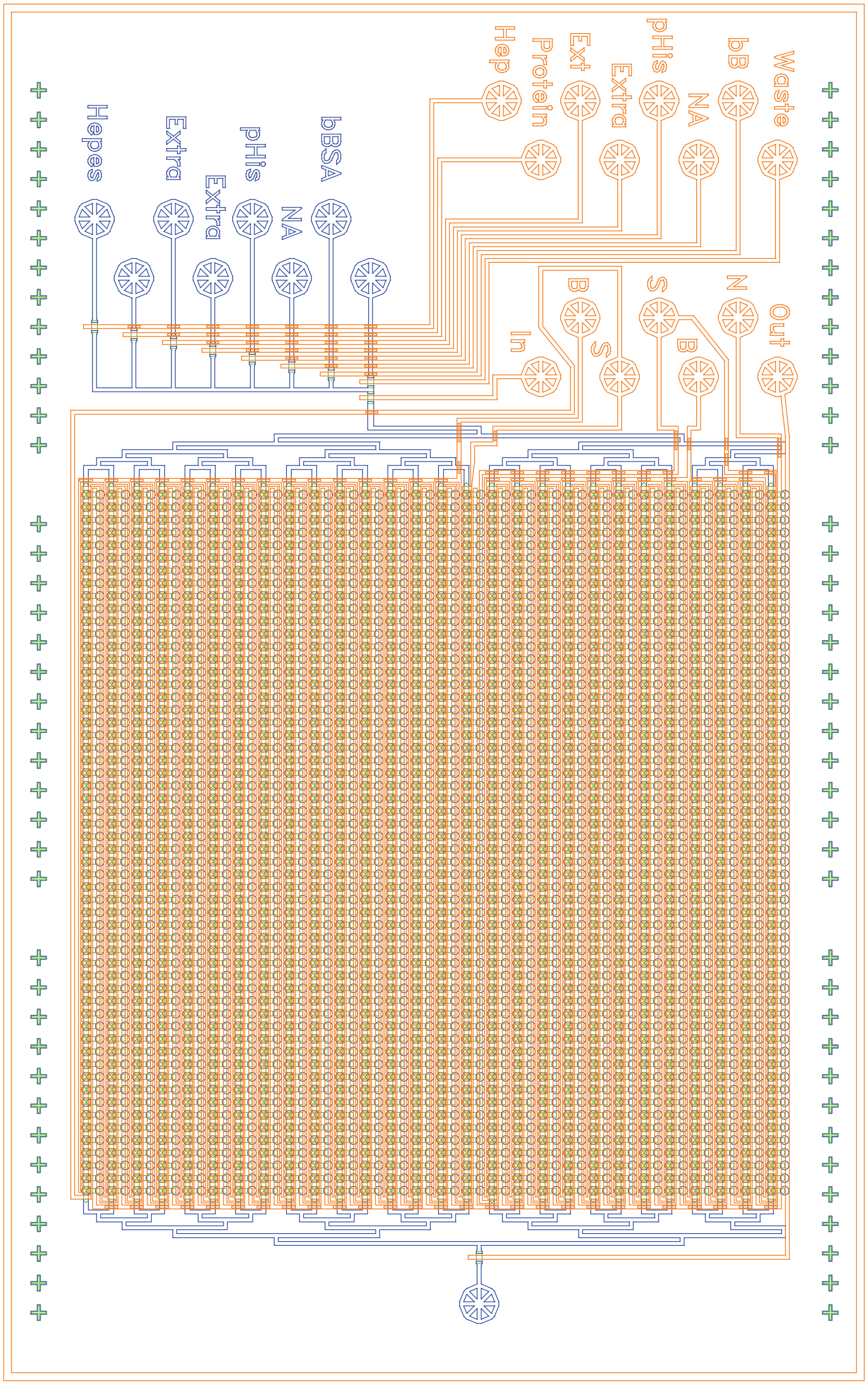

Figure 5: Cartoon schematic of flow channels (blue and green) and control channels (orange) for the devices used here for all MITOMI, STAMMP, and $k$-MITOMI experiments.

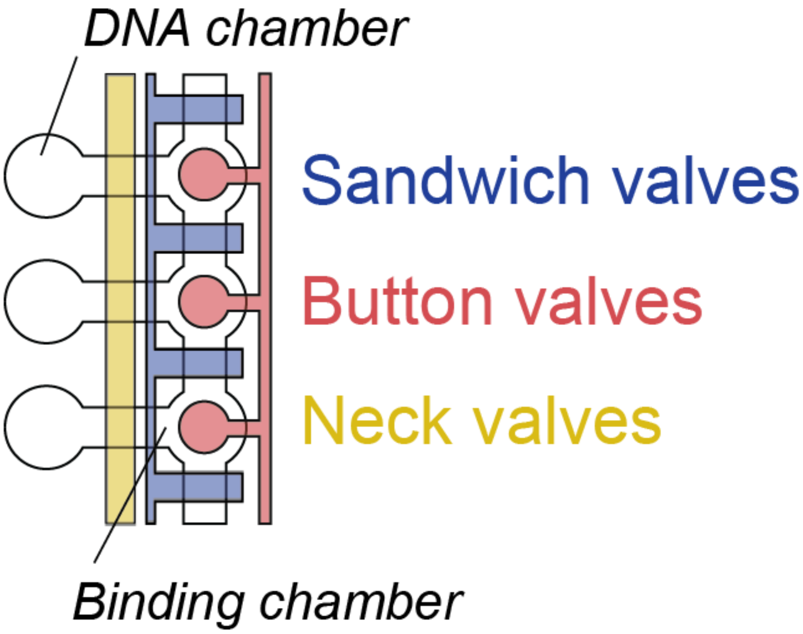

Figure 6: Cartoon schematic showing a detailed view of microfluidic valves and compartments within 3 unit cells.

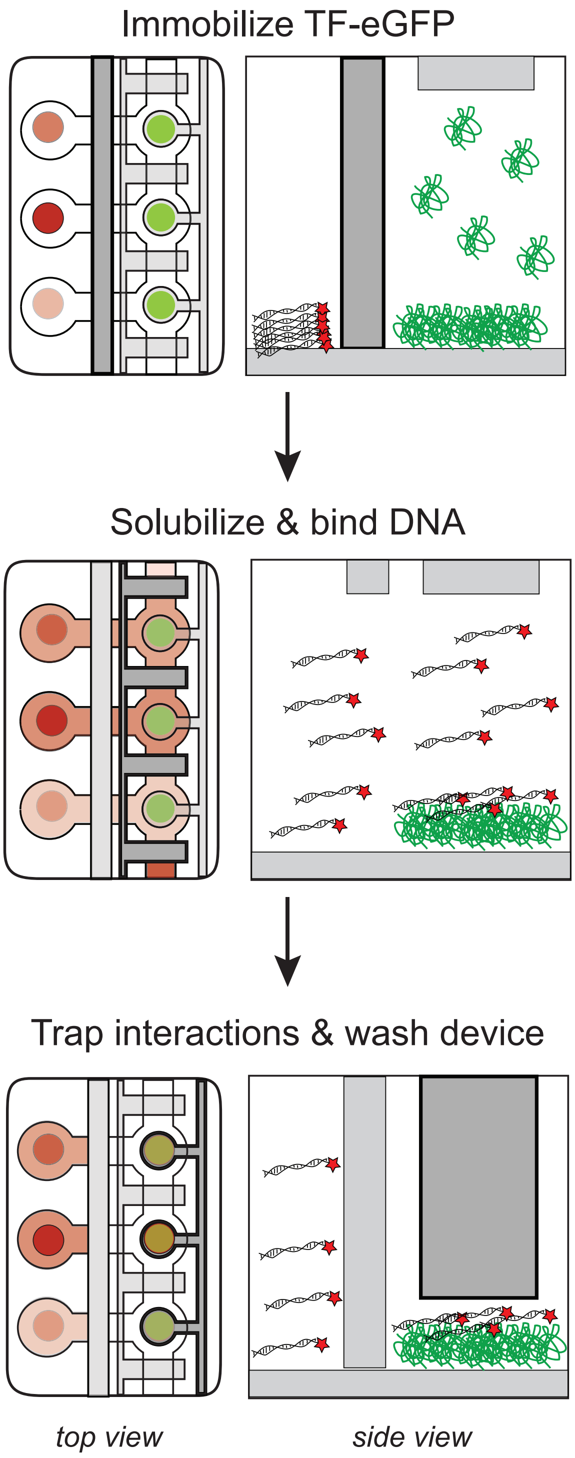

Figure 7: Cartoon schematic showing experimental pipeline for MITOMI experiments. **Top panel:** eGFP-tagged transcription factor is immobilized under the button valve with the sandwich valve (dark gray) closed. **Middle panel:** Alexa647-labeled dsDNA is solubilized and allowed to bind to immobilized TF. Adjacent unit cells are segregated by closing sandwich valves (dark gray). **Bottom panel:** Finally, button valves (dark gray) are closed to trap TF-DNA interactions. Unbound DNA is washed from the device prior to imaging.

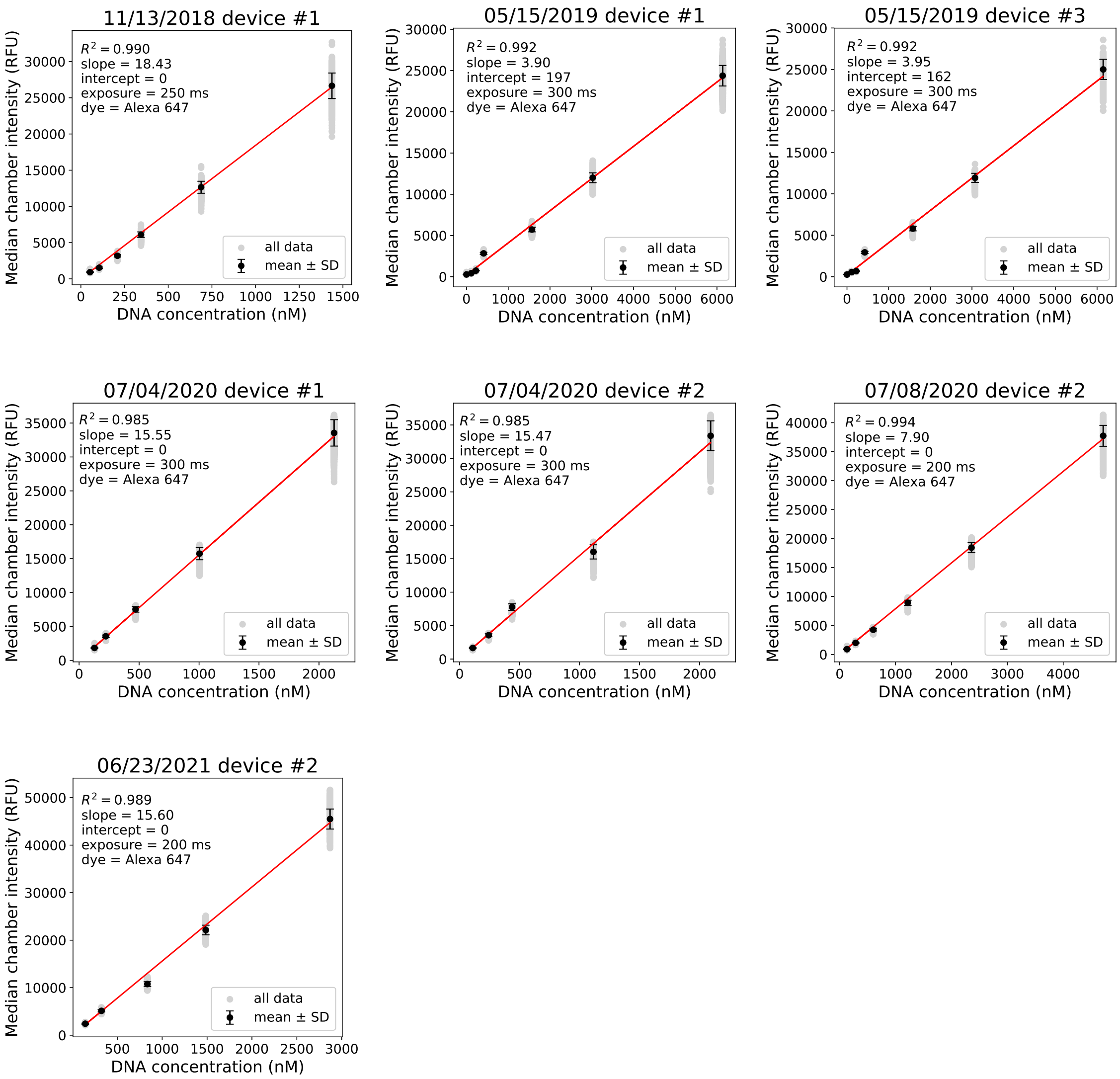

Figure 8: Calibration curves quantifying measured median Alexa-647 fluorescence intensity per chamber as a function of labeled DNA concentration. Light gray markers indicate per-chamber intensities, black markers indicate median value across all chambers, error bars indicate standard deviation, and red line shows the best fit linear regression. Fit parameters and exposure times are indicated within each panel.

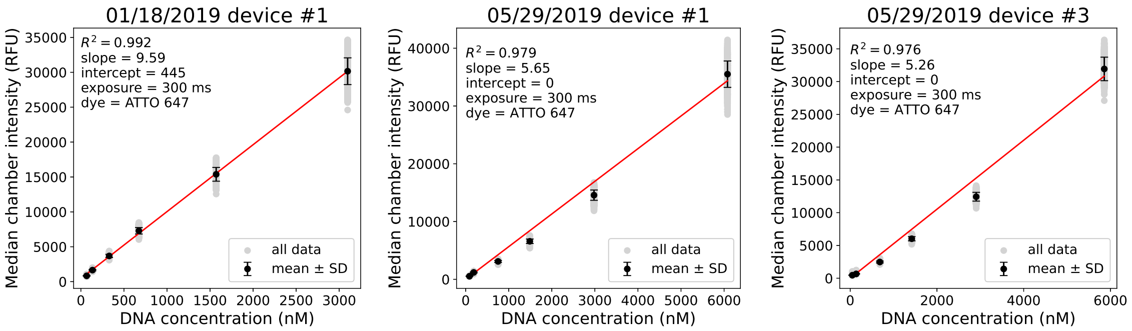

Figure 9: Calibraton curves quantifying measured median ATTO-647 fluorescence intensity per chamber as a function of labeled DNA concentration. Light gray markers indicate per-chamber intensities, black markers indicate median value across all chambers, error bars indicate standard deviation, and red line shows the best fit linear regression. Fit parameters and exposure times are indicated within each panel.

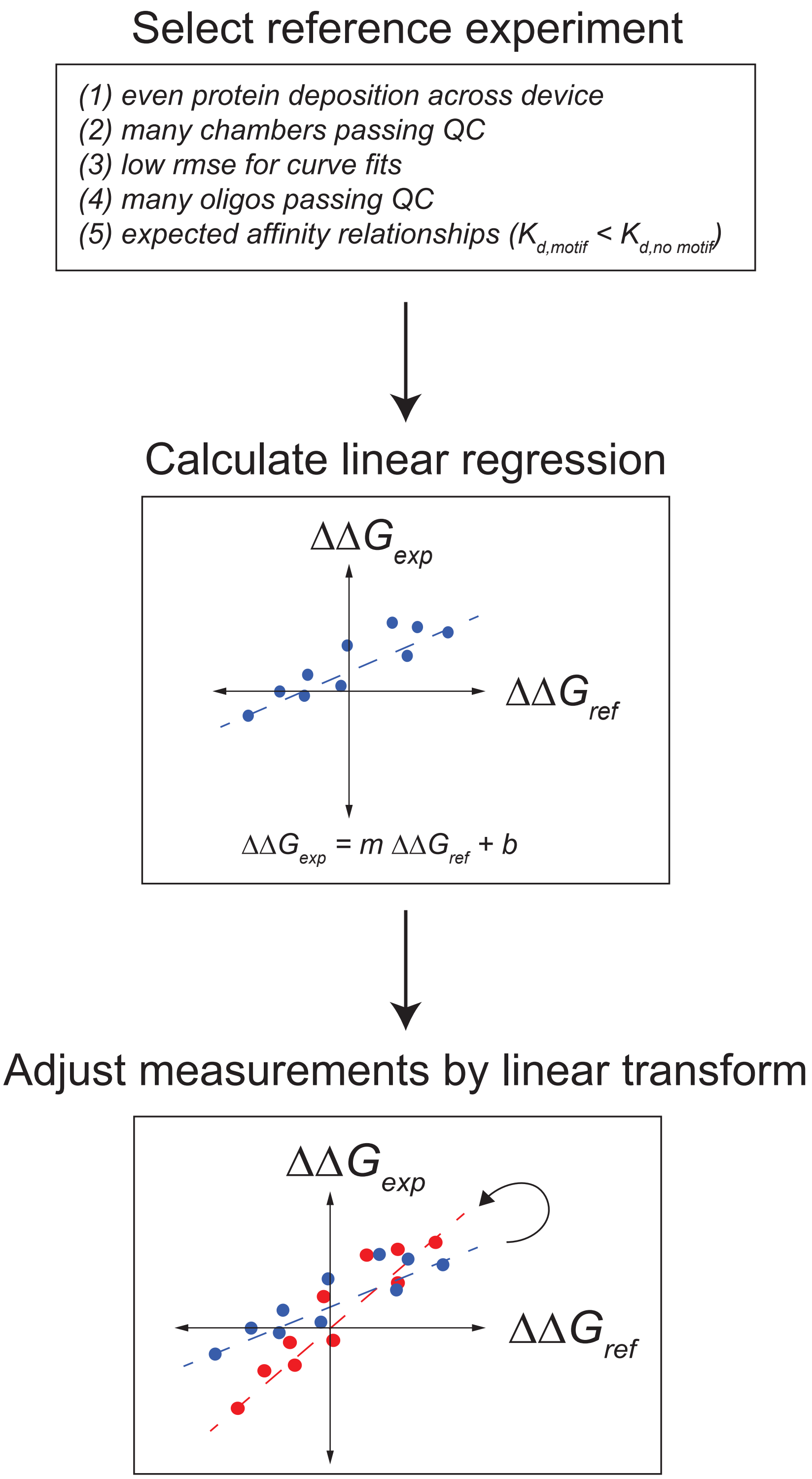

Figure 10: Cartoon schematic depicting computational pipeline for normalizing data across experimental replicates.

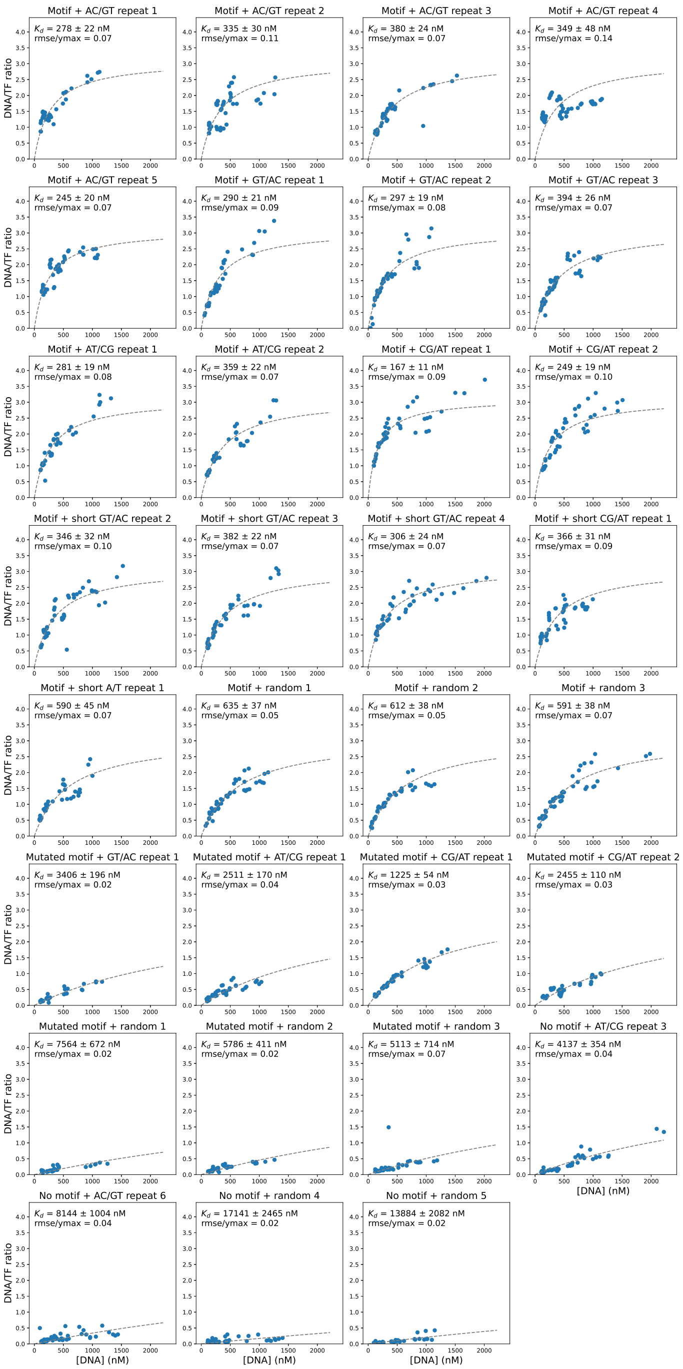

Figure 11: Example binding curves for MAX binding to DNA Library 1 from experiment on 01/10/2019. Markers indicate per-chamber fluorescence intensity ratios (Alexa-647/eGFP); black dashed line indicates fit to a Langmuir isotherm. Binding curves for other experiments are available on OSF (https://osf.io/gbxhz/).

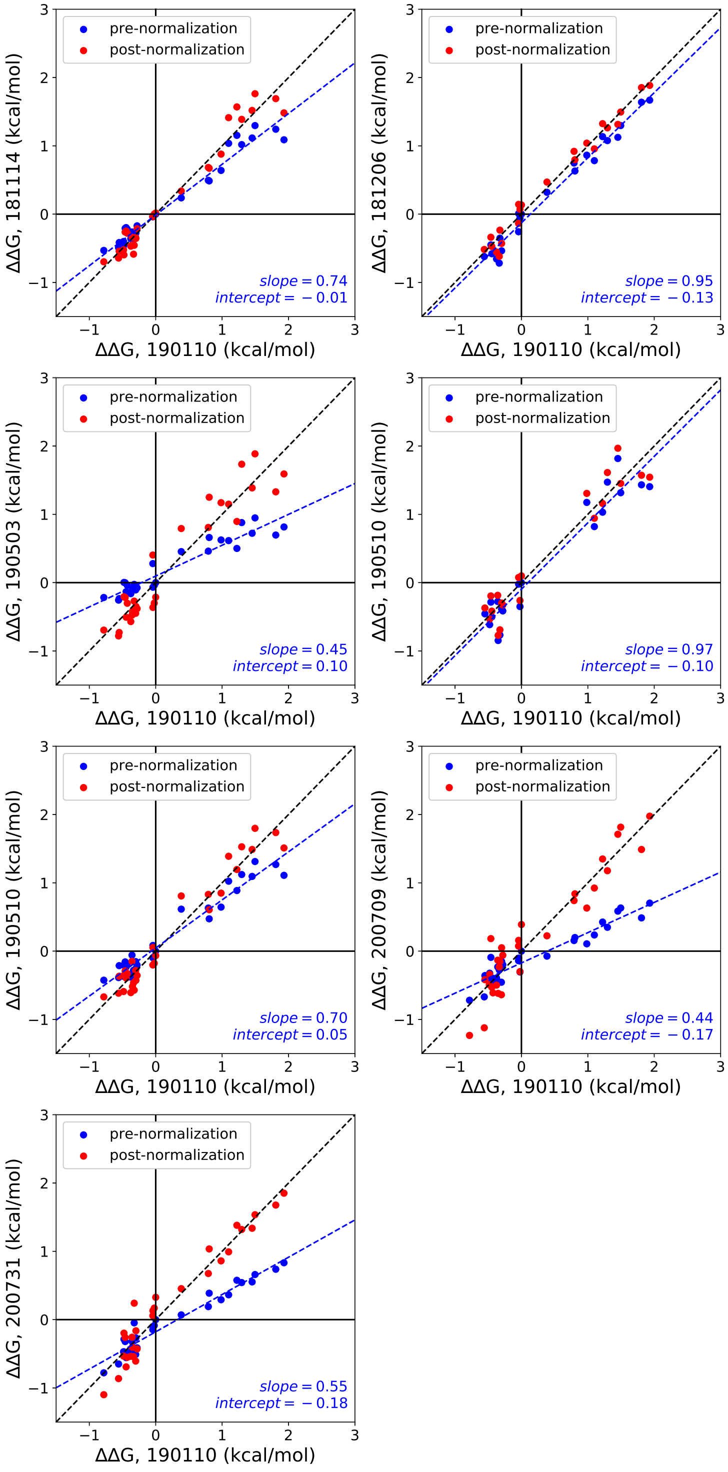

Figure 12: Comparisons between ΔΔG measurements for MAX binding to DNA Library 1 across 8 different experiments before (blue markers) and after (red markers) linear normalization (see Supplementary Methods). Blue line shows a linear regression to data prior to normalization; blue text indicates pre-normalization regression parameters; black dashed line indicates the 1:1 identity line.

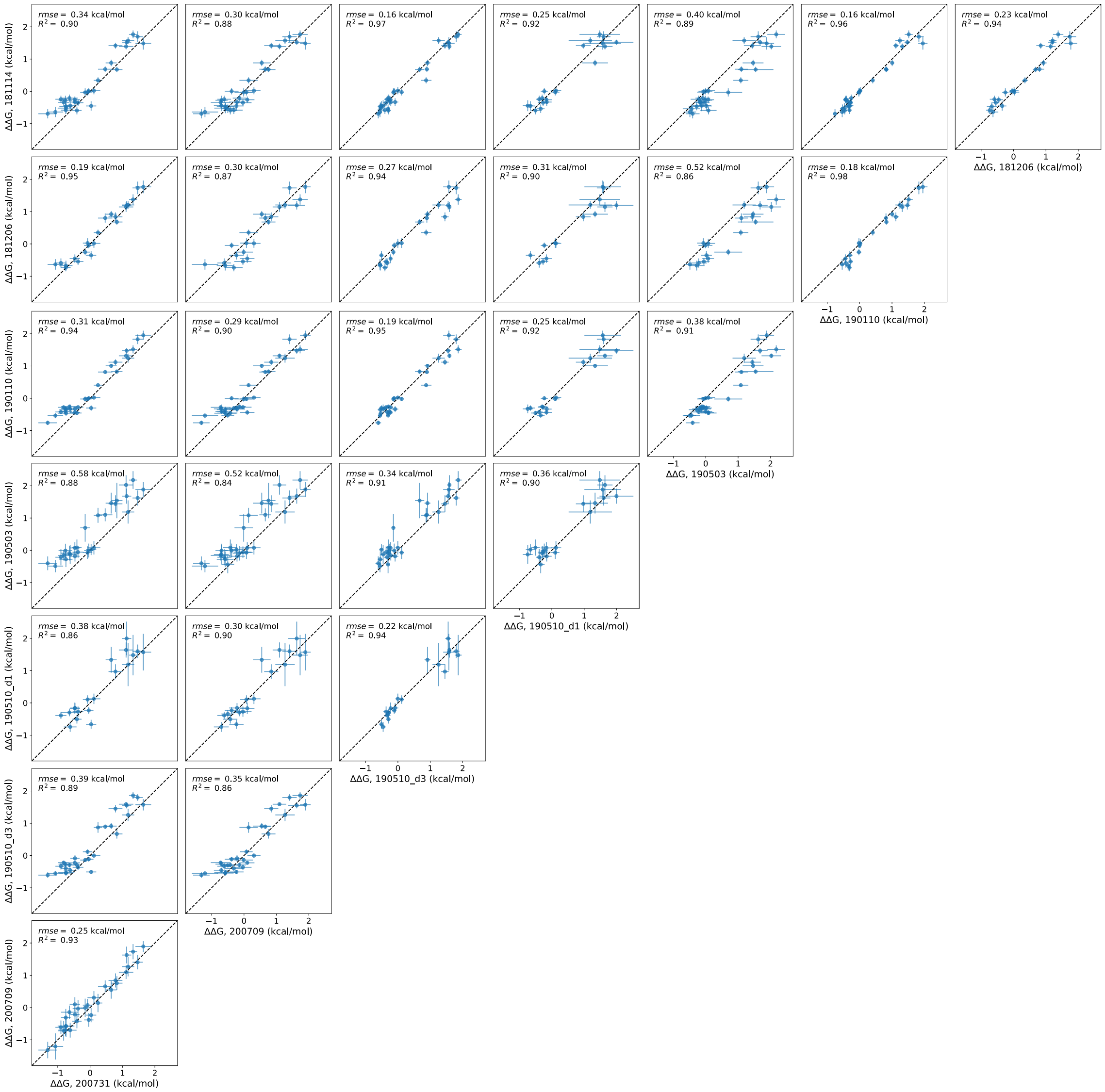

Figure 13: Pairwise comparisons of normalized ΔΔG values between all 8 experiments quantifying MAX binding to DNA Library 1.

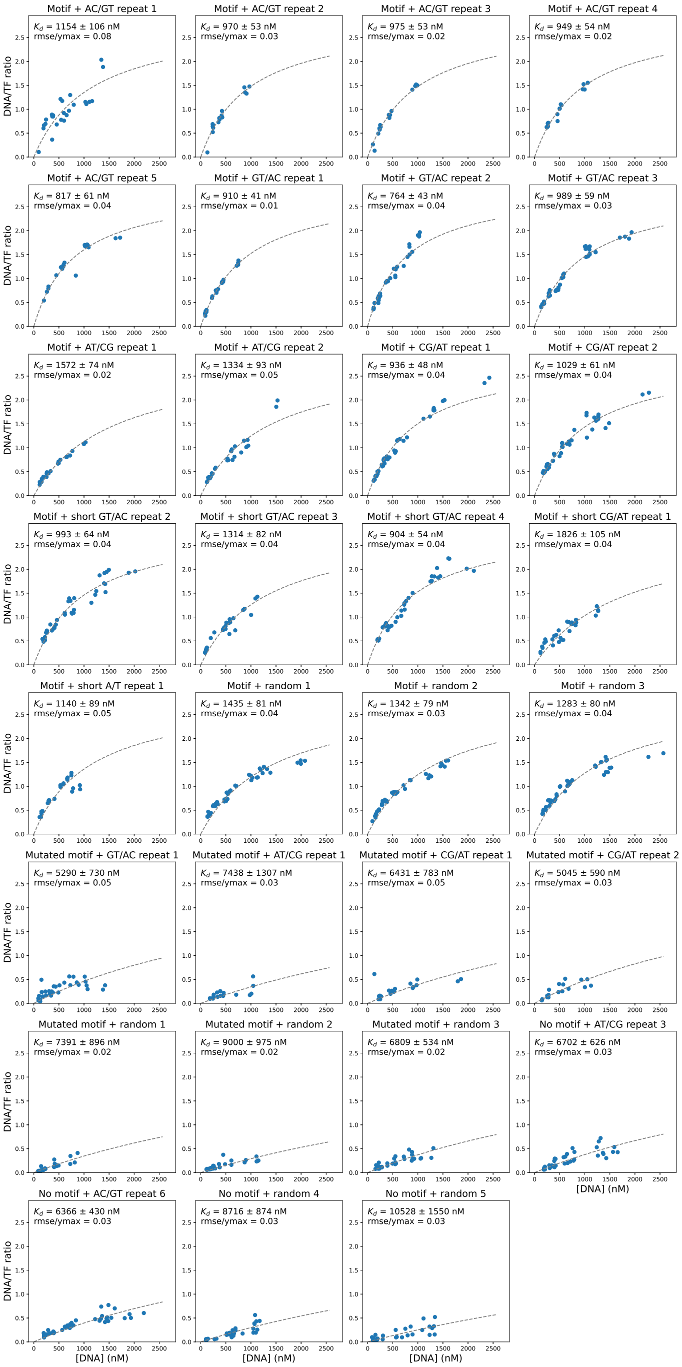

Figure 14: Example binding curves for Pho4 binding to DNA Library 1 from experiment on 07/10/2020 (device #2). Markers indicate per-chamber fluorescence intensity ratios (Alexa-647/eGFP); black dashed line indicates fit to a Langmuir isotherm. Binding curves for other experiments are available on OSF (https://osf.io/gbxhz/).

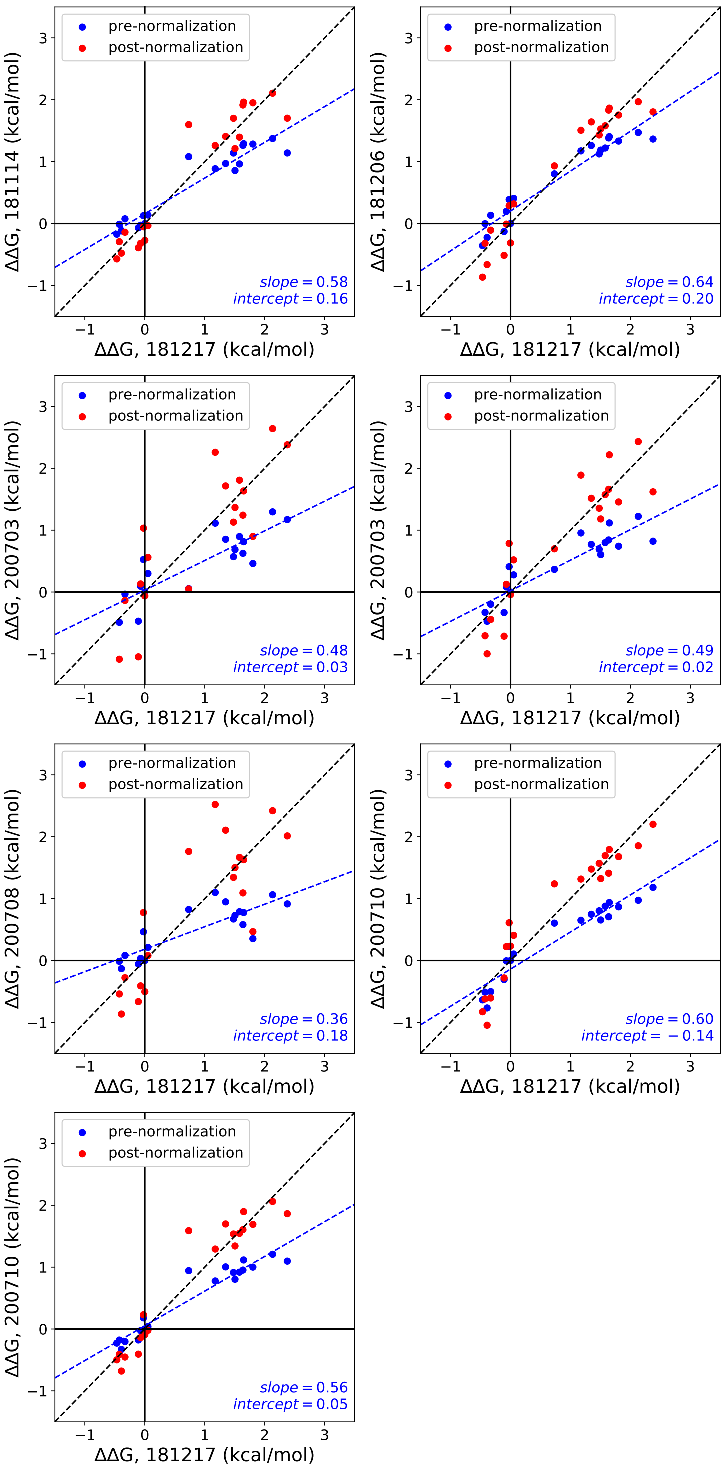

Figure 15: Comparisons between ΔΔG measurements for Pho4 binding to DNA Library 1 across 8 different experiments before (blue markers) and after (red markers) linear normalization (see Supplementary Methods). Blue line shows a linear regression to data prior to normalization; blue text indicates pre-normalization regression parameters; black dashed line indicates the 1:1 identity line.

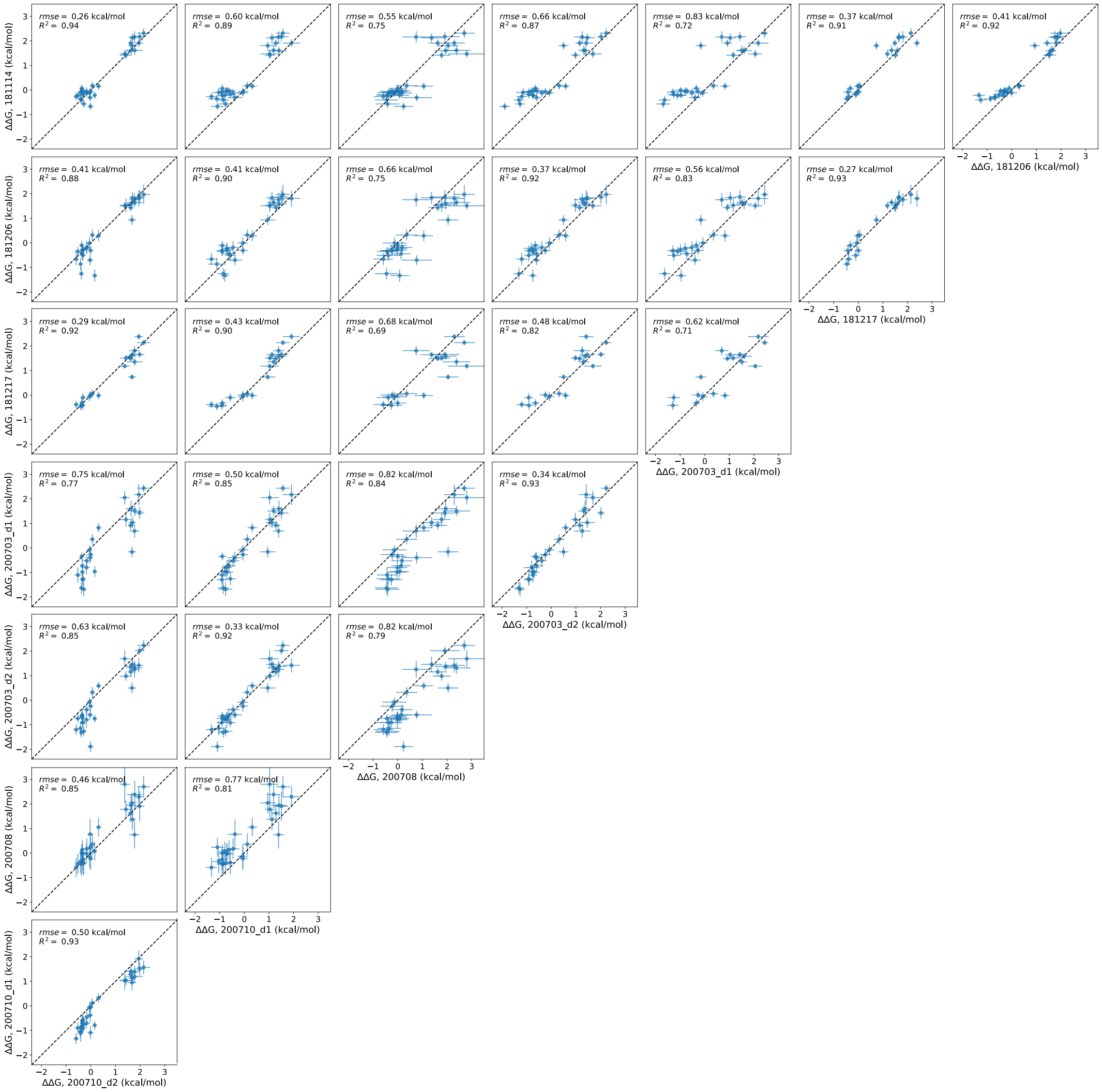

Figure 16: Pairwise comparisons of normalized ΔΔG values between all 8 experiments quantifying Pho4 binding to DNA Library 1.

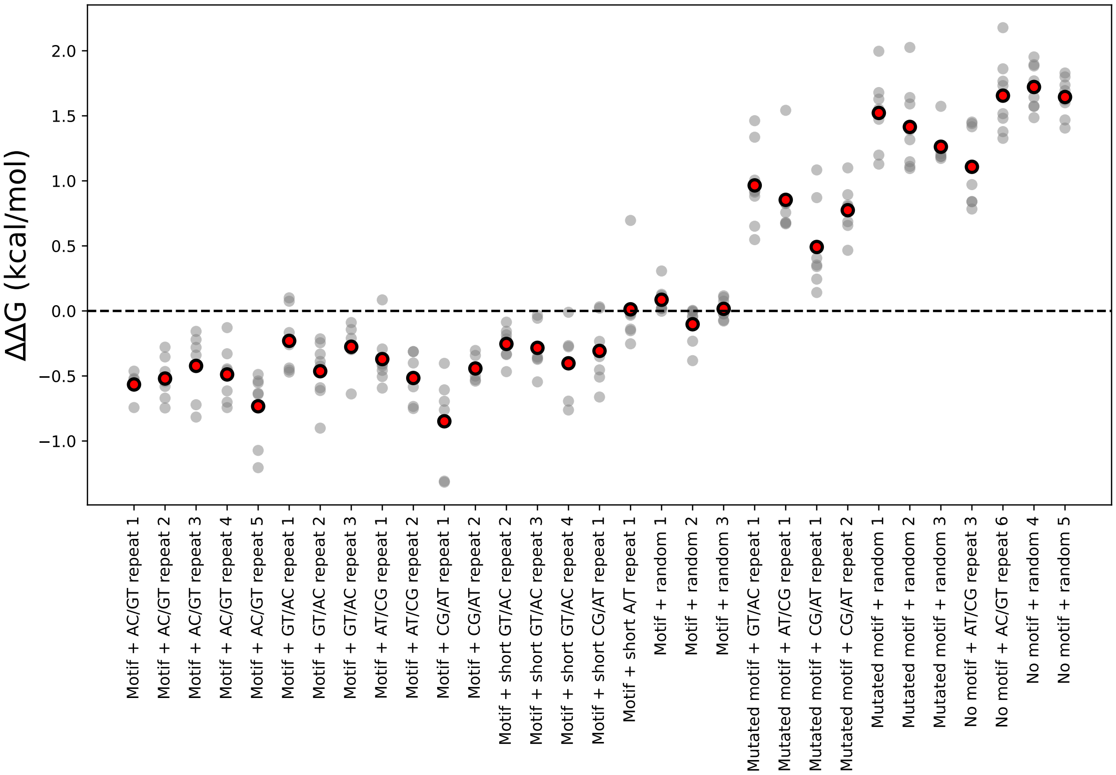

Figure 17: Normalized ΔΔGs across all experiments for MAX binding to each sequence from DNA Library 1. Gray dots indicate all measured ΔΔGs across all experiments; red dots indicate overall median. All ΔΔGs are calculated relative to the median affinity for all sequences containing a motif surrounded by random flanking sequences.

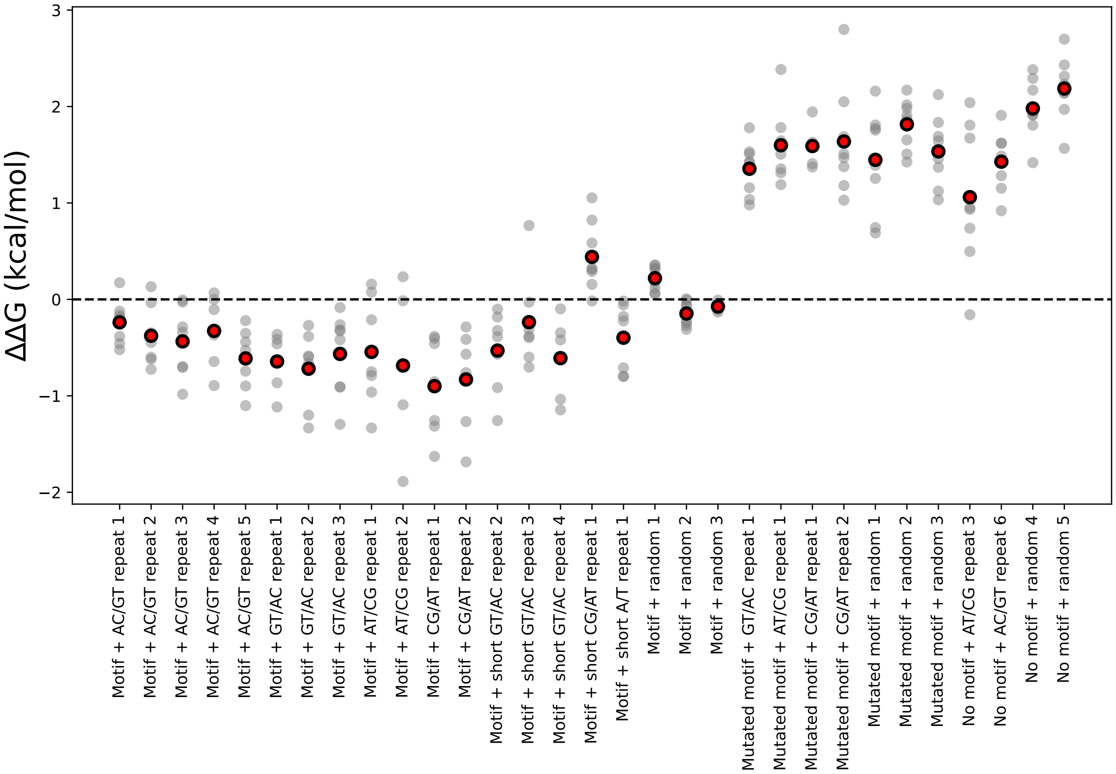

Figure 18: Normalized ΔΔGs across all experiments for Pho4 binding to each sequence from DNA Library 1. Gray dots indicate all measured ΔΔGs across all experiments; red dots indicate overall median. All ΔΔGs are calculated relative to the median affinity for all sequences containing a motif surrounded by random flanking sequences.

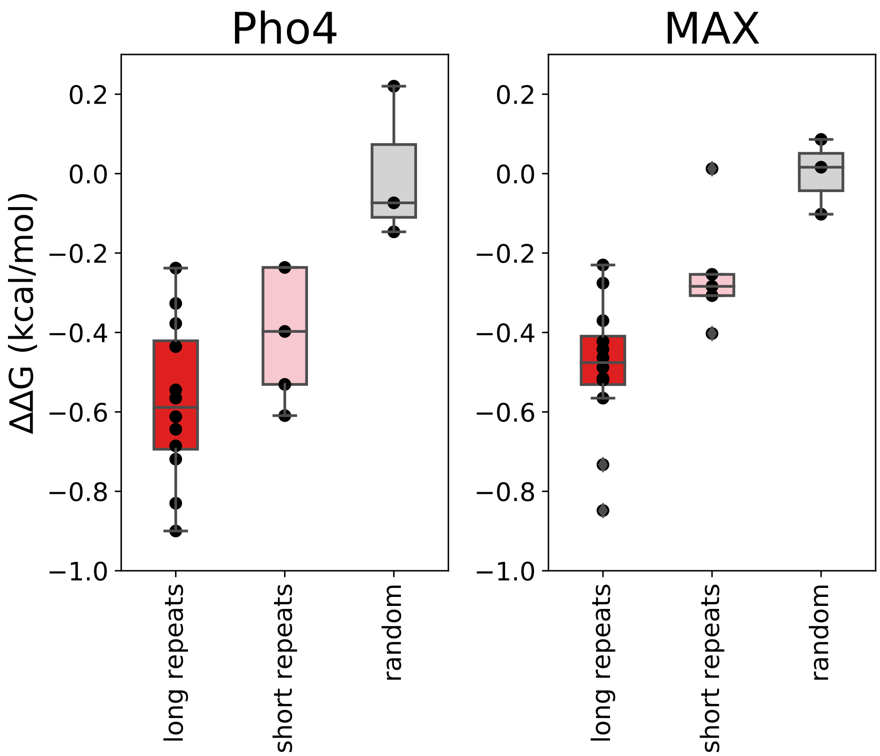

Figure 19: Distribution and box plot of median ΔΔGs for Pho4 (left) and MAX (right) binding all sequences with a motif surrounded by 'long repeats' (Motif + AC/GT repeat 1–5, Motif + GT/AC repeat 1–3, Motif + AT/CG repeat 1–2, Motif + CG/AT repeat 1–2; red, 13 bp repeats), 'short repeats' (Motif + short GT/AC repeat 2–4, Motif + short CG/AT repeat 1, and Motif + short A/T repeat 1; pink, 6–7 bp repeats and 6–7 bp random), or random sequence (Motif + random 1–3; gray, 13 bp random).

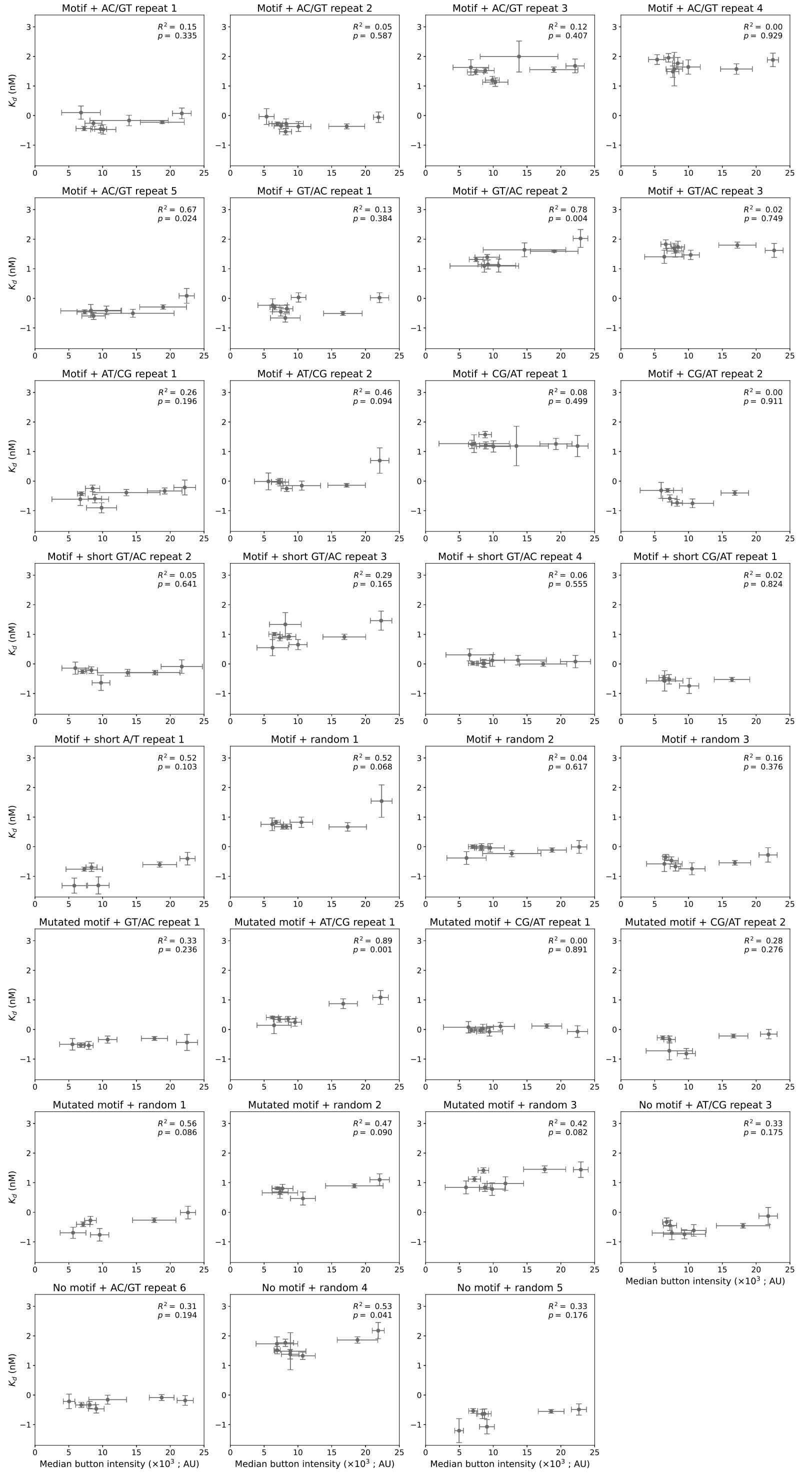

Figure 20: Measured $\Delta\Delta$G values for MAX do not depend on surface-immobilized TF concentration. Each panel shows measured $\Delta\Delta$G values *vs.* eGFP button intensities across all experiments for a given DNA Library 1 sequence. Markers indicate mean for all replicates within a given experiment; error bars indicate standard deviation.

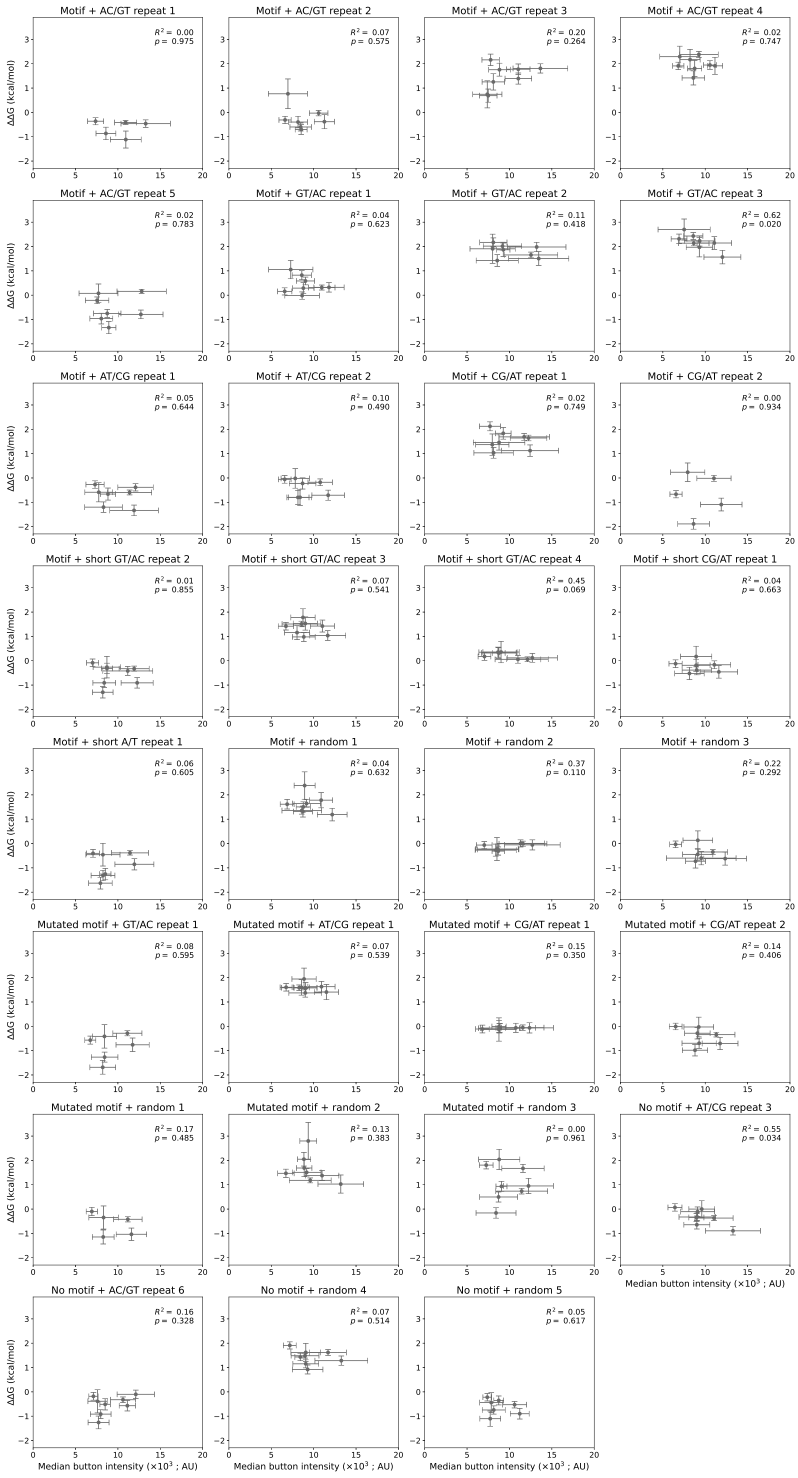

Figure 21: Measured ΔΔG values for Pho4 do not depend on surface-immobilized TF concentration. Each panel shows measured ΔΔG values *vs.* eGFP button intensities across all experiments for a given DNA Library 1 sequence. Markers indicate median for all replicates within a given experiment; error bars indicate standard deviation.

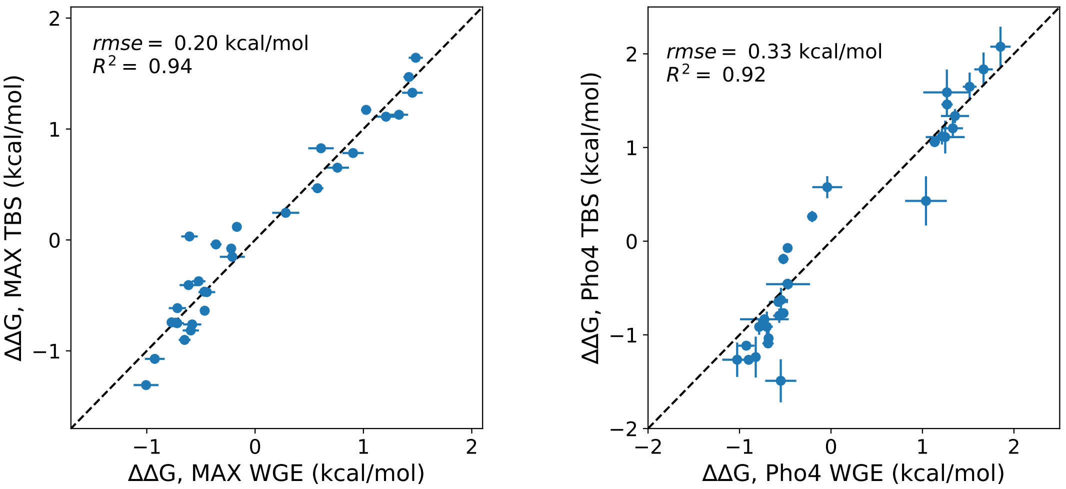

Figure 22: Measured ΔΔG values for DNA Library 1 in tris-buffered saline (TBS, $y$-axis) *vs.* wheat germ extract (WGE, $x$-axis) for MAX (left) and Pho4 (right). Markers indicate median values across all replicates for a given sequence; error bars indicate standard deviation; black dashed line indicates the 1:1 identity line.

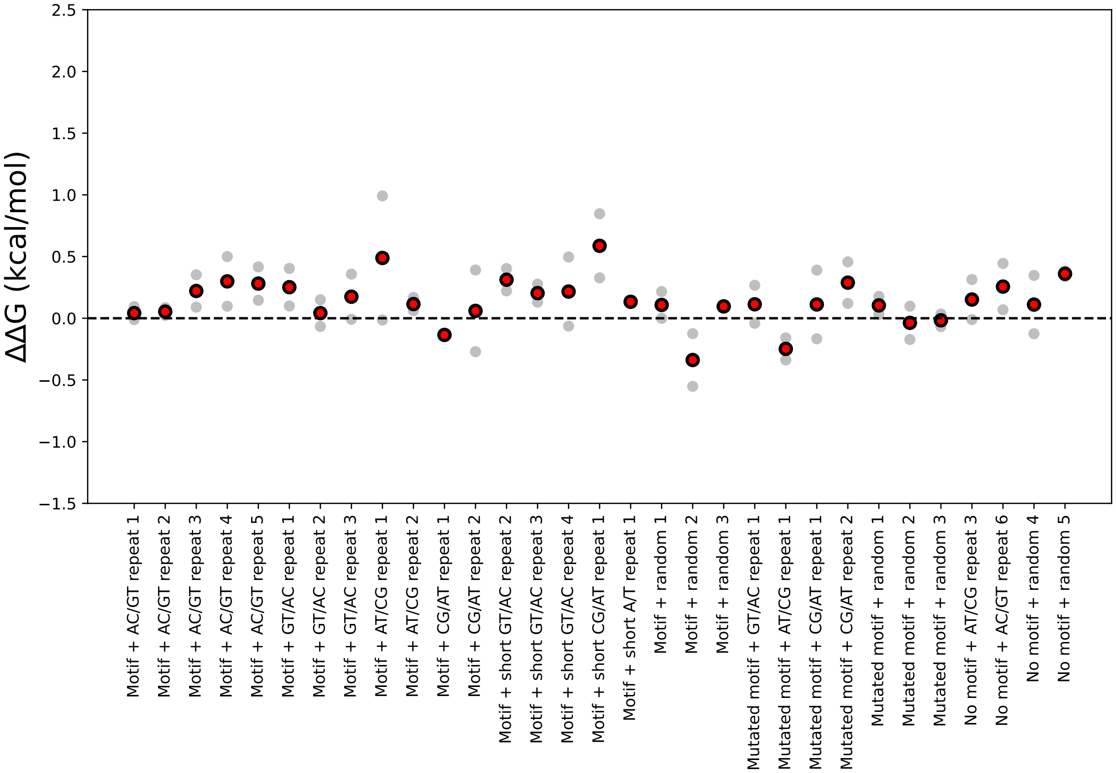

Figure 23: ΔΔGs across all experiments for an eGFP-only negative control binding to each sequence from DNA Library 1. Gray dots indicate all measured ΔΔGs across all experiments; red dots indicate overall median. All ΔΔGs are calculated relative to the median affinity for all sequences containing a motif surrounded by random flanking sequences.

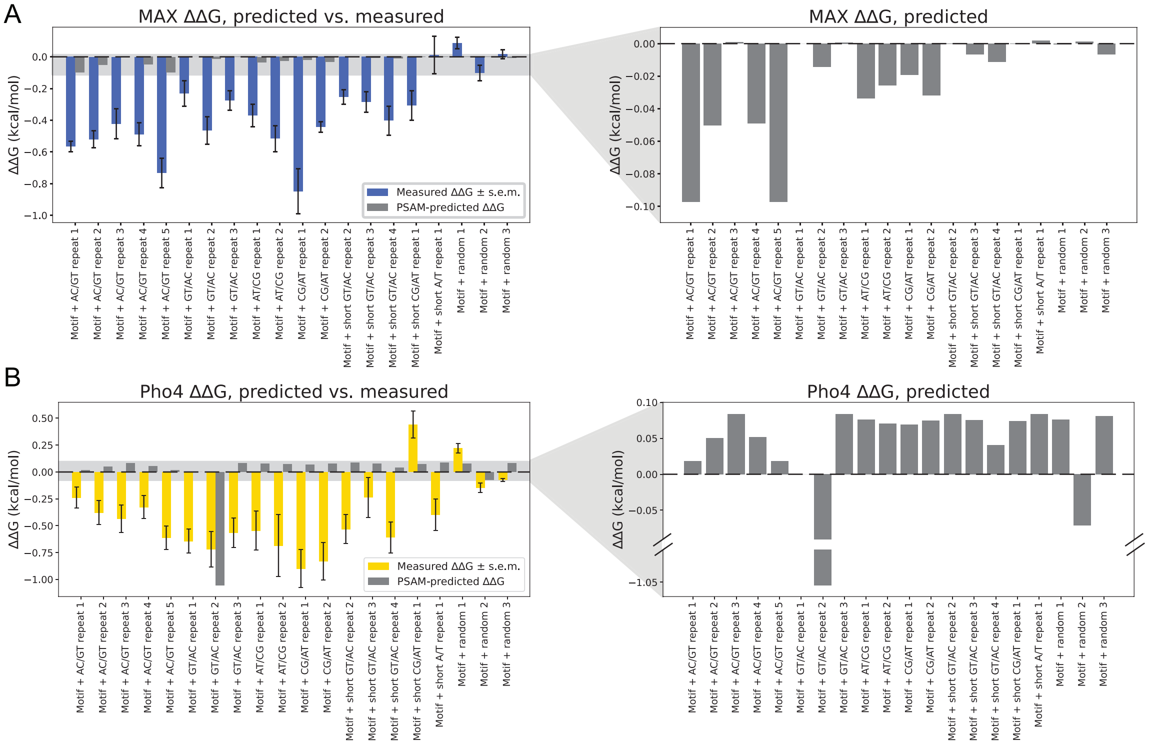

Figure 24: Detailed comparison of measured ΔΔGs *vs.* predicted binding calculated using a position-specific affinity matrix (PSAM) model. **(A)** Measured (blue bars) and predicted (gray bars) binding for MAX interacting with all DNA Library 1 sequences; error bars indicate the standard deviation across all experiments; translucent gray box indicates energy range highlighted at right. **(B)** Measured (yellow bars) and predicted (gray bars) binding for Pho4 interacting with all DNA Library 1 sequences; error bars indicate the standard deviation across all experiments; translucent gray box indicates energy range highlighted at right.

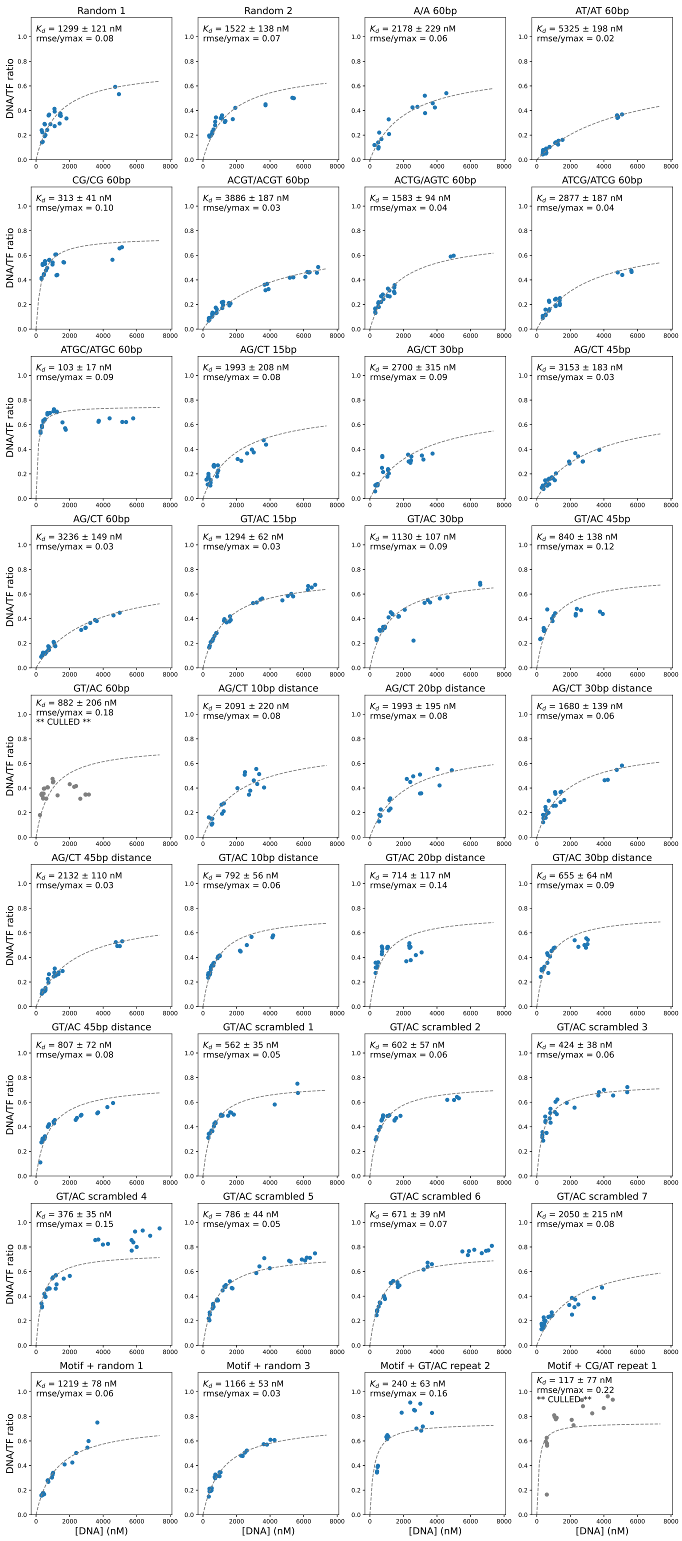

Figure 25: Example binding curves for MAX binding to DNA Library 2 from experiment on 10/10/2019. Markers indicate per-chamber fluorescence intensity ratios (Alexa-647/eGFP); dashed line indicates fit to a Langmuir isotherm; data that failed quality controls are shown in gray and marked as **CULLED**. Binding curves for other experiments are available on OSF (https://osf.io/gbxhz/).

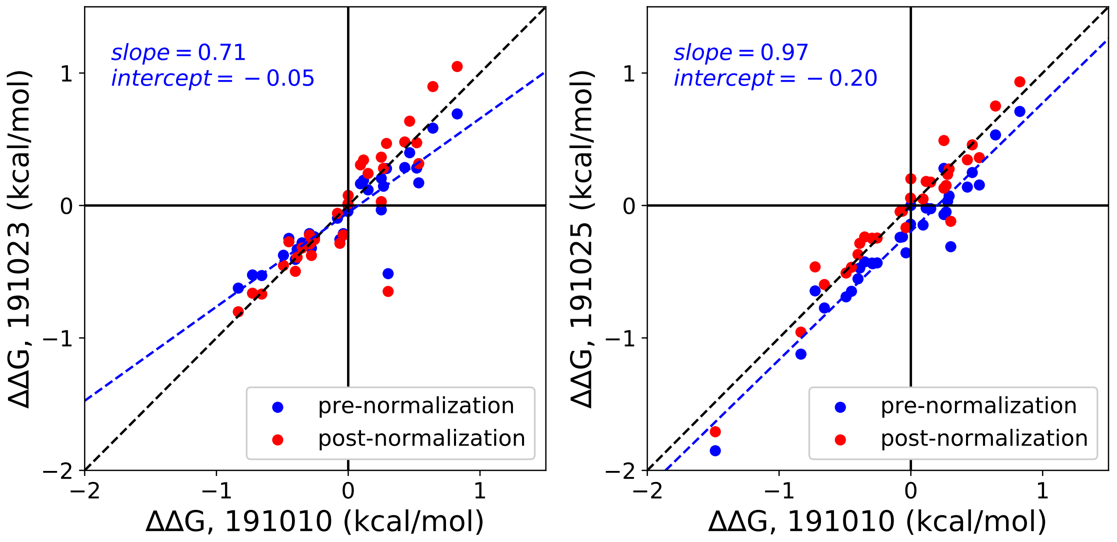

Figure 26: Comparisons between ΔΔG measurements for MAX binding to DNA Library 2 across 3 different experiments before (blue markers) and after (red markers) linear normalization (see Supplementary Methods). Blue line shows a linear regression to data prior to normalization; blue text indicates pre-normalization regression parameters; black dashed line indicates the 1:1 identity line.

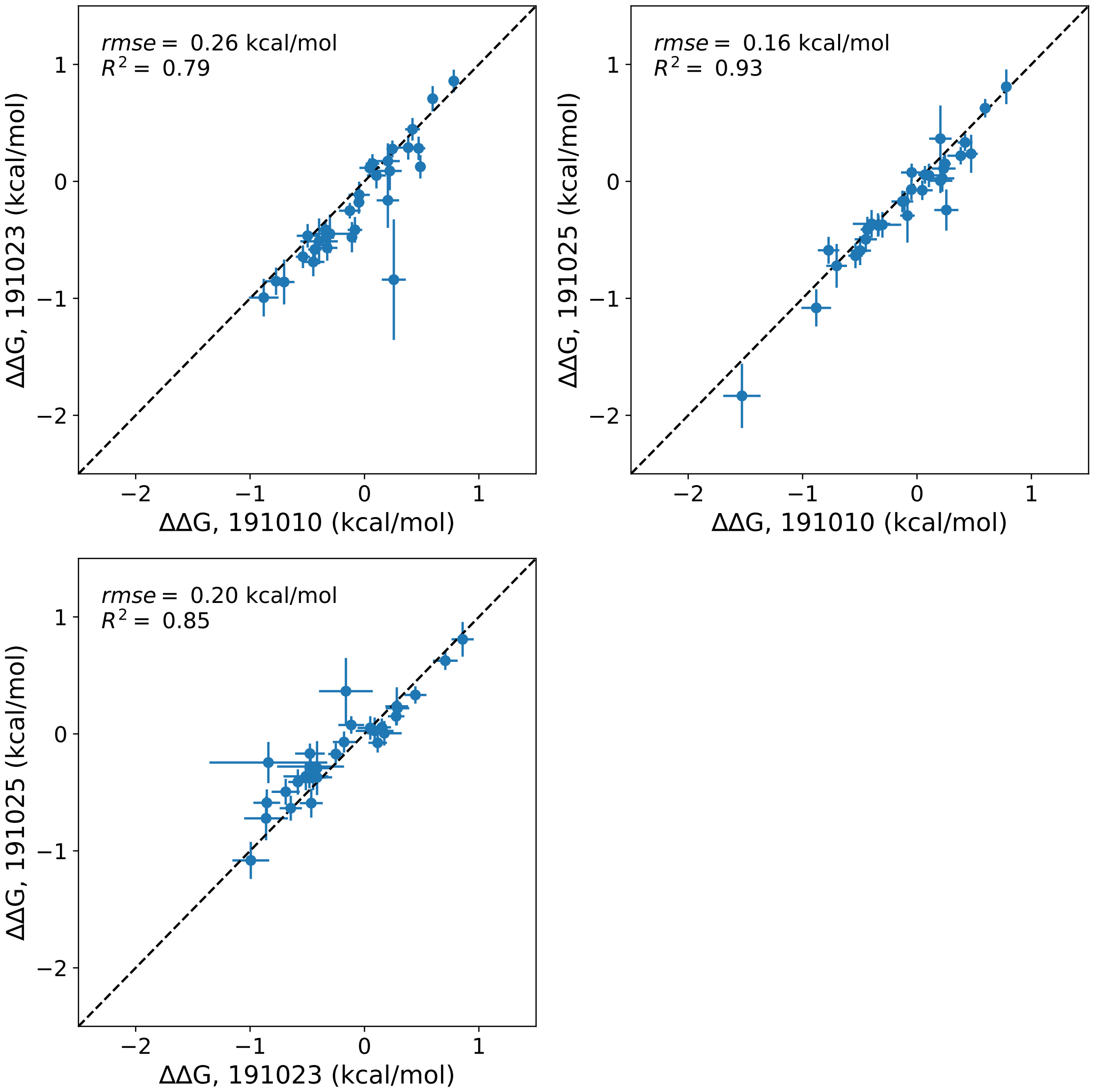

Figure 27: Pairwise comparisons of normalized ΔΔG values between all 3 experiments quantifying MAX binding to DNA Library 2.

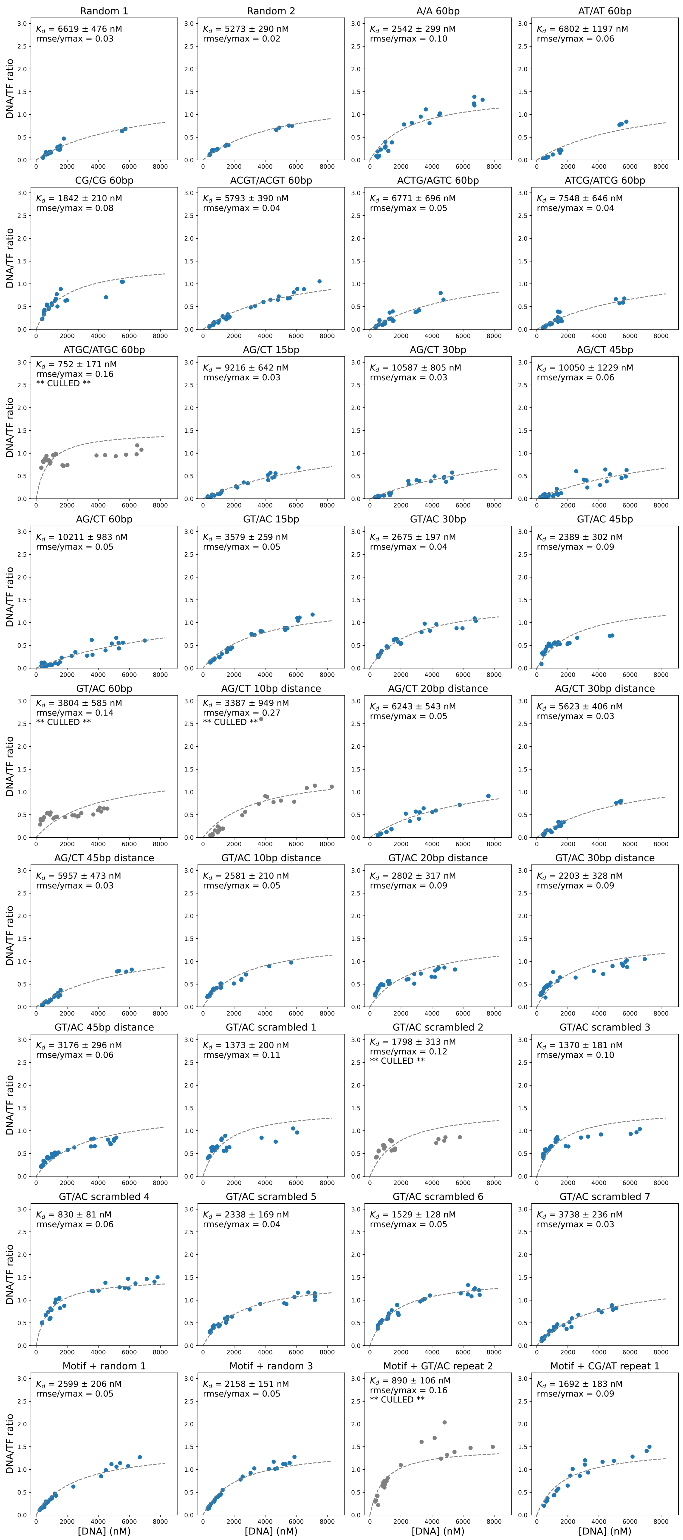

Figure 28: Example binding curves for Pho4 binding to DNA Library 2 from experiment on 11/04/2019. Markers indicate per-chamber fluorescence intensity ratios (Alexa-647/eGFP); dashed line indicates fit to a Langmuir isotherm; data that failed quality controls are shown in gray and marked as **CULLED**. Binding curves for other experiments are available on OSF (https://osf.io/gbxhz/).

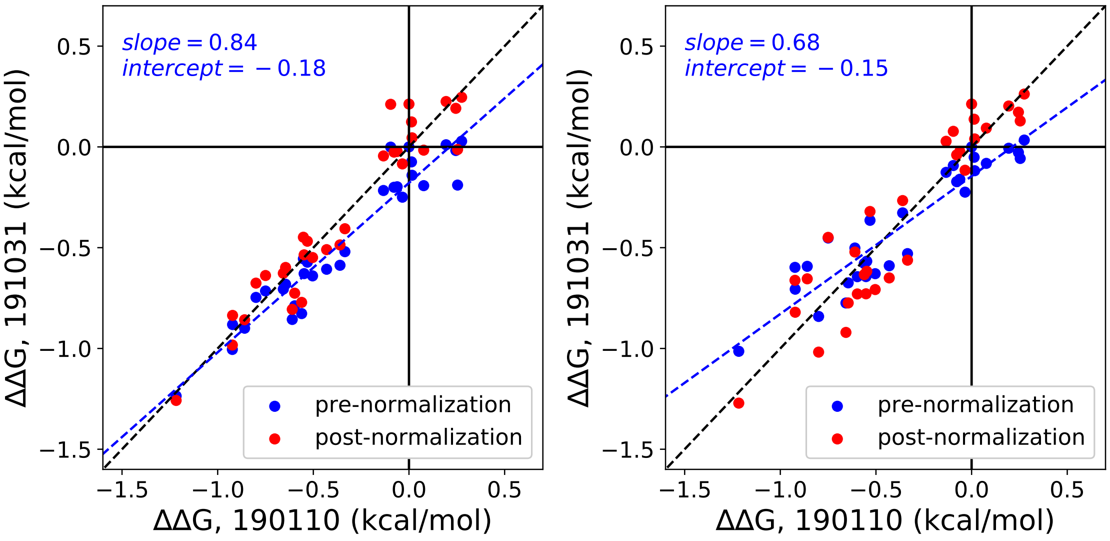

Figure 29: Comparisons between ΔΔG measurements for Pho4 binding to DNA Library 2 across 3 different experiments before (blue markers) and after (red markers) linear normalization (see Supplementary Methods). Blue line shows a linear regression to data prior to normalization; blue text indicates pre-normalization regression parameters; black dashed line indicates the 1:1 identity line.

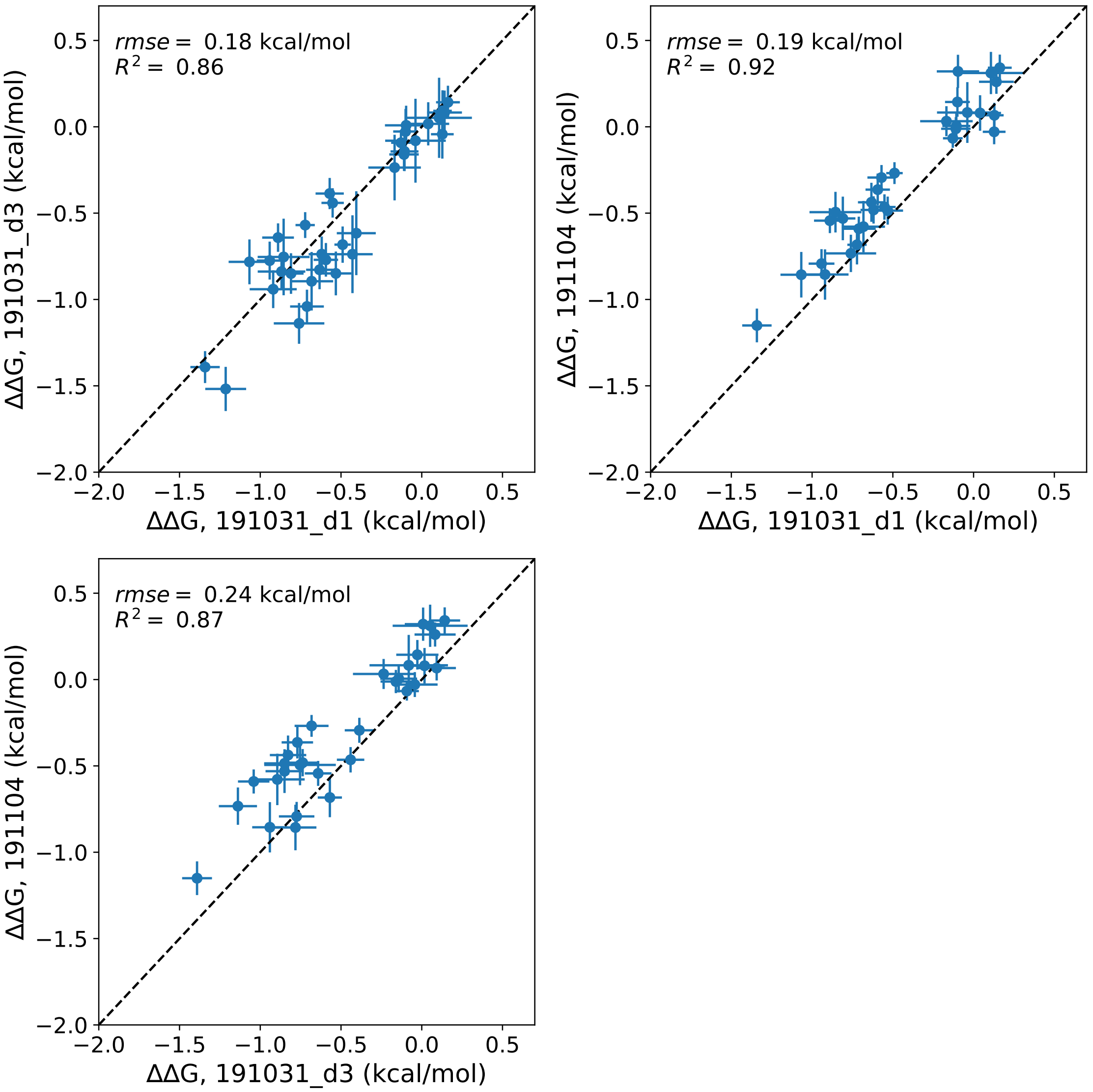

Figure 30: Pairwise comparisons of normalized ΔΔG values between all 3 experiments quantifying Pho4 binding to DNA Library 2.

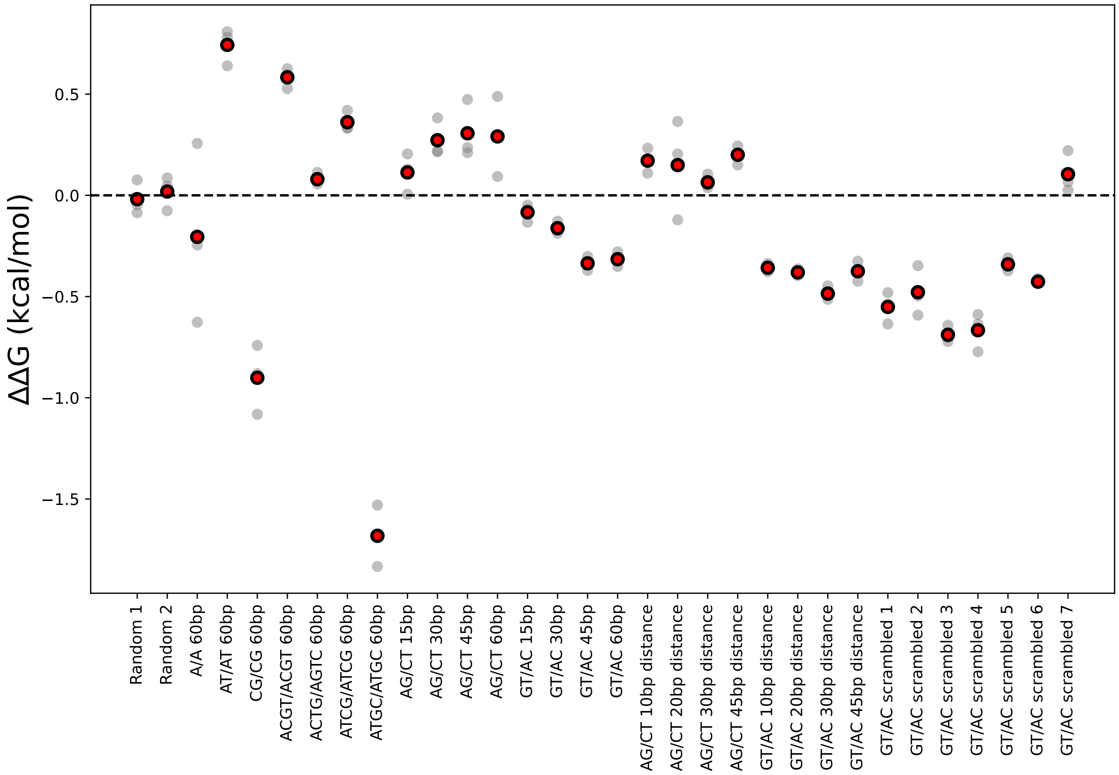

Figure 31: Normalized ΔΔGs across all experiments for MAX binding to each sequence from DNA Library 2. Gray dots indicate per-chamber ΔΔGs across all experiments; red dots indicate overall median. All ΔΔGs are calculated relative to the median affinity for all sequences containing a motif surrounded by random flanking sequences.

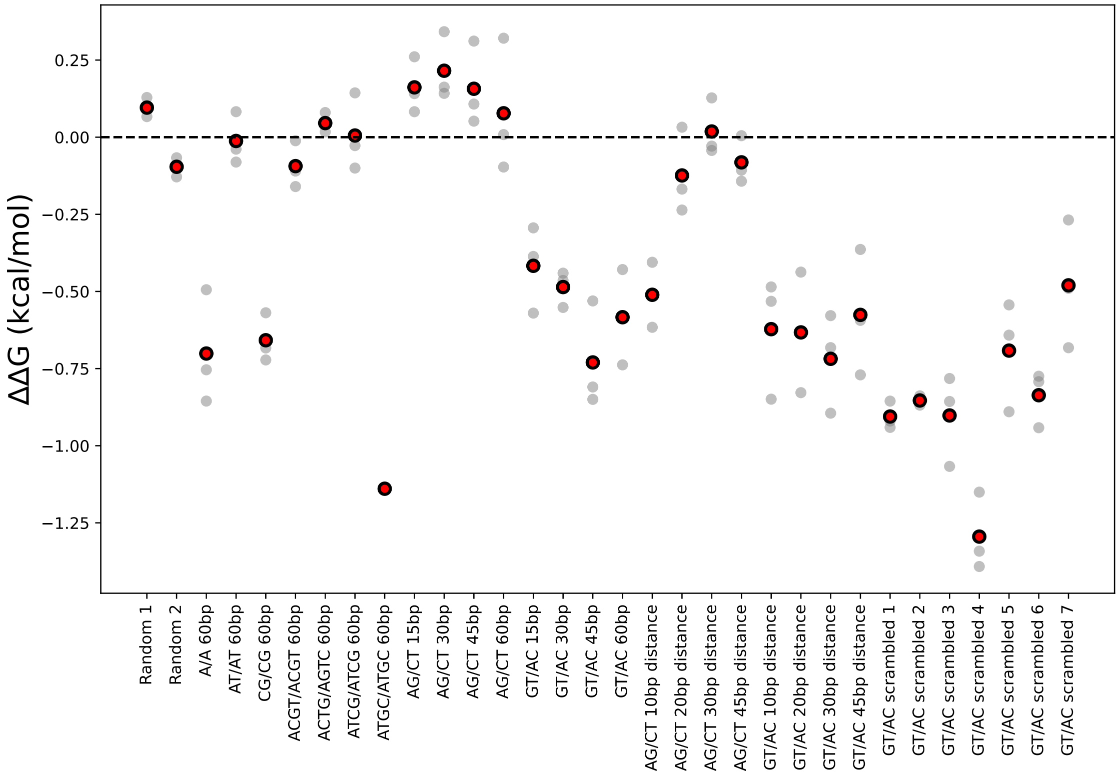

Figure 32: Normalized ΔΔGs across all experiments for Pho4 binding to each sequence from DNA Library 2. Gray dots indicate per-chamber ΔΔGs across all experiments; red dots indicate overall median. All ΔΔGs are calculated relative to the median affinity for all sequences containing a motif surrounded by random flanking sequences.

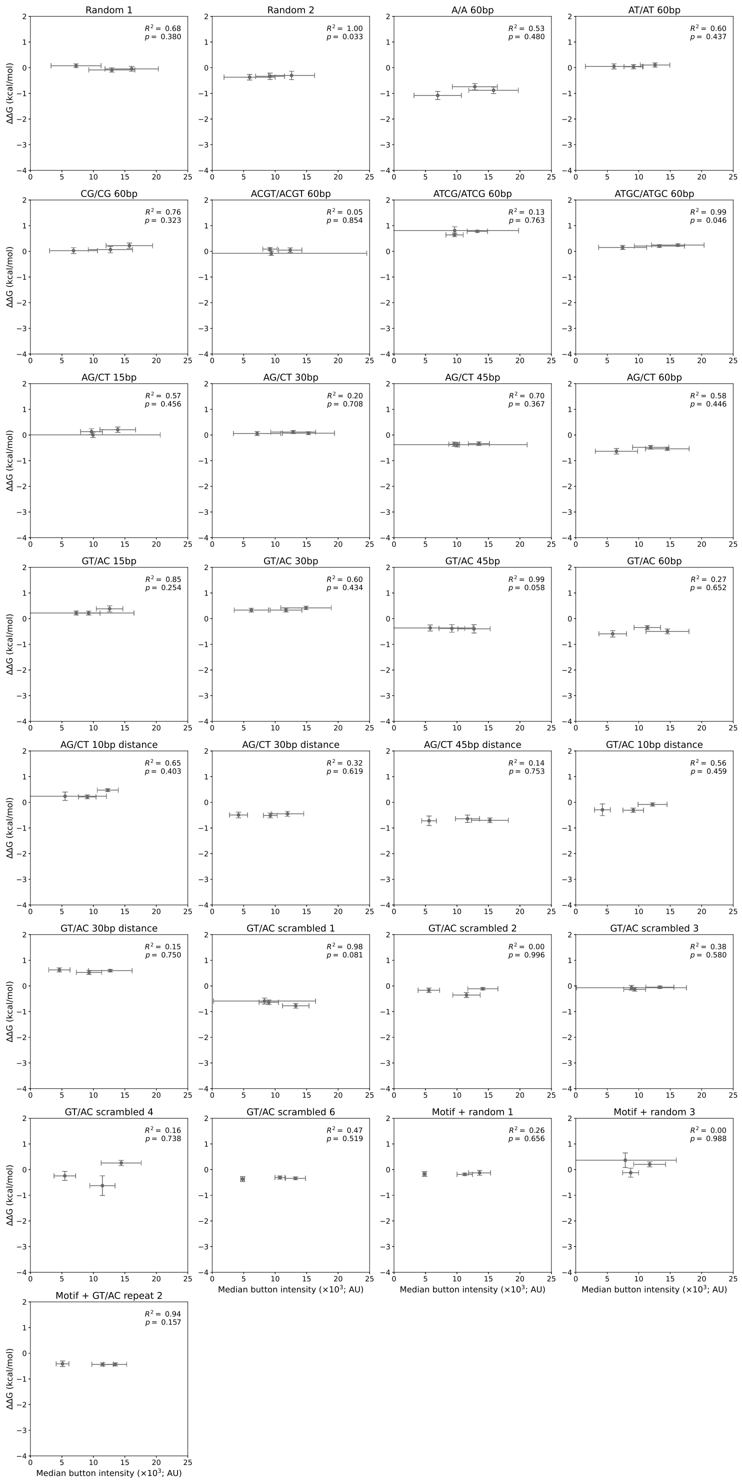

Figure 33: Measured DNA Library 2 ΔΔG values for MAX do not depend on surface-immobilized TF concentration. Each panel shows measured ΔΔG values *vs.* eGFP button intensities across all experiments for a given DNA Library 2 sequence. Markers indicate median for all replicates within a given experiment; error bars indicate standard deviation.

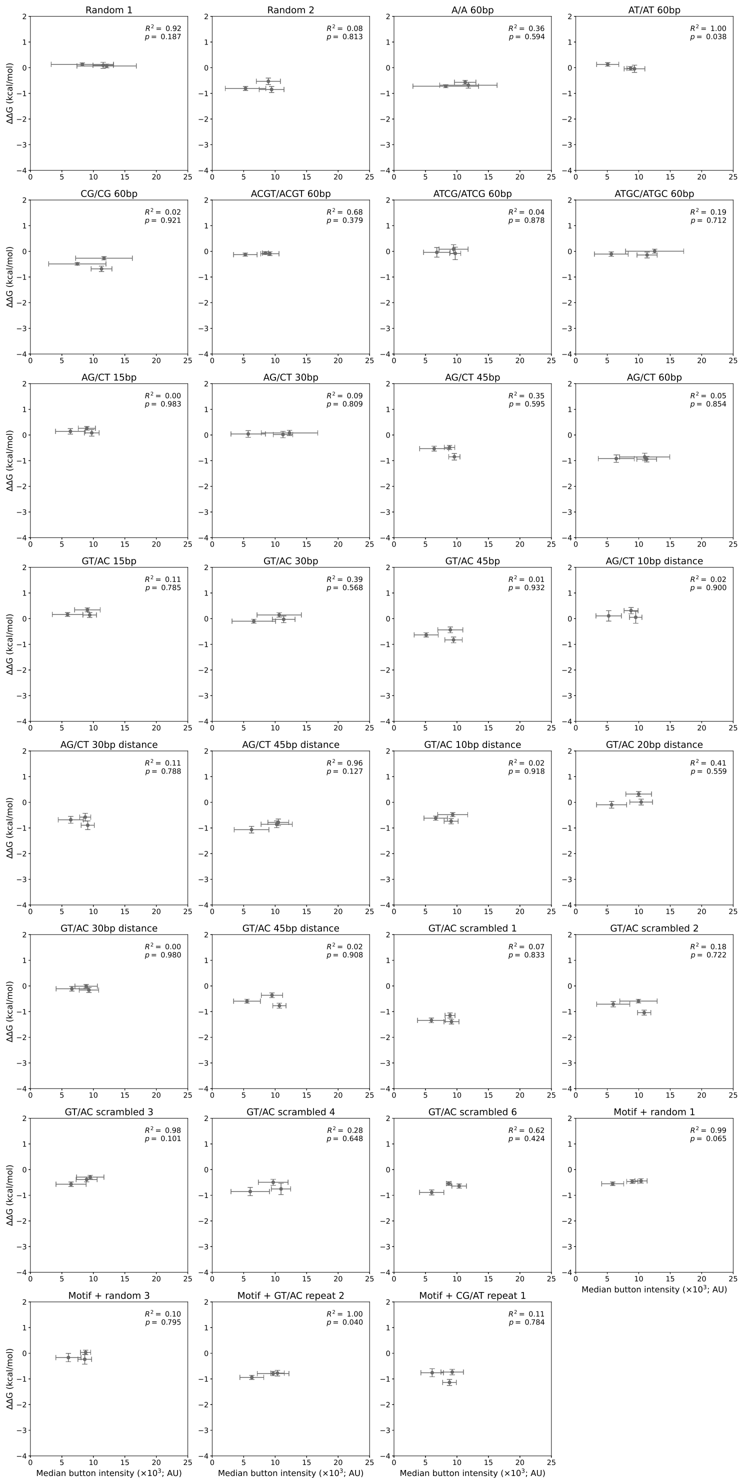

Figure 34: Measured DNA Library 2 ΔΔG values for Pho4 do not depend on surface-immobilized TF concentration. Each panel shows measured ΔΔG values *vs.* eGFP button intensities across all experiments for a given DNA Library 2 sequence. Markers indicate median for all replicates within a given experiment; error bars indicate standard deviation.

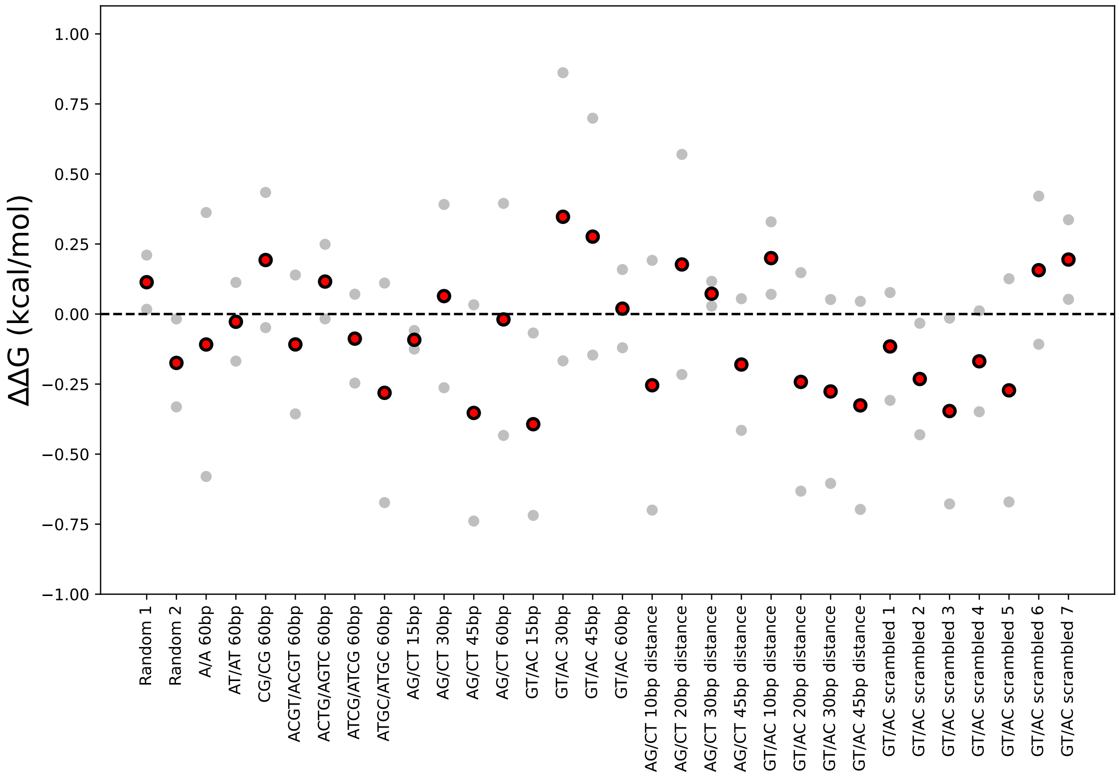

Figure 35: ΔΔGs across all experiments for an eGFP-only negative control binding to each sequence from DNA Library 2. Gray dots indicate per-chamber ΔΔGs across all experiments; red dots indicate overall median. All ΔΔGs are calculated relative to the median affinity for all sequences containing a motif surrounded by random flanking sequences.

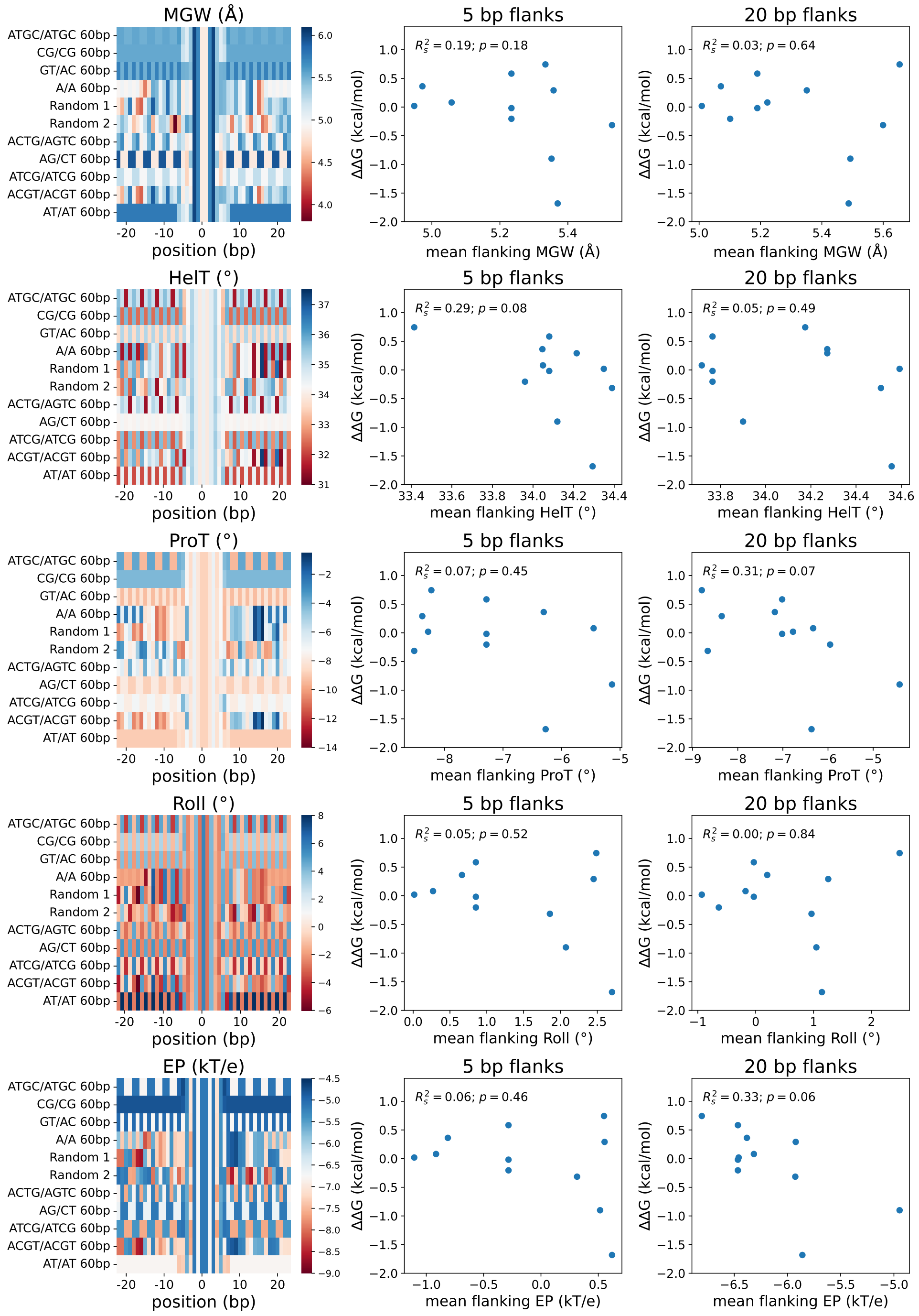

Figure 36: DNA shape parameters do not correlate with observed effects on DNA binding affinity for MAX. **(A)** Heat maps displaying calculated minor groove width (MGW), helical twist (HelT), propeller twist (ProT), roll (Roll), and electrostatic potential (EP) as a function of flanking nucleotide position for DNA Library 2 sequences. **(B)** Scatter plots showing measured ΔΔGs *vs.* calculated DNA shape parameters for each DNA Library 2 sequence. Left and right columns show calculations considering either the proximal 5 or 20 nucleotides on either side of the consensus motif, respectively. Sequences in heatmaps are sorted from top to bottom in order of highest to lowest affinity.

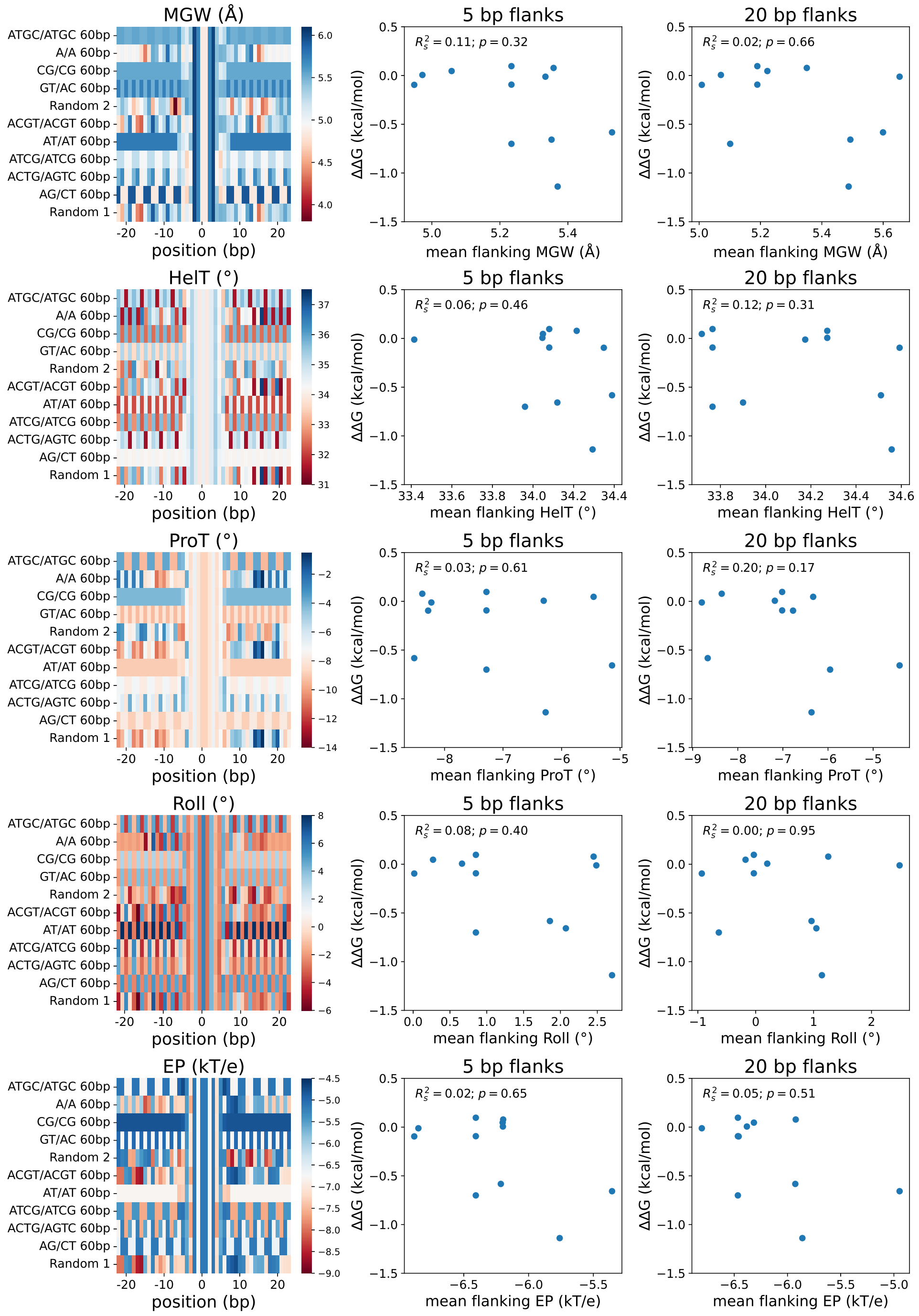

Figure 37: DNA shape parameters do not correlate with observed effects on DNA binding affinity for Pho4. **(A)** Heat maps displaying calculated minor groove width (MGW), helical twist (HelT), propeller twist (ProT), roll (Roll), and electrostatic potential (EP) as a function of flanking nucleotide position for DNA Library 2 sequences. **(B)** Scatter plots showing measured ΔΔGs *vs.* calculated DNA shape parameters for each DNA Library 2 sequence. Left and right columns show calculations considering either the proximal 5 or 20 nucleotides on either side of the consensus motif, respectively. Sequences in heatmaps are sorted from top to bottom in order of highest to lowest affinity.

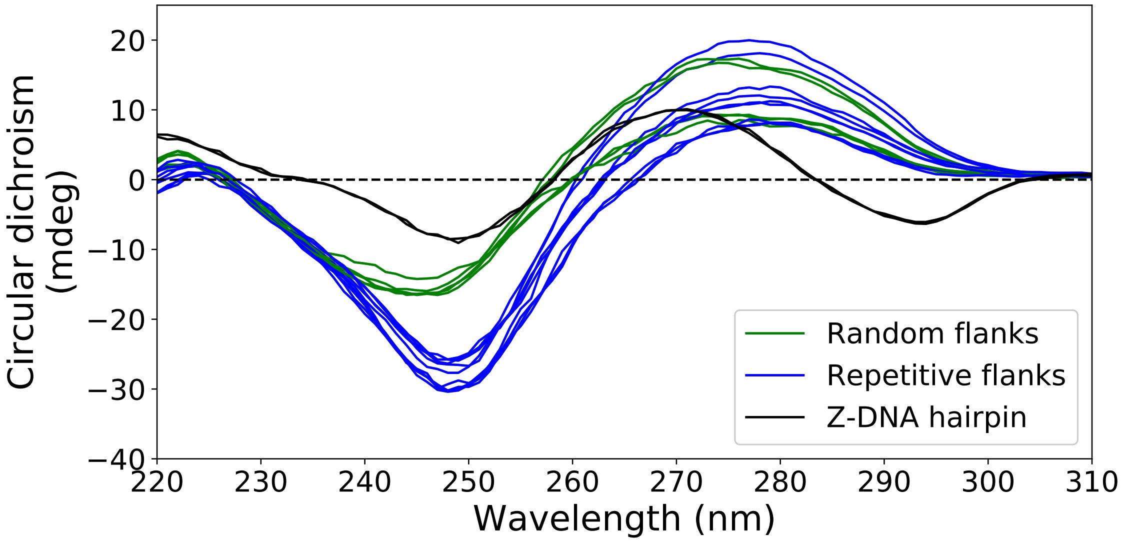

Figure 38: CD spectra for double-stranded DNA sequences containing a central E-box motif surrounded by either random (green lines) or repetitive (blue lines) flanking sequences; representative spectra for a GC hairpin sequence predicted to form Z-DNA are shown in black for comparison.

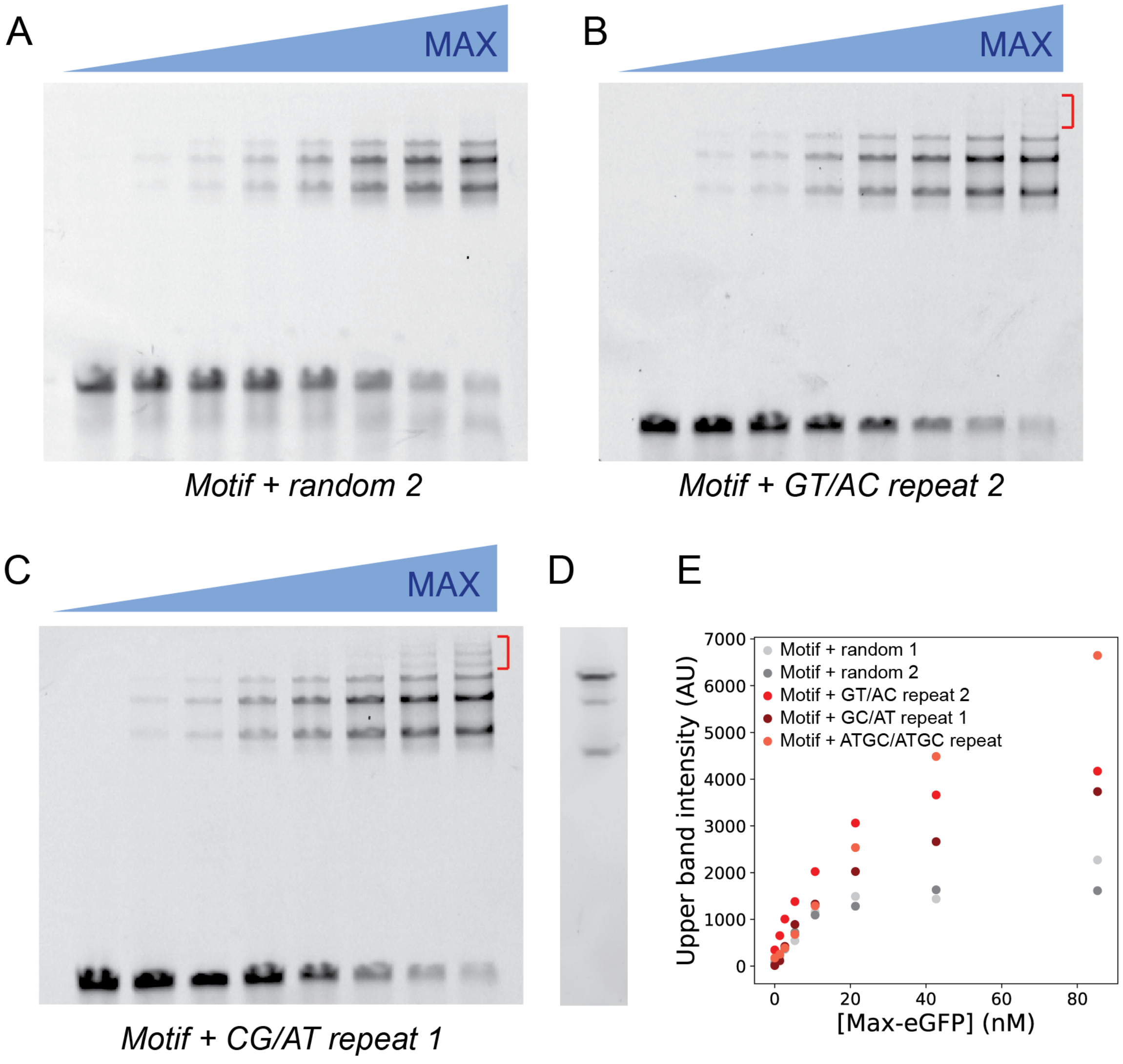

Figure 39: Additional EMSAs and quantification of gel band intensities. **(A)** EMSA for sequence containing a motif surrounded by random sequence, imaged in the Cy5 (DNA) channel. Concentration of MAX increases left to right. **(B)** EMSA for sequence containing a motif flanked on one side by GT repeats and on the other by AC repeats, imaged in the Cy5 (DNA) channel. GT/AC repeat flanks are favored over random flanks. Concentration of MAX increases left to right. Red bracket indicates DNA species bound by multiple TFs. **(C)** EMSA for sequence containing flanked on one side by CG repeats and on the other by AT repeats, imaged in the Cy5 (DNA) channel. CG/AT repeat flanks are favored over random flanks. Concentration of MAX increases left to right. Red bracket indicates DNA species bound by multiple TFs. **(D)** MAX-eGFP on a native protein gel runs with 3 bands, likely corresponding to MAX-eGFP dimers, MAX-eGFP monomers, and eGFP alone. Imaged in the eGFP channel. **(E)** Quantification of Alexa-647 signal in gel bands corresponding to MAX-bound DNA. Motif + repeat 1 and Motif + ATGC/ATGC repeat quantification correspond to the EMSAs shown in the main text, **Fig. 2D.**

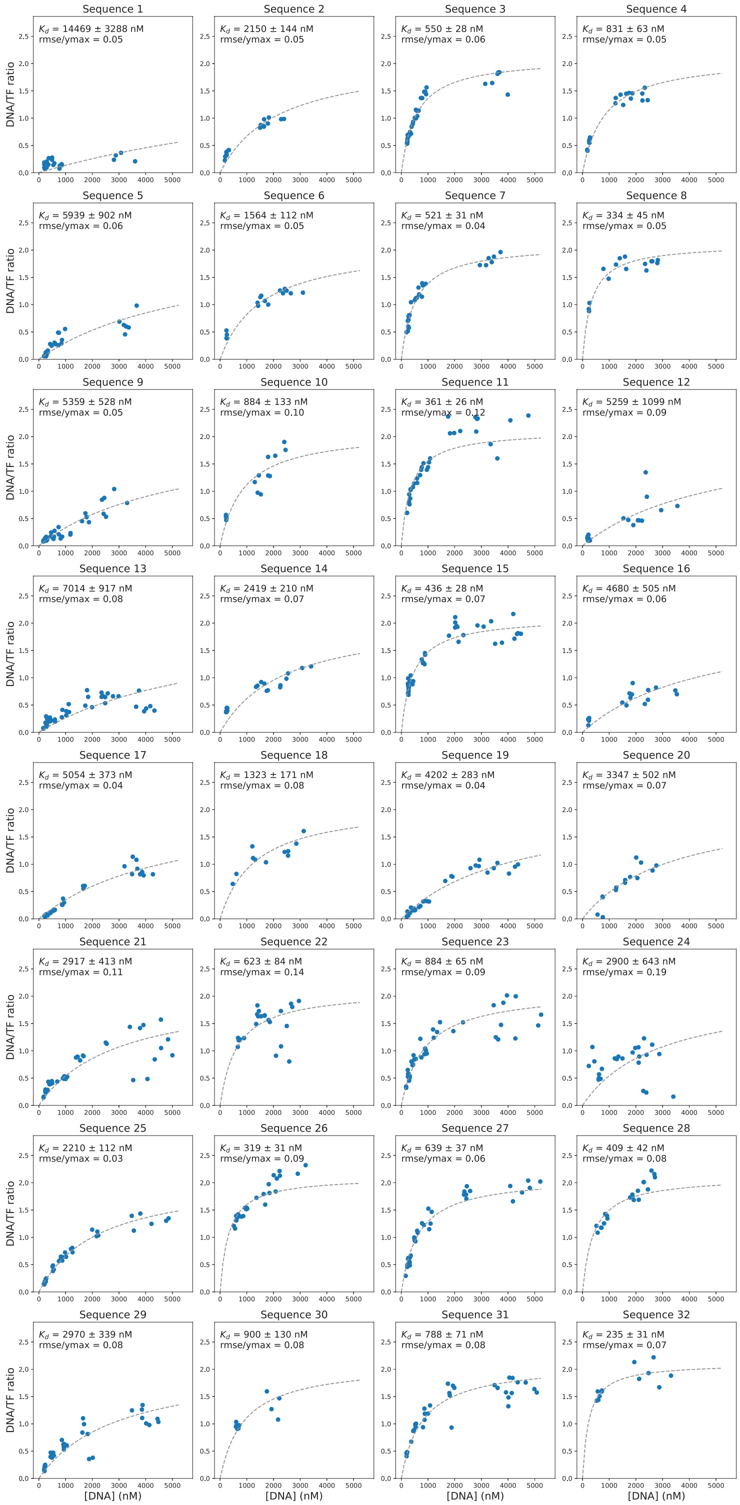

Figure 40: Binding curves for MAX binding to a selection of fluorescently-labeled dsDNA gcPBM probe sequences from Afek *et al.* [3]. Data from experiment on 03/25/2019. Markers indicate per-chamber fluorescence intensity ratios (Alexa-647/eGFP); dashed line indicates fit to a Langmuir isotherm.

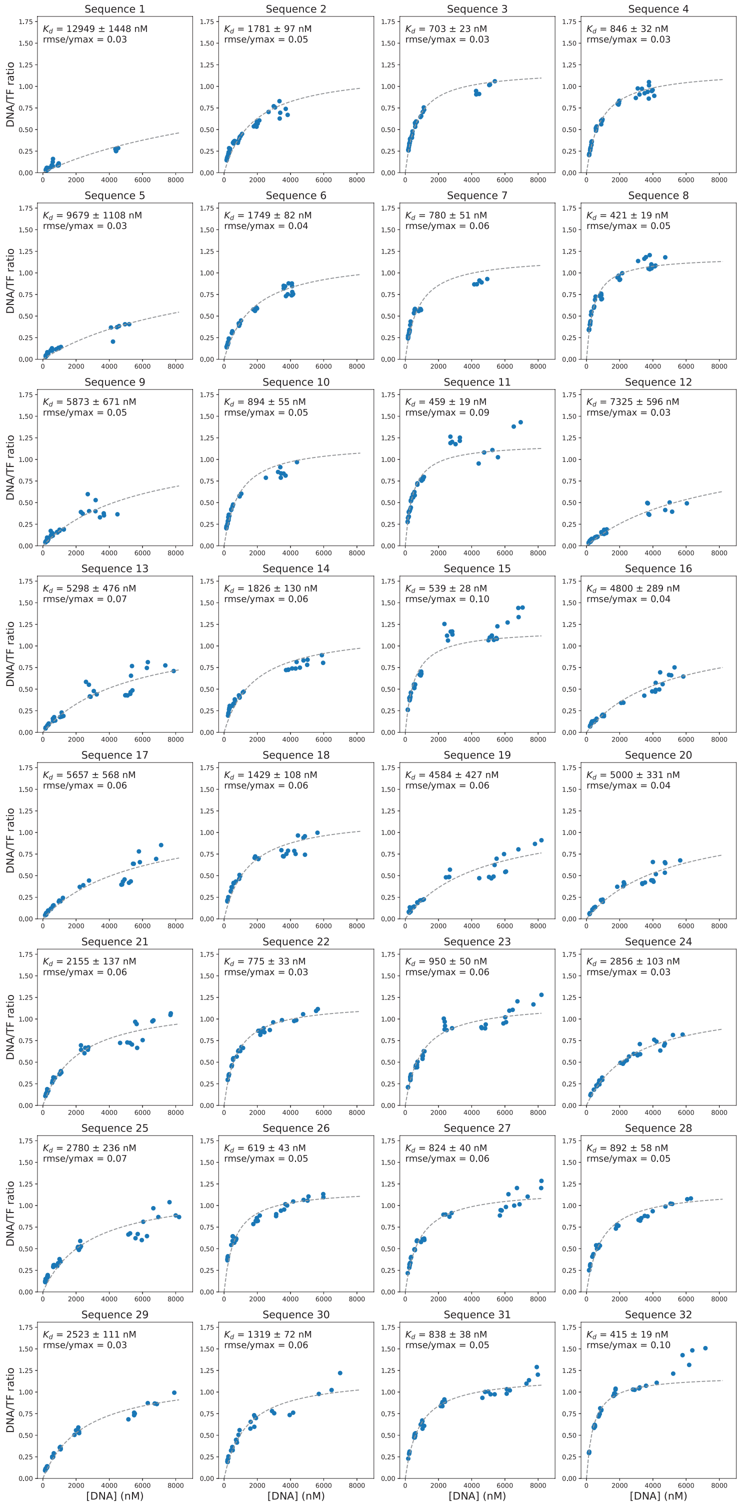

Figure 41: Binding curves for MAX binding to a selection of fluorescently-labeled dsDNA gcPBM probe sequences from Afek *et al.* [3]. Data from experiment on 04/11/2019. Markers indicate per-chamber fluorescence intensity ratios (Alexa-647/eGFP); dashed line indicates fit to a Langmuir isotherm. Binding curves for other experiments are available on OSF (https://osf.io/gbxhz/).

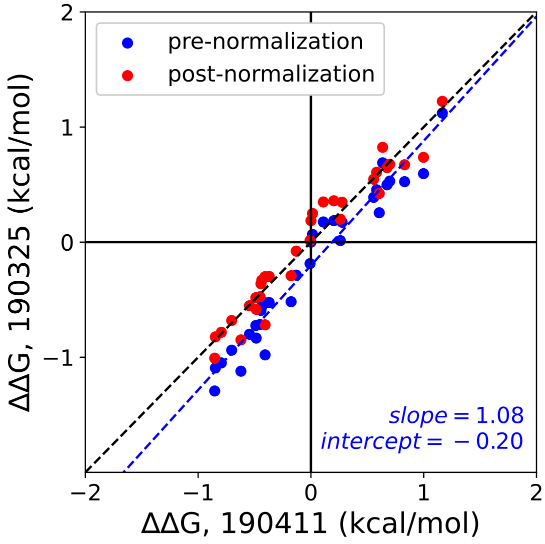

Figure 42: Comparisons between ΔΔG measurements for MAX binding to 32 gcPBM dsDNA sequences across 2 experimental replicates before (blue markers) and after (red markers) normalization (see Supplementary Methods). Blue line shows a linear regression to data prior to normalization; blue text indicates pre-normalization regression parameters; black dashed line indicates the 1:1 identity line.

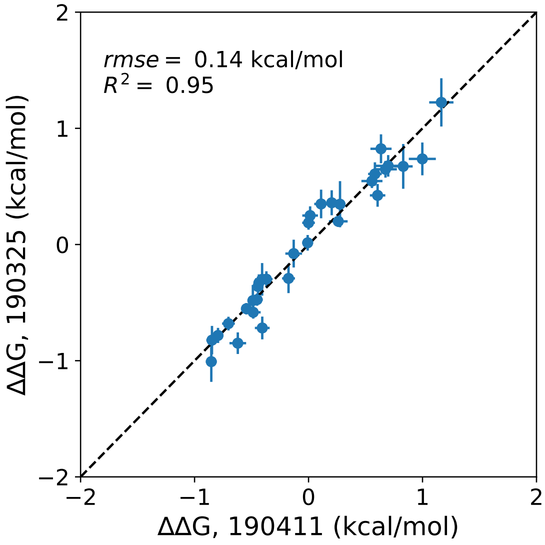

Figure 43: Pairwise comparisons of normalized ΔΔG values between experiments quantifying MAX binding to 32 gcPBM dsDNA sequences. Markers indicate median across all replicates within a given experiment; error bars indicate standard deviation of measured values.

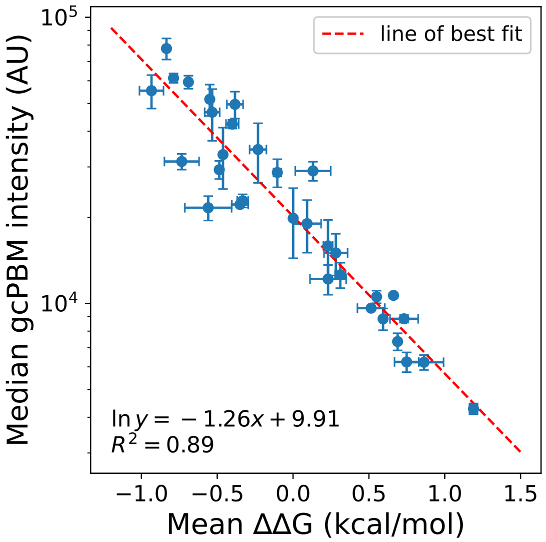

Figure 44: Comparison between MAX intensities in a gcPBM experiment *vs.* ΔΔGs measured via MITOMI. Markers indicate median intensities and ΔΔG values within a given experiment; error bars indicate standard deviation. Red line indicates a linear regression; returned fit parameters can be used to convert intensities to ΔΔGs for all remaining gcPBM probes.

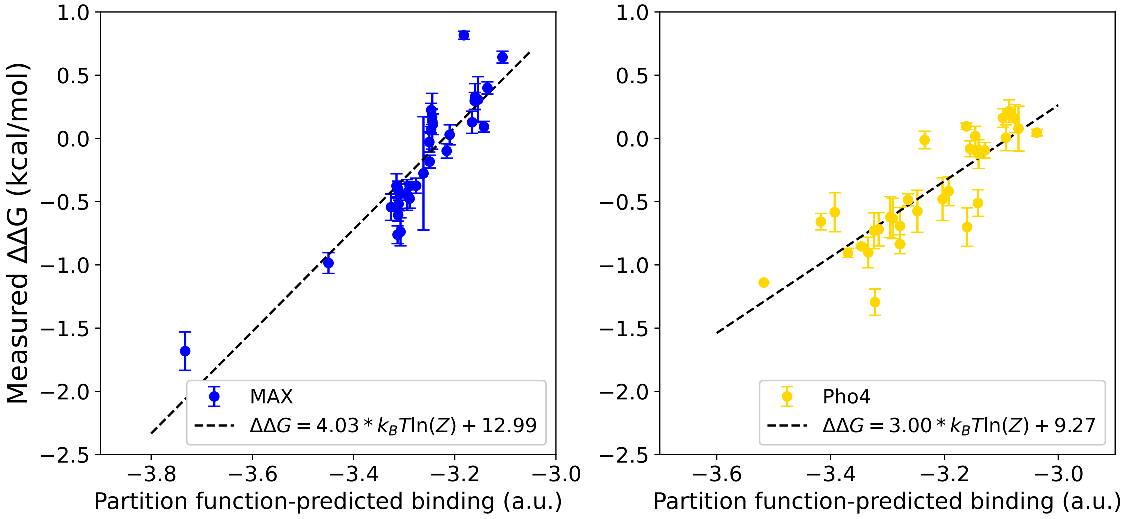

Figure 45: Measured binding of MAX (left) and Pho4 (right) binding to DNA Library 2 *vs.* uncalibrated partition function-based predictions of binding. Data are shown as mean ± standard deviation across experimental replicates. Dashed line indicates best fit, with slope and intercept of linear regression used for subsequent calibrates indicated.

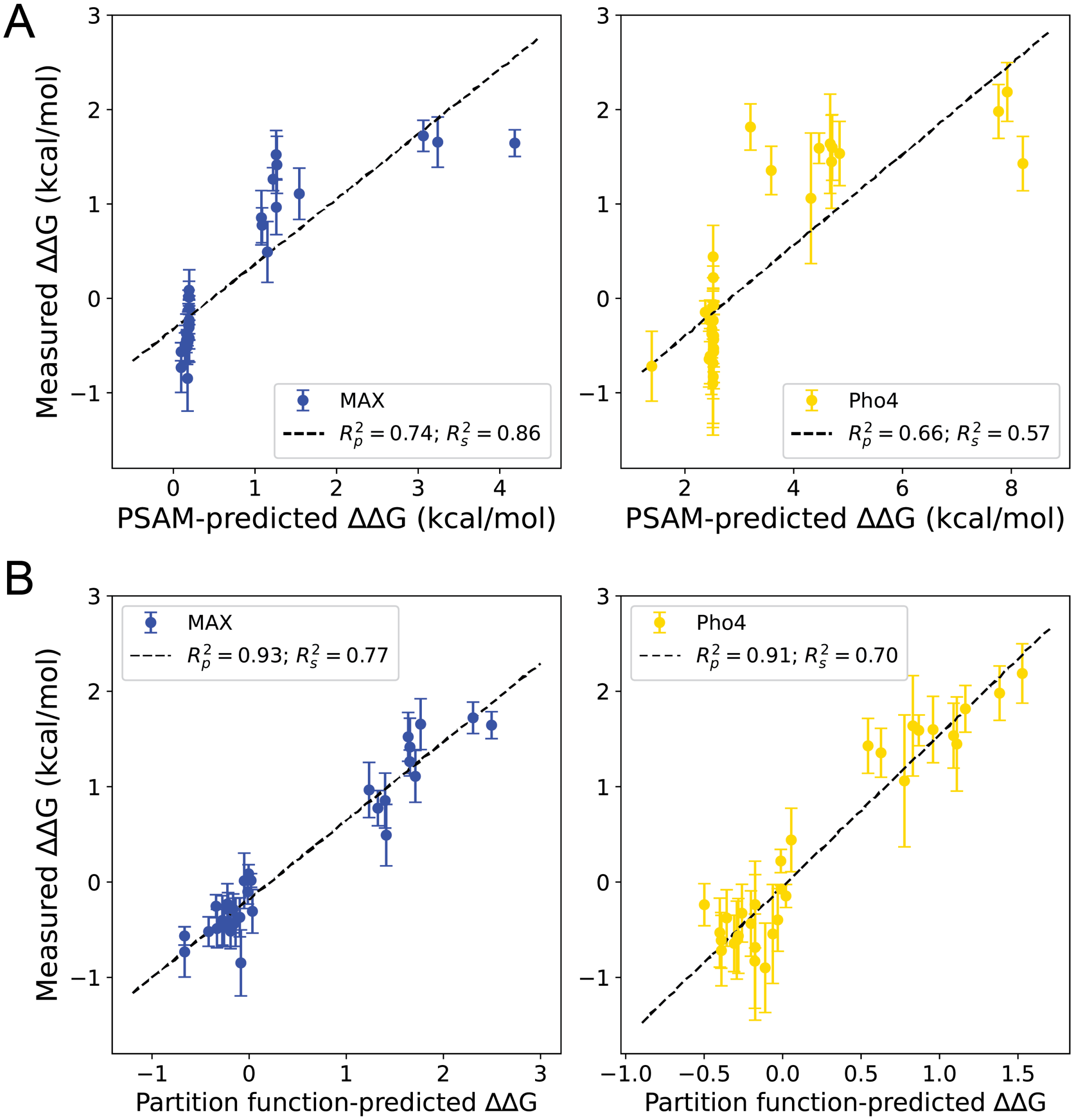

Figure 46: PSAM and partition-function based predictions of binding to DNA Library 1. **(A)** Measured binding of MAX (left) and Pho4 (right) binding to DNA Library 1 *vs.* PSAM-derived binding predictions. Data are shown as mean ± standard deviation across experimental replicates. Dotted line indicates best fit. **(B)** Measured binding of MAX (left) and Pho4 (right) binding to DNA Library 1 *vs.* partition function-derived predictions. Partition function-derived predictions were calibrated using linear regressions in Fig. S45. Data are shown as mean ± standard deviation across experimental replicates. Dotted line indicates best fit.

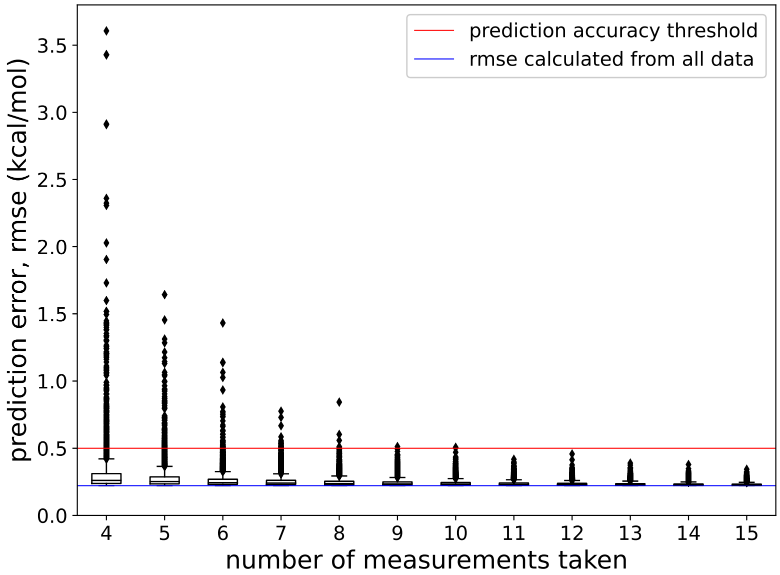

Figure 47: Results of simulation to estimate how many measurements are required for accurate calibration of partition function-derived binding energy predictions. For each value indicated on the $x$-axis, $n$ measurements were randomly sampled and used to calibrate predictions over 10,000 iterations. The rmse of predictions *vs.* measured binding is shown in a boxplot or outlier point for each of 10,000 iterations per $n$ measurements sampled. The rmse calculated from calibrating predictions with all 32 measurements is indicated in blue. An accuracy threshold of 0.5 kcal/mol is indicated in red.

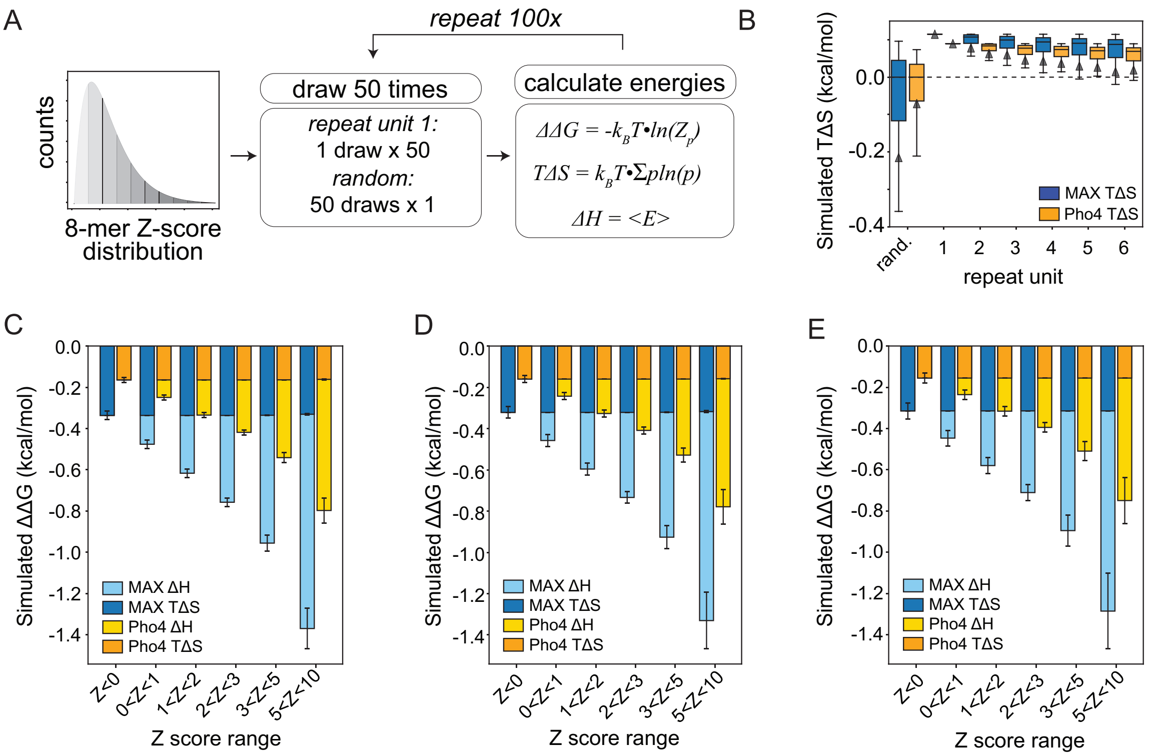

Figure 48: **(A)** Pipeline for simulating effects of repetitive *vs.* random flanking sequences on Gibbs free energy ($\Delta\Delta G$), entropy ($T\Delta S$), and enthalpy ($\Delta H$) as described in Sections 1.13–1.14. **(B)** Simulated entropy distributions as a function of repeat unit; box plots denote distribution median while triangles denote distribution mean. **(C)** Simulated entropy and enthalpy distributions for homopolymer repeat sequences (repeat unit = 1) for sequences with different Z-score ranges. **(D)** Simulated entropy and enthalpy distributions for dinucleotide repeat sequences (repeat unit = 2) for sequences with different Z-score ranges. **(E)** Simulated entropy and enthalpy distributions for tetranucleotide repeat sequences (repeat unit = 4) for sequences with different Z-score ranges.

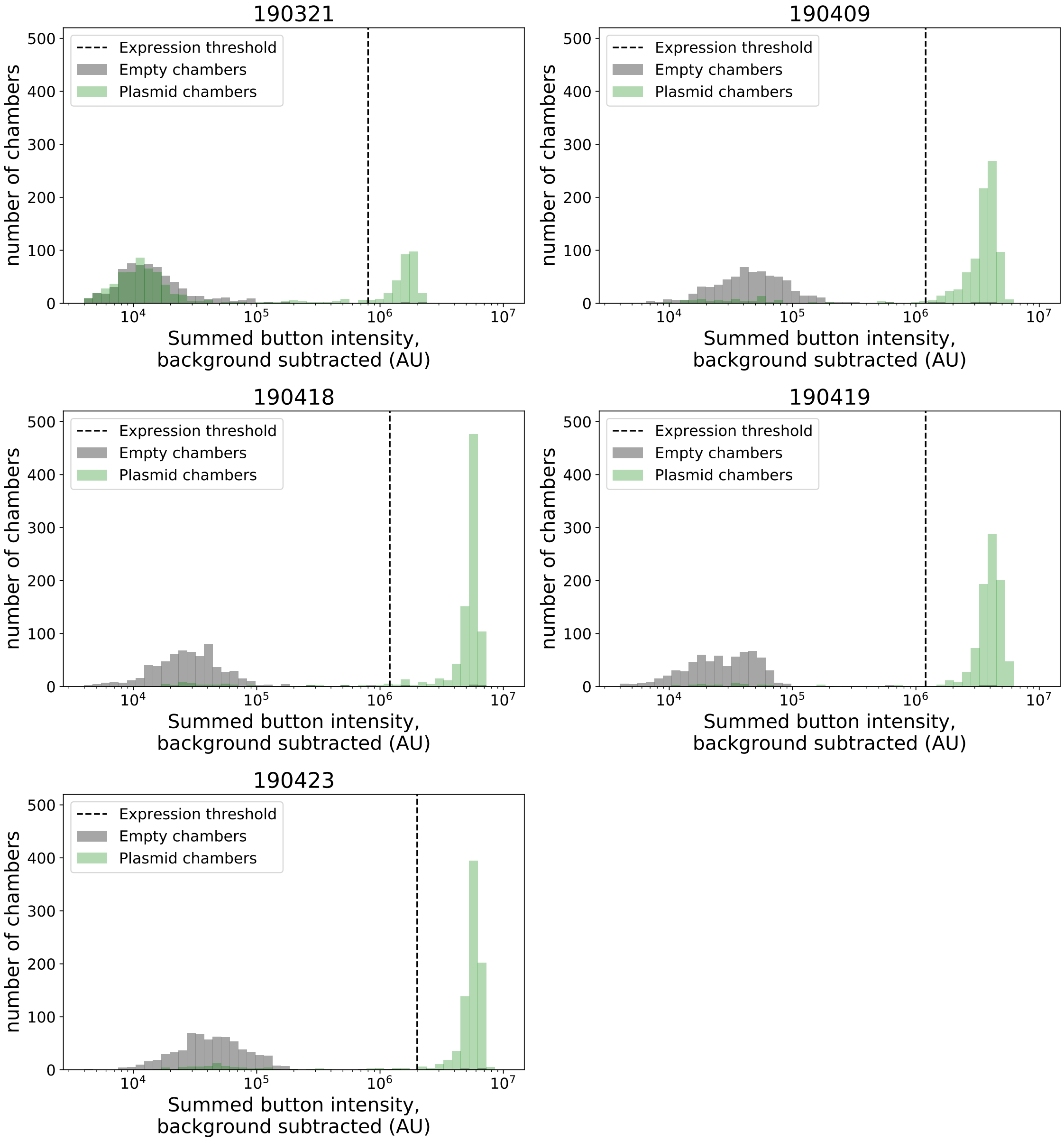

Figure 49: Distribution of measured per-chamber eGFP ‘button’ intensities for plasmid-containing (green bars) and empty (gray bars) chambers after on-chip *in vitro* transcription/translation in 5 STAMMP experiments quantifying binding of eGFP-tagged Pho4 to dsDNA containing an E-box motif surrounded by random sequence. Black dashed line indicates intensity threshold used to identify chambers with expressed Pho4 protein for downstream analysis.

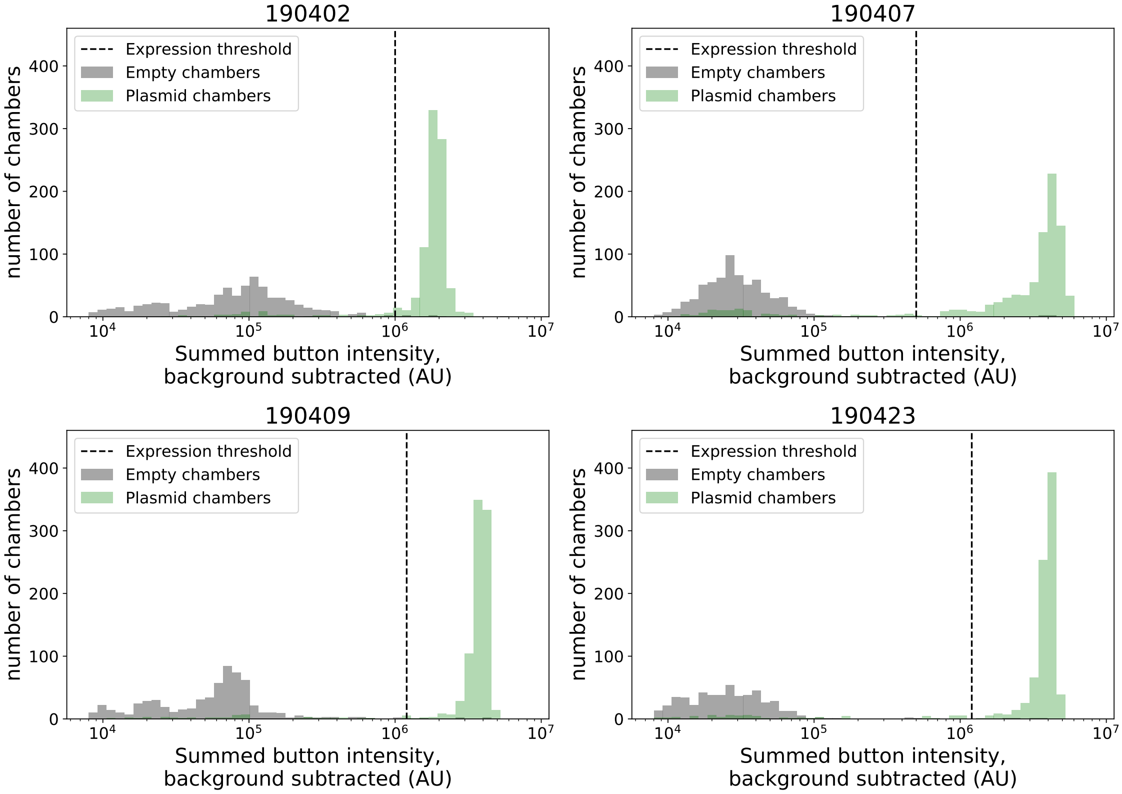

Figure 50: Distribution of measured per-chamber eGFP 'button' intensities for plasmid-containing (green bars) and empty (gray bars) chambers after on-chip *in vitro* transcription/translation in 4 STAMMP experiments quantifying binding of eGFP-tagged Pho4 to dsDNA containing an E-box motif surrounded by CG/AT repeats. Black dashed line indicates intensity threshold used to identify chambers with expressed Pho4 protein for downstream analysis.

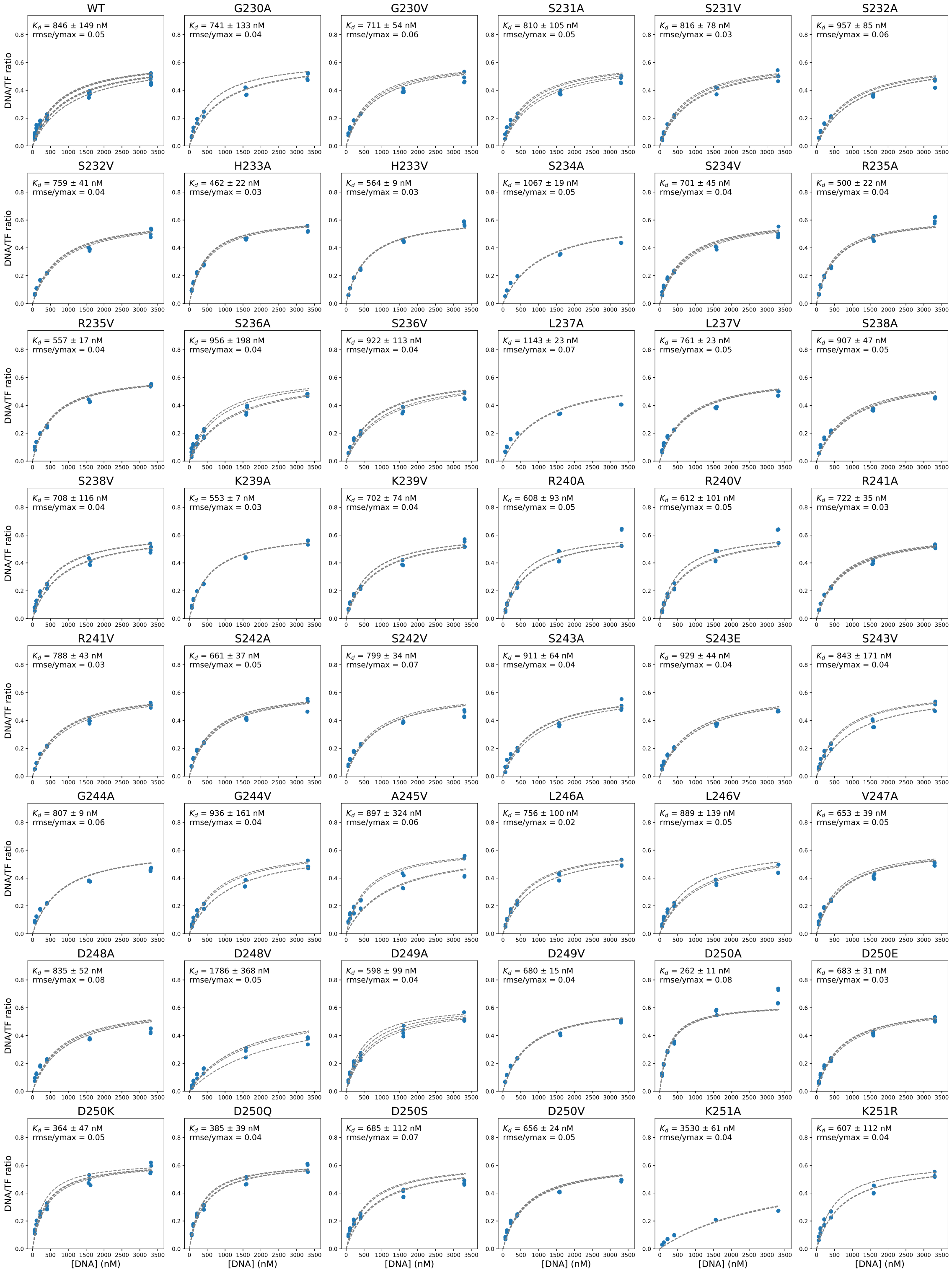

Figure 51: Example binding curves for STAMMP experiments assessing binding of 214 Pho4 variants passing QC to a DNA sequence containing a CACGTG E-box surrounded by random sequence from experiment on 04/23/2019. Markers indicate per-chamber fluorescence intensity ratios (Alexa-647/eGFP); dashed lines indicates per-chamber fits to a Langmuir isotherm. Binding curves for other experiments are available on OSF (https://osf.io/gbxhz/).

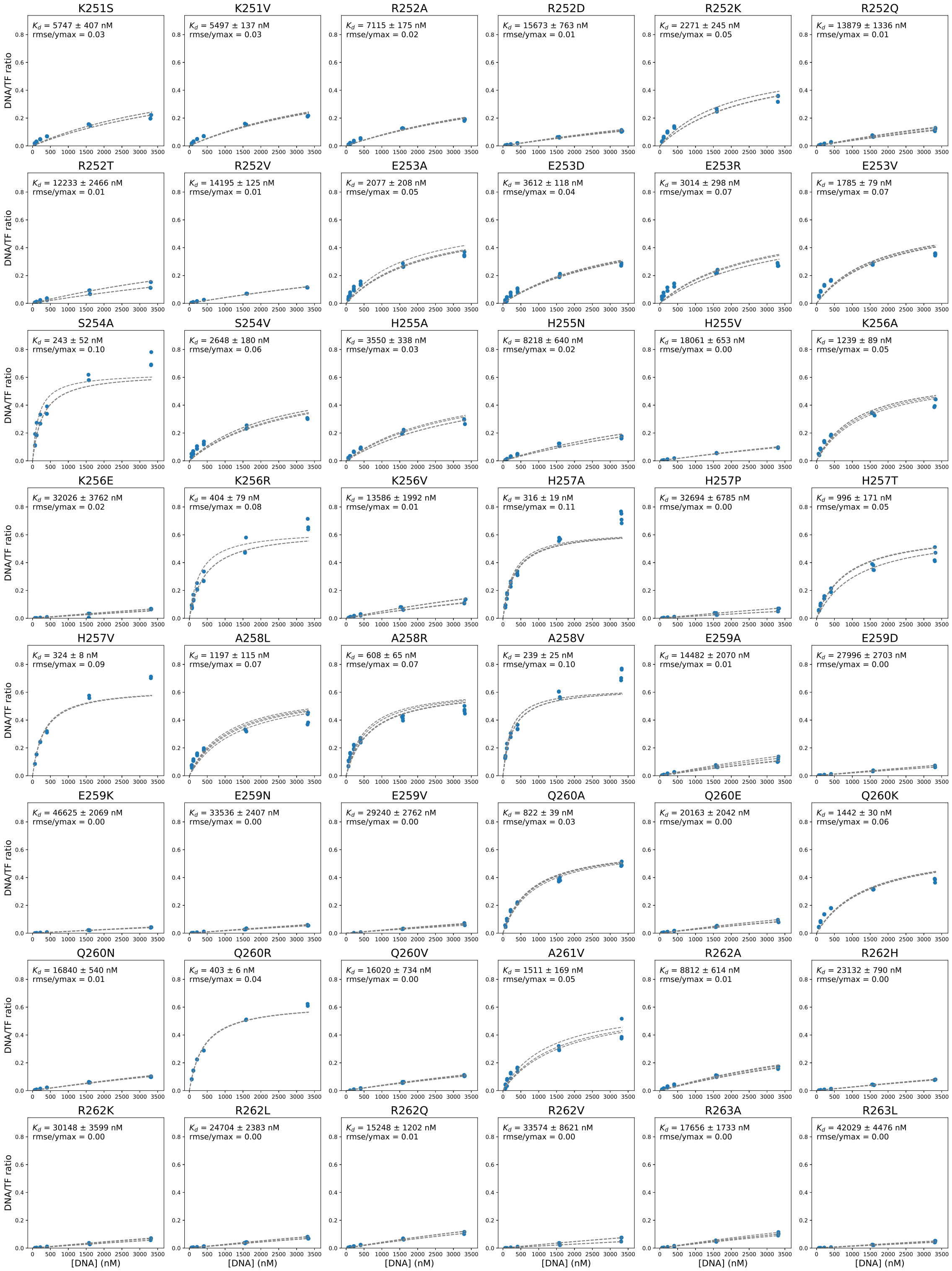

Figure 51: STAMMP binding curves (continued)

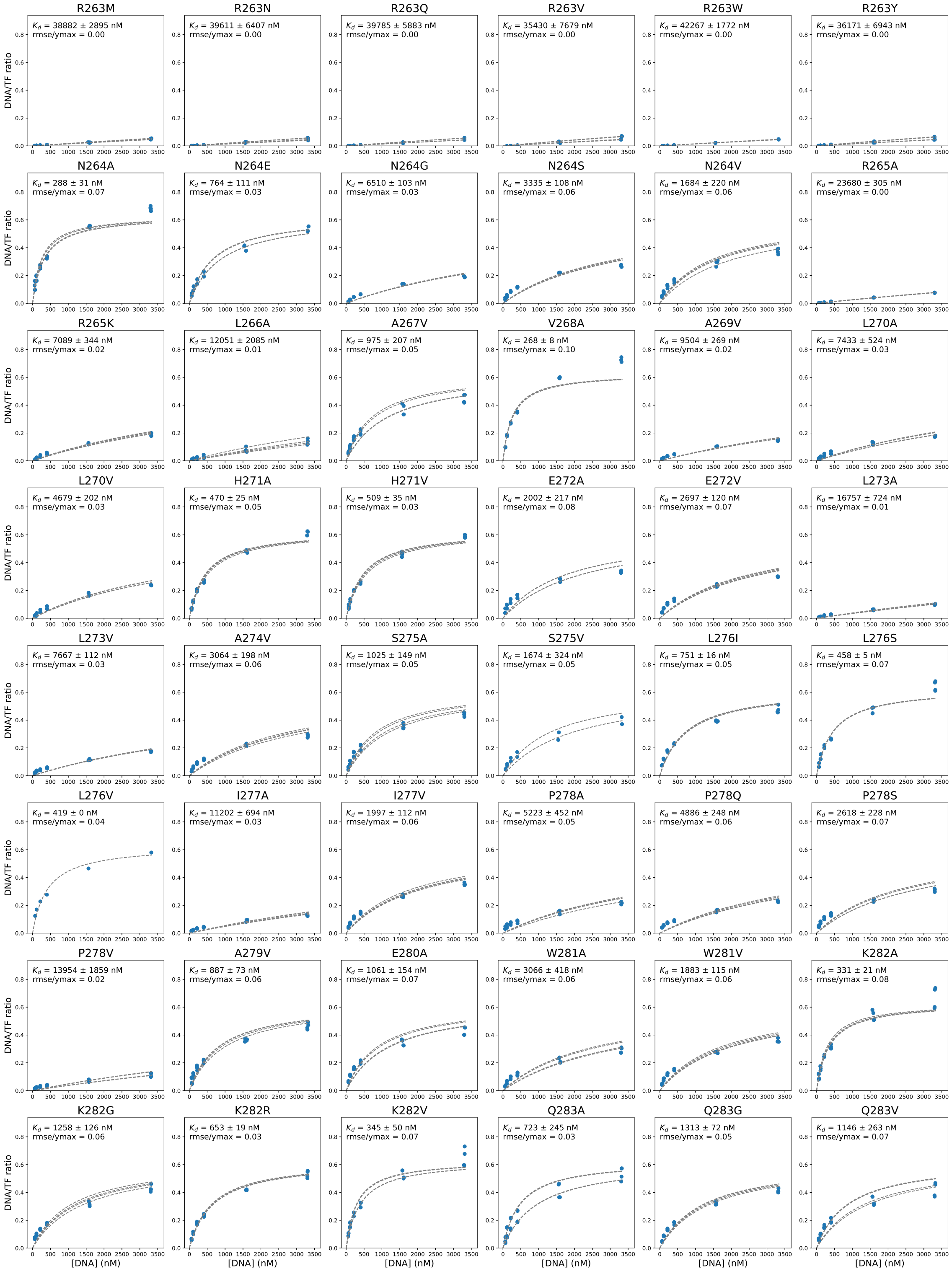

Figure 51: STAMMP binding curves (continued)

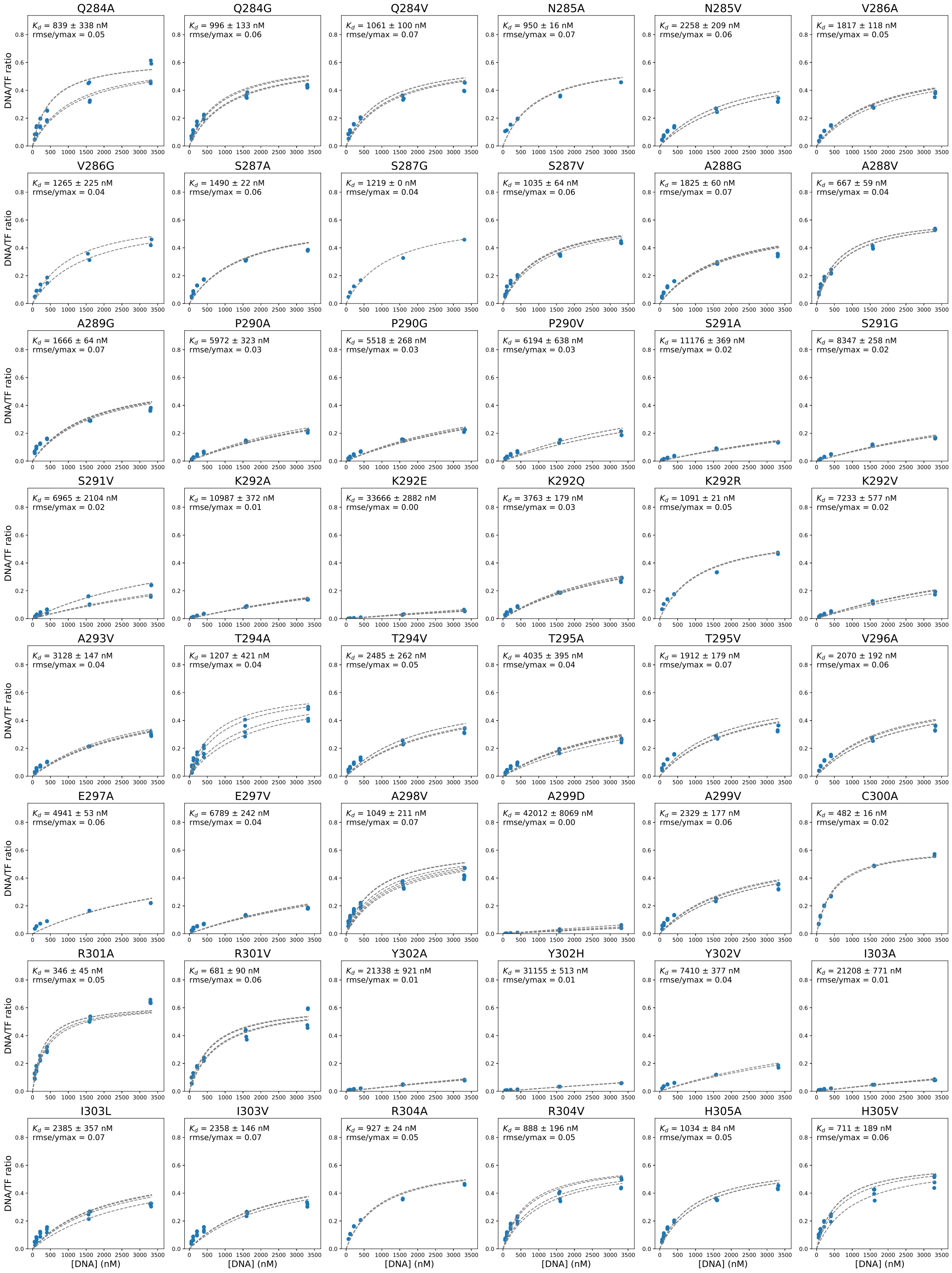

Figure 51: STAMMP binding curves (continued)

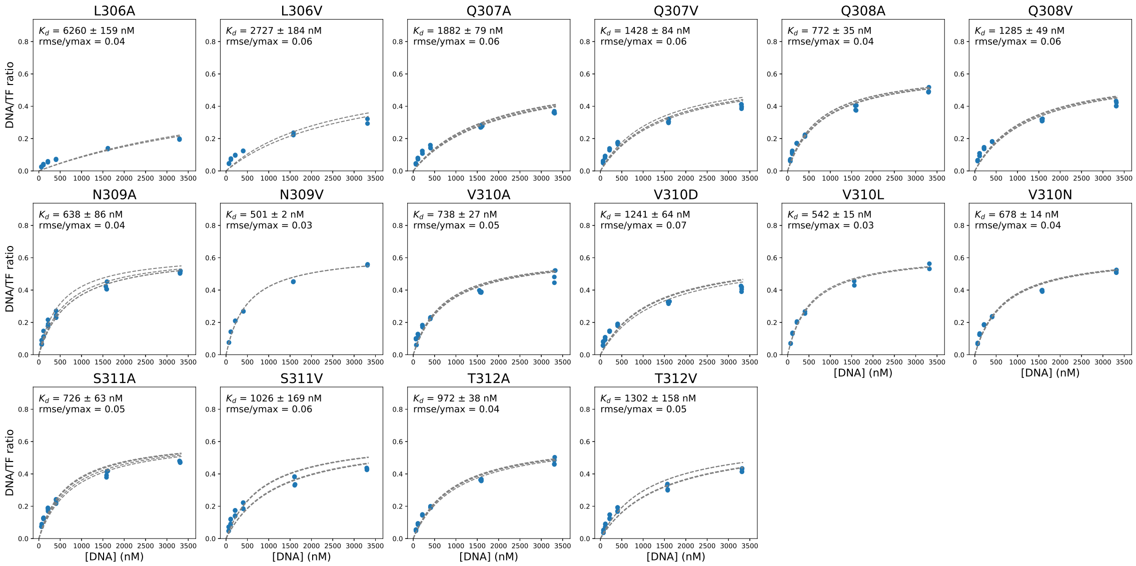

Figure 51: STAMMP binding curves (continued)

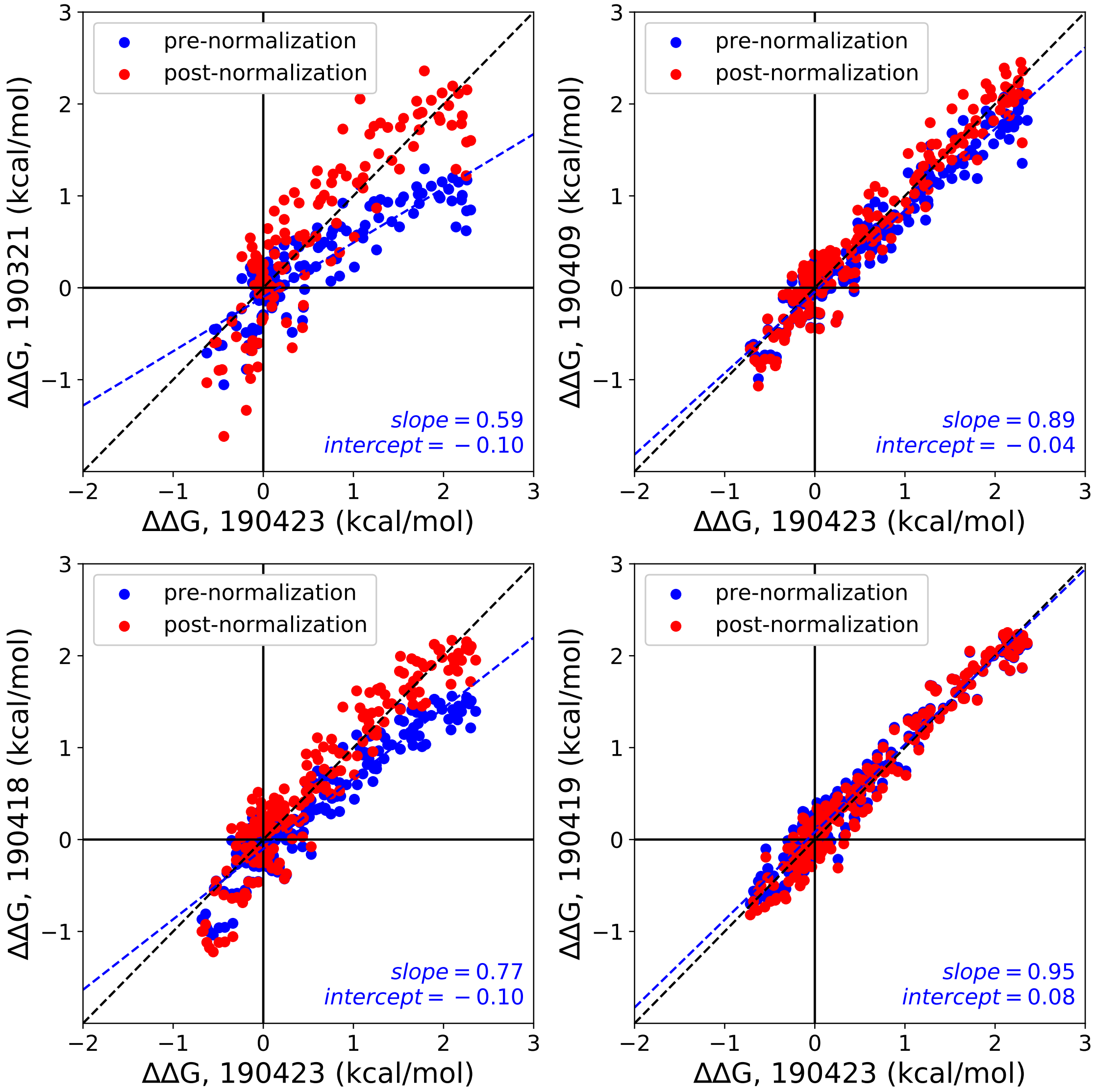

Figure 52: Comparisons between ΔΔG measurements for 214 Pho4 variants binding to a DNA sequence containing an E-box motif surrounded by random sequence across 5 different experiments before (blue markers) and after (red markers) linear normalization (see Supplementary Methods). Blue line shows a linear regression to data prior to normalization; blue text indicates pre-normalization regression parameters; black dashed line indicates the 1:1 identity line.

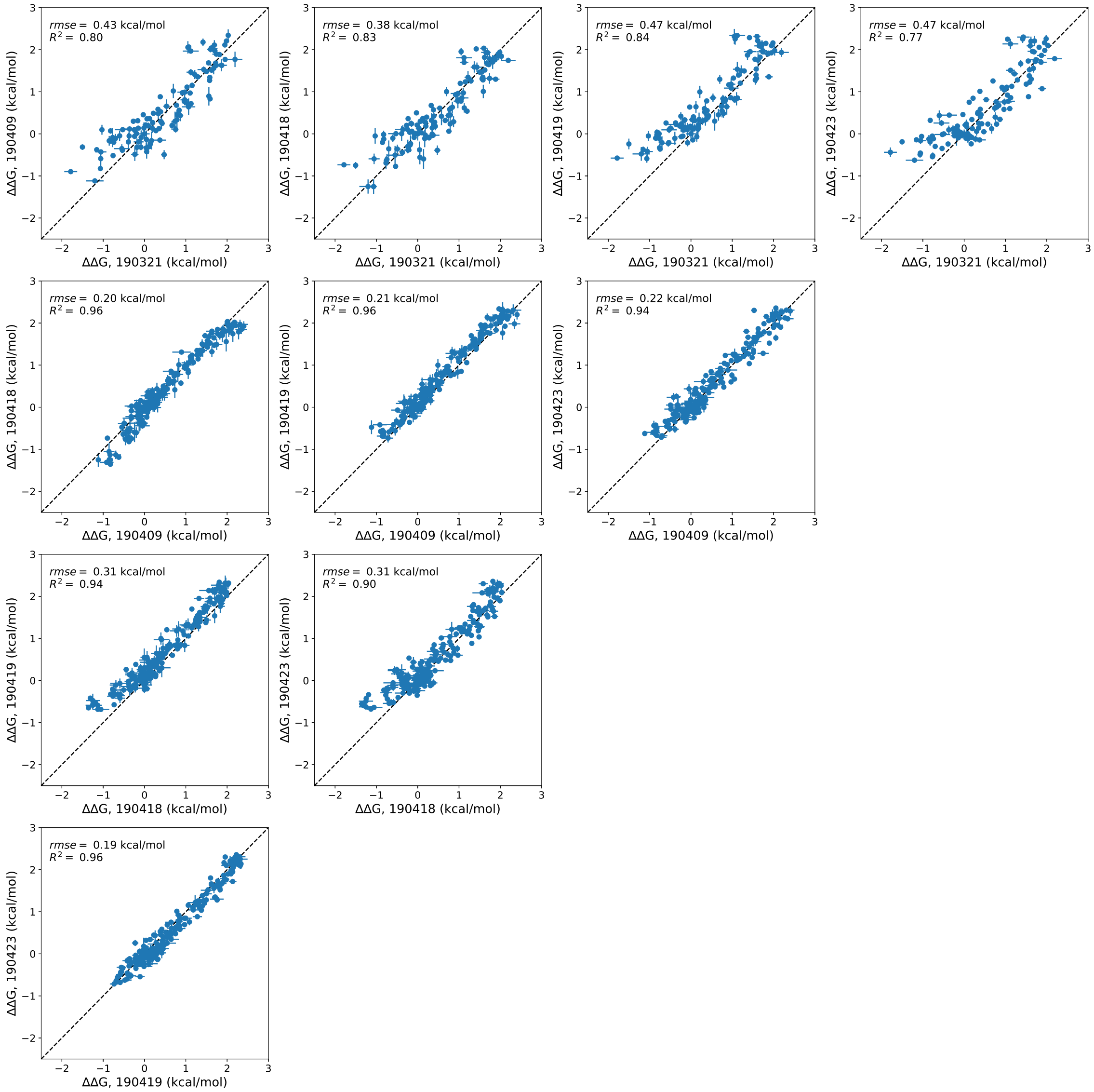

Figure 53: Pairwise comparisons of normalized ΔΔG values between 5 STAMMP experiments quantifying Pho4 mutants binding to a dsDNA sequence containing an E-box motif surrounded by random sequence. Markers indicate median across all replicates within a given experiment; error bars indicate standard deviation of measured values.

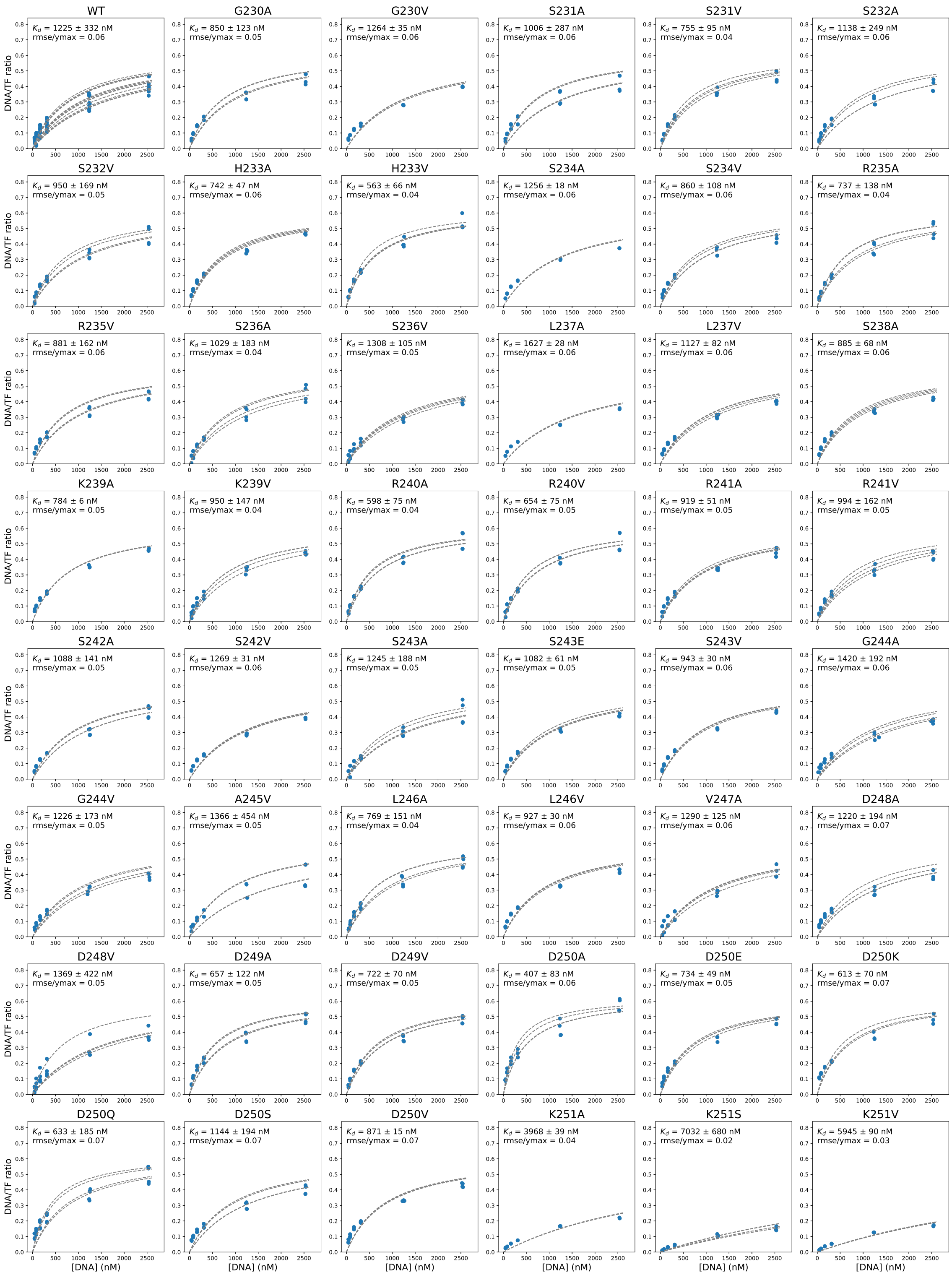

Figure 54: Example binding curves for STAMMP experiments assessing binding of 214 Pho4 variants passing QC to a DNA sequence containing a CACGTG E-box surrounded by CG/AT repeats sequence from experiment on 04/23/2019. Markers indicate per-chamber fluorescence intensity ratios (Alexa-647/eGFP); dashed lines indicates per-chamber fits to a Langmuir isotherm. Binding curves for other experiments are available on OSF (https://osf.io/gbxhz/).

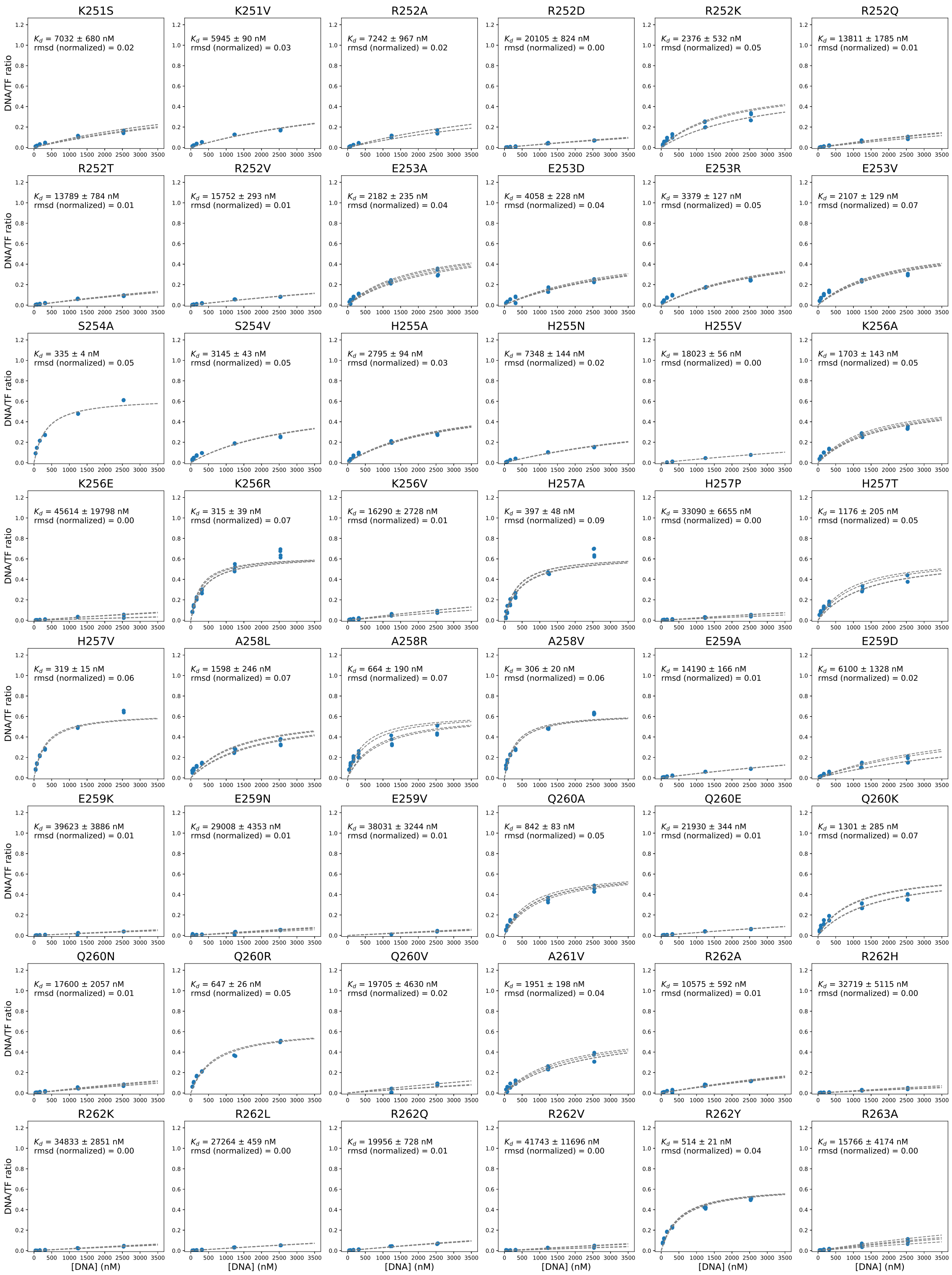

Figure 54: STAMMP binding curves (continued)

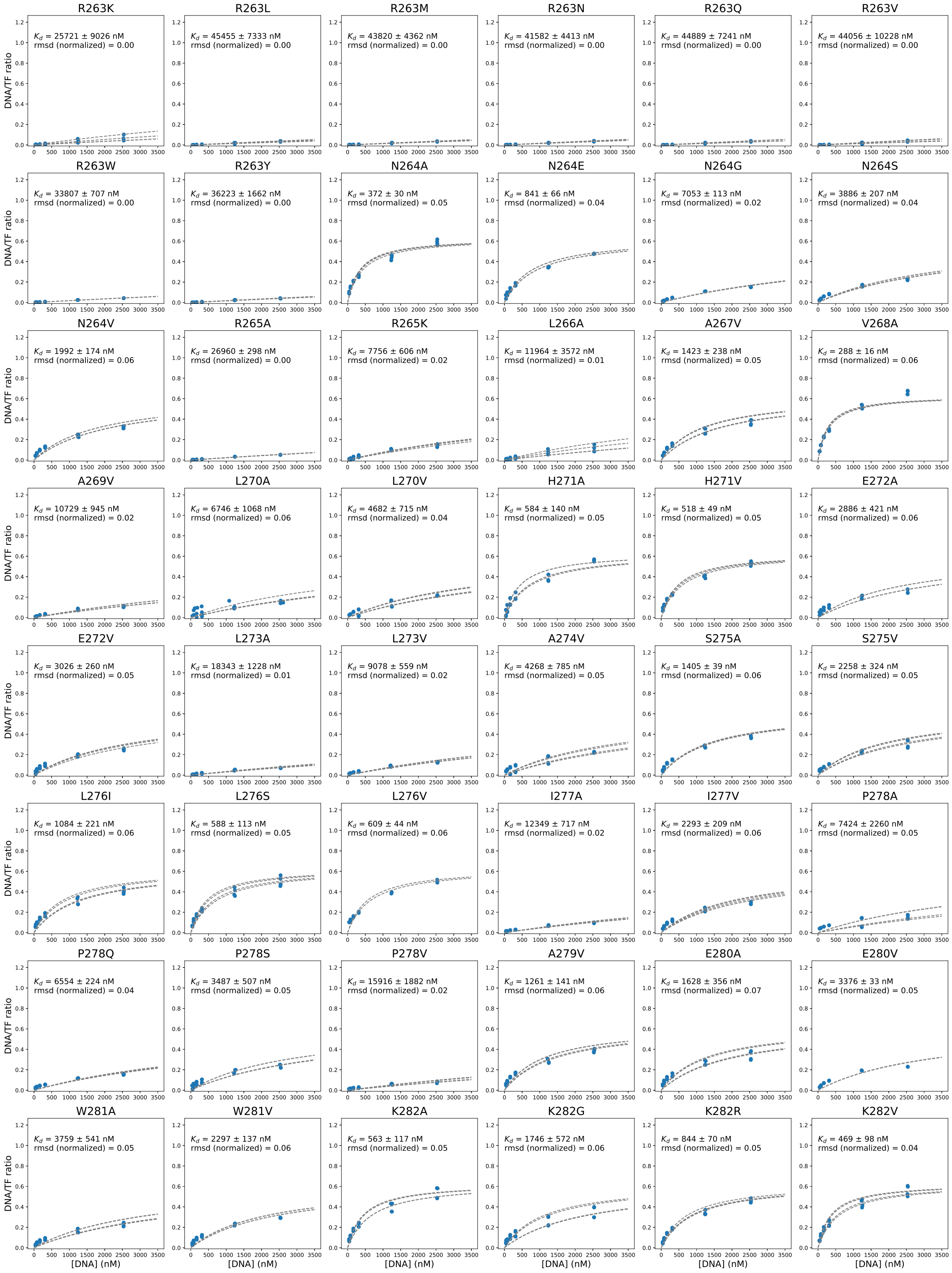

Figure 54: STAMMP binding curves (continued)

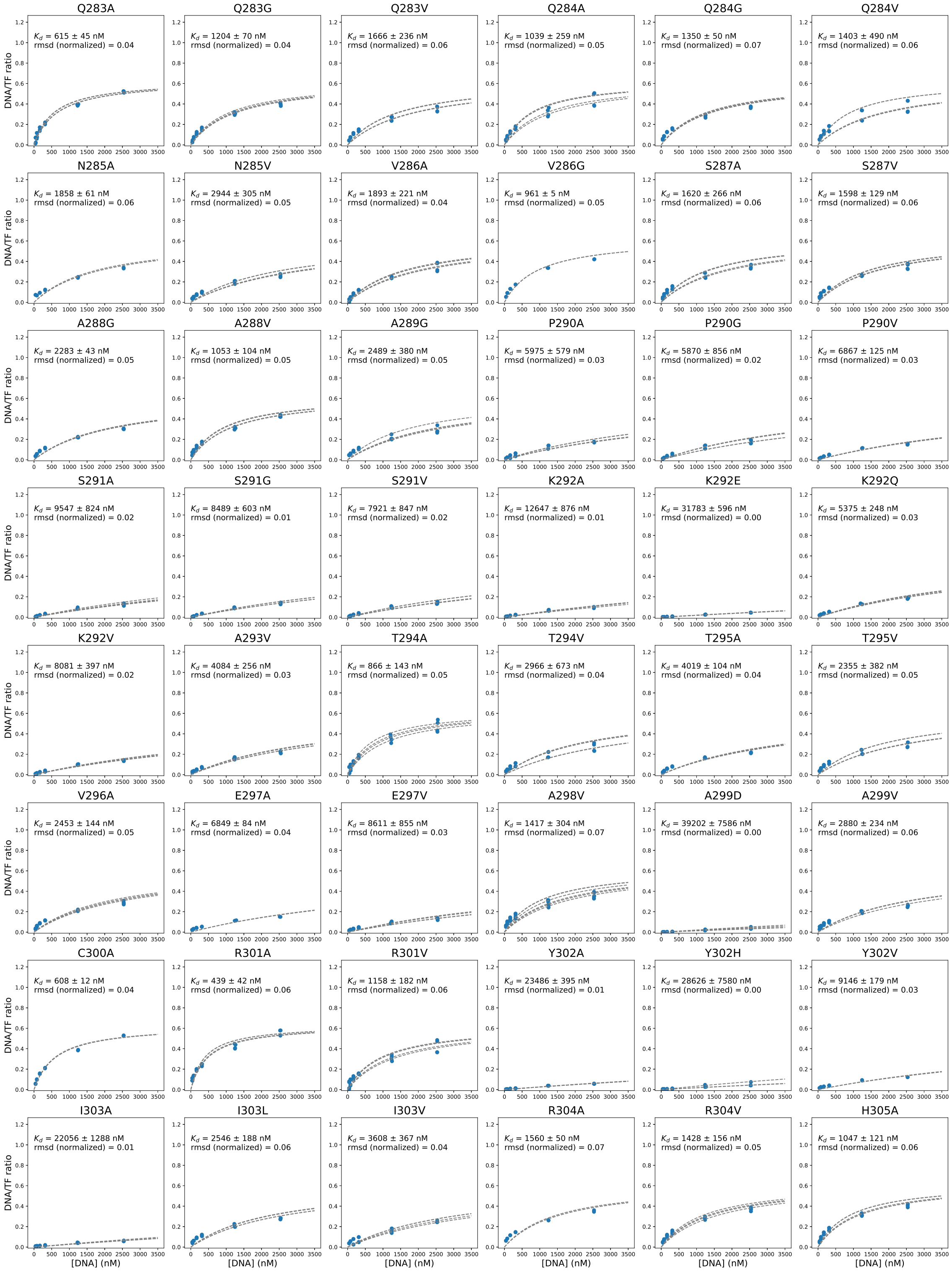

Figure 54: STAMMP binding curves (continued)

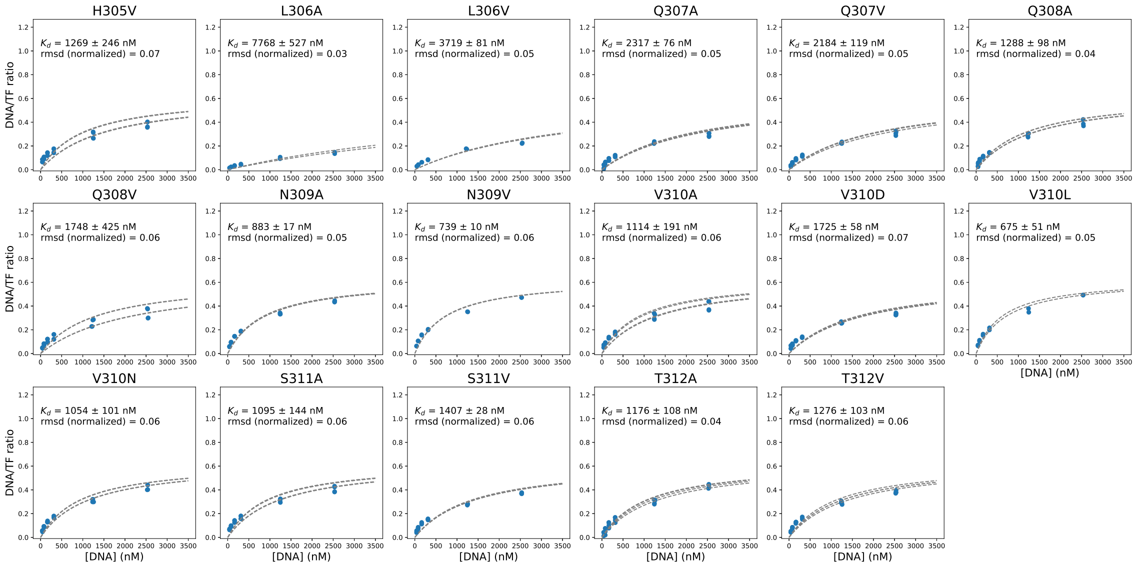

Figure 54: STAMMP binding curves (continued)

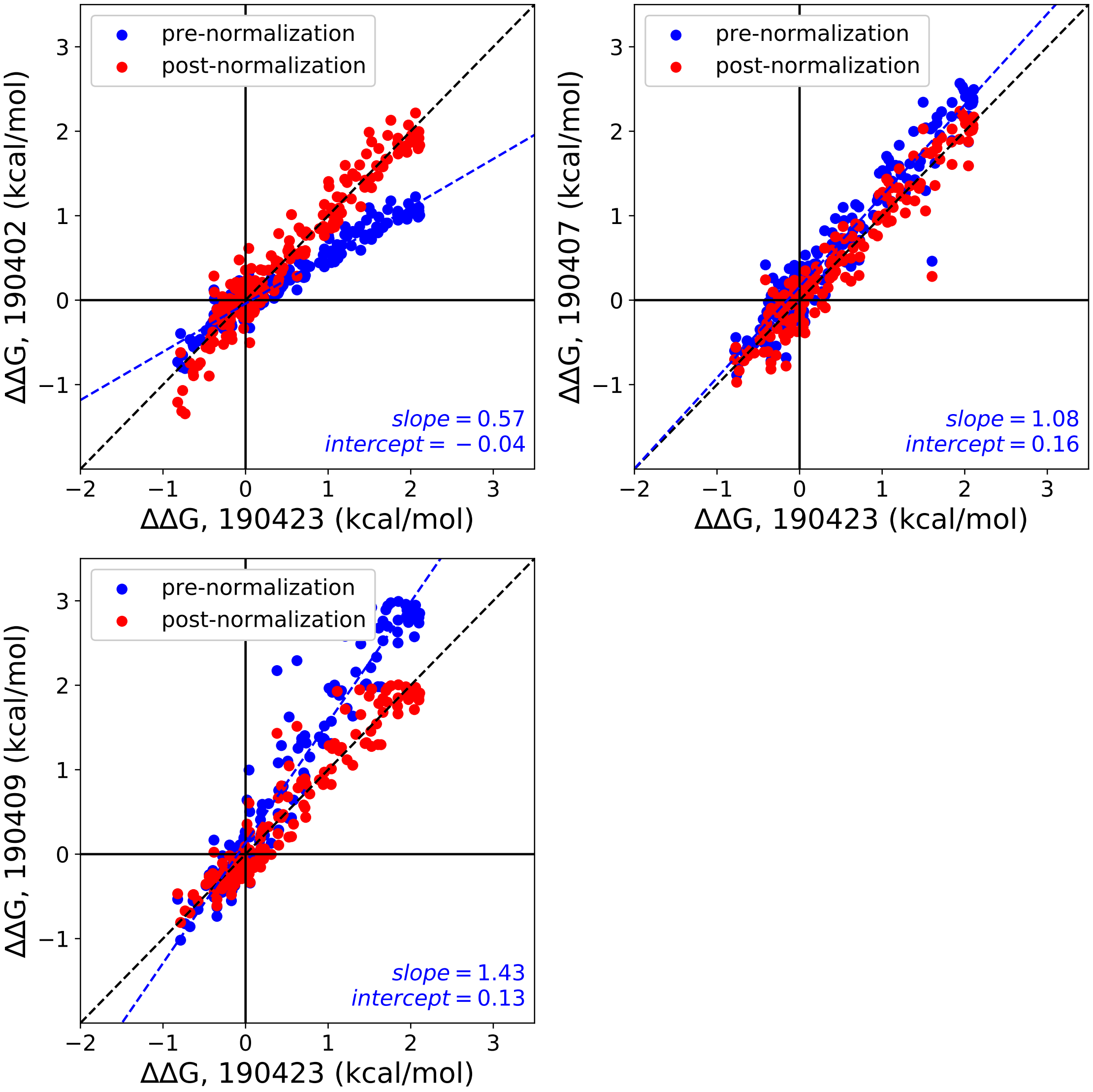

Figure 55: Comparisons between ΔΔG measurements for 214 Pho4 variants binding to a DNA sequence containing an E-box motif surrounded by CG/AT repeats across 4 different experiments before (blue markers) and after (red markers) linear normalization (see Supplementary Methods). Blue line shows a linear regression to data prior to normalization; blue text indicates pre-normalization regression parameters; black dashed line indicates the 1:1 identity line.

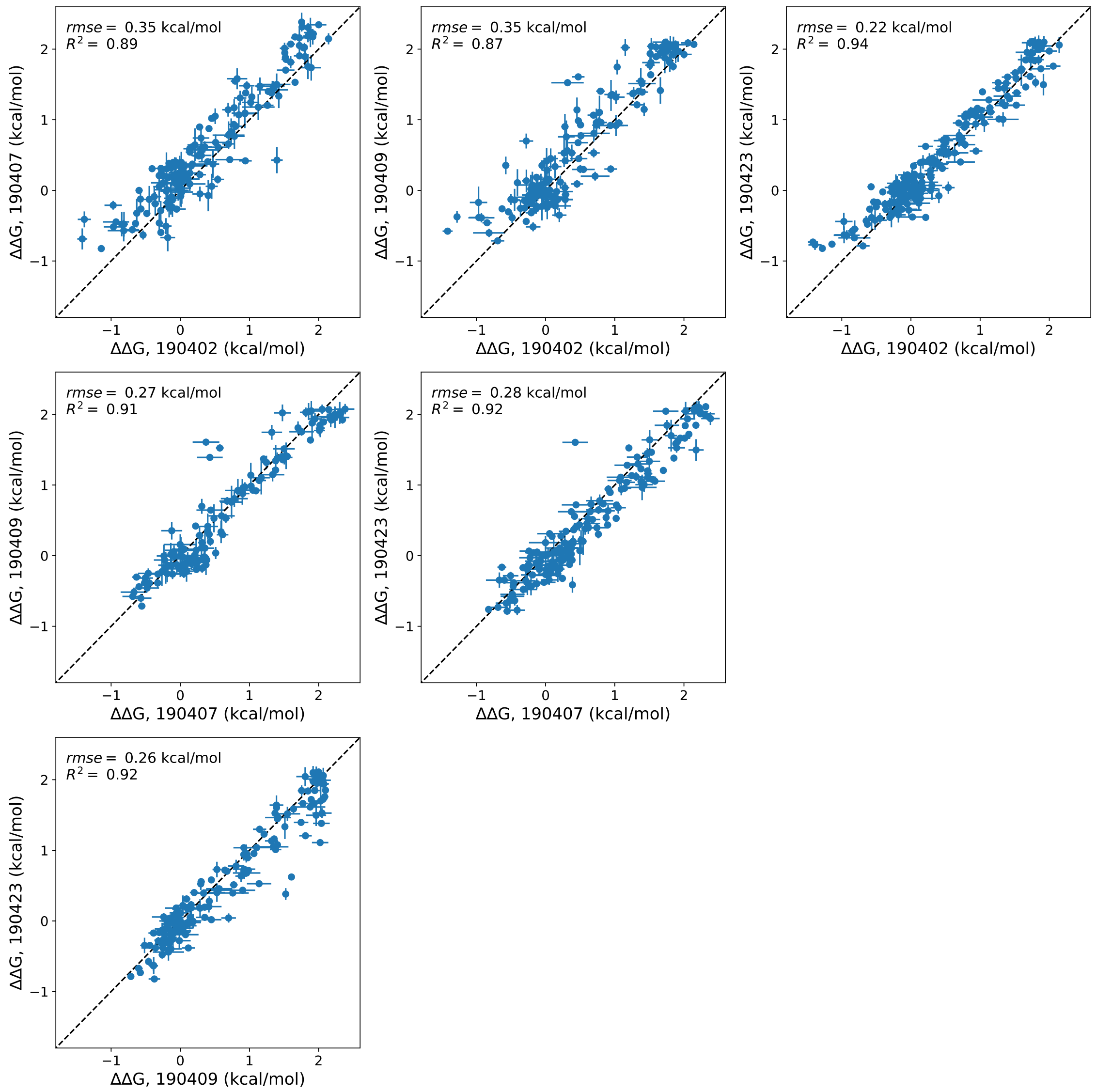

Figure 56: Pairwise comparisons of normalized ΔΔG values between 4 STAMMP experiments quantifying Pho4 mutants binding to a dsDNA sequence containing an E-box motif surrounded by GC repeats. Markers indicate median across all replicates within a given experiment; error bars indicate standard deviation of measured values.

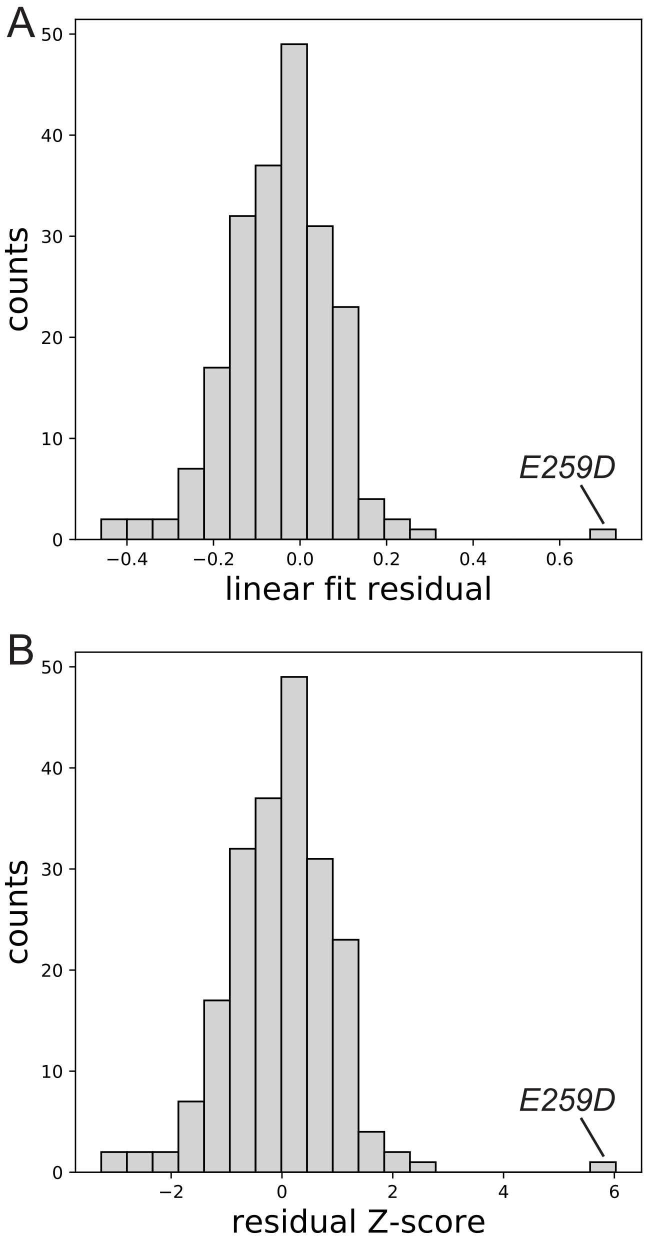

Figure 57: Calculated residuals **(A)** and residual Z-scores **(B)** from the 1:1 identity line for measured $\Delta\Delta$Gs for all Pho4 mutant variants interacting with DNA sequences containing a central E-box surrounded by either repetitive or random flanking sequences.

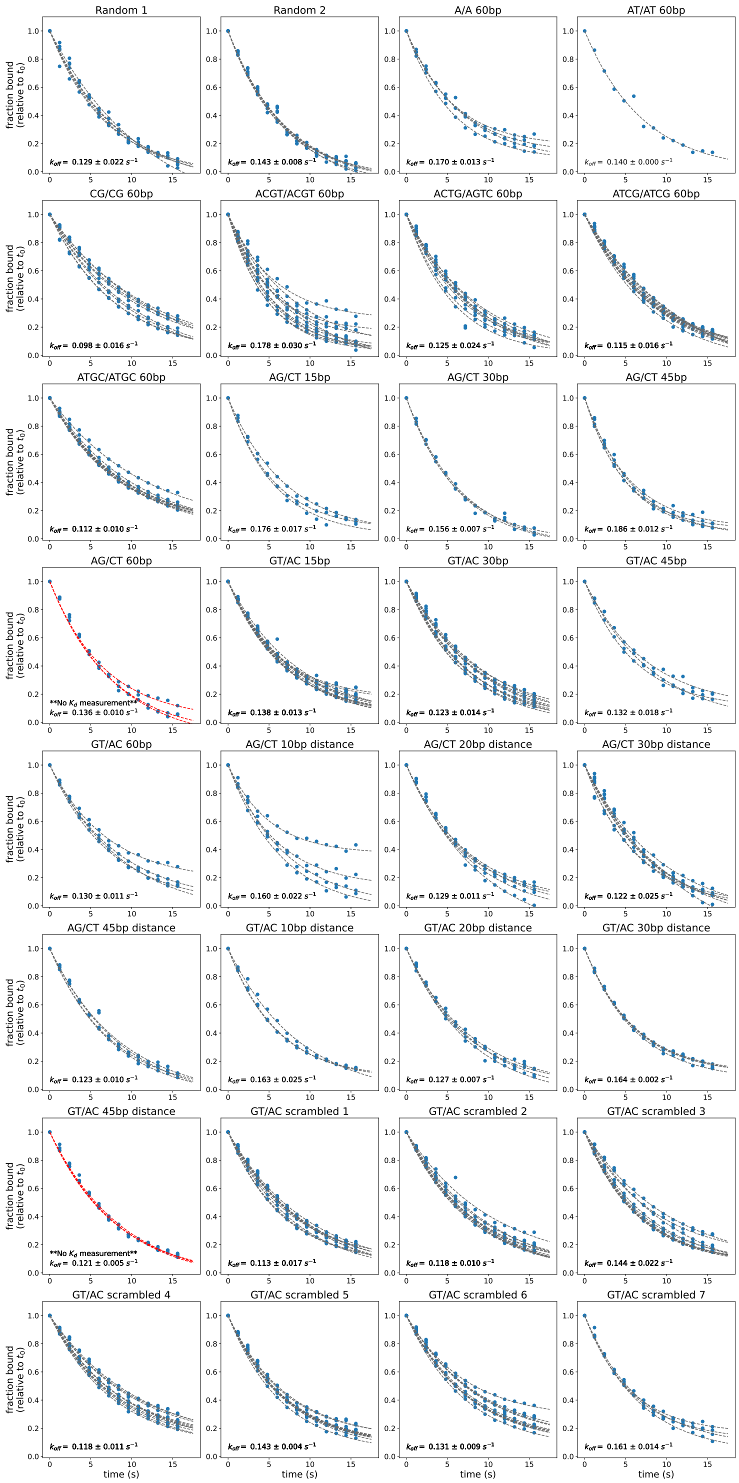

Figure 58: Example dissociation measurements for MAX interacting with DNA Library 2 sequences from experiment on 10/25/2019. Markers indicate per-chamber fluorescence ratio of Alexa647-labeled dsDNA to GFP-tagged TF; dashed lines indicate single exponential decay fits for data from each chamber. Sequences with no corresponding $K_d$ measurement from this experiment have fit lines shown in red. Dissociation curves for other experiments are available on OSF (https://osf.io/gbxhz/).

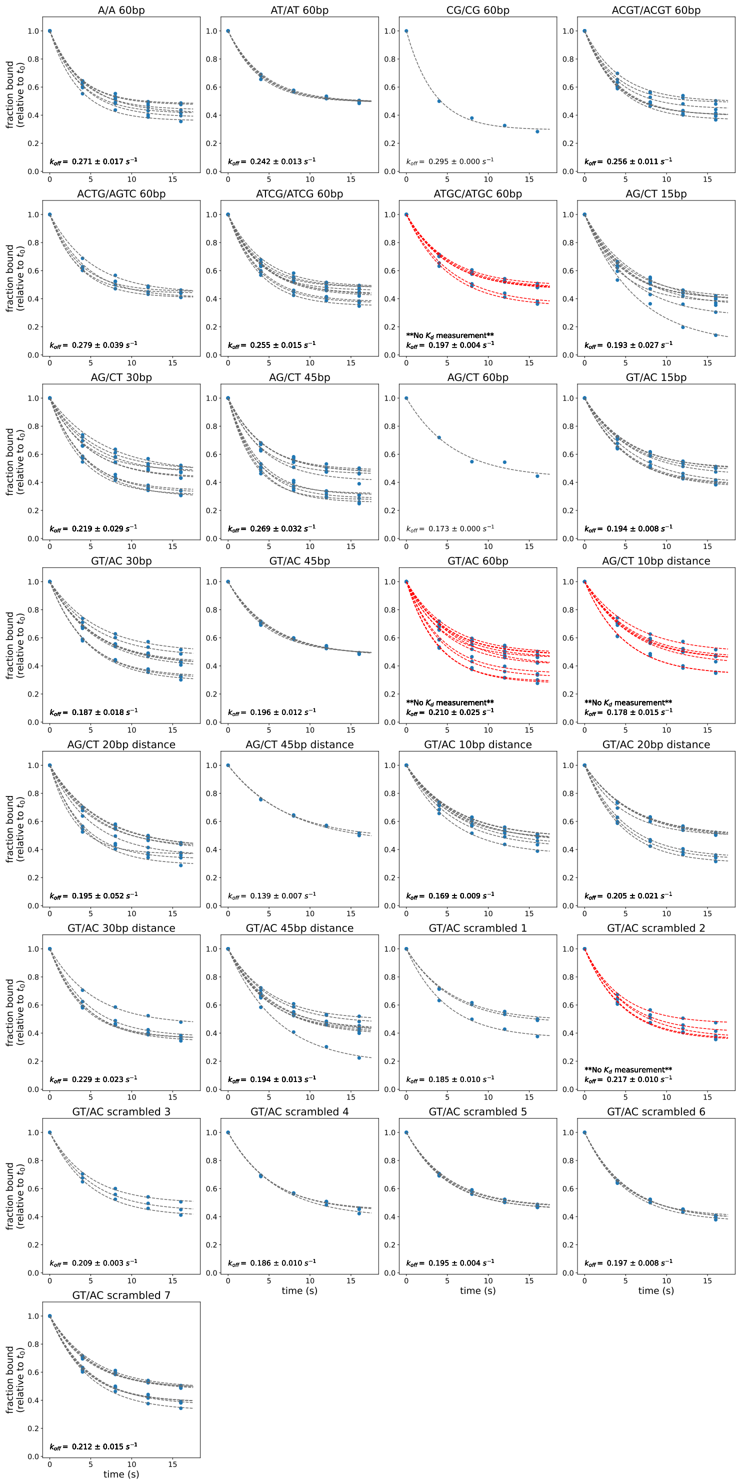

Figure 59: Example dissociation measurements for Pho4 interacting with DNA Library 2 sequences from experiment on 11/04/2019. Markers indicate per-chamber TF-bound Alexa-647 intensities; lines indicate single exponential decay fits for data from each chamber. Sequences with no corresponding $K_d$ measurement from this experiment have fit lines shown in red. Dissociation curves for other experiments are available on OSF (https://osf.io/gbxhz/).

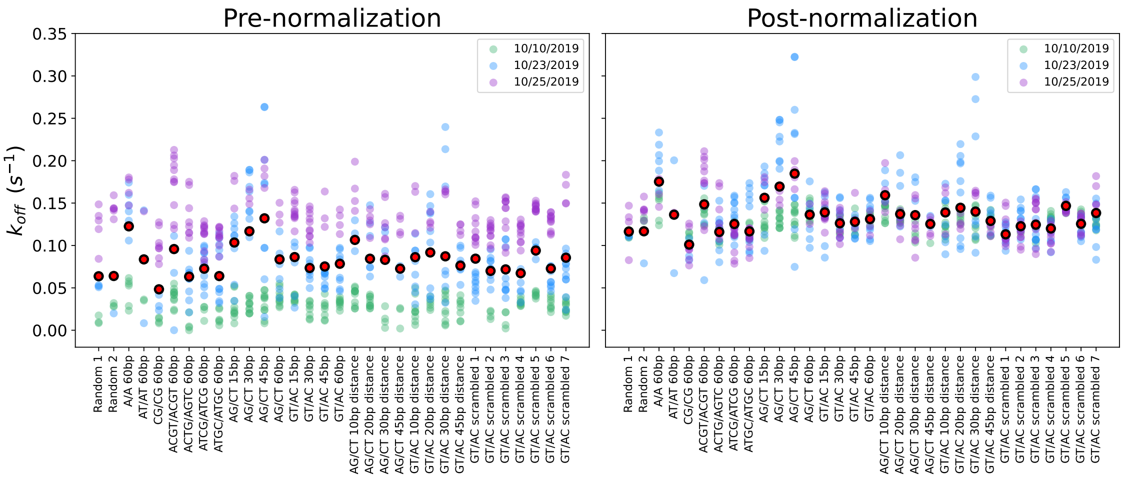

Figure 60: Comparisons between $k_{off}$ measurements for MAX binding to DNA Library 2 across 3 different experiments before and after linear normalization (see Supplementary Methods). Black points with red outline represent median $k_{off}$ across all experiments.

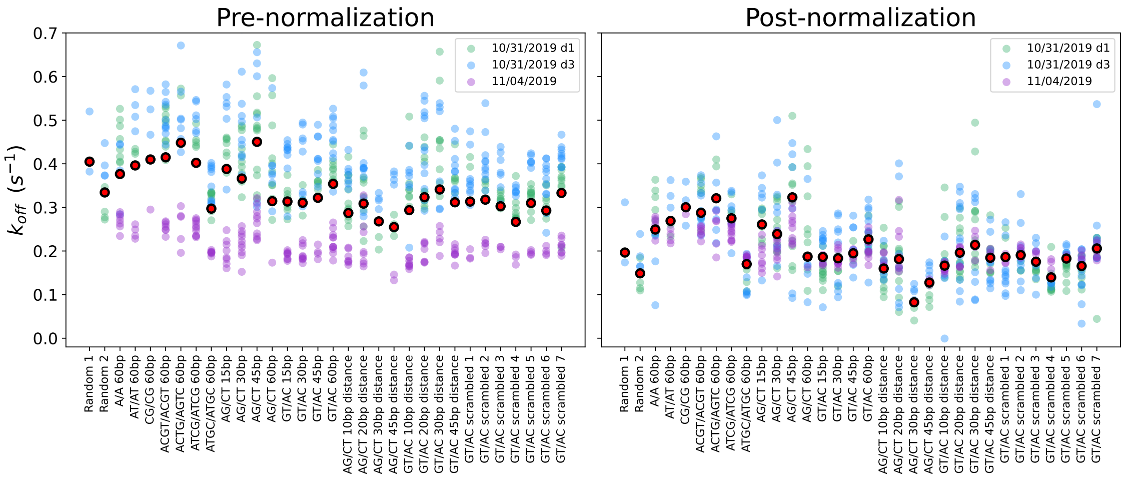

Figure 61: Comparisons between $k_{off}$ measurements for Pho4 binding to DNA Library 2 across 3 different experiments before (blue markers) and after (red markers) linear normalization (see Supplementary Methods). Black points with red outline represent median $k_{off}$ across all experiments.

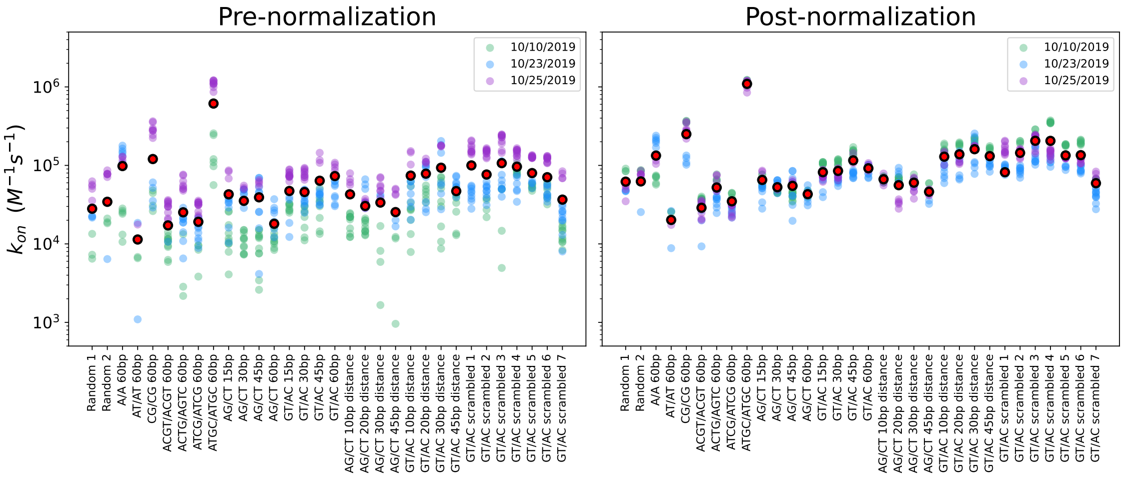

Figure 62: Comparisons between inferred $k_{on}$ values for MAX binding to DNA Library 2 across 3 different experiments before and after linear normalization of $k_{off}$ values (see Supplementary Methods). Black points with red outline represent median $k_{on}$ across all experiments.

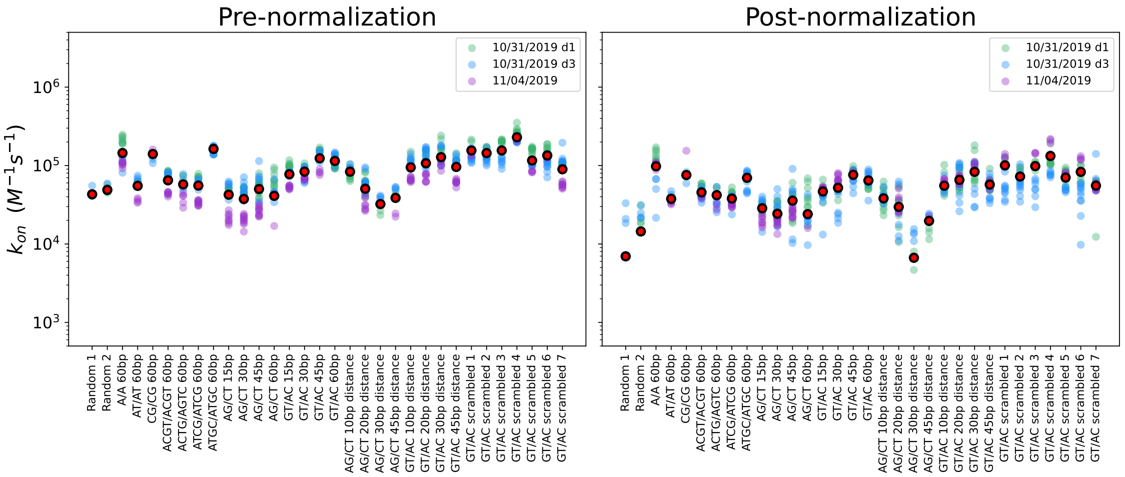

Figure 63: Comparisons between inferred $k_{on}$ values for Pho4 binding to DNA Library 2 across 3 different experiments before and after linear normalization of $k_{off}$ values (see Supplementary Methods). Black points with red outline represent median $k_{on}$ across all experiments.

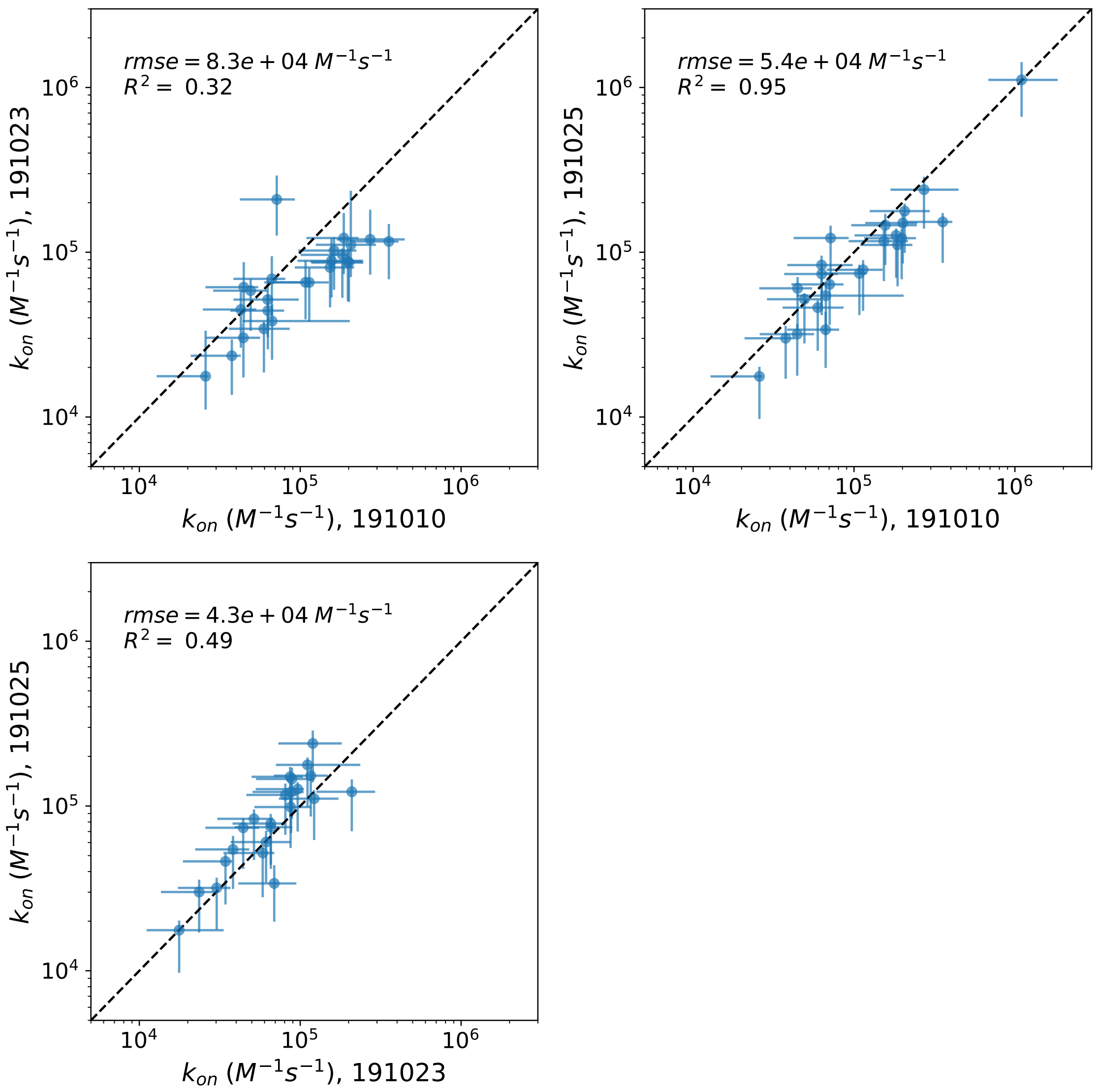

Figure 64: Pairwise comparisons of calculated $k_{on}$ values for MAX by sequence across 3 experiments quantifying MAX association rate to DNA Library 2. Markers indicate mean across all replicates within a given experiment; error bars indicate standard deviation of measured values. Lower bounds of error bars adjusted for log scale.

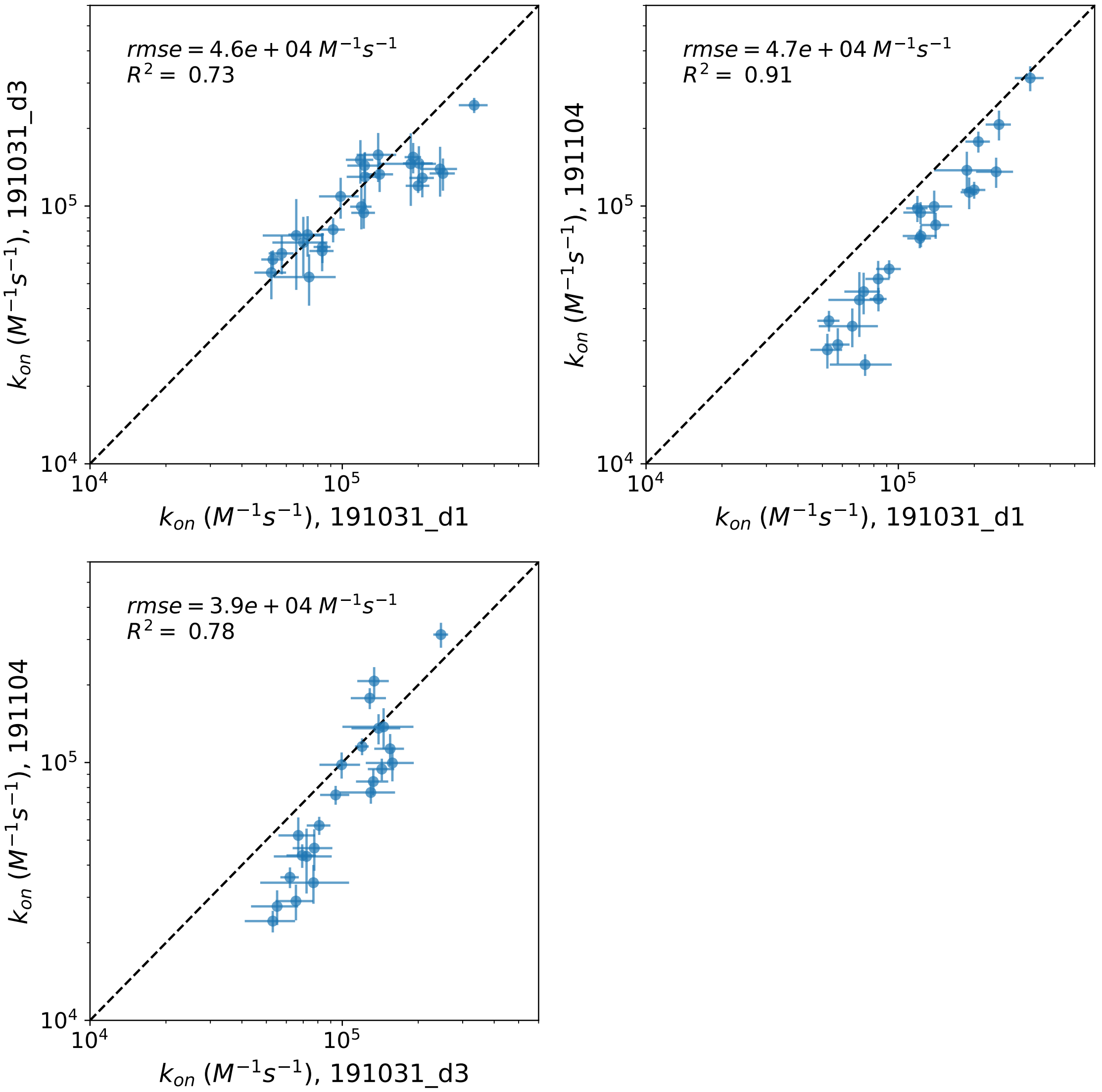

Figure 65: Pairwise comparisons of calculated $k_{on}$ values for Pho4 by sequence across 3 experiments quantifying Pho4 association rate to DNA Library 2. Markers indicate mean across all replicates within a given experiment; error bars indicate standard deviation of measured values.

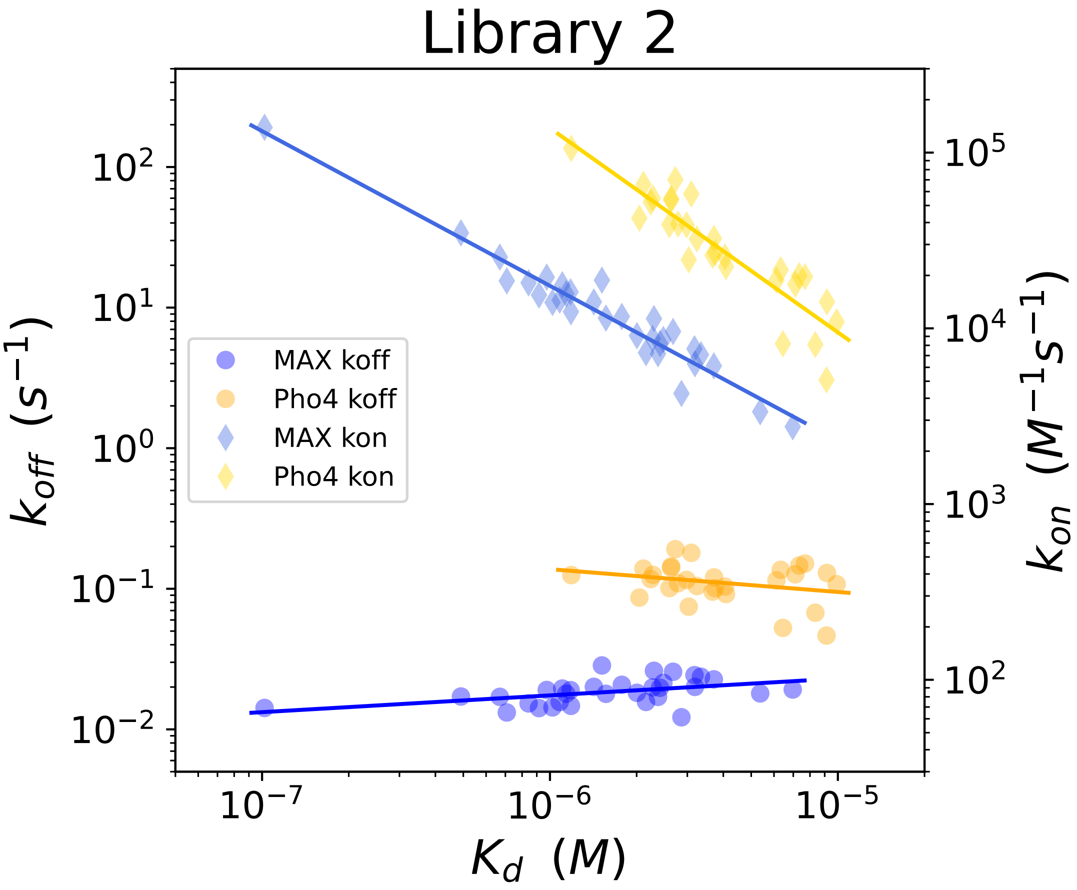

Figure 66: Average measured $k_{off}$ (circle markers, left axis) and calculated $k_{on}$ (diamond markers, right axis) values *vs.* measured affinities ($K_d$s) for Pho4 (yellow/orange) and MAX (blue) interacting with all sequences from DNA Library 2.

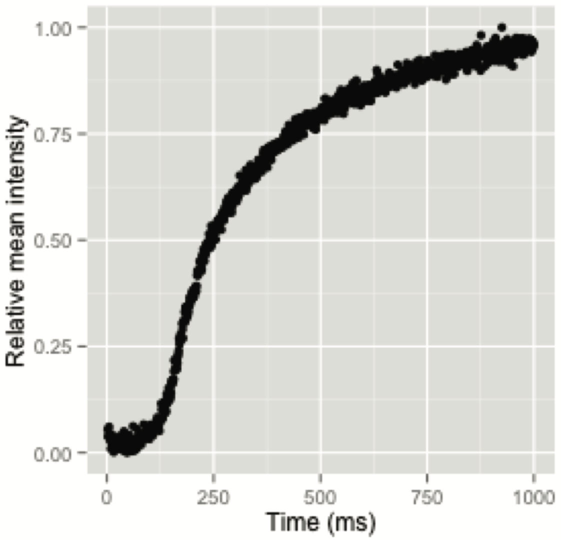

Figure 67: Quantification of MITOMI button valve opening kinetics demonstrates that button valves take approximately 1 second to open completely. As described in Section 1.16.5, 16 button valves were imaged continuously for 1 second at 1000 frames per second after sending a near-instantaneous signal to depressurize valves. Relative mean intensity is a measure of button state, where a value of 1 corresponds to a completely open button valve and a value of 0 corresponds to a completely closed valve. Each point is a measurement for a single button valve in a single imaging frame.

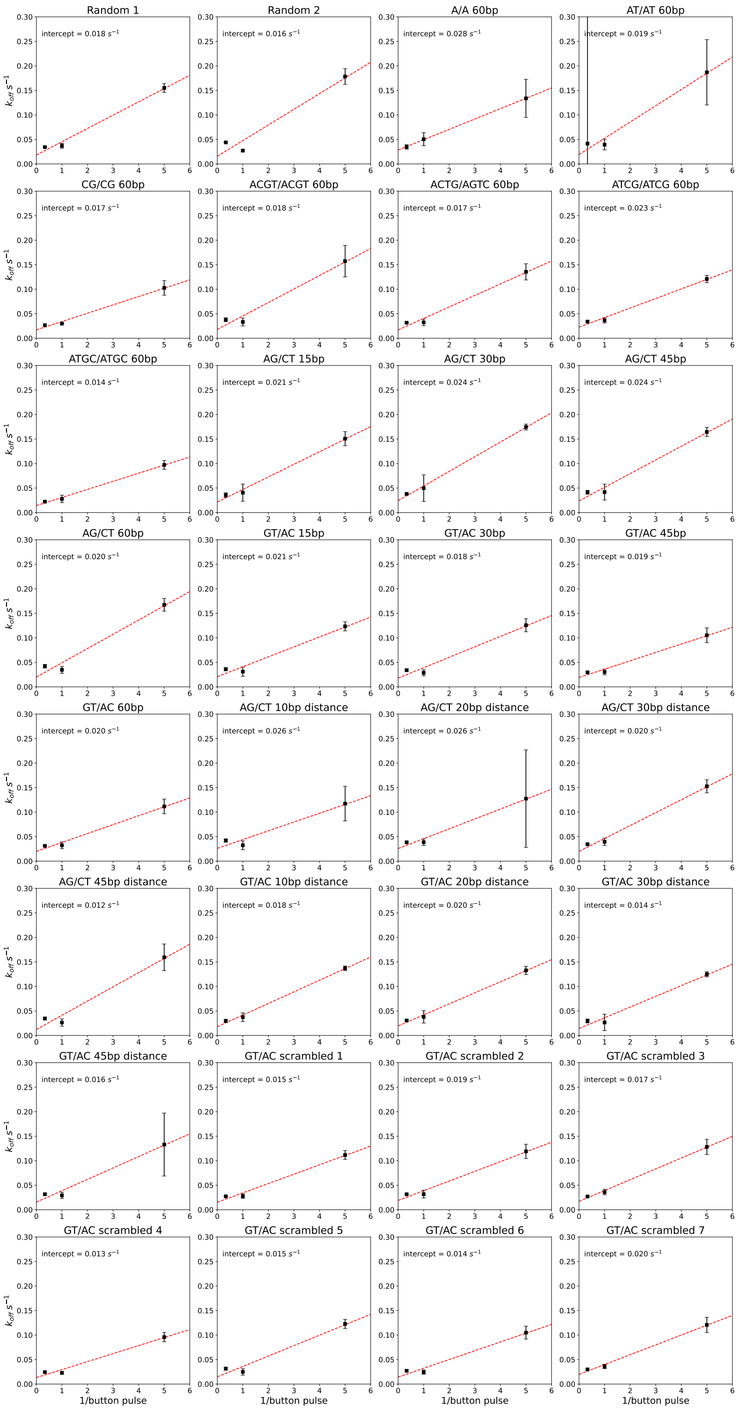

Figure 68: Measured $k_{off}$ values for MAX dissociating from oligonucleotides in DNA Library 2 *vs.* 1/button pulse, where button pulse is the length of time button valves are left open and oligos are allowed to dissociate. 1 second is subtracted from the programmed button pulse time to account for button opening delay, as shown in Figure S67. Points and error bars show median $k_{off}$ fits $\pm$ standard deviation for a given oligo per experiment. Red dashed lines show linear fits, where the $y$-intercept (indicated in top-left corner of each plot) is interpreted as the 'true' $k_{off}$ value in the absence of mechanical shear. This normalization procedure is described in Section 1.16.5.

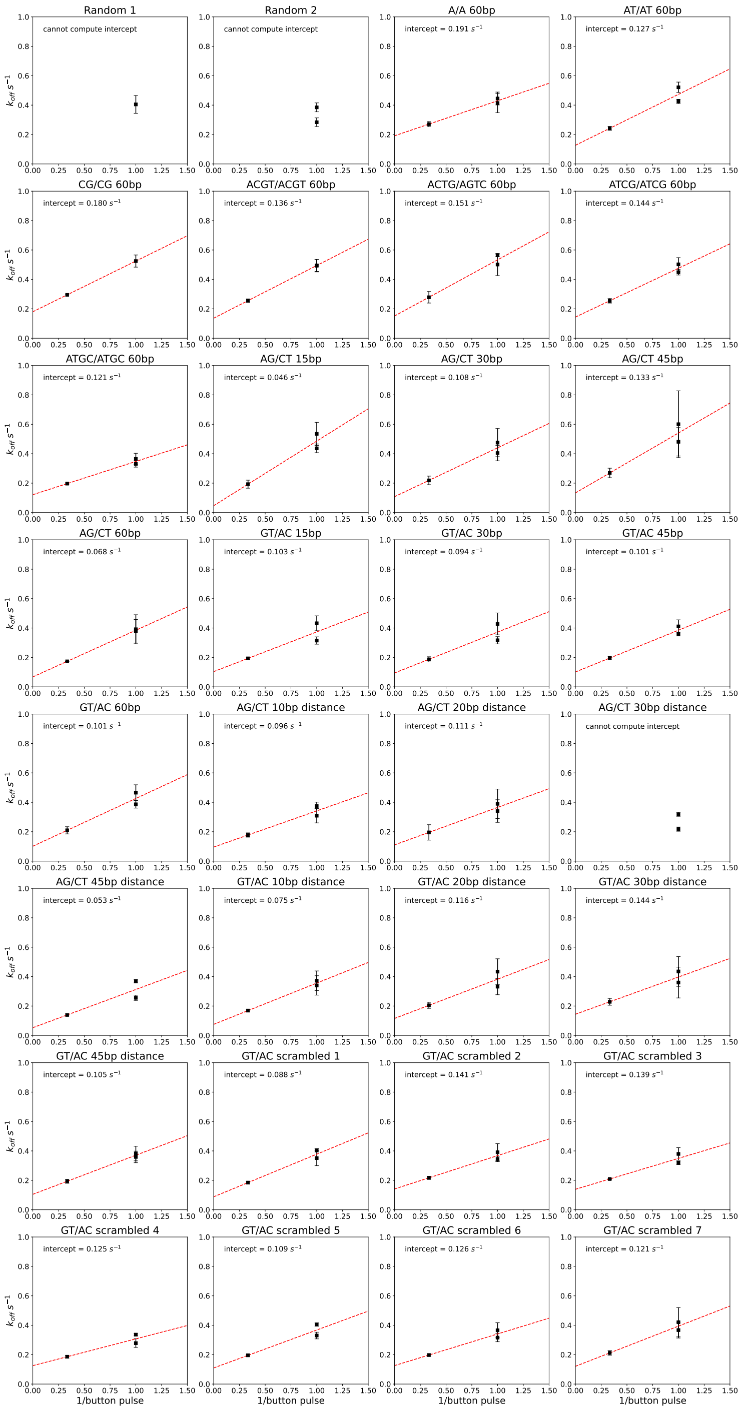

Figure 69: Measured $k_{off}$ values for Pho4 dissociating from oligonucleotides in DNA Library 2 *vs.* 1/button pulse, where button pulse is the length of time button valves are left open and oligos are allowed to dissociate. 1 second is subtracted from the programmed button pulse time to account for button opening delay, as shown in Figure S67. Points and error bars show median $k_{off}$ fits ± standard deviation for a given oligo per experiment. Red dashed lines show linear fits, where the $y$-intercept (indicated in top-left corner of each plot) is interpreted as the 'true' $k_{off}$ value in the absence of mechanical shear. This normalization procedure is described in Section 1.16.5.

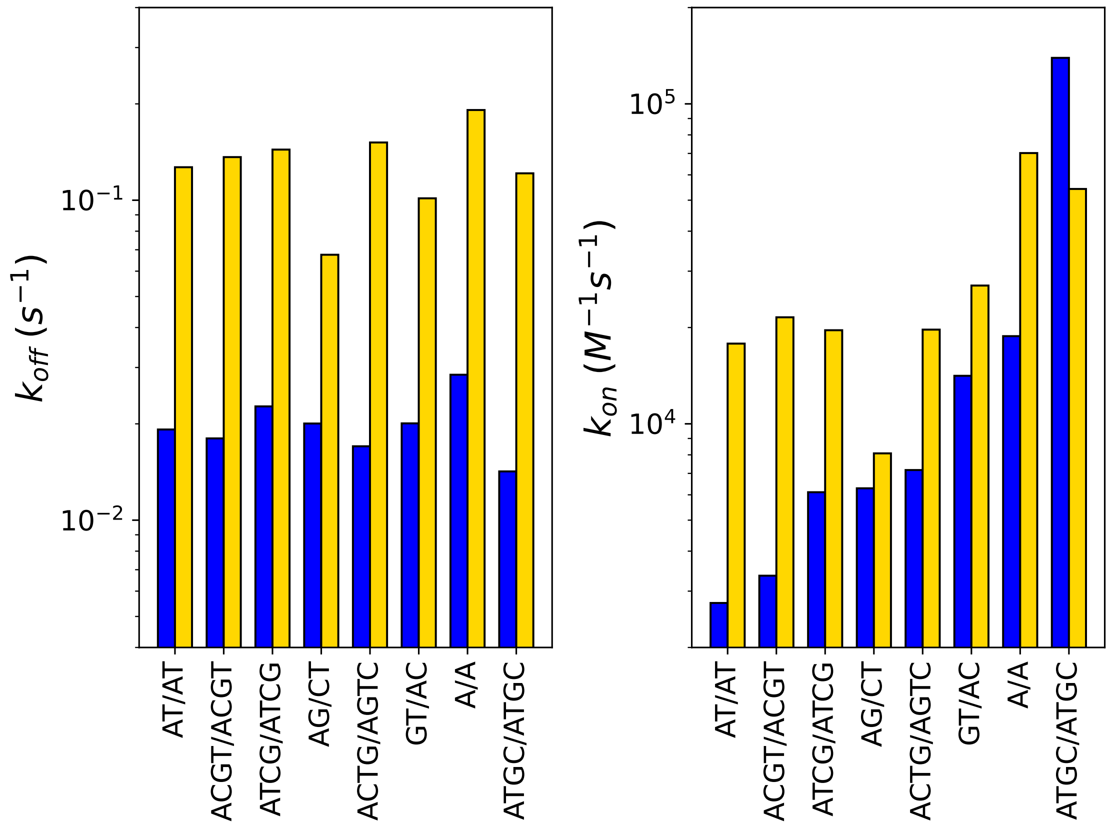

Figure 70: Measured $k_{off}$ (left, circle markers) and calculated $k_{on}$ (right, diamond markers) values as a function of flanking sequence for Pho4 (yellow) and MAX (blue) using an alternative normalization procedure described in Section 1.16.5.

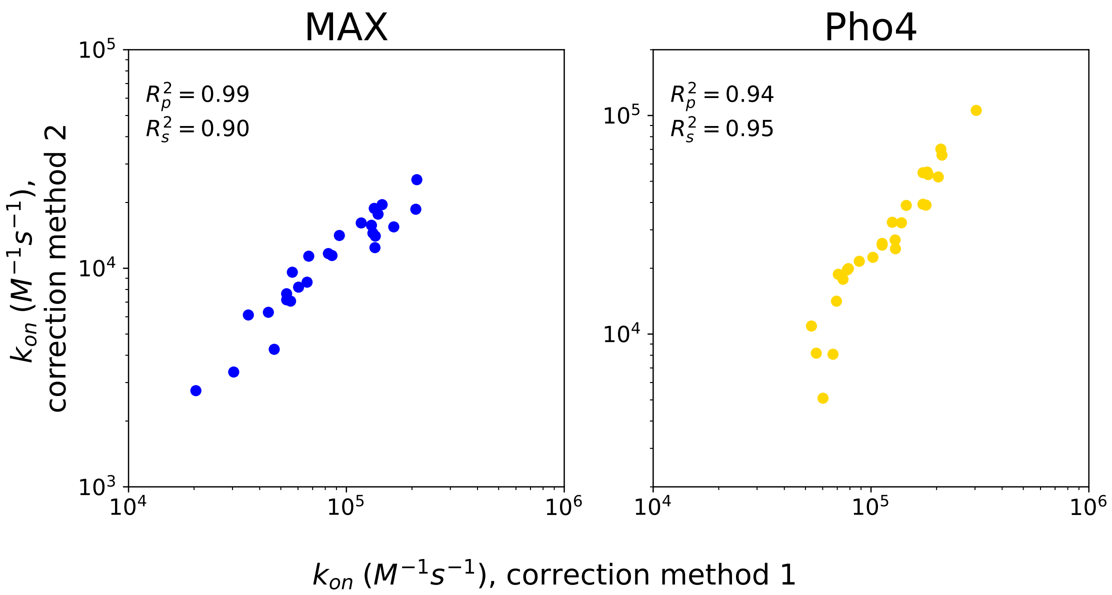

Figure 71: Different normalization strategies yield nearly identical inferred $k_{on}$ values for MAX (left) and Pho4 (right) binding to DNA Library 2. $x$-axis shows $k_{on}$ values calculated from normalization method described in Section 1.16.4 and shown in main text Fig. 4C. $y$-axis shows $k_{on}$ values calculated from normalization method described in Section 1.16.5 and shown in Fig. S70.

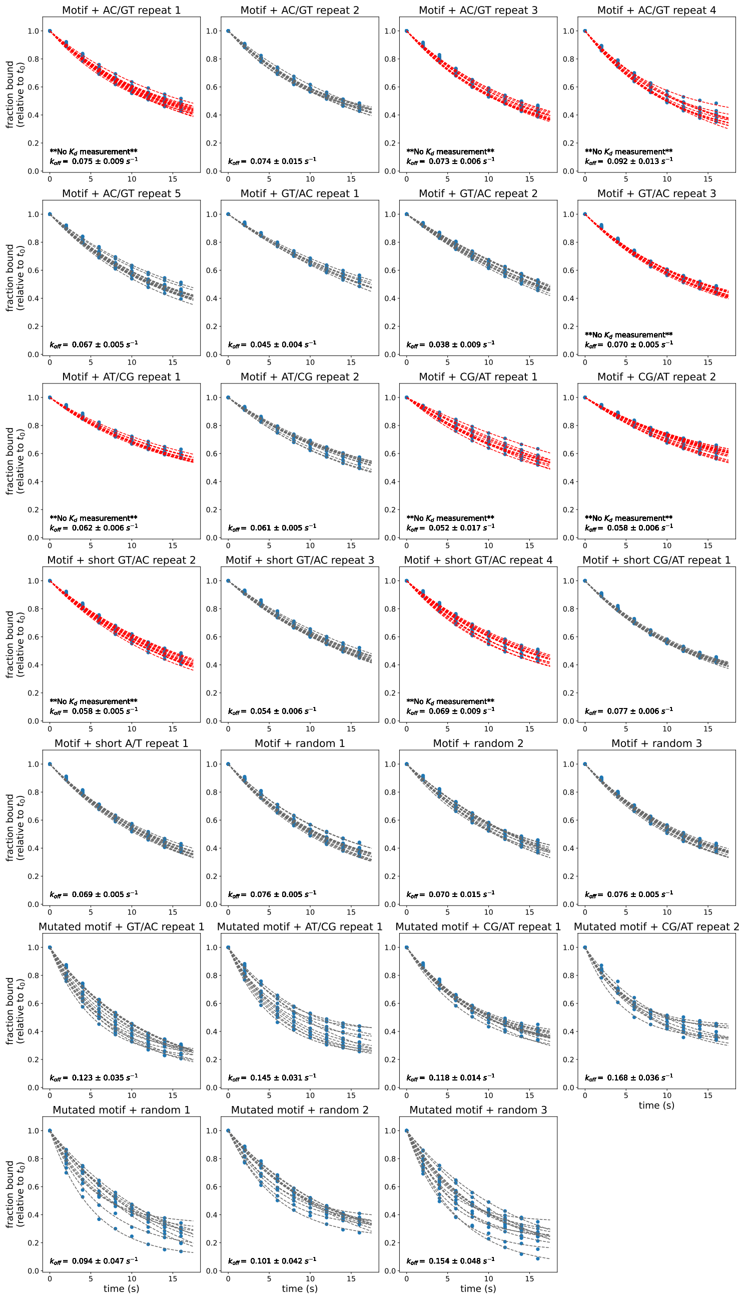

Figure 72: Example dissociation measurements for MAX interacting with DNA Library 1 sequences from experiment on 12/06/2018. Markers indicate per-chamber fluorescence ratio of Alexa647-labeled dsDNA to GFP-tagged TF; lines indicate single exponential decay fits for data from each chamber. Sequences with no corresponding $K_d$ measurement from this experiment have fit lines shown in red. Dissociation curves for other experiments are available on OSF (https://osf.io/gbxhz/).

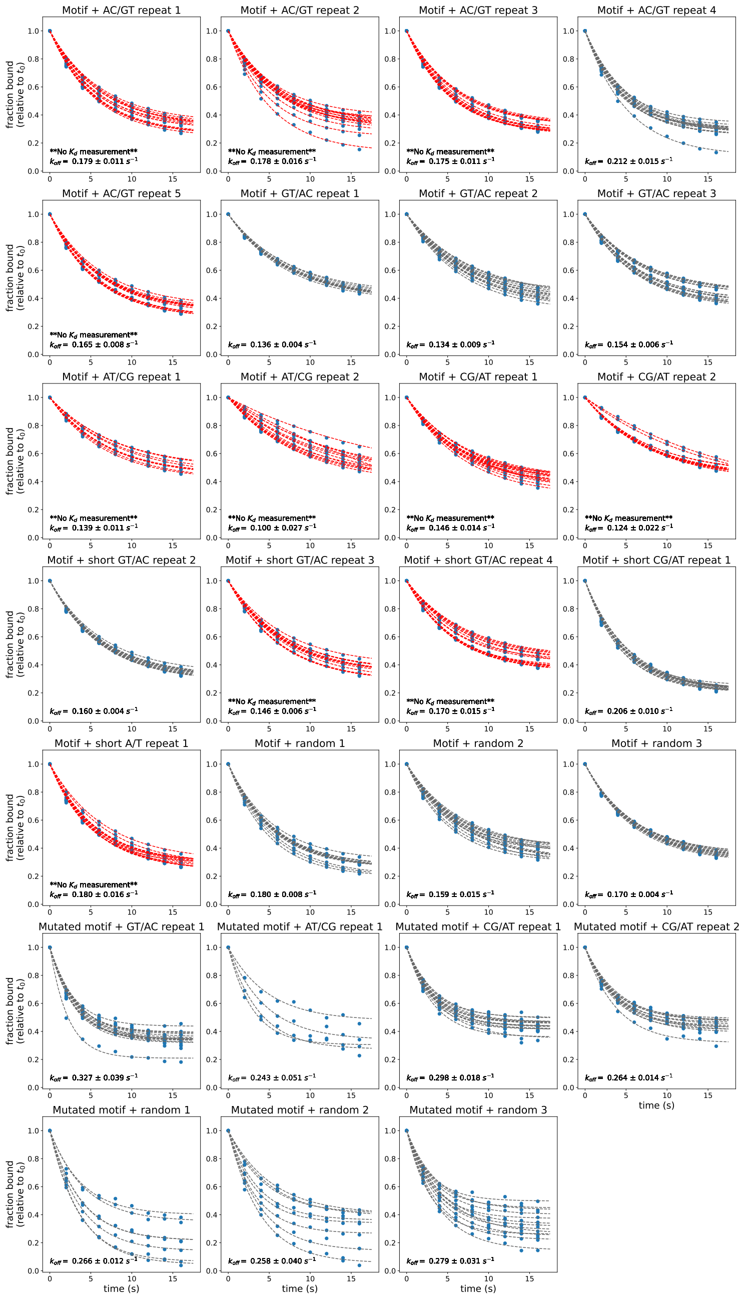

Figure 73: Example dissociation measurements for Pho4 interacting with DNA Library 1 sequences from experiment on 12/17/2018. Markers indicate per-chamber TF-bound Alexa-647 intensities; lines indicate single exponential decay fits for data from each chamber. Sequences with no corresponding $K_d$ measurement from this experiment have fit lines shown in red. Dissociation curves for other experiments are available on OSF (https://osf.io/gbxhz/).

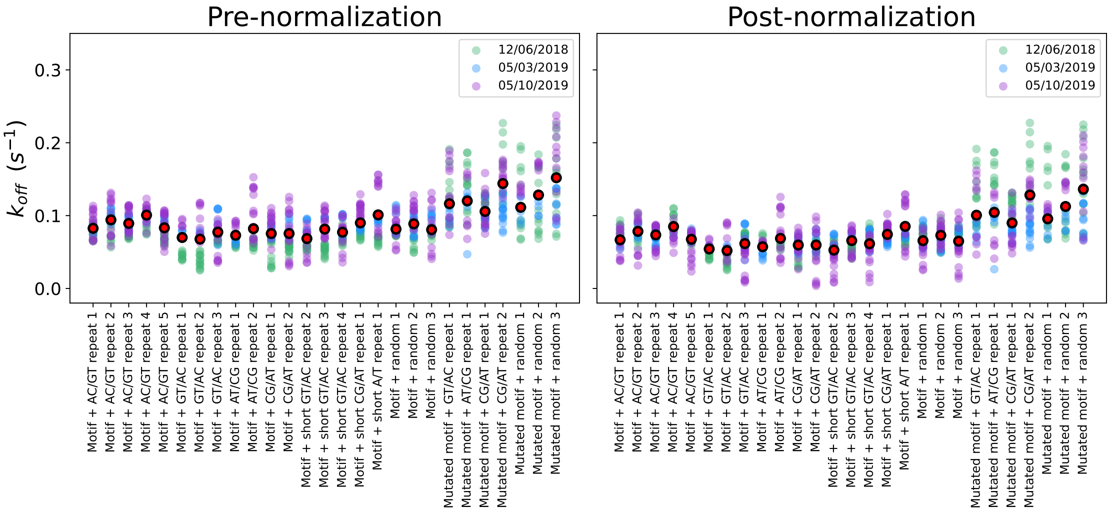

Figure 74: Comparisons between $k_{off}$ measurements for MAX binding to DNA Library 1 across 3 different experiments before and after linear normalization (see Supplementary Methods). Black points with red outline represent median $k_{off}$ across all experiments.

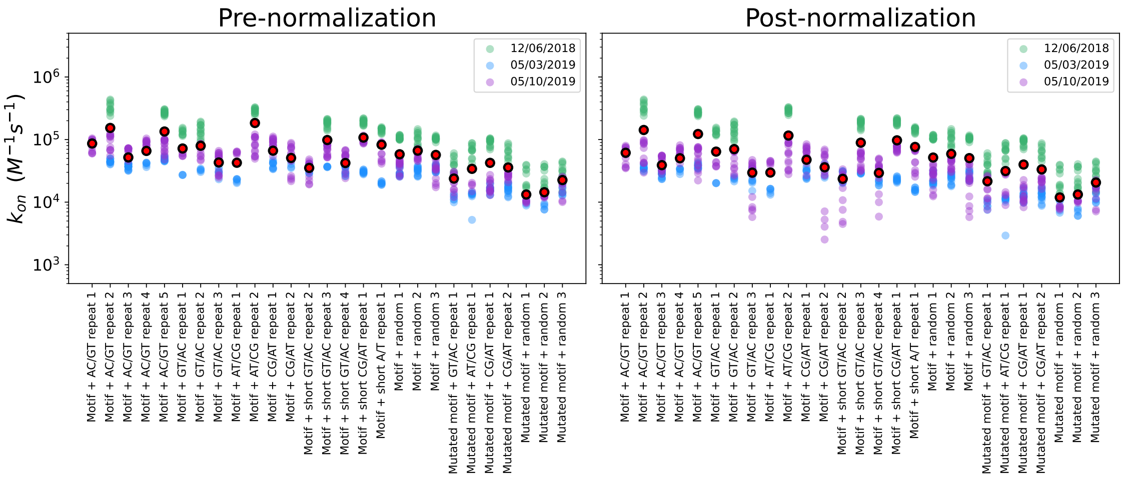

Figure 75: Comparisons between inferred $k_{on}$ values for MAX binding to DNA Library 1 across 3 different experiments before and after linear normalization of $k_{off}$ values (see Supplementary Methods). Black points with red outline represent median $k_{on}$ across all experiments.

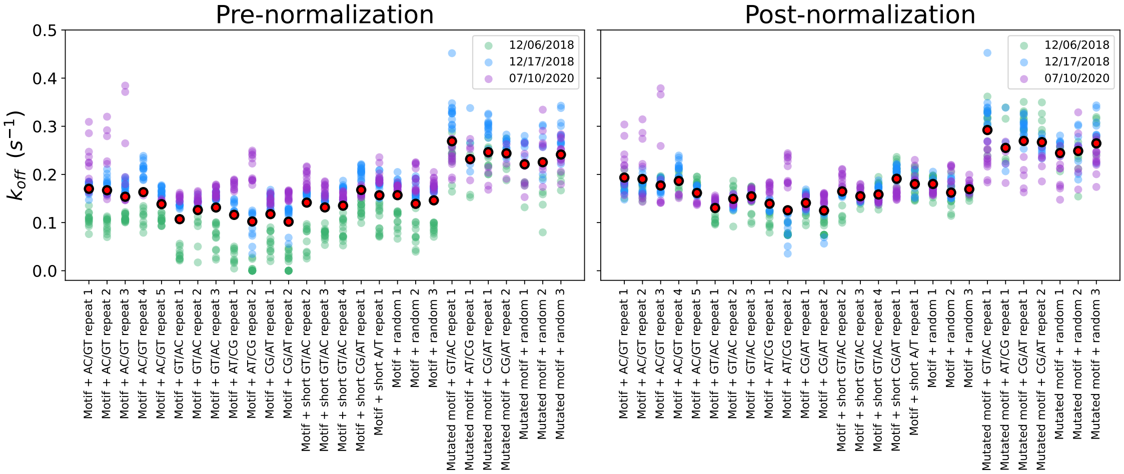

Figure 76: Comparisons between $k_{off}$ measurements for Pho4 binding to DNA Library 1 across 3 different experiments before and after linear normalization (see Supplementary Methods). Black points with red outline represent median $k_{off}$ across all experiments.

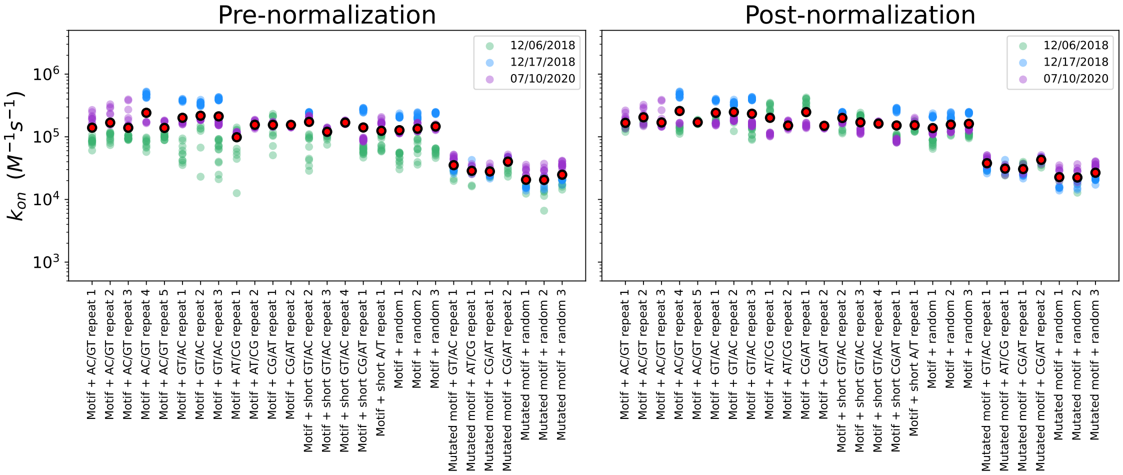

Figure 77: Comparisons between inferred $k_{on}$ values for Pho4 binding to DNA Library 1 across 3 different experiments before and after linear normalization of $k_{off}$ values (see Supplementary Methods). Black points with red outline represent median $k_{on}$ across all experiments.

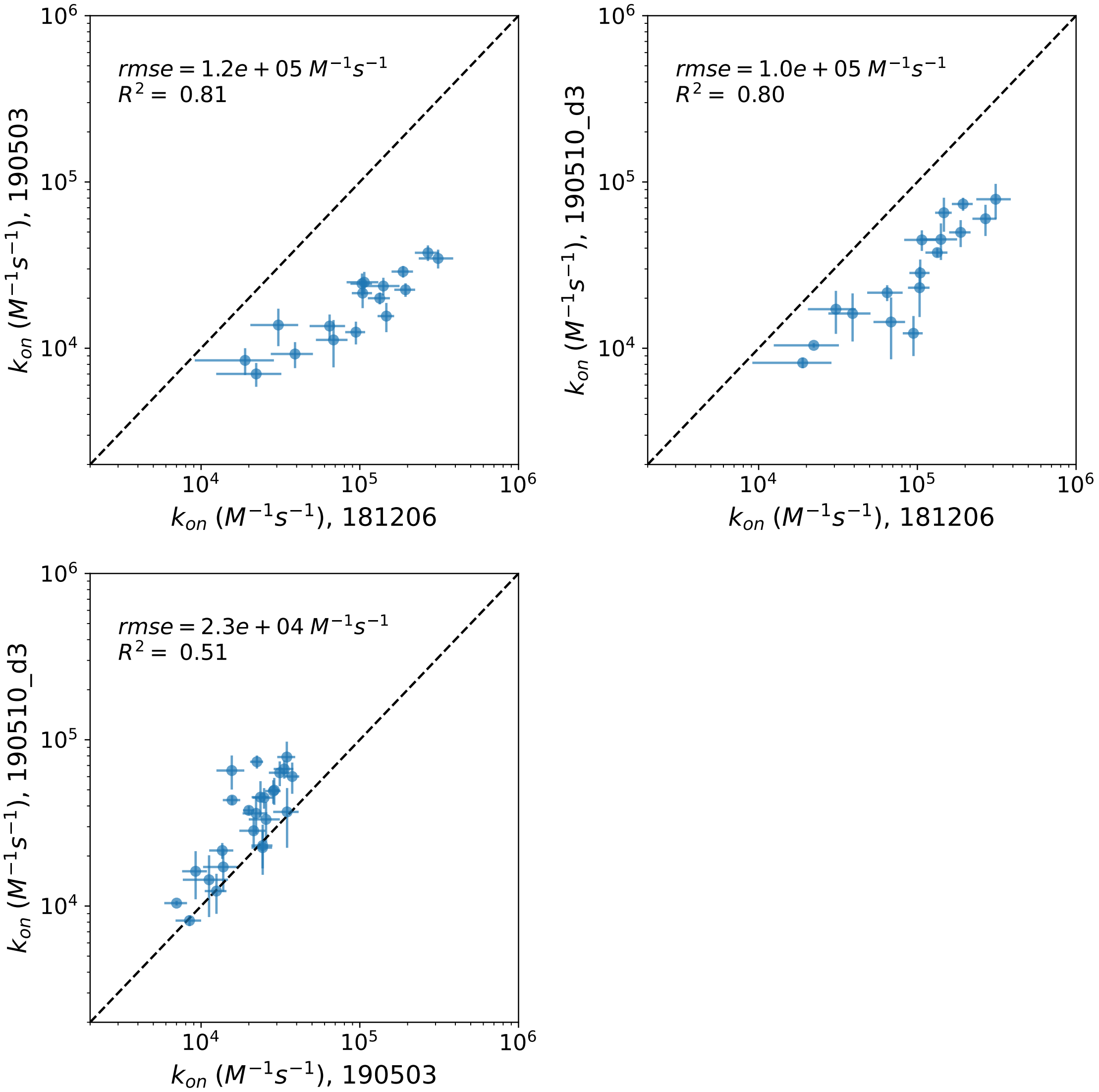

Figure 78: Pairwise comparisons of calculated $k_{on}$ values for MAX by sequence across 3 experiments quantifying MAX association rate to DNA library1. Markers indicate mean across all replicates within a given experiment; error bars indicate standard deviation of measured values.

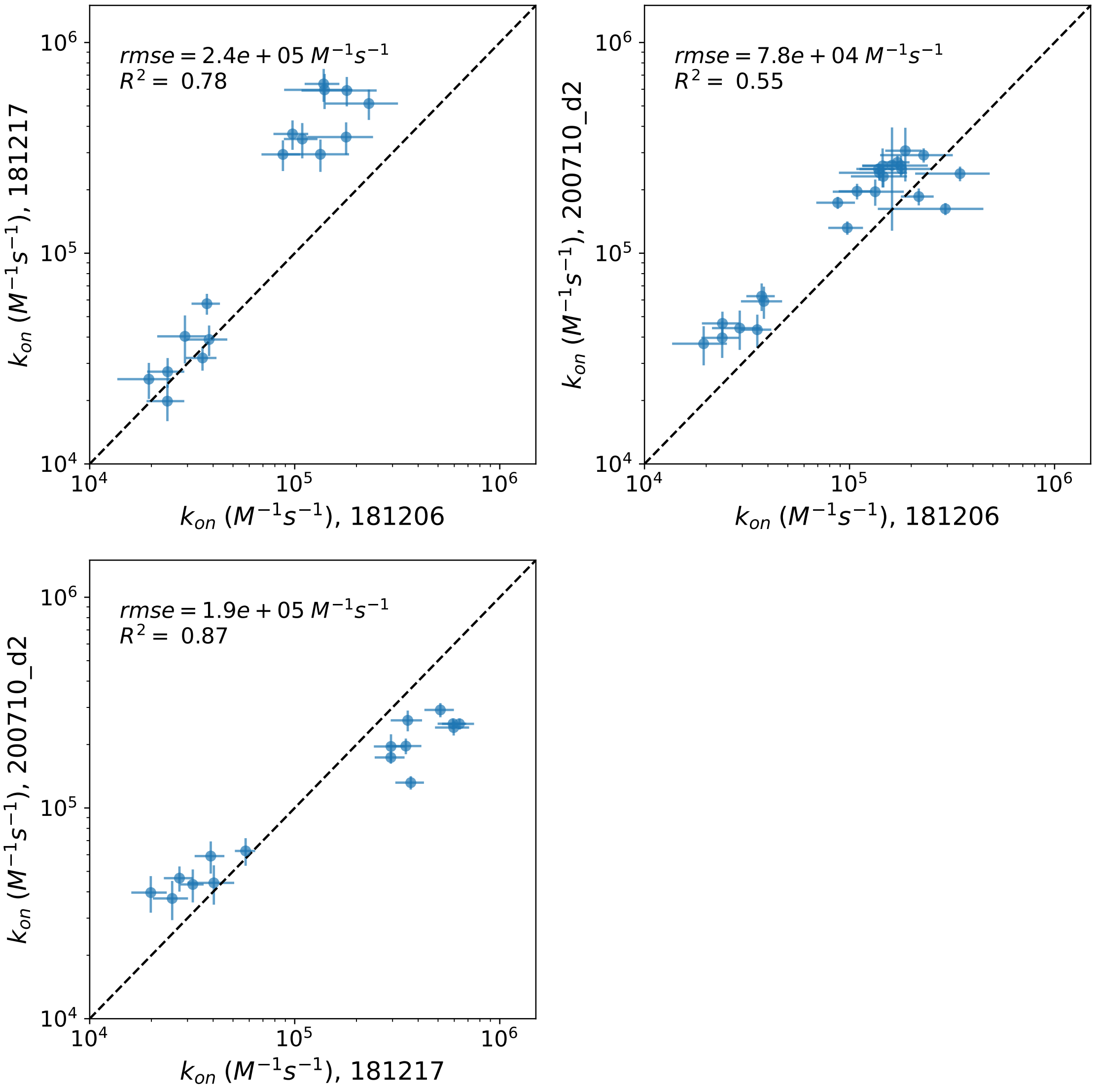

Figure 79: Pairwise comparisons of calculated $k_{on}$ values for Pho4 by sequence across 3 experiments quantifying Pho4 association rate to DNA Library 1. Markers indicate mean across all replicates within a given experiment; error bars indicate standard deviation of measured values.

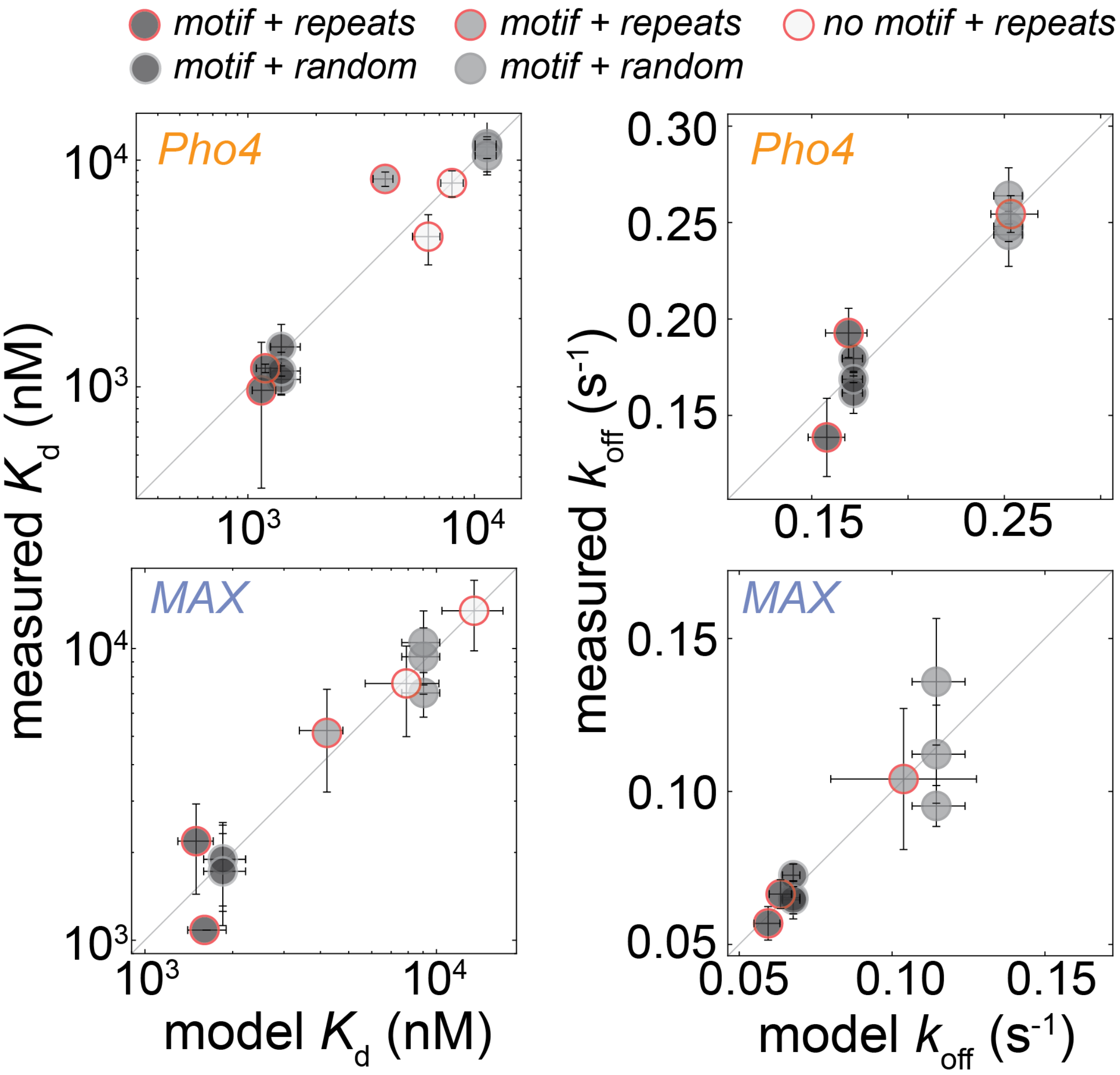

Figure 80: Measured vs. model-predicted affinities ($K_d$s, left) and dissociation rates ($k_{off}$s, right) across all sequences for Pho4 (top row) and MAX (bottom row)

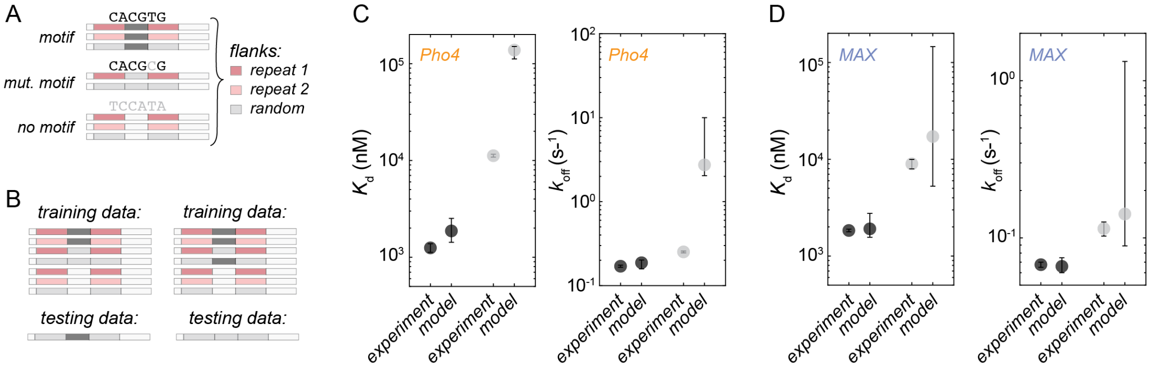

Figure 81: Continuous-time Markov Chain (CTMC) model fitting performance for train/test data sets. **(A)** Cartoon schematic of all sequence types with kinetic and equilibrium data used to train and test CTMC model. **(B)** Cartoon schematic of two train/test splits. Left: motif surrounded by random flank used for test (dark gray points in **(C)** and **(D)**). Right: mutated motif (CACGTG) surrounded by random flank used for test (light gray points in **(C)** and **(D)**). **(C)** Performance of model on test data for Pho4 $K_d$ and $k_{off}$. Error bars indicate standard deviation of bootstrapped distribution of fits. Differences between experimental and model values are likely due to known systematic over-estimation of $k_{off}$ and $K_d$ in MITOMI assays. **(D)** Performance of model on test data for MAX $K_d$ and $k_{off}$. Error bars indicate standard deviation of bootstrapped distribution of fits.

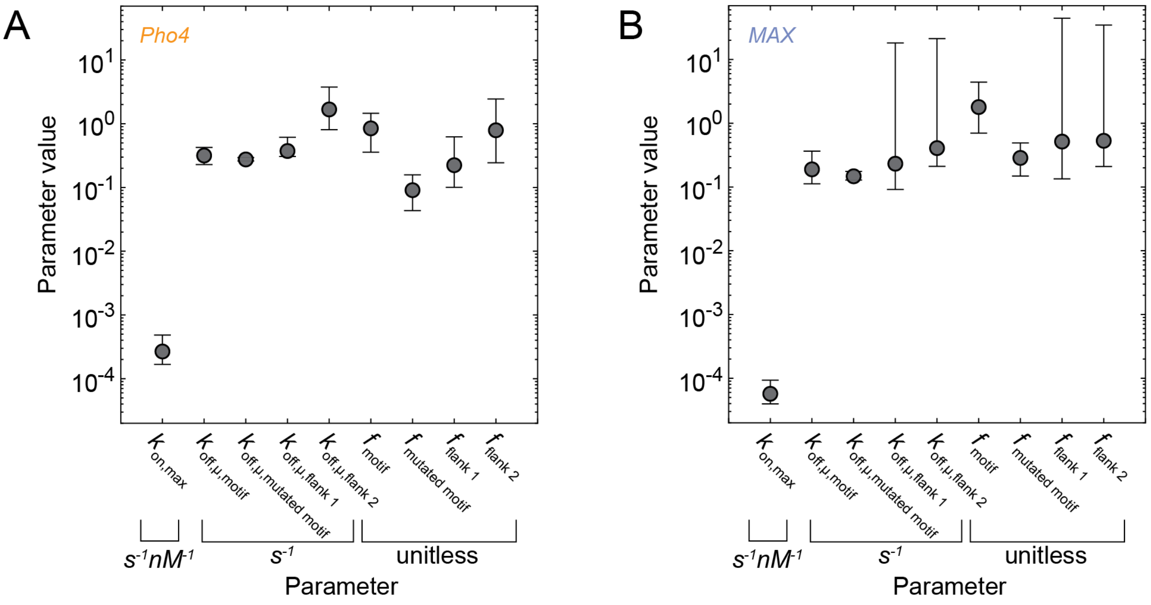

Figure 82: Parameters fit from continuous-time Markov Chain model for **(A)** Pho4 and **(B)** MAX.

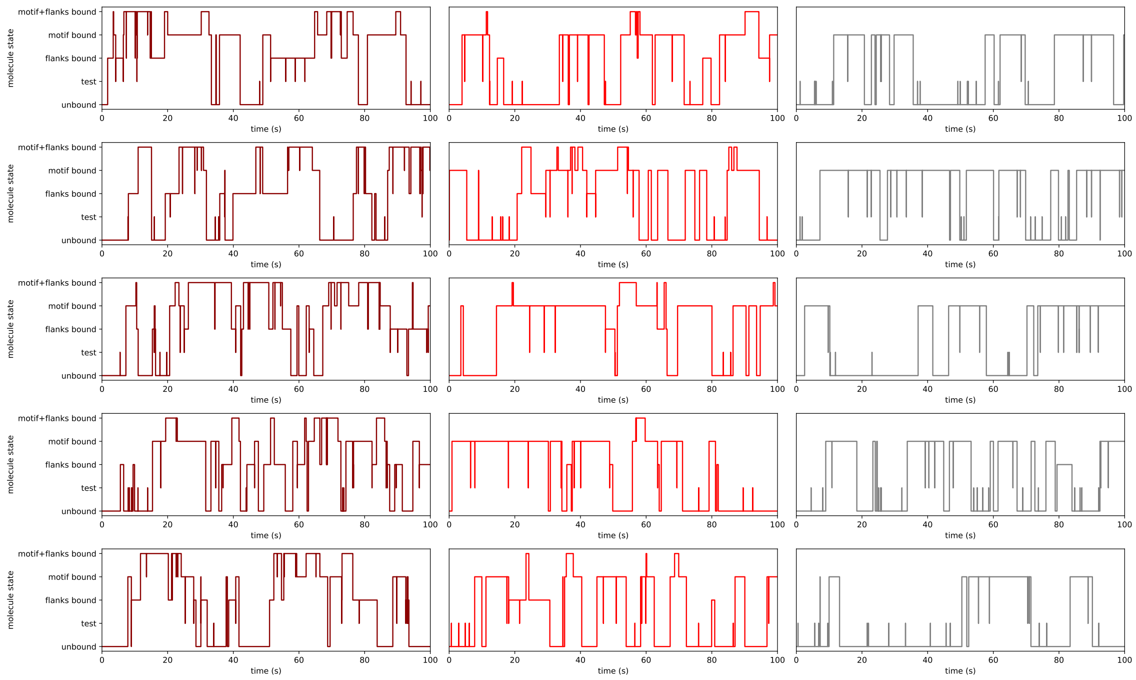

Figure 83: Example occupancy traces of a genomic binding site containing a motif and either CG repeats (maroon, left column), GT repeats (medium red, middle column), or random flanks (gray, right column) from individual iterations of a Gillespie simulation over 100 s. Each case models 2600 TF molecules. Note that flanking sequences can be occupied by up to 9 TFs at a time, while a motif can be occupied by only a single TF.

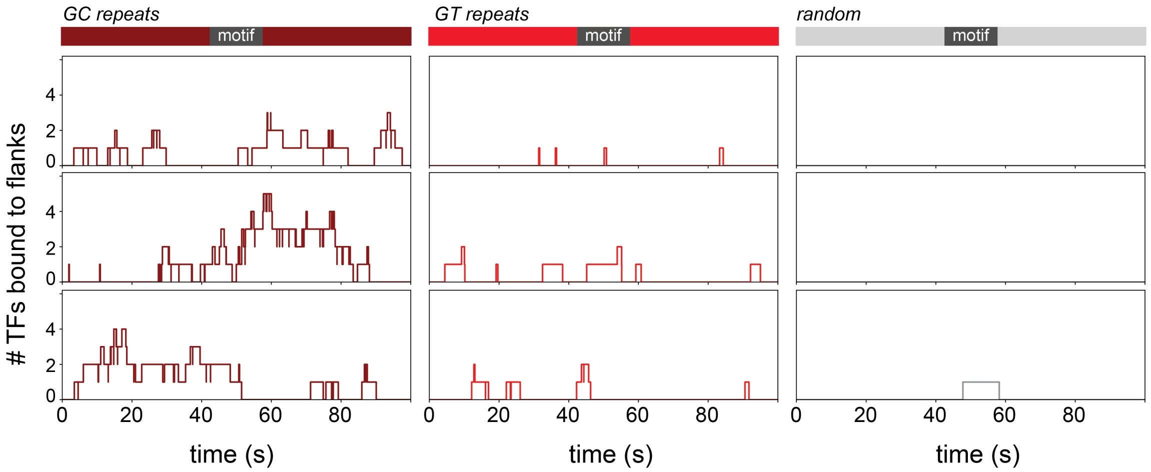

Figure 84: Example traces showing the number of TFs bound to flanking sequences surrounding a motif from individual iterations of a Gillespie simulation over 100 s. Flanking sequences are either CG repeats (maroon, left column), GT repeats (medium red, middle column), or random (gray, right column). Each case models 2600 TF molecules. Note that flanking sequences can be occupied by up to 9 TFs at a time, while a motif can be occupied by only a single TF.

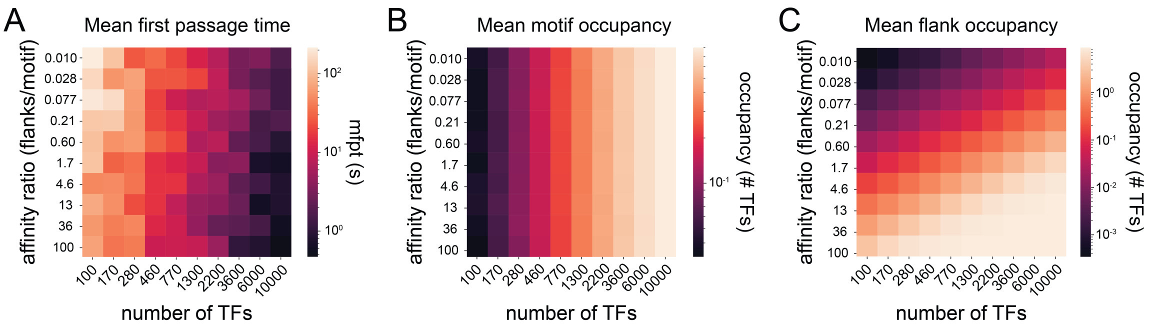

Figure 85: Sensitivity analysis for number of TFs acroos a range of flank/motif affinity ratios in Gillespie simulations. **(A)** Mean first passage time as a function of flank/motif affinity ratio and number of TFs. **(B)** Motif occupancy as a function of flank/motif affinity ratio and number of TFs. **(C)** Flank occupancy as a function of flank/motif affinity ratio and number of TFs. Each depicted value is averaged across 10 iterations of the simulation.

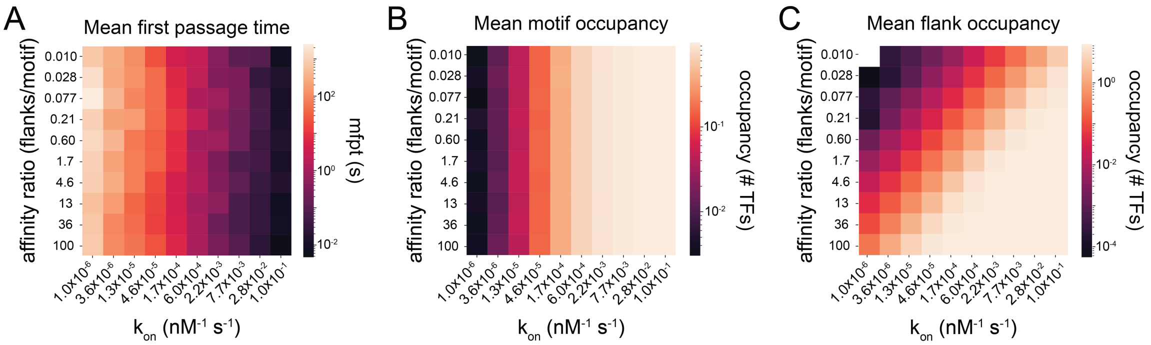

Figure 86: Sensitivity analysis for $k_{on}$ across a range of flank/motif affinity ratios in Gillespie simulations. **(A)** Mean first passage time as a function of flank/motif affinity ratio and $k_{on}$. **(B)** Motif occupancy as a function of flank/motif affinity ratio and $k_{on}$. **(C)** Flank occupancy as a function of flank/motif affinity ratio and $k_{on}$. Each depicted value is averaged across 10 iterations of the simulation.

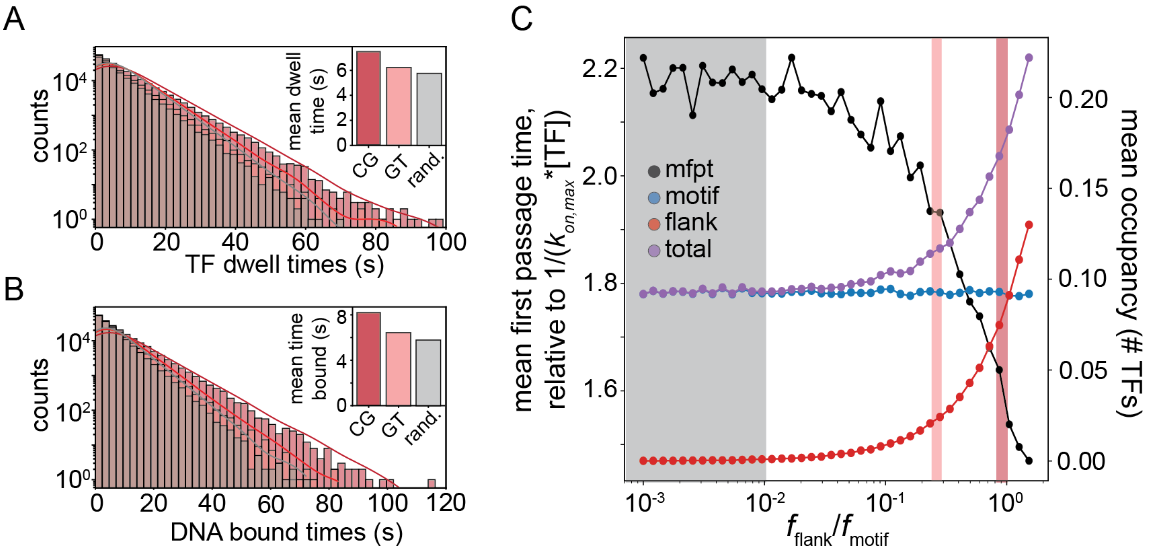

Figure 87: Supplement to Fig. 4G-I in the main text, where simulations were conducted with $k_{on,max} = 2.67 \times 10^5\ M^{-1}s^{-1}$. Here the same simulations are repeated with $k_{on,max} = 2.67 \times 10^4\ M^{-1}s^{-1}$. **(A)** Log-linear distribution of TF dwell times across 1000 simulations for sequences with a consensus motif flanked by CG repeats, GT repeats, or random sequence; inset shows mean dwell times by sequence. **(B)** Log-linear distribution of the fraction of time a DNA sequence is bound across 1000 simulations for sequences with a consensus motif flanked by GC repeats, GT repeats, or random sequence; inset shows mean time occupied by sequence. **(C)** Mean first passage time (black markers, left axis; units relative to fastest possible search time, 1/(kon,max*[TF])), mean motif occupancy (blue markers, right axis), mean flank occupancy (red markers, right axis), and mean total DNA occupancy (purple markers, right axis) as a function of the likelihood of binding flanking sequence; gray box indicates range of affinities for random flanks; pink and red boxes correspond to fflank values for GT and CG repeats, respectively.

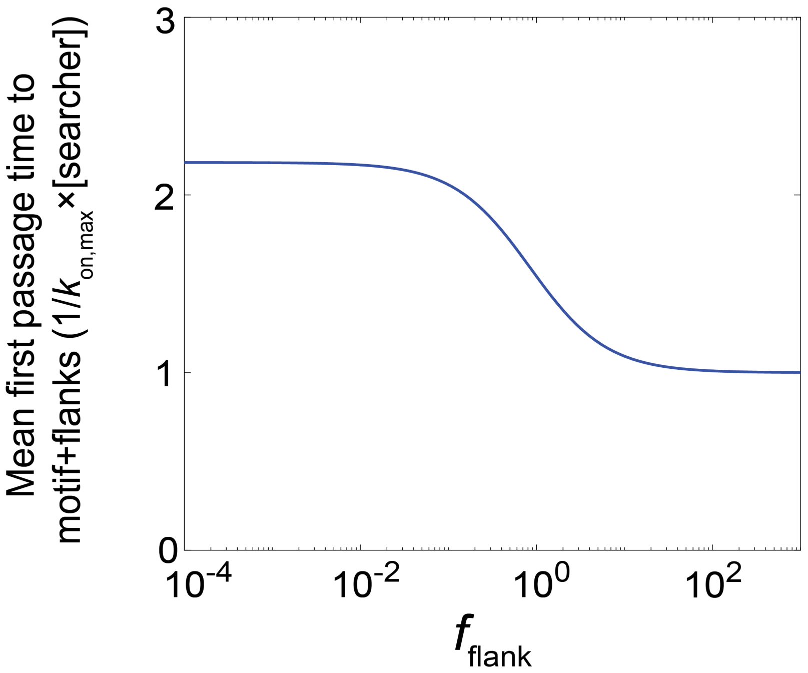

Figure 88: Mean first passage time computed from continuous-time Markov Chain model for Pho4 searching in the yeast nucleus as a function of $f_{flank}$, the likelihood of binding flanking sequence surrounding a motif.

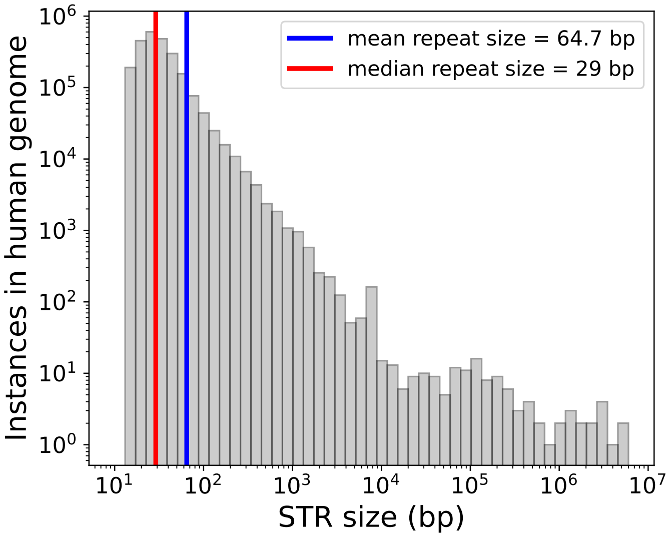

Figure 89: Distribution of all repeat sizes in the human genome with mean and median repeat sizes indicated.

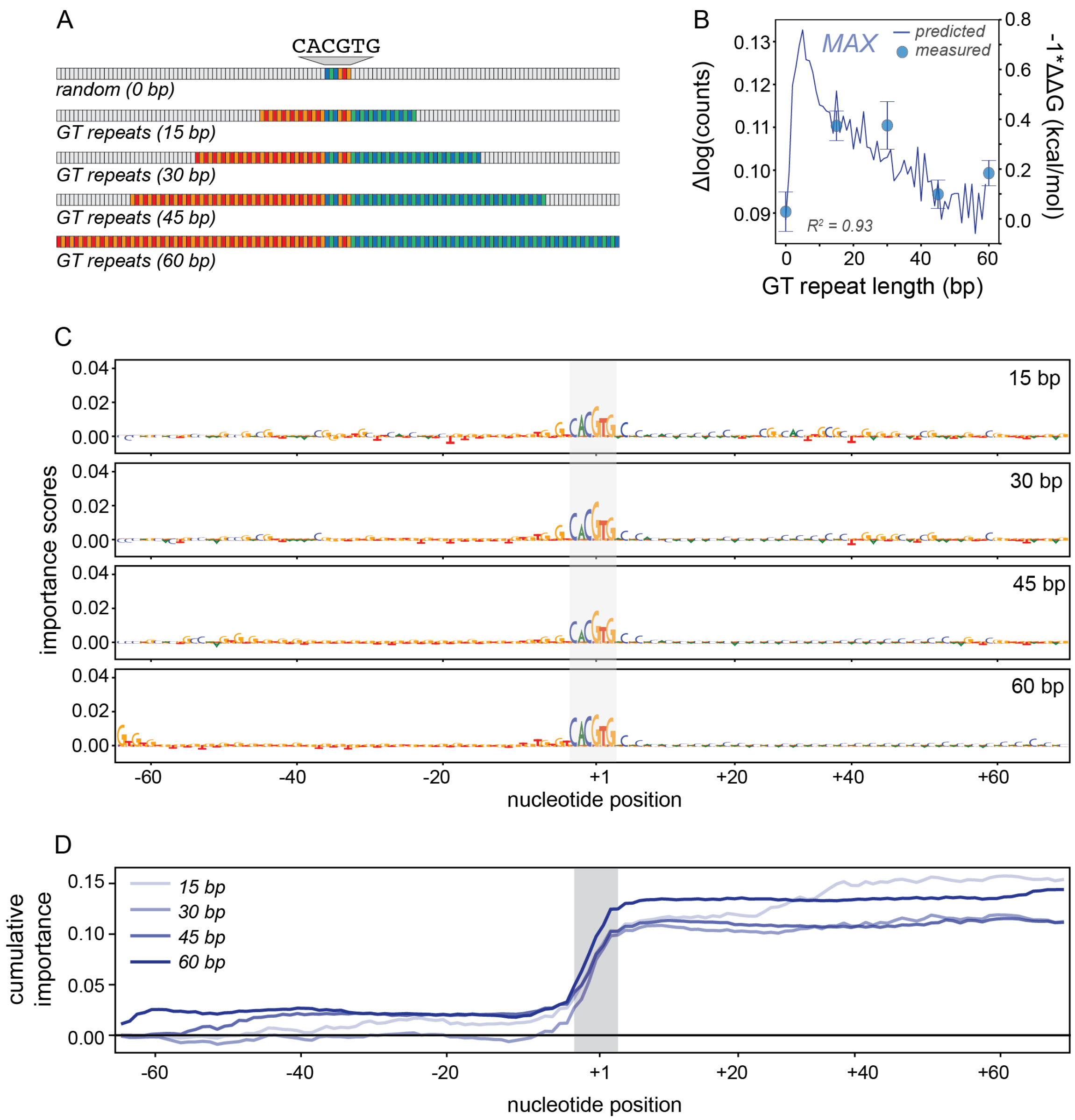

Figure 90: AffinityDistillation predictions for the effects of GT repeats on binding (trained on MAX ChIP-seq data) mirror *in vitro* measurements of MAX binding. **(A)** Schematic of sequences with E-box and 15, 30, 45, or 60 bp of favored GT repeats, measured in DNA Library 2. **(B)** AffinityDistillation-predicted change in log(counts) (blue line, left axis) and -1*MITOMI measured $\Delta\Delta G$s (blue markers, right axis) as a function of repeat length (relative to a sequence with a motif and random flanks). Markers and error bars show median and standard deviation across replicates. **(C)** DeepLIFT interpretations for sequences from **(A)** demonstrate non-uniform and subtle contributions of GT repeats to AffinityDistillation log($counts$) predictions. Gray box indicates motif position. **(D)** Cumulative importance scores as a function of position for sequences with E-box and 15, 30, 45, or 60 bp of GT repeats; gray box indicates motif position.

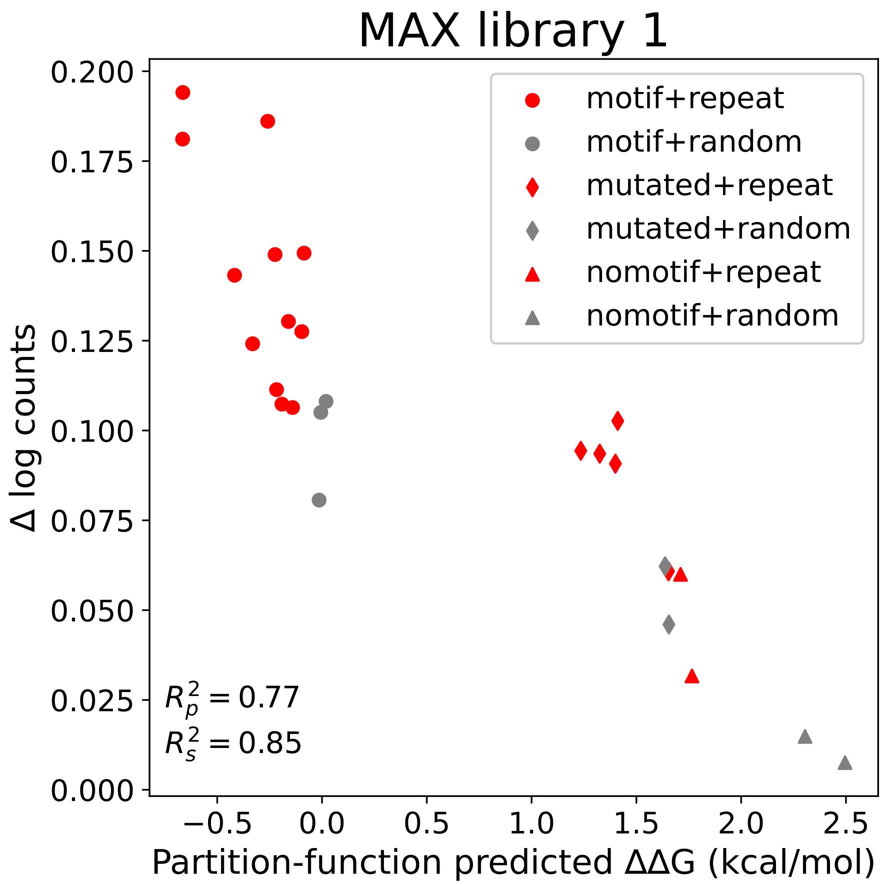

Figure 91: AffinityDistillation-predicted binding ($\Delta \log(counts)$) *vs.* partition-function predicted $\Delta\Delta G$s for 26 DNA sequences containing either an intact motif, a mutated motif, or scrambled sequence surrounded by either repetitive (red markers) or random (gray markers) flanking sequence.

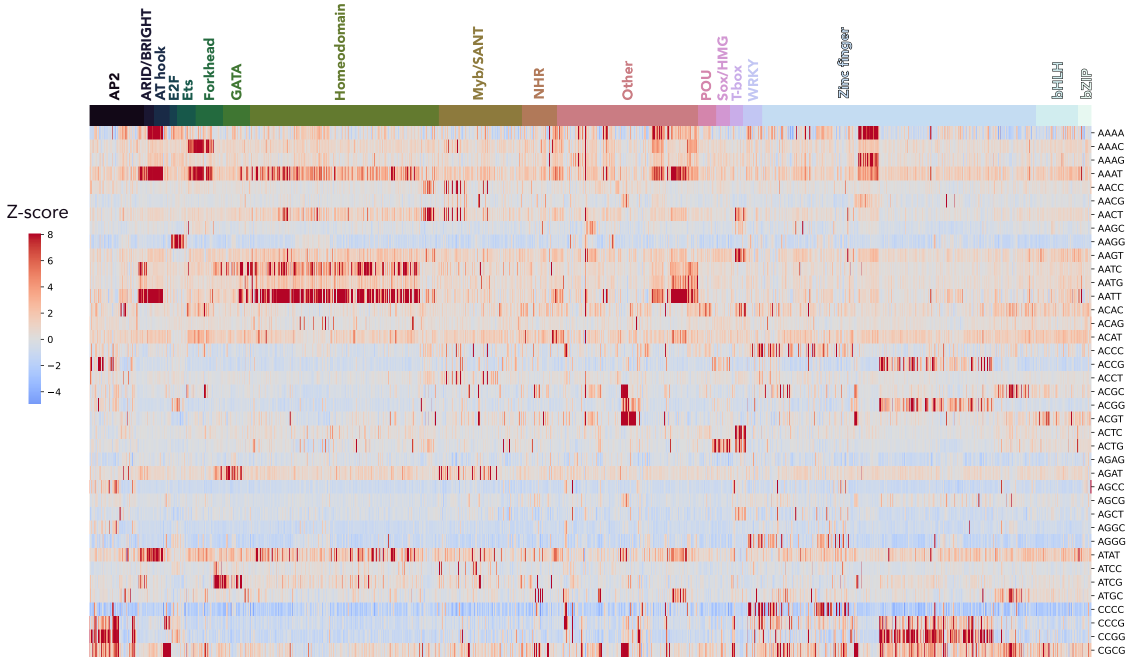

Figure 92: Heatmap showing the intensity Z-score for 1291 TFs interacting with 39 repeat types with TFs ordered by family.

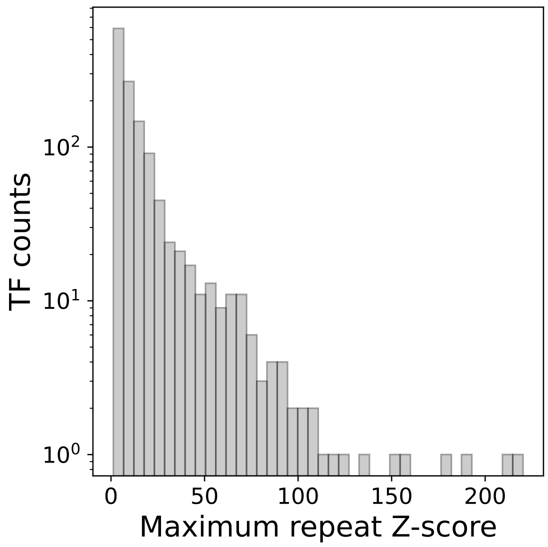

Figure 93: Distribution of the maximum repeat median intensity Z score across all 1291 TFs analyzed.

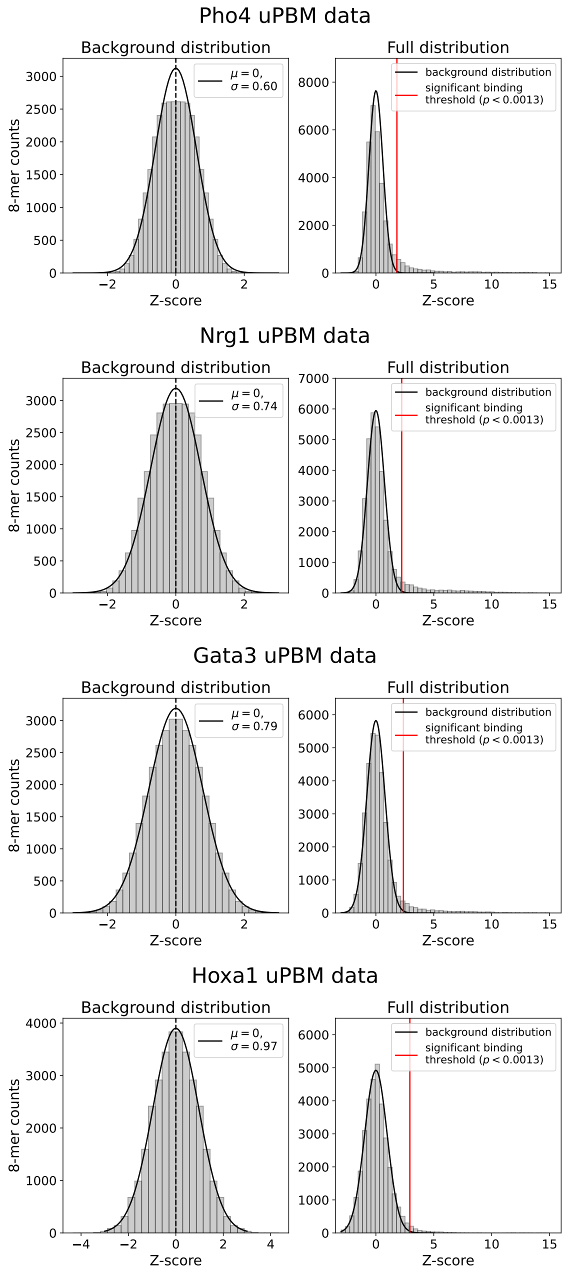

Figure 94: Median 8-mer intensity Z score distributions for uPBM experiments across 4 representative TFs (Pho4, Nrg1, Gata3, and Hoxa1). Background intensity Z-score distributions (left) were well-fit to a Gaussian centered around 0; 8-mer Z-scores with p-value less than Bonferroni-corrected threshold of $0.05/39 = 0.0013$ (indicated by the solid red light, right) were considered to be bound significantly above background. 39 is the number of repeat types and therefore the number of hypothesis tests performed.

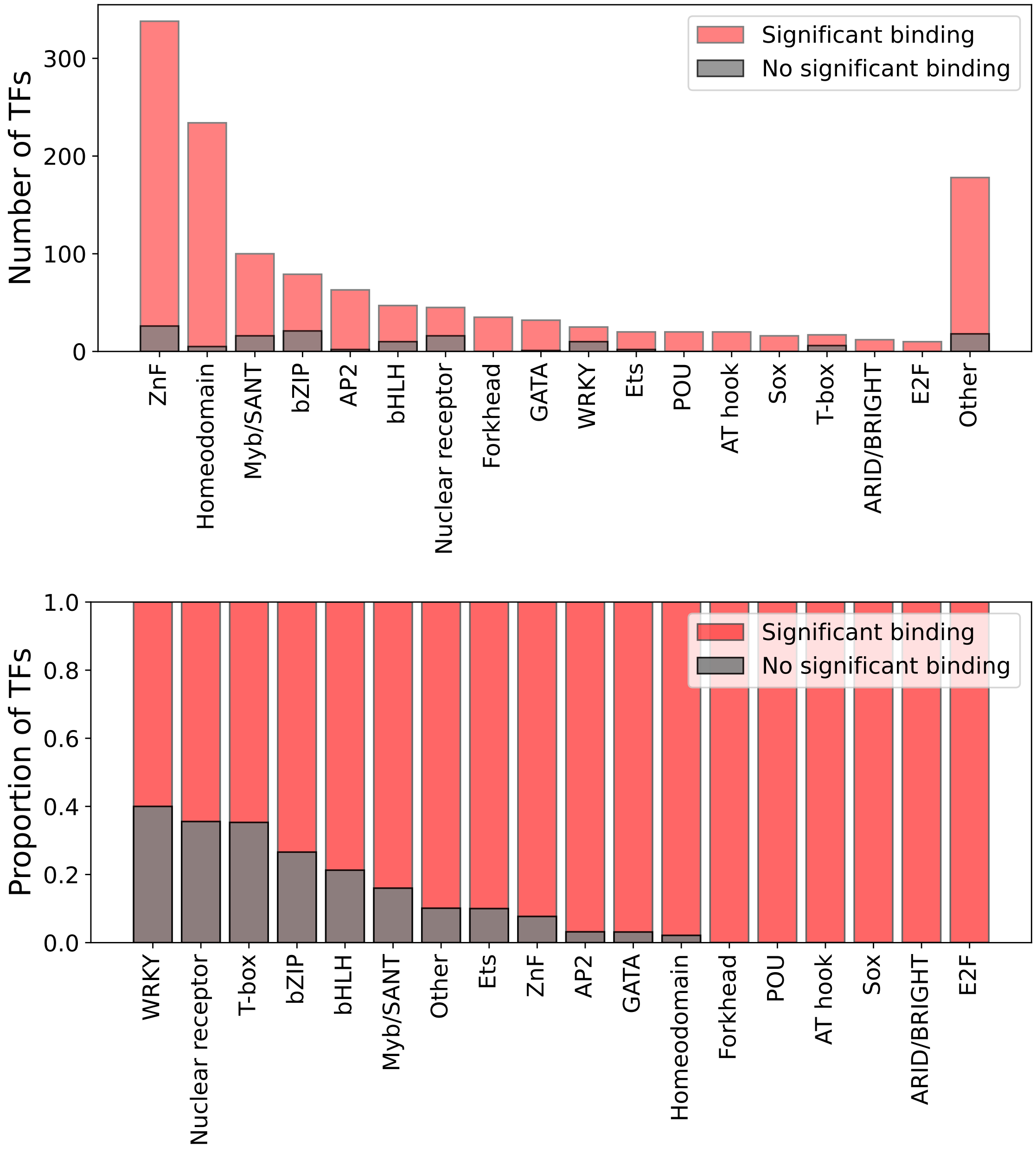

Figure 95: Number (top) and proportion (bottom) of TFs within TF families that show stronger binding to repeats than would be expected purely by chance.

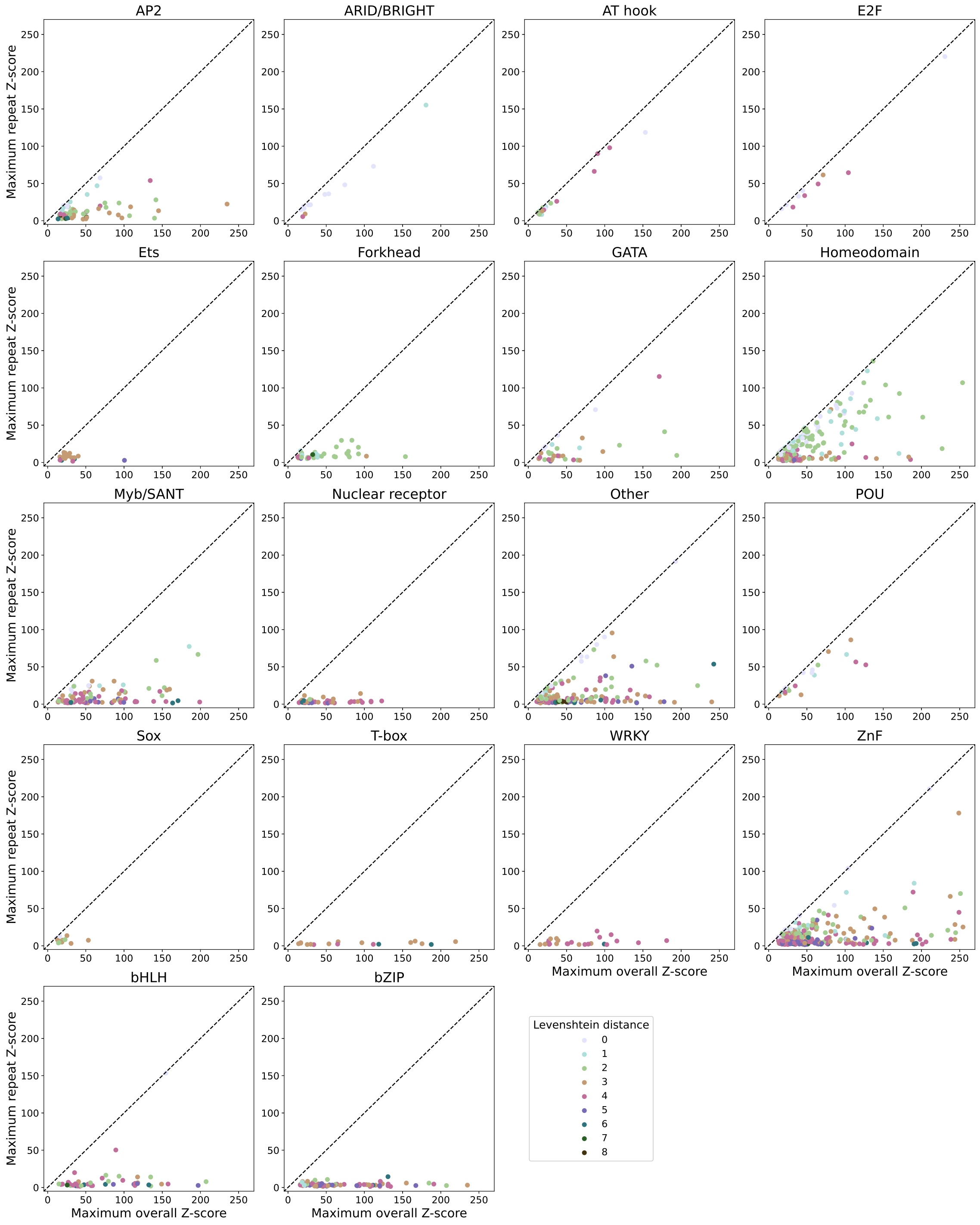

Figure 96: Distributions showing the maximum intensity Z score for a repetitive sequence *vs.* the maximum intensity Z score for all sequences for 18 different TF families; markers are colored according to the Levenshtein distance between the highest-scoring repeat and the highest-scoring sequence.

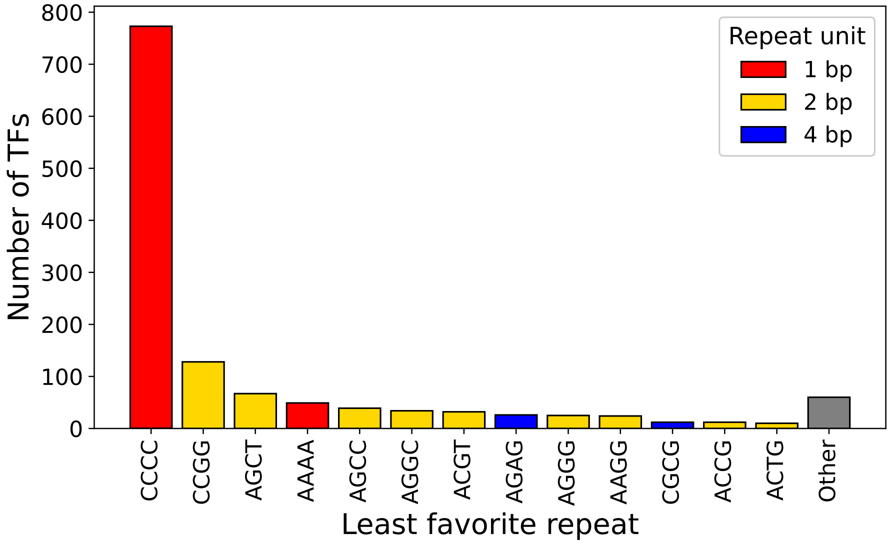

Figure 97: Repeats with lowest 8-mer Z-scores for each of 1,291 TFs. Mononucleotide repeats are shown in red, dinucleotide repeats are shown in yellow, and tetranucleotide repeats are shown in blue. All other repeats with fewer than 10 TFs are grouped into “Other.”

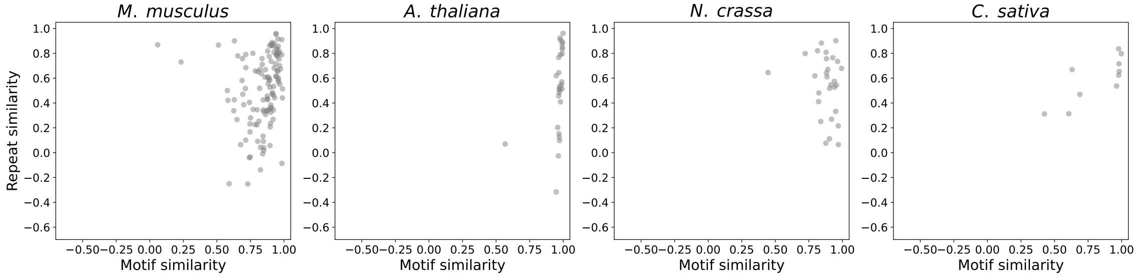

Figure 98: Pairwise comparison of motif and STR preferences, as measured by TF binding on universal PBMs, across basic helix-loop-helix (bHLH) paralogs within 4 different species. Here, we define paralogs as all TFs of the same structural class within a single species. Each point indicates the cosine similarity of motifs and of repeat preferences for a pairwise paralog comparison, with higher values indicating greater similarity. Each subplot depicts paralogs within a different species.

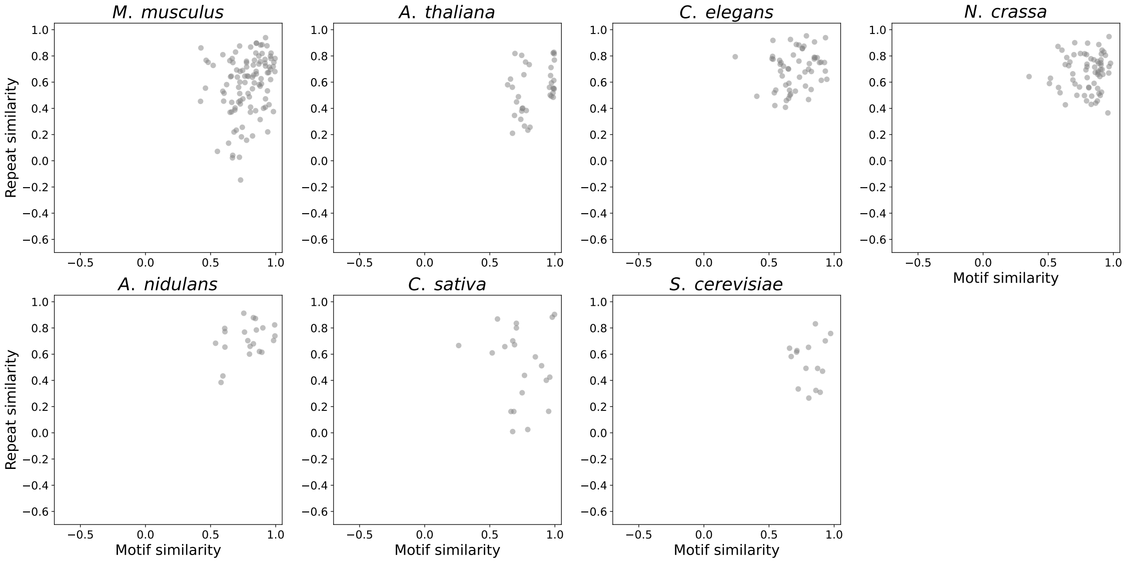

Figure 99: Pairwise comparison of motif and STR preferences, as measured by TF binding on universal PBMs, across basic leucine zipper (bZIP) paralogs within 7 different species. Here, we define paralogs as all TFs of the same structural class within a single species. Each point indicates the cosine similarity of motifs and of repeat preferences for a pairwise paralog comparison, with higher values indicating greater similarity. Each subplot depicts paralogs within a different species.

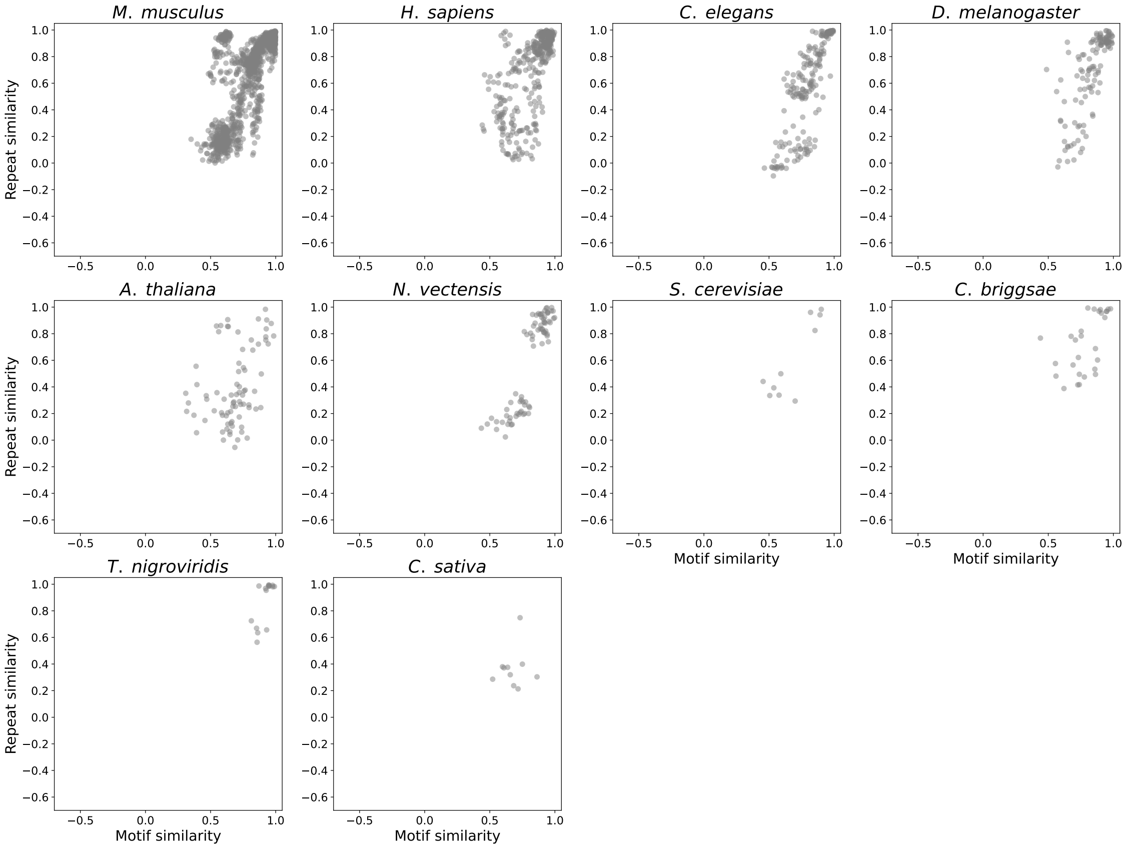

Figure 100: Pairwise comparison of motif and STR preferences, as measured by TF binding on universal PBMs, across homeodomain paralogs within 10 different species. Here, we define paralogs as all TFs of the same structural class within a single species. Each point indicates the cosine similarity of motifs and of repeat preferences for a pairwise paralog comparison, with higher values indicating greater similarity. Each subplot depicts paralogs within a different species.

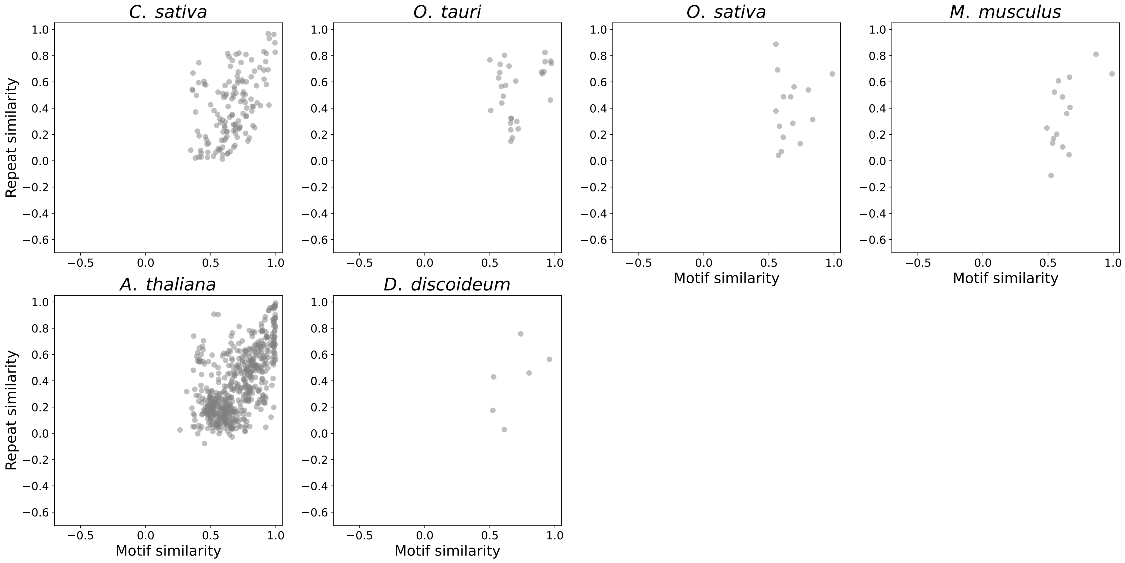

Figure 101: Pairwise comparison of motif and STR preferences, as measured by TF binding on universal PBMs, across Myb/SANT paralogs within 6 different species. Here, we define paralogs as all TFs of the same structural class within a single species. Each point indicates the cosine similarity of motifs and of repeat preferences for a pairwise paralog comparison, with higher values indicating greater similarity. Each subplot depicts paralogs within a different species.

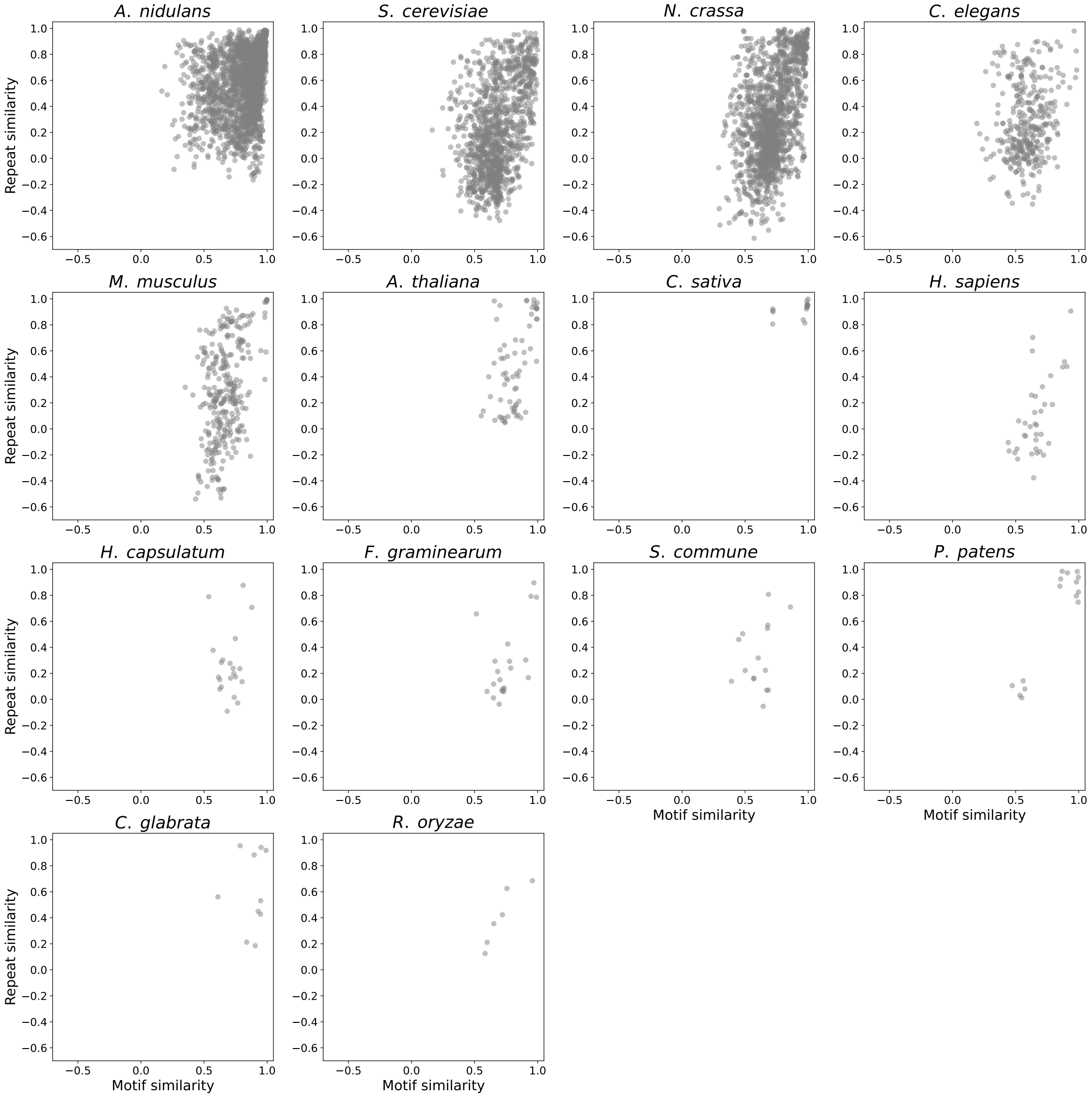

Figure 102: Pairwise comparison of motif and STR preferences, as measured by TF binding on universal PBMs, across zinc finger paralogs within 14 different species. Here, we define paralogs as all TFs of the same structural class within a single species. Each point indicates the cosine similarity of motifs and of repeat preferences for a pairwise paralog comparison, with higher values indicating greater similarity. Each subplot depicts paralogs within a different species.

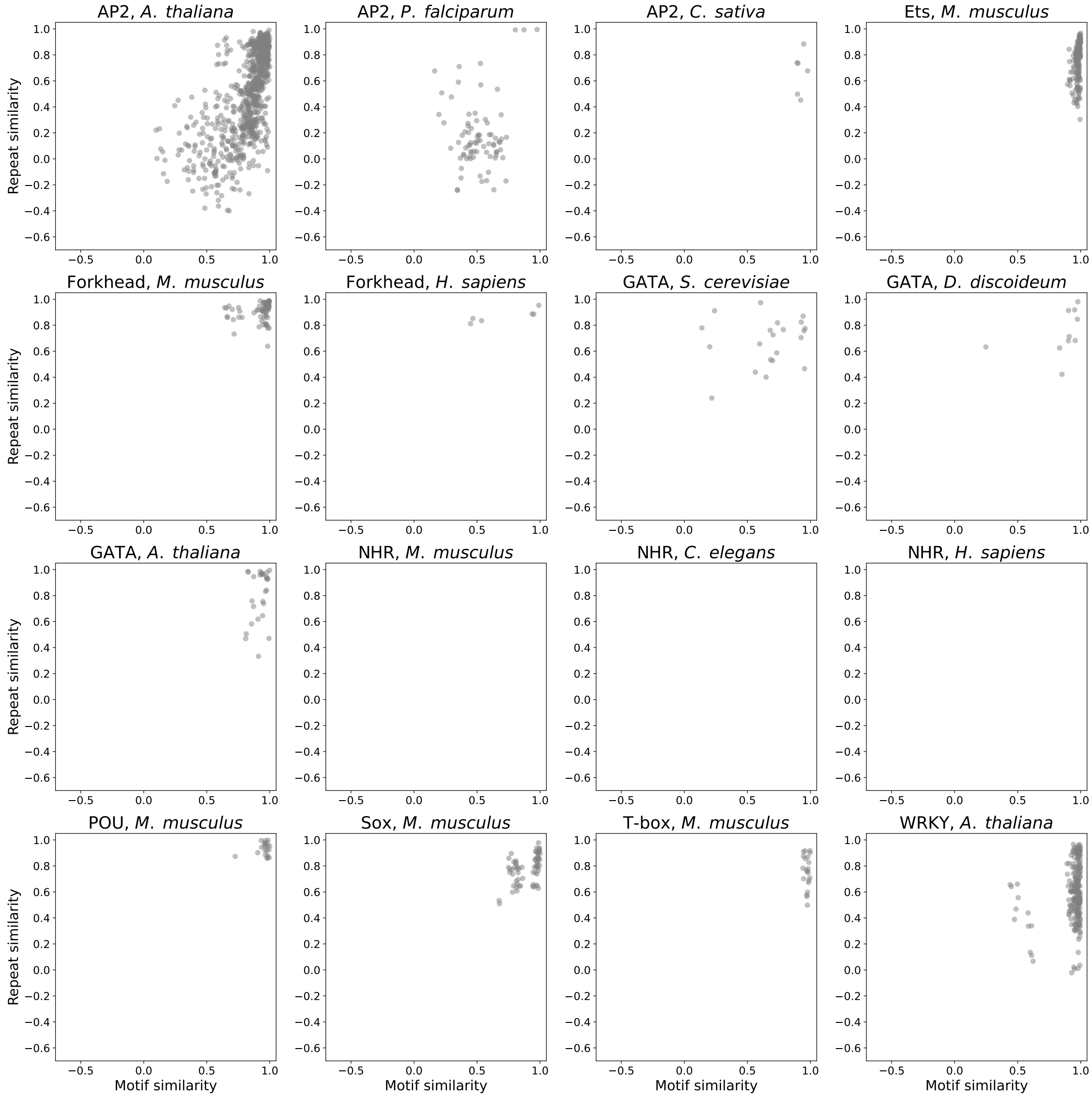

Figure 103: Pairwise comparison of motif and STR preferences, as measured by TF binding on universal PBMs, across various species and structural classes. Here, we define paralogs as all TFs of the same structural class within a single species. Each point indicates the cosine similarity of motifs and of repeat preferences for a pairwise paralog comparison, with higher values indicating greater similarity. Each subplot depicts paralogs of a specific structural class within a different species.

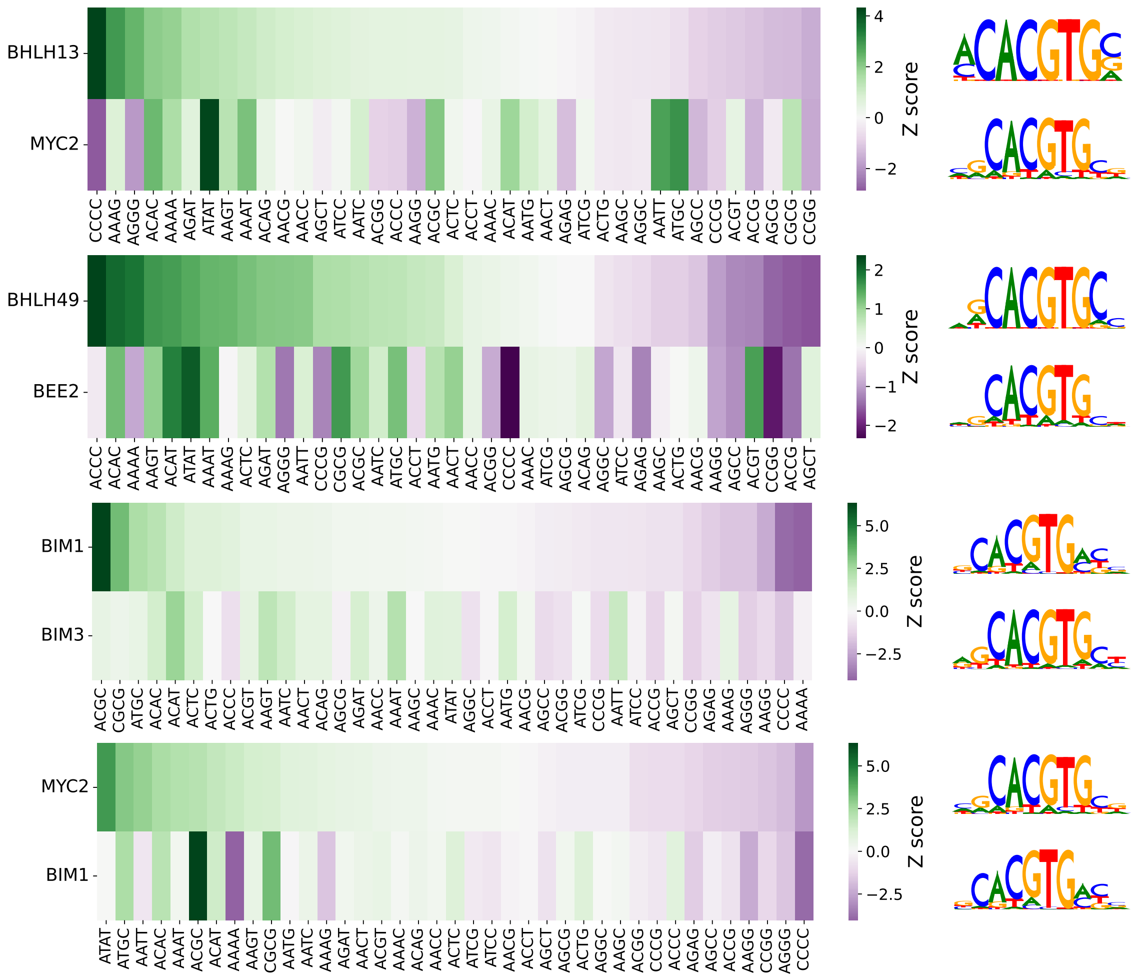

Figure 104: Pairwise comparison of STR preferences, as measured by TF binding on universal PBMs, across 4 pairs of basic helix-loop-helix (bHLH) paralogs in *Arabidopsis thaliana* with similar motif preferences. Heat maps show 8-mer Z-scores for 39 different repeat types. PWM logos show consensus motif preferences for each TF.

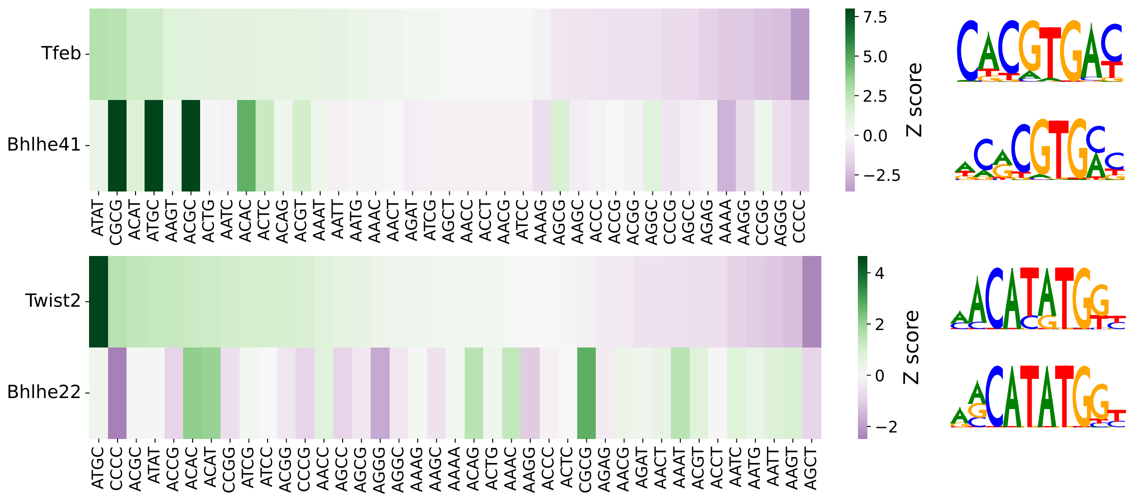

Figure 105: Pairwise comparison of STR preferences, as measured by TF binding on universal PBMs, across 2 pairs of basic helix-loop-helix (bHLH) paralogs in *Mus musculus* with similar motif preferences. Heat maps show 8-mer Z-scores for 39 different repeat types. PWM logos show consensus motif preferences for each TF.

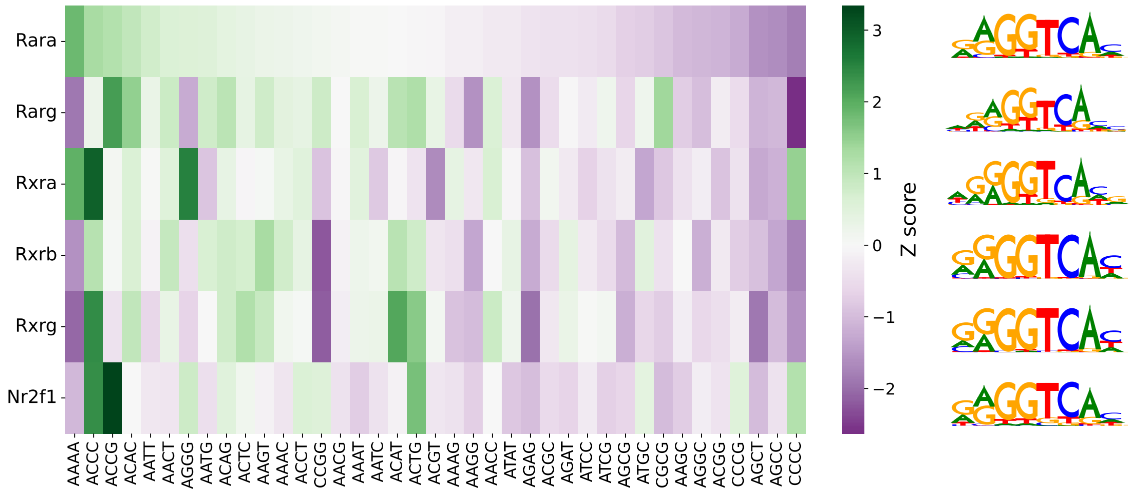

Figure 106: Comparison of STR preferences, as measured by TF binding on universal PBMs, across 6 pairs nuclear hormone receptor (NHR) paralogs in *Mus musculus* with similar motif preferences. Heat map shows 8-mer Z-scores for 39 different repeat types. PWM logos show consensus motif preferences for each TF.

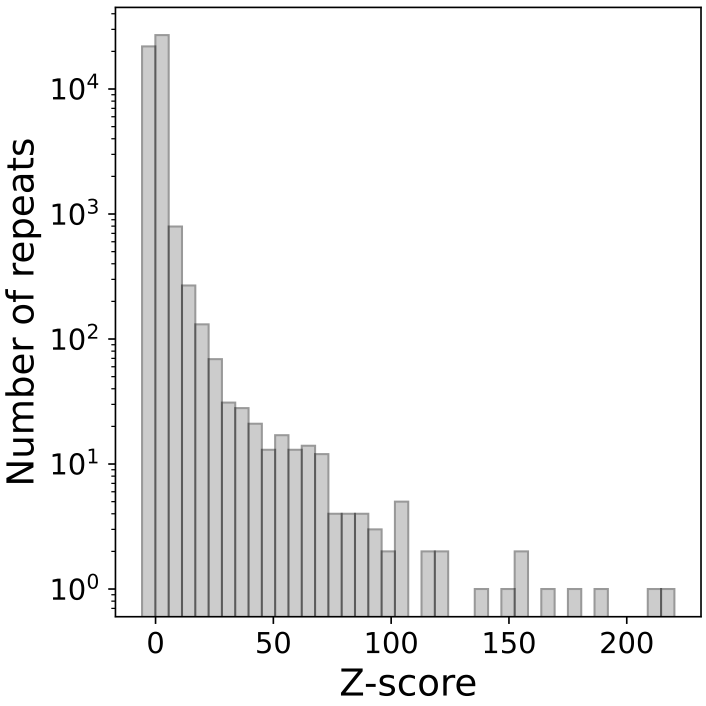

Figure 107: Distribution of all repeat Z-scores aggregated across 1,291 TFs.

### References

[1] Martín Abadi et al. *TensorFlow: Large-Scale Machine Learning on Heterogeneous Distributed Systems.* 2016. DOI: 10.48550/ARXIV.1603.04467. URL: https://arxiv.org/abs/1603.04467.

[2] Arjun K Aditham et al. “High-Throughput Affinity Measurements of Transcription Factor and DNA Mutations Reveal Affinity and Specificity Determinants.” In: *Cell Systems* 12.2 (Feb. 2021), 112–127.e11. DOI: 10.1016/j.cels.2020.11.012. URL: http://dx.doi.org/10.1016/j.cels.2020.11.012 (visited on 12/30/2020).

[3] Ariel Afek et al. “Protein-DNA binding in the absence of specific base-pair recognition.” In: *Proceedings of the National Academy of Sciences of the United States of America* 111.48 (Dec. 2014), pp. 17140–17145. DOI: 10.1073/pnas.1410569111. URL: http://dx.doi.org/10.1073/pnas.1410569111 (visited on 10/20/2018).

[4] Amr M. Alexandari et al. “De novo inference of thermodynamic binding energies using deep learning models of in vivo transcription factor binding”. In: *Manuscript in preparation* ().

[5] Nicolas Altemose. “A classical revival: Human satellite DNAs enter the genomics era”. In: *Seminars in Cell Developmental Biology* (2022). ISSN: 1084-9521. DOI: https://doi.org/10.1016/j.semcdb.2022.04.012. URL: https://www.sciencedirect.com/science/article/pii/S1084952122001379.

[6] Robin Andersson et al. “An atlas of active enhancers across human cell types and tissues.” In: *Nature* 507.7493 (Mar. 2014), pp. 455–461. DOI: 10.1038/nature12787. URL: http://dx.doi.org/10.1038/nature12787 (visited on 03/16/2020).

[7] Žiga Avsec et al. “Base-resolution models of transcription-factor binding reveal soft motif syntax.” In: *Nature Genetics* 53.3 (Mar. 2021), pp. 354–366. DOI: 10.1038/s41588-021-00782-6. URL: http://dx.doi.org/10.1038/s41588-021-00782-6 (visited on 11/19/2021).

[8] Gwenael Badis et al. “Diversity and complexity in DNA recognition by transcription factors.” In: *Science* 324.5935 (June 2009), pp. 1720–1723. ISSN: 1095-9203. DOI: 10.1126/science.1162327. URL: http://dx.doi.org/10.1126/science.1162327 (visited on 12/30/2018).

[9] Luis A Barrera et al. “Survey of variation in human transcription factors reveals prevalent DNA binding changes.” In: *Science* 351.6280 (Mar. 2016), pp. 1450–1454. DOI: 10.1126/science.aad2257. URL: http://dx.doi.org/10.1126/science.aad2257 (visited on 12/28/2021).

[10] G Benson. “Tandem repeats finder: a program to analyze DNA sequences.” In: *Nucleic Acids Research* 27.2 (Jan. 1999), pp. 573–580. DOI: 10.1093/nar/27.2.573. URL: http://dx.doi.org/10.1093/nar/27.2.573 (visited on 10/29/2019).

[11] Michael F Berger et al. “Compact, universal DNA microarrays to comprehensively determine transcription-factor binding site specificities.” In: *Nature Biotechnology* 24.11 (Nov. 2006), pp. 1429–1435. DOI: 10.1038/nbt1246. URL: http://dx.doi.org/10.1038/nbt1246 (visited on 04/17/2019).

[12] Kara Brower et al. “An Open-Source, Programmable Pneumatic Setup for Operation and Automated Control of Single- and Multi-Layer Microfluidic Devices.” In: *HardwareX* 3 (Apr. 2018), pp. 117–134. ISSN: 24680672. DOI: 10.1016/j.ohx.2017.10.001. URL: https://linkinghub.elsevier.com/retrieve/pii/S2468067217300433 (visited on 07/11/2021).

[13] Tsu-Pei Chiu et al. "DNAshapeR: an R/Bioconductor package for DNA shape prediction and feature encoding". In: *Bioinformatics* 32.8 (Dec. 2015), pp. 1211–1213. ISSN: 1367-4803. DOI: 10.1093/bioinformatics/btv735. eprint: https://academic.oup.com/bioinformatics/article-pdf/32/8/1211/16921293/btv735.pdf. URL: https://doi.org/10.1093/bioinformatics/btv735.

[14] François Chollet et al. *Keras*. https://keras.io. 2015.

[15] ENCODE Project Consortium. "An integrated encyclopedia of DNA elements in the human genome." In: *Nature* 489.7414 (Sept. 2012), pp. 57–74. DOI: 10.1038/nature11247. URL: http://dx.doi.org/10.1038/nature11247 (visited on 07/14/2016).

[16] Carrie A Davis et al. "The Encyclopedia of DNA elements (ENCODE): data portal update." In: *Nucleic Acids Research* 46.D1 (Jan. 2018), pp. D794–D801. DOI: 10.1093/nar/gkx1081. URL: http://dx.doi.org/10.1093/nar/gkx1081 (visited on 02/16/2021).

[17] Arthur D Edelstein et al. "Advanced methods of microscope control using $\mu$Manager software." In: *Journal of biological methods* 1.2 (2014). DOI: 10.14440/jbm.2014.36. URL: http://dx.doi.org/10.14440/jbm.2014.36 (visited on 07/11/2021).

[18] Polly M Fordyce et al. "De novo identification and biophysical characterization of transcription-factor binding sites with microfluidic affinity analysis." In: *Nature Biotechnology* 28.9 (Sept. 2010), pp. 970–975. DOI: 10.1038/nbt.1675. URL: http://dx.doi.org/10.1038/nbt.1675 (visited on 10/20/2018).

[19] Tianshun Gao and Jiang Qian. "EnhancerAtlas 2.0: an updated resource with enhancer annotation in 586 tissue/cell types across nine species." In: *Nucleic Acids Research* 48.D1 (Jan. 2020), pp. D58–D64. DOI: 10.1093/nar/gkz980. URL: http://dx.doi.org/10.1093/nar/gkz980 (visited on 12/20/2020).

[20] Marcel Geertz, David Shore, and Sebastian J Maerkl. "Massively parallel measurements of molecular interaction kinetics on a microfluidic platform." In: *Proceedings of the National Academy of Sciences of the United States of America* 109.41 (Oct. 2012), pp. 16540–16545. DOI: 10.1073/pnas.1206011109. URL: http://dx.doi.org/10.1073/pnas.1206011109 (visited on 10/25/2018).

[21] Daniel T Gillespie. "A general method for numerically simulating the stochastic time evolution of coupled chemical reactions". In: *Journal of Computational Physics* 22.4 (Dec. 1976), pp. 403–434. ISSN: 00219991. DOI: 10.1016/0021-9991(76)90041-3. URL: http://linkinghub.elsevier.com/retrieve/pii/0021999176900413 (visited on 01/11/2022).

[22] A S Hinrichs et al. "The UCSC Genome Browser Database: update 2006." In: *Nucleic Acids Research* 34.Database issue (Jan. 2006), pp. D590–8. DOI: 10.1093/nar/gkj144. URL: http://dx.doi.org/10.1093/nar/gkj144 (visited on 06/16/2020).

[23] Brandon Ho, Anastasia Baryshnikova, and Grant W Brown. "Unification of Protein Abundance Datasets Yields a Quantitative Saccharomyces cerevisiae Proteome." In: *Cell Systems* 6.2 (Feb. 2018), 192–205.e3. ISSN: 24054712. DOI: 10.1016/j.cels.2017.12.004. URL: http://linkinghub.elsevier.com/retrieve/pii/%7BS240547121730546X%7D (visited on 03/11/2021).

[24] Inga Jarmoskaite et al. "How to measure and evaluate binding affinities." In: *eLife* 9 (Aug. 2020). ISSN: 2050-084X. DOI: 10.7554/{eLife}.57264. URL: https://elifesciences.org/articles/57264 (visited on 08/12/2020).

[25] Brian H. Johnston. “[7] Generation and detection of Z-DNA”. In: *DNA structures part A: synthesis and physical analysis of DNA*. Vol. 211. Methods in Enzymology. Elsevier, 1992, pp. 127–158. ISBN: 9780121821128. DOI: 10.1016/0076-6879(92)11009-8. URL: https://linkinghub.elsevier.com/retrieve/pii/0076687992110098 (visited on 09/30/2019).

[26] Diederik P. Kingma and Jimmy Ba. *Adam: A Method for Stochastic Optimization.* 2014. DOI: 10.48550/ARXIV.1412.6980. URL: https://arxiv.org/abs/1412.6980.

[27] Daniel D Le et al. “Comprehensive, high-resolution binding energy landscapes reveal context dependencies of transcription factor binding.” In: *Proceedings of the National Academy of Sciences of the United States of America* 115.16 (Apr. 2018), E3702–E3711. ISSN: 0027-8424. DOI: 10.1073/pnas.1715888115. URL: http://www.pnas.org/lookup/doi/10.1073/pnas.1715888115 (visited on 10/20/2018).

[28] Marko Lõoke, Kersti Kristjuhan, and Arnold Kristjuhan. “Extraction of genomic DNA from yeasts for PCR-based applications.” In: *Biotechniques* 50.5 (May 2011), pp. 325–328. DOI: 10.2144/000113672. URL: http://dx.doi.org/10.2144/000113672 (visited on 10/08/2019).

[29] Scott Lundberg and Su-In Lee. *A Unified Approach to Interpreting Model Predictions.* 2017. DOI: 10.48550/ARXIV.1705.07874. URL: https://arxiv.org/abs/1705.07874.

[30] Michael Lynch et al. “A genome-wide view of the spectrum of spontaneous mutations in yeast.” In: *Proceedings of the National Academy of Sciences of the United States of America* 105.27 (July 2008), pp. 9272–9277. ISSN: 1091-6490. DOI: 10.1073/pnas.0803466105. URL: http://dx.doi.org/10.1073/pnas.0803466105 (visited on 03/10/2017).

[31] Sebastian J Maerkl and Stephen R Quake. “A systems approach to measuring the binding energy landscapes of transcription factors.” In: *Science* 315.5809 (Jan. 2007), pp. 233–237. DOI: 10.1126/science.1131007. URL: http://dx.doi.org/10.1126/science.1131007 (visited on 11/28/2018).

[32] C J Markin et al. “Revealing enzyme functional architecture via high-throughput microfluidic enzyme kinetics.” In: *Science* 373.6553 (July 2021). ISSN: 0036-8075. DOI: 10.1126/science.abf8761. URL: https://www.sciencemag.org/lookup/doi/10.1126/science.abf8761 (visited on 05/04/2022).

[33] Emil Marklund et al. “Sequence specificity in DNA binding is mainly governed by association”. In: *Science* 375.6579 (Jan. 2022), pp. 442–445. ISSN: 0036-8075. DOI: 10.1126/science.abg7427. URL: https://www.science.org/doi/10.1126/science.abg7427 (visited on 02/06/2022).

[34] T N Marriage et al. “Direct estimation of the mutation rate at dinucleotide microsatellite loci in Arabidopsis thaliana (Brassicaceae).” In: *Heredity* 103.4 (Oct. 2009), pp. 310–317. DOI: 10.1038/hdy.2009.67. URL: http://dx.doi.org/10.1038/hdy.2009.67 (visited on 01/23/2022).

[35] G G Maul and L Deaven. “Quantitative determination of nuclear pore complexes in cycling cells with differing DNA content.” In: *The Journal of Cell Biology* 73.3 (June 1977), pp. 748–760. DOI: 10.1083/jcb.73.3.748. URL: http://dx.doi.org/10.1083/jcb.73.3.748 (visited on 01/11/2022).

[36] Ron Milo and Rob Phillips. *Cell biology by the numbers.* Garland Science, Dec. 2015. ISBN: 9780429258770. DOI: 10.1201/9780429258770. URL: https://www.taylorfrancis.com/books/9781317230694 (visited on 05/04/2022).

[37] Michael Model. “Intensity calibration and flat-field correction for fluorescence microscopes.” In: *Current protocols in cytometry / editorial board, J. Paul Robinson, managing editor ... [et al.]* 68 (Apr. 2014), pp. 10.14.1–10. DOI: 10.1002/0471142956.cy1014s68. URL: http://dx.doi.org/10.1002/0471142956.cy1014s68 (visited on 07/11/2021).

[38] Sergey Nurk et al. “The complete sequence of a human genome”. In: *Science* 376.6588 (2022), pp. 44–53. DOI: 10.1126/science.abj6987. eprint: https://www.science.org/doi/pdf/10.1126/science.abj6987. URL: https://www.science.org/doi/abs/10.1126/science.abj6987.

[39] Stephan Ossowski et al. “The rate and molecular spectrum of spontaneous mutations in Arabidopsis thaliana.” In: *Science* 327.5961 (Jan. 2010), pp. 92–94. ISSN: 1095-9203. DOI: 10.1126/science.1180677. URL: http://dx.doi.org/10.1126/science.1180677 (visited on 01/23/2022).

[40] Johannes Schindelin et al. “Fiji: an open-source platform for biological-image analysis.” In: *Nature Methods* 9.7 (June 2012), pp. 676–682. DOI: 10.1038/nmeth.2019. URL: http://dx.doi.org/10.1038/nmeth.2019 (visited on 07/11/2021).

[41] Avanti Shrikumar, Peyton Greenside, and Anshul Kundaje. “Learning Important Features Through Propagating Activation Differences”. In: (2017). DOI: 10.48550/ARXIV.1704.02685. URL: https://arxiv.org/abs/1704.02685.

[42] Avanti Shrikumar, Peyton Greenside, and Anshul Kundaje. “Reverse-complement parameter sharing improves deep learning models for genomics”. In: *bioRxiv* (2017). DOI: 10.1101/103663. eprint: https://www.biorxiv.org/content/early/2017/01/27/103663.full.pdf. URL: https://www.biorxiv.org/content/early/2017/01/27/103663.

[43] T J Thomas, U B Gunnia, and T Thomas. “Polyamine-induced B-DNA to Z-DNA conformational transition of a plasmid DNA with (dG-dC)n insert.” In: *The Journal of Biological Chemistry* 266.10 (Apr. 1991), pp. 6137–6141. URL: https://www.ncbi.nlm.nih.gov/pubmed/1848849 (visited on 09/27/2019).

[44] Jing Wang et al. “HACER: an atlas of human active enhancers to interpret regulatory variants.” In: *Nucleic Acids Research* 47.D1 (Jan. 2019), pp. D106–D112. DOI: 10.1093/nar/gky864. URL: http://dx.doi.org/10.1093/nar/gky864 (visited on 12/20/2020).

[45] J Matthew Watson et al. “Germline replications and somatic mutation accumulation are independent of vegetative life span in Arabidopsis.” In: *Proceedings of the National Academy of Sciences of the United States of America* 113.43 (Oct. 2016), pp. 12226–12231. DOI: 10.1073/pnas.1609686113. URL: http://dx.doi.org/10.1073/pnas.1609686113 (visited on 01/23/2022).

[46] Matthew T Weirauch et al. “Determination and inference of eukaryotic transcription factor sequence specificity.” In: *Cell* 158.6 (Sept. 2014), pp. 1431–1443. DOI: 10.1016/j.cell.2014.08.009. URL: http://dx.doi.org/10.1016/j.cell.2014.08.009 (visited on 09/12/2014).

[47] Yimeng Yin et al. “Impact of cytosine methylation on DNA binding specificities of human transcription factors.” In: *Science* 356.6337 (May 2017). ISSN: 0036-8075. DOI: 10.1126/science.aaj2239. URL: http://www.sciencemag.org/lookup/doi/10.1126/science.aaj2239 (visited on 04/12/2020).

[48] David A Zacharias et al. "Partitioning of lipid-modified monomeric GFPs into membrane microdomains of live cells." In: *Science* 296.5569 (May 2002), pp. 913–916. ISSN: 1095-9203. DOI: 10.1126/science.1068539. URL: http://dx.doi.org/10.1126/science.1068539 (visited on 08/13/2020).

[49] Cong Zhu et al. "High-resolution DNA-binding specificity analysis of yeast transcription factors." In: *Genome Research* 19.4 (Apr. 2009), pp. 556–566. DOI: 10.1101/gr.090233.108. URL: http://dx.doi.org/10.1101/gr.090233.108 (visited on 02/16/2021).
